# Diverse chemotypes drive biased signaling by cannabinoid receptors

**DOI:** 10.1101/2020.11.09.375162

**Authors:** Tamara Miljuš, Franziska M. Heydenreich, Thais Gazzi, Atsushi Kimbara, Mark Rogers-Evans, Matthias Nettekoven, Elisabeth Zirwes, Anja Osterwald, Arne C. Rufer, Christoph Ullmer, Wolfgang Guba, Christian Le Gouill, Jürgen Fingerle, Marc Nazaré, Uwe Grether, Michel Bouvier, Dmitry B. Veprintsev

## Abstract

Cannabinoid CB1 and CB2 receptors are members of the G protein-coupled receptor family, which is the largest class of membrane proteins in the human genome. As part of the endocannabinoid system, they have many regulatory functions in the human body. Their malfunction therefore triggers a diverse set of undesired conditions, such as pain, neuropathy, nephropathy, pruritus, osteoporosis, cachexia and Alzheimer’s disease. Although drugs targeting the system exist, the molecular and functional mechanisms involved are still poorly understood, preventing the development of better therapeutics with fewer undesired effects. One path toward the development of better and safer medicines targeting cannabinoid receptors relies on the ability of some compounds to activate a subset of pathways engaged by the receptor while sparing or even inhibiting the others, a phenomenon known as biased signaling. To take advantage of this phenomenon for drug development, a better profiling of the pathways engaged by the receptors is required. Using a BRET-based signaling detection platform, we systematically analyzed the primary signaling cascades activated by CB1 and CB2 receptors, including 9 G protein and 2 β-arrestin subtypes. Given that biased signaling is driven by ligand-specific distinct active conformations of the receptor, establishing a link between the signaling profiles elicited by different drugs and their chemotypes may help designing compounds that selectively activate beneficial pathways while avoiding those leading to undesired effects. We screened a selection of 35 structurally diverse ligands, including endocannabinoids, phytocannabinoids and synthetic compounds structurally similar or significantly different from natural cannabinoids. Our data show that biased signaling is a prominent feature of the cannabinoid receptor system and that, as predicted, ligands with different chemotypes have distinct signaling profiles. The study therefore allows for better understanding of cannabinoid receptors signaling and provides the information about tool compounds that can now be used to link signaling pathways to biological outcomes, aiding the design of improved therapeutics.

## Introduction

G protein-coupled receptors (GPCRs) transduce external signals to intracellular responses upon activation by ligands such as small molecules, hormones, neurotransmitters, chemokines, and photons^1^. Ligand binding on the extracellular side of the transmembrane receptor results in conformational rearrangements in the transmembrane domain^2^, leading to the recruitment and activation of G proteins^3,4^, phosphorylation by G protein-coupled receptor kinases (GRKs), interaction with arrestins and eventually receptor desensitization and internalization, recycling or degradation^5,6^. The type of G protein activated by the receptor determines which intracellular signaling cascades are triggered, regulating downstream physiological responses. While some GPCRs activate only one or a few G proteins, others interact with several members of multiple G protein subfamilies, as well as multiple β-arrestin subtypes^7–9^. Once engaged by the receptors, the β-arrestins can also lead to intracellular responses^10^.

The human endocannabinoid system consists of cannabinoid CB1 and CB2 receptors, along with additional receptors, endocannabinoids and enzymes involved in their transport and metabolism^11^. CB2 regulates immune responses and inflammatory pathways. While being expressed mostly in cells of the immune system with the highest expression in B-lymphocytes, it is also found in the peripheral nervous system^12^. In contrast, CB1 is one of the most abundant GPCRs in the brain^13^, where it regulates neurotransmitter release from presynaptic neurons and is responsible for synaptic plasticity and psychoactive responses associated with marijuana consumption^14^.

The endocannabinoid system is involved in neuromodulatory activity, cardiovascular tone, energy metabolism, bone formation, immunity, and reproduction, whereas its malfunction triggers various inflammatory conditions resulting in atherosclerosis^15^, neuropathy^16^, nephropathy, pruritus, osteoporosis^17^, Alzheimer’s disease^18^, autoimmune diseases^19^ and cancer^20^. It is reported that both cannabinoid receptors and endogenous ligands are upregulated in tumor tissue compared with non-tumor tissue, where CB2 has a strong anti-inflammatory effect and plays an important role in cell homeostasis^21^. Therefore, the spectrum of physiological activities makes the endocannabinoid system a promising target for treatment of various diseases. Cannabinoid receptors in spinal chord are targeted for neuropathic pain^22^, while cannabinoid inverse agonists are used as appetite suppressors to treat obesity^23^.

Both receptors were reported to couple mainly to G_i/o_ proteins, modulating intracellular signaling pathways, such as inhibition of cAMP-production, activation of extracellular receptor kinases (ERKs) and G protein-coupled inward rectifying K^+^-channels (GIRKs), and to recruit and activate β-arrestins^24–27^. In addition, there is evidence for activation of members of other G protein families under certain conditions^28–30^, receptor dimerisation^31^, internalized^26,32^, and recycling^30^.

Despite the fact that phytocannabinoids have been in use for millennia, the molecular basis of their pleiotropic actions is still not well understood. A major hurdle for effective treatment with natural products is the lack of selectivity of known natural cannabinoids and endocannabinoids. It belongs to the peculiarities of natural cannabinoid receptor ligands, which are mainly agonists, that they exist in high numbers as endocannabinoids, as well as phytocannabinoids, while none of them seems to be specific for one single cannabinoid receptor. Due to the high homology between the ligand binding domains, they activate both CB1 and CB2 as well as some additional receptors, such as GPR55, TRPV1 channels and 5-HT_3_ receptors. The pleiotropy of the cannabinoid system extends to their signaling cascade as it has been reported that CB1 and CB2 can engage members of different G protein families and that some ligands seem to show biased activity toward these different effectors^30,33–40^. The structural diversity of naturally occurring cannabinoids thus makes them an ideal playground to unravel their specific signaling pathways and to search for artificial ligands with specific properties, both in receptor activation/inactivation and biases in signaling. As positive effects are likely achieved exclusively through the regulation of one of the receptors and may even be associated with a specific signaling pathway, this approach may lead to better pharmacological understanding and therapy. For instance, CB2 selective compounds would allow targeting of CB2-mediated pathologies without the undesired psychoactive effects mainly resulting form the co-activation of the neuronally expressed CB1 receptors. Achieving such selectivity is a major hurdle for the development of CB2-targeting drugs. Similarly, a better mapping of the signaling pathways that can be triggered by different subclass of cannabinoid ligands could help linking specific pathways to beneficial or adverse biological outcomes.

The ability of certain ligands to preferentially activate subsets of the signaling pathways over others engaged by a given receptor, which is know as biased signaling or functional selectivity^9,41,42^, is pervasive within the family of GPCRs and generated hope for the development of better and safer drugs^43^. Yet, the lack of tool compounds allowing for linking specific pathways to desired therapeutic responses, combined with the lack of knowledge concerning the nature of the ligand–receptor interactions that drive biases, has hampered the development of functionally selective drugs demonstrating clinical advantages. Given that signaling bias is believed to result from ligand-specific distinct active conformations of the receptor^44–50^, profiling the signaling of ligands with distinct chemotypes should inform us on the relationship between chemical structures and receptor responses, in addition to provide tool compounds to directly test the impact of specific biases on biological functions. Here we assessed the CB1/CB2 selectivity and signaling profile of structurally diverse ligands, including endocannabinoids, phytocannabinoids and synthetic compounds structurally similar or significantly different from natural cannabinoids.

## Results

### Selecting structurally diverse library of CB ligands

The 35 cannabinoid receptor ligands we selected for this study originate from several structurally diverse chemotypes (Table S1). SR144528, AM630, AM1241, WIN55912-2 and CP55940 are well characterized synthetic cannabinoid ligands^38,51^. SR144528 and AM630 are CB2 inverse agonists, AM1241 is a CB2 agonist, whereas WIN55912-2 and CP55940 are mixed CB1 and CB2 agonists. Δ^9^-THC (henceforth referred to as THC), cannabinol, HU210, nabilone, HU308 and JWH133 are phytocannabinoid-derived CB modulators^38,51,52^. THC, cannabinol, HU210 and nabilone have been described as mixed CB1/CB2 agonists, while HU308 and JWH133 are CB2-selective agonists. The endogenous cannabinoid ligands 2-arachidonoyl glycerol (2-AG) and anandamide, both non-selective activators of CB1 and CB2^38,51^, were also included in the screening library. A more recently discovered CB2 selective triazolopyrimdine-based scaffold is also represented in the screening set by RO6435559, RO6844395^53^, RO6869094^54^, RO6871487^54^ and RO6878558^54,55^. Common characteristics of these ligands are that they carry three exit vectors at positions 3, 5 and 7 of the triazolopyrimidine core. While the position 3 residue is kept constant using a (2-chlorophenyl)methyl moiety the other two positions are modified. Further structural diversity in the screening set was added by including a pyridine/pyrazine derived scaffold which carries generally 3 substituents in positions 2, 5 and 6 of the heteroaromatic core (RO6843766^56^, RO6844112, RO5135445, RO6892033, RO7032019^57^, FMP7234690, FMP7234691, FMP7234694, FMP7234698 and FMP7234699). For increasing the chance of engaging different signaling pathways, molecules with variations at all three exit vectors have been included. At position 2 the common denominator is a carboxylic acid amide group or as in case of pyridine RO6892033, a respective amide isoester. FMP7234690, FMP7234691, FMP7234694, FMP7234698 and FMP7234699 are decorated with different linker-nitrobenzofurazan elements in position 2 which allows for investigating the influence of molecules with different sizes on signaling bias. To complete the structurally diverse set of probes, a 2,4,5-trisubstituted pyridine derived class of novel CB2 ligands possessing a distinct and unique structure activity relationship as compared to the 2,5,6-trisubstituted pyridines/pyrazines was added (RO6850007^58^, RO6853457^58^ and RO6926274^59^). The commonality of these ligands is a cyclopropyl ring in position 5 whereas the residues in positions 2 and 4 are different. RO6926274 carries on top a chlorine atom in position 6.

### *In vitro* pharmacology data

Prior to studying their biased signaling fingerprint, the ligand test set has been subjected to in depth *in vitro* pharmacological profiling, including binding data, maximal ligand-induced response (E_max_) and half-maximal response concentration (as pEC50) in cAMP production and β-arrestin recruitment assays for both human CB1 and CB2 receptors (Table S2). These data were complemented by the respective mouse CB2 values in order to get an idea on potential species differences which are highly important for translating results from mouse *in vivo* pharmacology studies to humans. Most ligands have a very high affinity for the human CB2 receptor as assessed by competition binding studies using [^3^H]-CP55940 as a radioligand tracer. HU210 showed the highest affinity (hCB2 pK_i_ = 9.78), closely followed by the structurally completely unrelated pyridine RO6853973 (hCB2 pK_i_ = 9.45). The endogenous ligand anandamide exhibits the lowest affinity (hCB2 pK_i_ = 6.91) for the human CB2 among the entire ligand set. Triazolopyrimdine RO6878558 is the second weakest human CB2 binder with a pK_i_ of 6.94. Generally, pK_i_ values for human and mouse CB2 are within the same range (Δ(pK_i_(hCB2) - pK_i_(mCB2)) = ± 0.5). A few ligands however show a significantly higher affinity for mouse than for human CB2 (cf. SR144528: pK_i_(hCB2) = 7.88 vs. pK_i_(mCB2) = 10.7; and RO6850007: pK_i_(hCB2) = 7.65 vs. pK_i_(mCB2) = 9.21). Interestingly, several CB2 binders with an opposite behavior are included in the compound set, having a higher affinity for human than for mouse CB2 receptor (cf. WIN55912-2: pK_i_(hCB2) = 8.57 vs. pK_i_(mCB2) = 7.28; and RO6844395: pK_i_(hCB2) = 7.67 vs. pK_i_(mCB2) = 6.87). With respect to the closely related human CB1, the library covers a broad range of affinities ranging from molecules with basically no interaction with human CB1 (cf. (*S*)-AM1241, HU308 and RO7032019 with pK_i_(hCB1) values < 5) to high affinity CB1 ligands such as THC (pK_i_(hCB1) = 8.48), HU210 (pK_i_(hCB1) = 9.55) and trisubstituted pyridine RO6853973 (pK_i_(hCB1) = 8.21). Some molecules show a preference for binding the human CB1 over the human CB2 receptor (cf. CP55940 10^pKi(hCB2)-pKi(hCB1)^ = 0.2). The endocannabinoid 2-AG is a balanced human CB1 and CB2 binder (10^pKi(hCB2)-pKi(hCB1)^ = 1). However, the majority of tested ligands show a preference for human CB2. Highest CB2 binding selectivity is found for triazolopyrimidine RO6871487 (10^pKi(hCB2)-pKi(hCB1)^ = 436). Pyridine RO6843766 also exhibits a strong preference for human CB2 (10^pKi(hCB2)-pKi(hCB1)^ = 416), while HU308 is the most CB2 selective cannabinoid-derived ligand (10^pKi(hCB2)-pKi(hCB1)^ = 278). The CB2 ligand library was furthermore profiled in cellular systems for functional read-outs, such as modulation of cAMP levels in CB2-overexpressing Chinese hamster ovary (CHO) cell lines. Several molecules from different chemotypes exhibited subnanomolar potencies on human CB2 (cf. HU210: cAMP pEC50(hCB2) = 9.45 and CP55940: cAMP pEC50(hCB2) = 10.33). The weakest tested ligand was again anandamide yielding a cAMP pEC50(hCB2) value of 5.93.

### Set of ligands with a range of activities

To enhance the chance for getting a maximally diverse biased signaling fingerprint, the compound library was chosen in a way that it included agonists, partial agonists, as well as antagonists and inverse agonists (Table S2). For example, HU308 exhibits a cAMP E_max_(hCB2) value of 98% and is therefore considered to be a full agonist on human CB2 receptor (the effect of agonists is normalized to the effect of 10 μM CP55940 used as the reference full agonist). In addition, pyridine-derived ligands were able to achieve full efficacy (cf. FMP7234691: cAMP E_max_(hCB2) = 100%), as well as the endocannabinoids 2-AG and anandamide (cAMP E_max_(hCB2) = 94% and 87%, respectively). In contrast, the efficacy of triazolopyrimidine RO6871487 reaches only 52% of the maximal effect and is therefore classified as partial agonist. Tetrasubstituted pyridine RO6926274 is an even weaker partial agonist, with a 38% maximal effect value. The strongest inverse agonism was found for AM630 (cAMP E_max_(hCB2) = −152%) and SR144528 (cAMP E_max_(hCB2) = −151%). Pyridine RO7032019 was also classified as an inverse agonist (cAMP E_max_(hCB2) = −92%). Potency for cAMP production by the human CB1 was low for many of the ligands (cf. JWH133: cAMP pEC50(hCB1) < 5; and RO6850007: cAMP pEC50(hCB1) < 5). In contrast, HU210, CP55940, and RO6853973 exhibit single digit nanomolar or even subnanomolar EC50 values (cAMP pEC50(hCB1) = 9.58, 9.73 and 8.12, respectively). All tested ligands behave either as full CB1 agonists – with pyridines FMP7234691 and FMP7234690 showing even higher E_max_ values (cAMP E_max_(hCB1) = 123% and 152%, respectively) than CP55940 at 10 μM – or partial agonists at the highest tested concentration of 10 μM. No antagonism nor inverse agonism for human CB1 was found.

### Receptor subtype variations of ligand activity

With the exception of HU210 (10^cAMP pEC50(hCB2) - cAMP pEC50(hCB1)^ = 1), all molecules show at least some functional selectivity when tested at the human CB1 and CB2 receptors. cAMP selectivity ratios, defined by 10^cAMP pEC50(hCB2) - cAMP pEC50(hCB1)^, reached values up to > 3388 (HU308). Functional selectivity was generally higher than binding selectivity (for exception cf. RO6843766: 10^cAMP^ pEC50(hCB2) - cAMP pEC50(hCB1) = 168 vs. 10^(pKi(hCB2)-pKi(hCB1)^ = 416). Most significant difference between cAMP and binding selectivity was observed for AM630 and WIN55912-2 (10^cAMP pEC50(hCB2) - cAMP pEC50(hCB1)^ / 10^pKi(hCB2)-pKi(hCB1)^ > 45 and 44, respectively). Potencies in the mouse cAMP assay range from highly active molecules (cf. CP55940: cAMP pEC50(mCB2) = 10.17; and RO6892033: cAMP pEC50(mCB2) = 8.39) to inactive ligands (cf. RO7032019: cAMP pEC50(mCB2) < 5). Generally, pEC50 data for human and mouse CB2 receptors are within the same order of magnitude (pEC50(hCB2)-pEC50(mCB2) = 0 ± 1), especially when the molecules behave as strong partial or full agonists for the human receptor (cAMP E_max_(hCB2) > 70%). Overall, efficacy for mouse CB2 tends to be lower than efficacy for human CB2. In particular, partial agonists for human CB2 are associated with lower potency for mouse CB2 (cf. RO6926274: cAMP pEC50(hCB2) = 8.15 vs. cAMP pEC50(mCB2) < 5). Such partial agonists for human CB2 tend to be weak partial agonists or neutral antagonists for mouse CB2 (cf. RO6853973: cAMP E_max_(hCB2) = 45 vs. cAMP E_max_(mCB2) = 23; and RO6926274: cAMP E_max_(hCB2) = 38 vs. cAMP E_max_(mCB2) = 7). In several cases even a switch from agonism at human CB2 toward inverse agonism at mouse CB2 has been observed (cf. (*R*)-AM1241: cAMP E_max_(hCB2) = 65 vs. cAMP E_max_(mCB2) = −22; and RO6850007: cAMP E_max_(hCB2) = 63 vs. cAMP E_max_(mCB2) = −48). Such changes can be of utmost importance when it comes to the interpretation of cAMP-dependent efficacy mouse data *in vivo*. Interestingly, for the two enantiomers of AM1241, only (*R*)-AM1241 showed a switch from agonism at human to inverse agonism at mouse CB2, while (*S*)-AM1241 showed partial agonism at human and mouse CB2 (cAMP E_max_(hCB2) = 78 vs. cAMP E_max_(mCB2) = 35). Apart from cAMP read-outs, functional potency, efficacy and selectivity data in a β-arrestin recruitment assay were generated for the library of ligands. β-arrestin pEC50 values at human CB2 range from 8.39 (CP55940) to < 5 (cf. (*rac*)-AM1241). Potency for most molecules in hCB2 β-arrestin recruitment assay is lower than in the hCB2 cAMP assay (cf. WIN55912-2: cAMP pEC50(hCB2) = 9.50 vs. β-arrestin pEC50(hCB2) = 7.93). With regard to efficacy, once again the whole spectrum ranging from inverse agonism (cf. AM630: β-arrestin E_max_(hCB2) = −124%), passing by partial agonism (cf. (*R*)-AM1241: β-arrestin E_max_(hCB2) = 29%), toward full agonism (cf. CP55940: β-arrestin E_max_(hCB2) = 80%) is covered by the ligand set. Between the two endocannabinoids, 2-AG behaves as a stronger partial agonist than anandamide (cf. 2-AG: β-arrestin E_max_(hCB2) = 38%; 2-AG: β-arrestin E_max_(hCB2) = 80%). Based on the available data, inverse agonism in the human CB2 cAMP assay translates into inverse agonism in the human CB2 β-arrestin recruitment assay. A similar trend was observed for agonism. However, full agonists in the human CB2 cAMP assay possess, in most cases, a lower E_max_ value in the human CB2 β-arrestin recruitment assay (cf. (*S*)-AM1241: cAMP E_max_(hCB2) = 78% vs. β-arrestin E_max_(hCB2) = 29%). In the human CB1 β-arrestin recruitment assay, CP55940 was the most potent (β-arrestin pEC50(hCB1) = 7.96), while several ligands possess pEC_50_ values below 5 (cf. (*S*)-AM1241 and HU308). CP55940 represented the most efficacious agonist (β-arrestin E_max_(hCB1) = 94%) of the test set, and JWH133 was a partial agonist (β-arrestin E_max_(hCB1) = 40%). Although SR144528 and AM630 did not have an effect on CB1-mediated cAMP signaling, they showed inverse agonism in β-arrestin recruitment assay (SR144528 β-arrestin E_max_(hCB1) = −82%; AM630: β-arrestin E_max_(hCB1) = −120%) The selectivity of CB2 vs. CB1 β-arrestin recruitment ranges from poorly selective CP55940 (10^β^-^arrestin pEC50(hCB2) - β^-^arrestin pEC50(hCB1)^ = 3), to highly selective SR144528 (10^β^-^arrestin pEC50(hCB2) - β^-^arrestin pEC50(hCB1)^ = 316). Representative mouse CB2 β-arrestin recruitment data range from molecules with pEC50 values of 8.02 (CP55940) to values below 4.5 (cf. SR144528). For all tested molecules, potency at the mouse CB2 β-arrestin recruitment assay was lower than at the human CB2, as well as at mouse CB2 cAMP assay. Most efficacious agonist was CP55940 (β-arrestin E_max_(mCB2) = 96). In summary, the ligand test set provides maximal diversity, not only with regard to the chemical structure of the molecules, but also to its *in vitro* pharmacology profile. Ligands with subnanomolar affinities for both human and mouse CB2 receptors are included, as well as molecules with weaker affinity. Binders with a slight preference for CB1, and highly selective CB2 ligands are part of the library. Regarding functional potency, a broad range of pEC50 values, both for cAMP and β-arrestin recruitment, is covered. Importantly, the complete efficacy space, ranging from inverse agonism to full agonism, was displayed for both read-outs. Therefore, the ligand test set is ideally suited for identifying different biases in signaling fingerprints.

### Early absorption, distribution, metabolism and excretion (ADME) profile

To assess the diversity of the ligand test set with regard to its physicochemical properties and as a preparation for potential *in vivo* follow-up studies for exploring whether differences in biased signaling result in different pharmacodynamic effects, early adsorption, distribution, metabolism and excretion (ADME) data were generated (Table S3). Exposing CB2 in the respective target tissue after systemic routes of administration is a pre-requisite for carrying out such *in vivo* studies. To achieve high exposures, especially favourable lipophilicity, membrane permeation and microsomal stability values are of high relevance and were, therefore, key elements of this assessment. The ligands are highly diverse in their molecular weight (MW), covering a range from 310.4 (cf. cannabinol) to 841.0 g/mol for the CB2 ligand FMP7234698 which, besides a CB2 recognition element, also contains a spacer and a fluorescent nitrobenzofurazan moiety, while optimal MW for achieving high bioavailabilities is below 500 g/mol. In addition, the ligand set is extremely diverse in their polar surface area (PSA). The almost pure hydrocarbon molecule JWH133 exhibits the lowest PSA value of 7.9 Å^2^, while polyethylenglycol-derived FMP7234698 possesses a very high PSA of 176.9 Å^2^.

The maximum number of hydrogen bond donors (HBDs) within the library is two (cf. anandamide and RO6853457). Several of the selected molecules do not contain a hydrogen bond donating element at all (cf. AM630 and RO6435559). In case where barriers, such as the blood brain barrier or the blood retina barrier, need to be crossed, molecules with a PSA < 75 Å^2^ and a low number of HBDs will be preferred^60,61^.

Since the endogenous cannabinoid receptor ligands, 2-AG and anandamide, are fatty acid derivatives, their lipophilicity is high. This is reflected by their calculated octanol/water partition coefficients (K_ow_ clogP) of 6.7 and 6.3, respectively. The corresponding values of the synthetic ligands SR144528 (9.2), HU308 (9.0) and especially pyridine FMP7234699 (11.3), which contains a hydrocarbon spacer between nitrobenzofurazan dye and CB2 recognition element, are higher. Interestingly, CB2 tolerates a tremendous range of polarities, since the highly polar pyridine bis-amide RO7032019 (K_ow_ clogP = 1.5) is, like FMP7234699, a tight binder. Experimentally generated octanol/water distribution coefficients (log*D*) are often lower than its corresponding calculated K_ow_ clogP values, ranging from 2.5 (RO7032019) to 4.29 (HU308). Because charges affect measured log*D* values, but are not reflected in predicted K_ow_ clogP numbers, differences between log*D* and K_ow_ clogP values are generally higher for molecules that are charged under physiological conditions (pH 7.4) e.g. as found for (*S*)-AM1241 (log*D* = 3.64 vs. K_ow_ clogP = 5.7) carrying a basic piperidine moiety (pK_a_ = 7.8). Most of the molecules in our ligand set are either uncharged or carry a basic center, whereas only cannabinol, CP55940 and pyridine RO6850007 contain weakly acidic phenol or amide moieties. Compound solubility was assessed using lyophilisation solubility assay (LYSA)^62^. Kinetic solubility in aqueous buffer is the highest for pyridine derived bis-amide RO6853457 (LYSA = 92 μg/mL). For the endocannabinoids and the cannabinoid-derived ligands LYSA values are generally poor (cf. 2-AG: LYSA = < 0.5 μg/mL; or JWH133: LYSA = < 0.4 μg/mL). The molecules’ potential for passive membrane permeation was investigated using the parallel artificial membrane permeability assay (PAMPA) model^63^. Highest permeation coefficients *P*_eff_ were found for the two highly drug-like pyridines RO7032019 (*P*_eff_ = 6.4 × 10^−6^ cm/s) and RO6853457 (*P*_eff_ = 3.9 × 10^−6^ cm/s), with their efficient passive permeation also reflected by a high propensity for permeating to the acceptor compartment (%Acc.) value of 12% for both molecules. Anandamide was also able to passively permeate membranes (*P*_eff_ = 0.26 × 10^−6^ cm/s). In contrast, molecules with higher K_ow_ clogP, such as SR144528 and CP55940, had a high propensity for sticking to the membrane (%Mem. = 54% and 47%, respectively) rather than permeating to the acceptor compartment (%Acc. = 0% for both molecules), leading to a permeation coefficient value of zero for both molecules.

Clearance rates of the CB2 ligands were determined in co-incubation experiments with liver microsomes, providing first predictions of doses needed for reaching sufficient target tissue exposure for future *in vivo* follow-up studies (low to medium clearances (CL) translate into low doses). In general, clearance values in human are lower than in rodent liver microsomes. Highest human microsomal CL was obtained for (*R*)-AM1241 (304 μL/min/kg). In mouse microsomes WIN55912-2 and CP55940 were least stable (CL = 745 μL/min/kg for both molecules). The most stable molecule in human, as well as in mouse microsomes, is pyridine RO7032019, which originates from a drug-discovery program (CL = 10 μL/min/kg in both species). CB2 inverse agonist AM630 and agonists HU308 and JWH133 have been exploited in various mouse disease models. While mouse hepatocyte CL values are high for AM630 (101 μL/min/million cells), low clearances were determined for HU308 (5 μL/min/million cells) and JWH133 (8 μL/min/million cells). In human hepatocytes all three molecules exhibit low to medium clearances. In case of AM630 and JWH133, clearance in hepatocytes is lower than in microsomes (AM630: hum. mic. CL = 82 μL/min/kg vs. hum. hep. CL = 18 μL/min/million cells; and JWH133: mouse mic. CL = 63 μL/min/kg vs. hum. hep. CL = 10 μL/min/million cells).

This observation can likely be attributed to the tight protein binding being reflected by the low free fractions found in both the human and the mouse plasma protein binding assays (AM630: free fraction (%) human/mouse = <0.1/0.4; and JWH133: free fraction (%) human/mouse = <0.1/<0.1). Free fraction usually increases with higher molecule polarity, and this trend can be observed in the cannabinoid receptor ligand test set, where for more polar (*S*)-AM1241 (K_ow_ clogP = 5.7, log*D* = 3.64) a free fraction of 0.5% in human and of 1.2% in mouse plasma protein binding assay was determined, whereas less polar JWH133 (K_ow_ clogP = 8.5, log*D* > 3) exhibits free fractions below 0.1% in both species. Neither (*S*)-AM1241 nor JWH133 are substrates for the mouse P-glycoprotein (P-gp) transporter ((*S*)-AM1241: (P-gp) mediated efflux ratio mouse = 1.4; JWH133: (P-gp) mediated efflux ratio mouse = 1.7), making them capable for crossing biological barriers such as the blood brain barrier, something that might become important when signaling bias is investigated in mouse models of neuroinflammatory diseases. In contrary, SR144528 and HU308 are interacting with the P-gp transporter (SR144528: (P-gp) mediated efflux ratio mouse = 5.6; HU308: (P-gp) mediated efflux ratio mouse = 7.3) and are, therefore, less suitable for being applied in brain disease models as they would require higher doses to compensate for the efflux. Overall, the biased signaling fingerprint ligand test set covers an extremely broad range of physicochemical, as well as early ADME properties.

### Signaling fingerprint of CB1 and CB2

Using a BRET-based signaling platform^64–66^, we assessed the signaling repertoire of both CB1 and CB2 toward 13 G protein and the 2 β-arrestin subtypes in response to the agonists CP55940 for CB1 and JWH133 for CB2. G_s_ protein activation was assessed by monitoring decrease of BRET signal upon dissociation of Gα_s_ and Gβ_1_γ2 subunits^64,67^, while activation of other G protein subtypes was measured as an increase in BRET signal upon recruitment of a subtype-specific signaling effector to the plasma membrane^66^. Similarly, recruitment of β-arrestins to the receptors was monitored as β-arrestin translocation from cytosol to the plasma membrane^65^. Both cannabinoid receptors couple primarily to the G_i/o_ family of G proteins and recruit β-arrestin2 more strongly than β-arrestin1 upon activation (Figure 1; Table S7). In addition, CB1 activates the atypical G_i/o_ family member G_z_ and well as G_12/13_ and G_15_. No activation of the G_s_ or any members of G_q_ family apart from G_15_ was observed. Both CB1 and CB2 only negligibly promote β-arrestin trafficking to endosomes upon receptor internalization as assessed by enhanced bystander BRET^65^, in contrast with the strong translocation promoted by the vasopressin V2 receptor (Figure 2).

**Figure 1.**
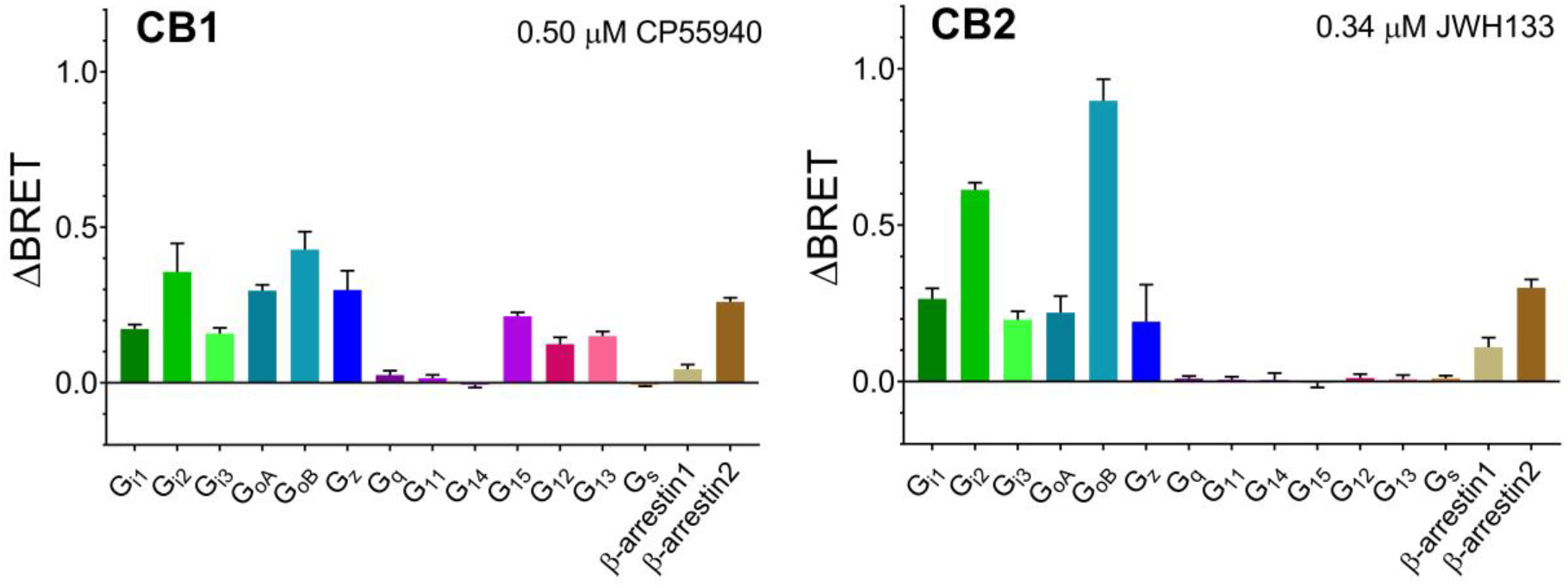
G protein activation and β-arrestin recruitment by CB1 and CB2: maximal ligand-induced response (E_max_) in ΔBRET signal upon receptor stimulation with agonist (0.5 μM CP55940 for CB1, 0.34 μM JWH133 for CB2). Positive ΔBRET indicates activation of a specific pathway. For G protein activation and β-arrestin recruitment, relocation of signaling effector to the plasma membrane was monitored. For activation of G_s_ protein, dissociation of Gα subunit from Gβγ was monitored. Mean and standard error of mean are shown, data from Table S7. CB1 couples to G_i/o_ family, G_z_, G_15_ and G_12/13_. CB2 couples to G_i/o_ family of G proteins. Both receptors recruit β-arrestin2 more strongly than β-arrestin1.

**Figure 2.**
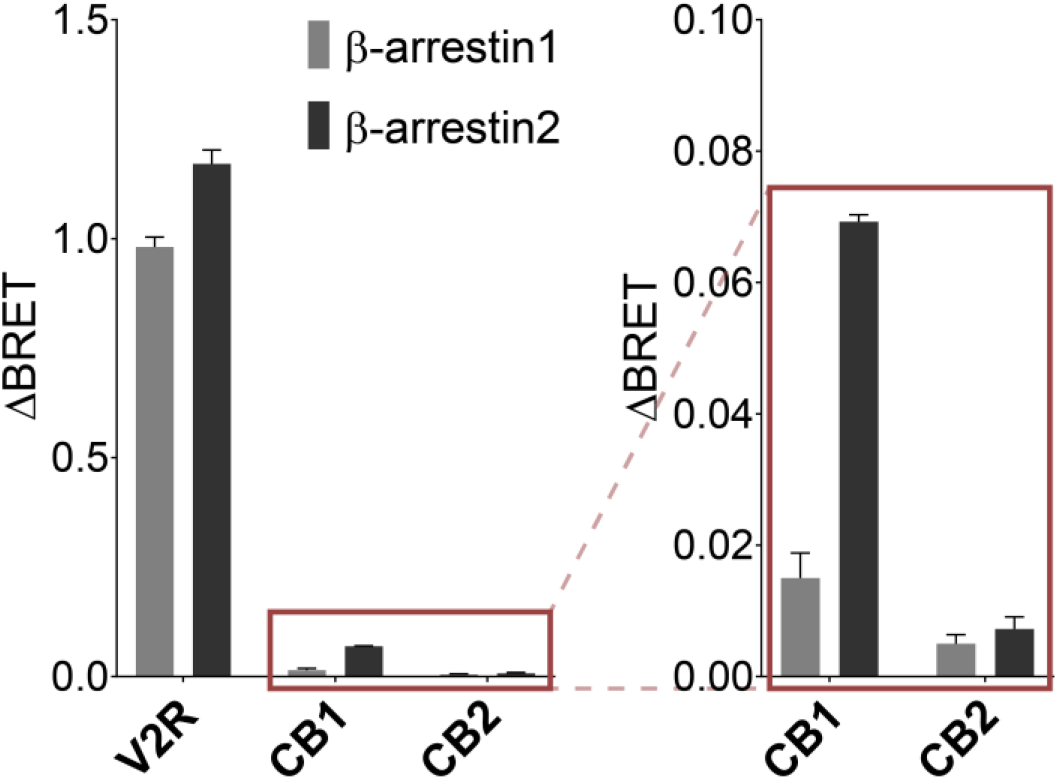
β-arrestin trafficking to the endosomes upon receptor internalization, monitored as relocation of β-arrestin to the endosomal membrane. Both cannabinoid receptors show negligible β-arrestin recruitment in endosomes when compared to vasopressin V2 receptor (V2R), a strong β-arrestin recruiter.

For each of the pathways activated by cannabinoid receptors, a concentration-dependent response was measured for a selection of ligands. While 35 ligands were tested for CB2-mediated signaling, only eight ligands, which were not CB2 selective, were tested for CB1-mediated signaling. Maximal ligand-induced response (E_max_) and half-maximal response concentration (as pEC50) were obtained for each ligand (Table S8 and S9). In order to allow comparison of each ligands’ effect on different signaling pathways, the E_max_ value for each ligand was normalized to the value of the reference ligand WIN55212-2. The same reference ligand was used to calculate bias factors using the operational model of bias agonism^68^ (Table S15 and S17).

### CB1 signaling

When considering the maximal ligand-promoted β-arrestin recruitment (E_max_), all compounds were more efficacious for β-arrestin2 than β-arrestin1, except for 2-AG where β-arrestin1 response is about 40% higher than that of β-arrestin2 (Figure 3). Interestingly, 2-AG is also the only compound that generated a larger response for the two β-arrestins than for the G proteins, indicating a clear preference for β-arrestin signaling. Regarding G protein activation, most compounds had similar relative efficacies (E_max_) toward the different subtypes (Figure 3, left). Notable exceptions includes cannabinol, which, in contrast to all other compounds, acts as an inverse agonist on G_15_ response. Other differences include the G_i1_ response to THC and cannabinol, which was generally low for the other compounds, but is among the most efficacious compared to the other G protein subtypes for these two ligands. Response of G_15_ and G_12_ was also generally lower, except for 2-AG and CP55940. When looking at potencies (pEC50 values, Figure 3, right), a general trend of higher pEC50 for the G_i/o_ family of G proteins is observed across the tested ligands, with a clear separation across 3 logarithmic units visible for 2-AG: G_i/o_ < G_12/13_ = G_15_ < β-arrestins. A wide range of potencies is also observed for cannabinol-induced signaling via CB1, although without clear separation between potencies for activation of distinct G protein subtypes (Figure 3, right).

**Figure 3.**
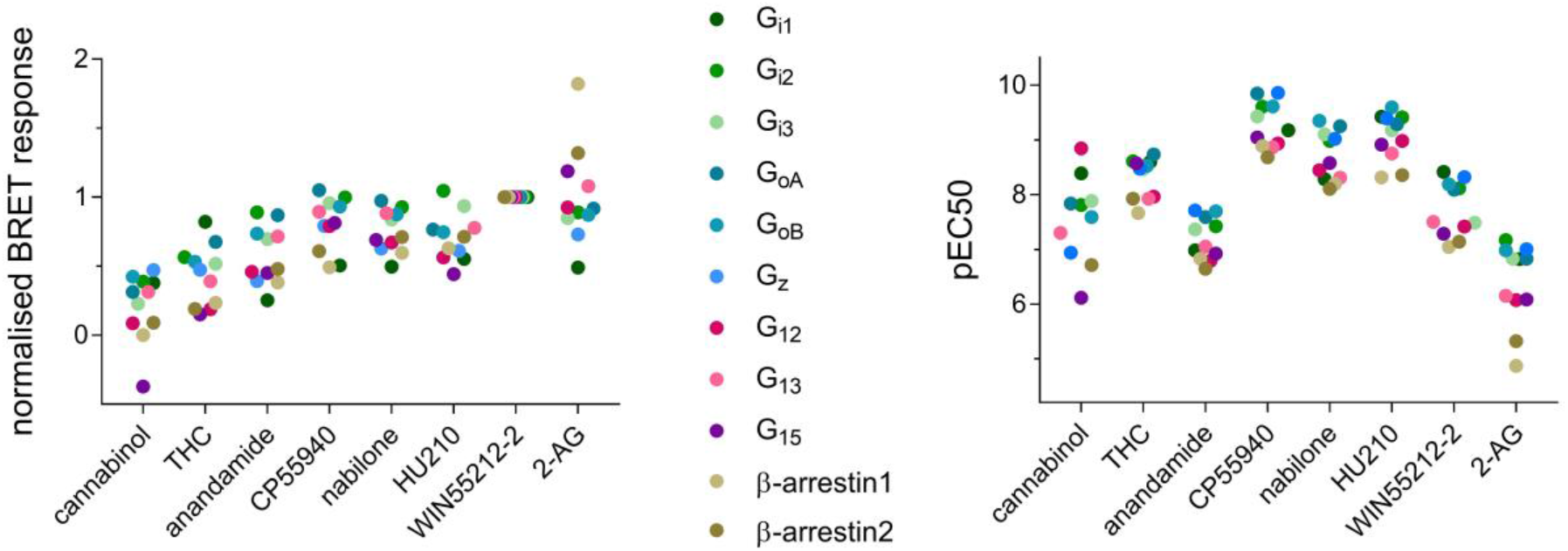
Characterization of ligand-induced CB1 signaling: dot-charts showing maximal relative ligand-induced response values (E_max_) normalized to WIN55212-2 (left) and negative logarithm of half-maximal response concentration (pEC50) for all the tested pathways. Only mean values of three independent experiments are shown, data from Table S8.

Based on efficacy only, 2-AG showed slight preference towards recruitment of β-arrestins than activation of G proteins, clearly visible from the obtained concentration-response curves (Figure 4). Thus, even when a maximal activation of G proteins is reached (1-10 μM), further increase in 2-AG concentration can lead to a change in signaling output due to more efficacious β-arrestin activation. In contrast, another endocannabinoid anandamide showed higher efficacy for activation of G_i2_, G_oB_ and G_13_ than other tested pathways (Figure 4).

**Figure 4.**
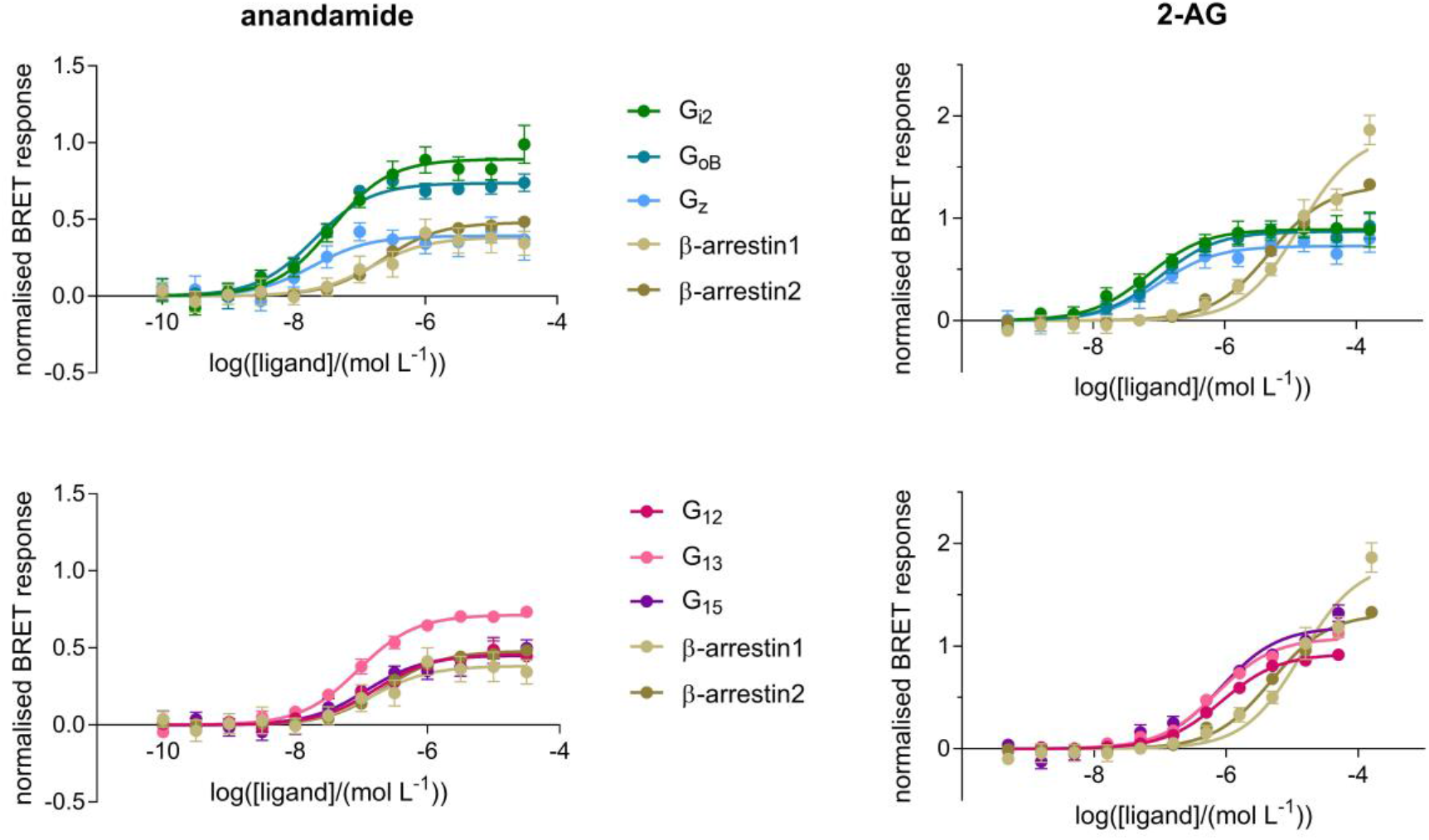
Concentration-dependent CB1-mediated activation of G proteins and β-arrestins by endocannabinoids anandamide (left) and 2-AG (right). Signal response is normalized to reference ligand WIN55,212-2. Mean and standard error of mean are shown, data from Table S10 and Table S11.

Based on bias factors calculated using WIN55212-2 as a reference, all the tested ligands are generally balanced (Table S15). CP55940, nabilone, and anandamide are slightly biased towards G_i2_ than G_i1_, and cannabinol preferentially activates β-arrestin1: β-arrestin1 > G_12_ > G_i3_ = G_i1_ = G_13_ = G_i2_ = G_oA_ = G_oB_ > β-arrestin2 > G_z_. Although modest differences in relative efficacies toward different G proteins were observed among the compounds, calculated bias factors vs. WIN55212-2 did not reach statistical significance (Table S15).

### CB2 signaling

In contrast to CB1 receptor signaling, larger differences across the signaling repertoire of the CB2 were observed for the ligands tested (Figure 5).

**Figure 5.**
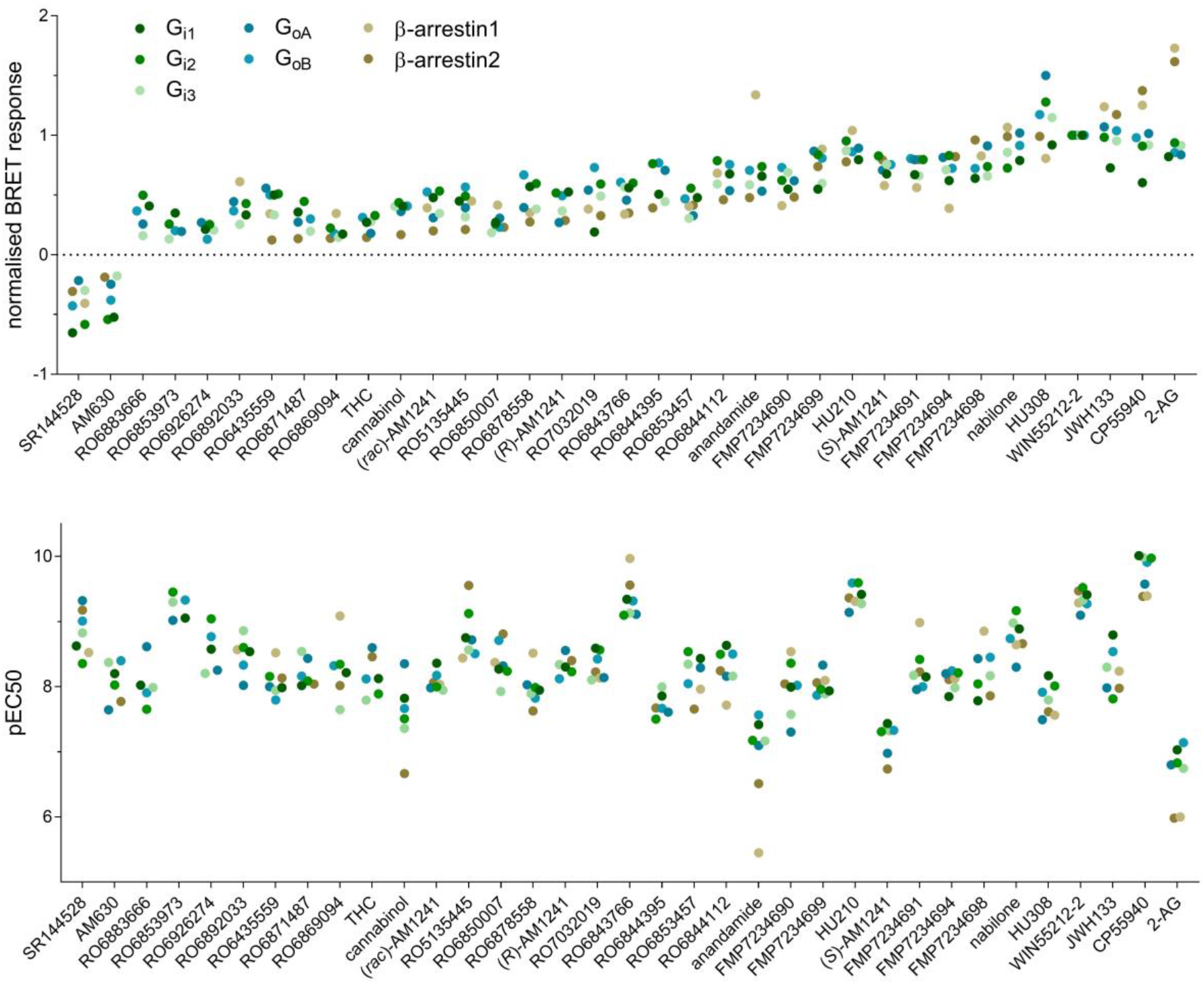
Characterization of ligand-induced CB2 signaling: dot-charts showing maximal relative ligand-induced response values (E_max_) normalized to WIN55212-2 (top) and negative logarithm of half-maximal response concentration (pEC50, bottom) for all the tested pathways. Mean values of three independent experiments are shown, data from Table S9.

Focusing on efficacies and potencies of compounds (Figure 5), most ligands activating all tested pathways. However, several ligands activated only G proteins, without detectable β-arrestin recruitment: RO6883666, RO6853973 and RO6926274. Regarding G protein activation, G_i1_ is sometimes recruited with lower efficacy than other G proteins (Figure 5). 2-AG, anandamide, RO6892033 and to a lower extent CP55940, HU210 and JWH133 evoke stronger response at β-arrestin than G proteins. While pyridine-derived RO6892033 does not recruit β-arrestin2, β-arrestin1 might be slightly preferred over G proteins (pEC50 similar, but more efficacious). In addition, RO6869094 showed preference towards β-arrestin1 than any other pathway, based on both E_max_ and pEC50.

When considering the maximal ligand-induced CB2-mediated β-arrestin1 and β-arrestin2 response (Figure 5), 2-AG produces similar E_max_ values for β-arrestin1 and β-arrestin2, whereas anandamide evokes a robust β-arrestin1, but only a minor β-arrestin2 response. Furthermore, endocannabinoid anandamide is less potent for β-arrestin1 than G proteins and especially β-arrestin2 (Figure 5, Table S9). The signaling preference of endocannabinoids 2-AG and anandamide is well illustrated in the concentration-response curves presented in Figure 6. Whereas 2-AG clearly showed preferences towards both β-arrestins vs. G proteins, anandamide showed preference only towards β-arrestin1 in comparison with other tested pathways (Figure 6).

**Figure 6.**
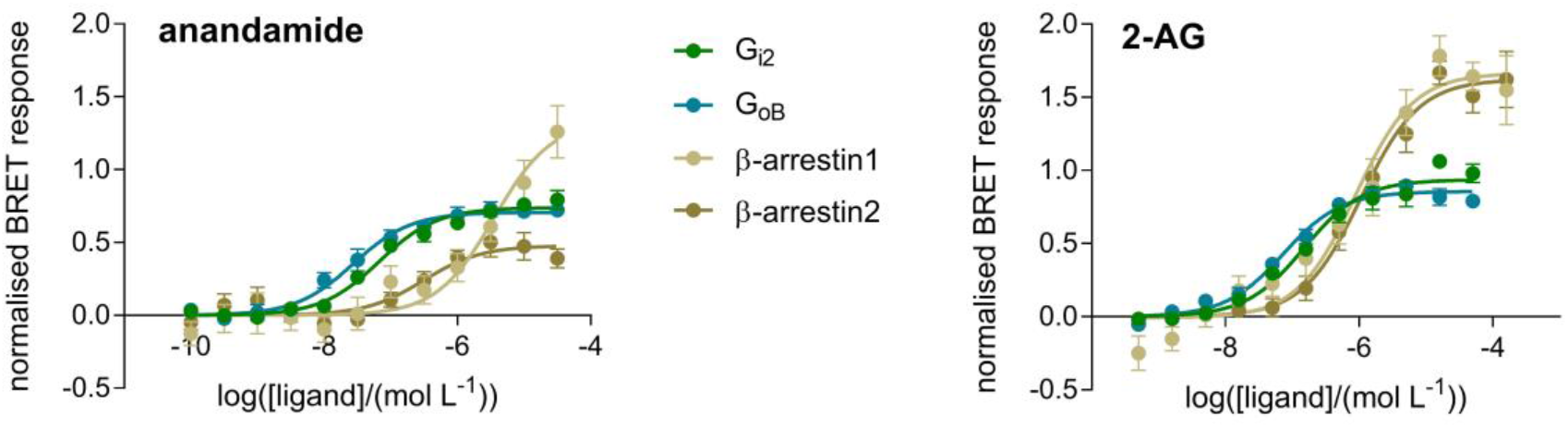
Concentration-dependent CB2-mediated activation of G proteins and β-arrestins by endocannabinoids anandamide (left) and 2-AG (right). Signal response is normalized to reference ligand WIN55,212-2. Mean and standard error of mean are shown, data from Table S12 and Table S13.

Interesting signaling profiles were observed for distinct enantiomers and racemic AM1241. While (*R*)-AM1241 recruits only β-arrestin2 (E_max_ = 29%), (*S*)-AM1241 recruits both β-arrestins with a minor preference towards β-arrestin2 (β-arrestin1 E_max_ = 58%, β-arrestin2 E_max_ = 79%). However, racemic AM1241 activates β-arrestin1 stronger than β-arrestin2 (β-arrestin1 E_max_ = 39%, β-arrestin2 E_max_ = 20%), with β-arrestin2 response being lower than for either of the enantiomers. This observation is in accordance with the findings of Soethoudt *et al.*^38^

As for CB1, relative effectivenesses (Table S16) and bias factors using WIN55212-2 as a reference (Table S17) were calculated for CB2. Three groups of ligands could be identified by hierarchical clustering of the calculated bias factors of the ligands (Figure S1). The first group have similar signaling properties as the reference ligand WIN55212-2, including HU210 and nabilone. The second group included ligands with signaling reduced more or less uniform across the observed signaling pathways (G_i1-3_, G_oA_, G_oB_, β-arrestin1 and β-arrestin2). This group included HU308 and JWH133, among other ligands. The third group that included THC, cannabinol, 2-AG and anandamide among others had a more pronounced biased signaling profile. All of these ligands had β-arrestin signaling affected more strongly compared to G protein signaling, effectively making them G protein biased. In this group, compounds RO6843766, RO6926274 and RO6853973 did not recruit arrestin, while RO7234687 was active only towards G_o_ family.

Several of the tested ligands show a bias towards β-arrestin1 over β-arrestin2 when compared to the reference ligand WIN55212-2: RO6871487 > RO6869094 ≈ RO6892033 > cannabinol ≈ RO6878558 ≈ FMP7234698 ≈ (*R*)-AM1241 ≈ RO6435559. While cannabinol, anandamide, RO6892033 and 2-AG are slightly biased towards G_oB_ over β-arrestin2, RO6883666 and cannabinol are biased towards G_oB_ over G_oA_. JWH133 is slightly biased towards G_i1_ vs. G_i2_ and towards G_oB_ vs. G_i2_, RO6883666 and cannabinol are biased towards G_oA_ over G_i2_, while none of the ligands show any bias between G_i2_ and G_i3_. RO6853457, cannabinol and RO6892033 are slightly biased towards G_i2_ over β-arrestin2, cannabinol and anandamide towards G_i2_ over β-arrestin1. A few ligands are biased towards β-arrestin1 vs. G_i2_: RO6871487 ≈ RO6853973 > RO6869094 ≈ FMP7234698 ≈ (*R*)-AM1241 ≈ RO6843766 ≈ JWH133.

### CB2 signaling can be biased by mutations in the ligand binding site

In addition to using structurally different ligands to induce biased signaling response upon activation of CB2, we tested the hypothesis that mutations within the ligand binding pocket can influence the receptor signaling. Using Arpeggio web server for calculating interatomic interactions in protein structures^69^, we detected and visualized interactions between CB2 model and a potent agonist HU210, identifying bulky and hydrophobic residues making the most extensive contacts with the receptor ligand binding pocket (F87, F94, F183, W194, F281; Figure S2). Those residues were mutated into alanine, and the effect of mutations on the receptor signaling profile upon activation with HU210 was monitored using the BRET biosensors (Figure 7). Mutations had dramatic and distinct effects on the signaling profile of CB2 suggesting a role for some of these residues in directing functional selectivity. Mutation W194A completely abolished receptor signaling via G_i2_ and β-arrestin2. F87A lowered the receptor’s affinity for G protein activation more than for β-arrestin recruitment, making the receptor preferentially signal via β-arrestin. Two mutations, F183A and F281A, affected only β-arrestin2 and not β-arrestin1, either via reduced maximal ligand-induced response (F281A) or reduced ligand potency (F183A) (Figure 7). F94A did not alter CB2 signaling.

**Figure 7.**
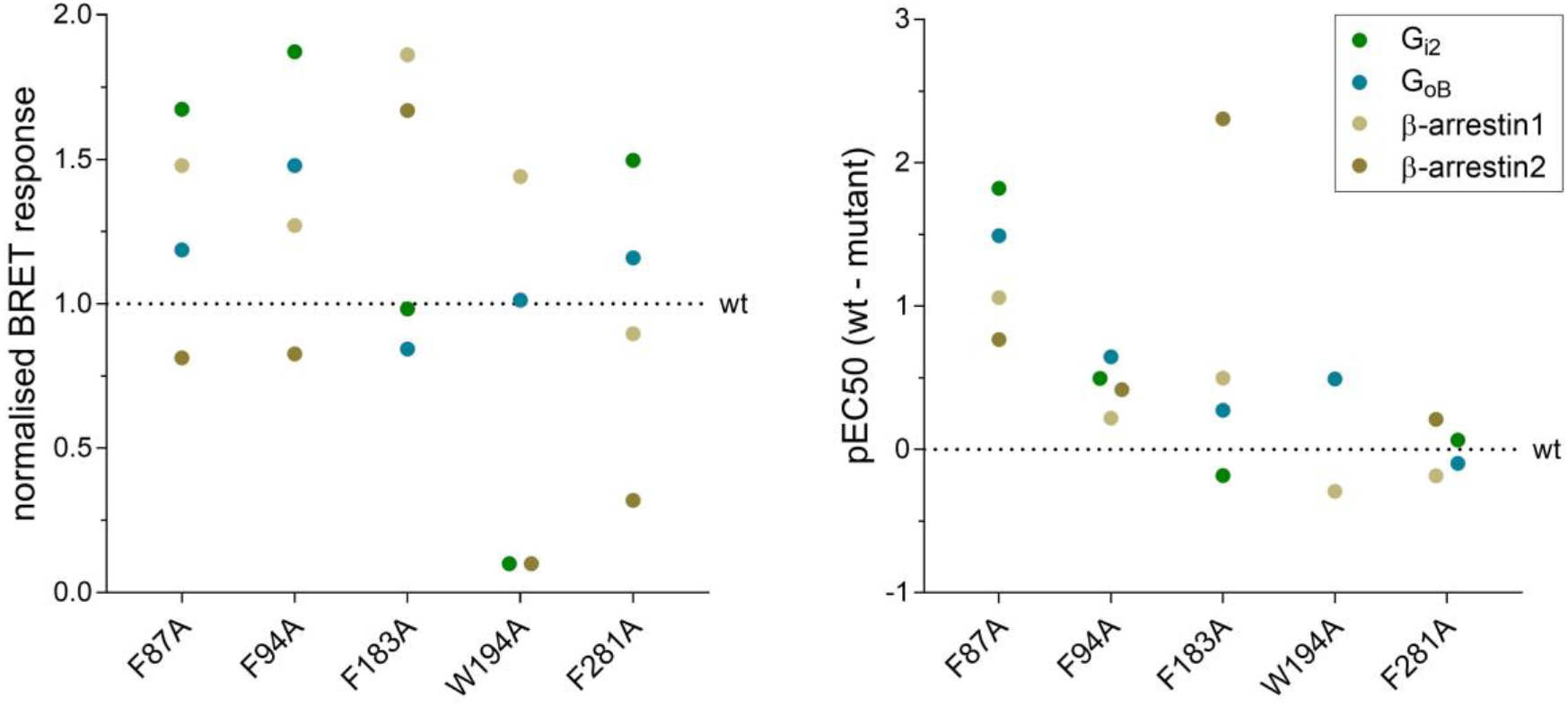
Agonist-induced signaling of CB2 mutants: dot-charts showing maximal relative HU210-induced response values (E_max_) normalized to the wild-type receptor (left) and negative logarithm of half-maximal response concentration (pEC50) subtracted from wild-type CB2 value for each pathway (right). Mean values of three independent experiments are shown. Concentration-dependent signaling shown in Figure S4.

## Discussion

The set of 35 ligands we used contains representatives from 9 different chemotypes which are very diverse not only with regard to their chemical structure, but also with regard to their *in vitro* pharmacology profile and physicochemical properties.

We confirmed that both cannabinoid receptors activate the G_i/o_ family of G proteins. In addition, CB1 also activated G_z_, G_12/13_, and G_15_ although we did not observe activation of other members of the G_q_ family (G_q_, G_11_, G_14_) upon stimulation by CP55940. We also have not observed activation of the G_s_ reported previously^28^. As G_s_ signaling was only observed at a very high receptor density in the previous study, the most likely explanation for this difference is that G_s_ coupling is weak and could not be observed at receptor expression levels used in our study. Both receptors preferentially recruit β-arrestin2 over β-arrestin1, and β-arrestins only weakly traffic to the endosomes with the receptors upon internalization. This is consistent with both CB1 and CB2 classified as class A (in arrestin-recruitment) GPCRs interacting transiently with β-arrestins when compared to V2R, which is a class B GPCR interacting stably with β-arrestins^70^. A highly discussed question regards β-arrestin recruitment by classical cannabinoids containing dibenzopyran ring, which are either phytocannabinoids or their synthetic analogues. While Dhopeshwarkar and Mackie^32^ observed CB2-mediated β-arrestin recruitment only by nonclassical cannabinoids, Soethoudt *et al.*^38^ claiming that both classical and nonclassical cannabinoids recruit β-arrestins to a certain extent, although both studies used PathHunter^®^ β-arrestin assay in CHO cells (DiscoverX). Our data confirm that both classical and nonclassical cannabinoids are capable of recruiting β-arrestins, where an extensively used classical phytocannabinoid-derived ligand, JWH133, showed significant β-arrestin recruitment via CB2, which is comparable to that of non-classical CP55940.

Compared to THC, JWH133 is lacking a hydroxyl group at position 1 of dibenzopyran ring and has a branched and by one CH2 moiety shorter aliphatic tail at position 3 (Table S1). However, in our study phytocannabinoids THC and cannabinol only weakly recruited β-arrestin2, without measurable β-arrestin1 recruitment. HU210 is structurally very similar to JWH133 in having the same branched aliphatic tail at position 3, but has the aliphatic tail two carbons longer than the THC (Table S12). The branched aliphatic groups interact with the residues 87, 183 and 281 that are important for arrestin vs. G protein signaling, and also makes strong interaction with the W194 at the tip of the aliphatic chain – perhaps explain the dramatic difference in activity between THC and HU210. This suggests that small variations in the ligand structure can have a significant impact on the ability of CB receptors to recruit arrestin, reflecting the observed weaker arrestin recruitment by CB receptors compared to the “strong” recruiters, such as V2R.

Most ligands were more potent in activation of G_i/o_ proteins than recruitment of β-arrestins. Although THC weakly recruited β-arrestins, it was one of the strongest G_i1_ activators via CB1. In case of CB1, which also activates other G protein subtypes, ligands were generally less potent in activating those (eg. G_12_ and G_15_) than G_i/o_ protein members, but more potent than in β-arrestin recruitment. In case of CB2, synthetic cannabinoids HU210, HU308, WIN55212-2, JWH133 and CP55940, as well as the endocannabinoid 2-AG were among the highest-efficacy ligands. Only weak recruitment of β-arrestins was detected by cannabinol, reflecting its structural similarity to THC. Interestingly, phytocannabinoid-derived cannabinol acted as an inverse agonist on CB1-mediated G_15_ activation, while being an agonist for other pathways.

Furthermore, we found the signaling of the endogenous ligand 2-AG to prefer β-arrestin recruitment over G protein activation at both cannabinoid receptors, as judged by the amplitude of the response (efficacy). Although bias factors based on the operational model indicate weakened signaling due to lower potency at signaling via β-arrestins compared to G proteins, efficacy at higher 2-AG concentrations drove the signaling towards β-arrestin. It is important to note that operational model does prioritize potency shifts over efficacy, and therefore may not accurately reflect activity of the weak affinity ligands such as endocannabinoids. Similar behavior occurring only at CB2-mediated β-arrestin1 recruitment was also observed with another endogenous ligand, anandamide. Reported 2-AG and anandamide concentrations in human blood are in the nanomolar range^71^, with possibly higher local concentrations around the transmembrane receptors due to their high lipophilicity. Therefore, the range of ligand concentrations used within our study corresponds to the physiological endocannabinoid concentrations. Given the importance of GPCR desensitization via arrestin-dependent internalization, as well as the arrestin-mediated signaling, the observation of endocannabinoids preferentially activating β-arrestins over G proteins at higher ligand concentrations as judged by higher amplitudes of the responses could hold answers to physiological regulation of endocannabinoid system signaling.

The two tested CB2 inverse agonists, AM630 and SR144528, evoked stronger inhibition on signaling via G_i_ proteins than G_o_ proteins and β-arrestins, suggesting that CB2 constitutive activity is composed mostly of G_i_ protein signaling, or that those inverse agonists are less able to inhibit signaling via other G proteins. Although the inverse agonists mainly inhibited G_i_ signaling, constitutive activity of CB2 via β-arrestin pathway was also suppressed, indicating receptor’s constant desensitization and/or constant β-arrestin mediated signaling.

Relative effectiveness and bias factors were calculated for all the tested ligands and pathways using the operational model of biased agonism (Tables S14–S17). Although this method has an advantage of comparing ligand bias independently of the biological system used to asses signaling^72^, one needs to be mindful when interpreting the results due to the large error propagation of the calculated bias factors (Table S15 and Table S17). We have observed classification of ligands into three broad classes of full agonists, agonists with reduced activity across board (eg. partial agonists) and agonists with “unbalanced” activity changes, eg. biased agonists (Figure S1). While strong correlation was observed among G_i_ family members, somewhat reduced correlation was observed between G_i_ vs. G_o_, G_i_ vs. β-arrestin and among G_o_ family responses (Figure S3). The weakest correlation was observed between the two β-arrestin responses (Figure S3). This suggests that, while it is nearly impossible to evoke a bias between G_i1_ and G_i2_, for example, it is possible to develop a G_i_ vs. arrestin or even G_i_ vs. G_o_ biased ligands.

## Conclusion

Our detailed analysis of G protein activation and β-arrestin recruitment by CB1 and CB2 receptors showed an unexpected pleiotropy of signaling preferences by different ligands. This observation is consistent with the results of Wouters *et al.*^39^, who demonstrated that CB1 readily recruits arrestin with a set of 21 synthetic CB1 ligands, and that its signaling can be biased depending on the structure of the ligand used. For example, we show that the alkyl groups that are specific to HU210 relative to THC interact with the residues involved in biased signaling. Importantly, we showed that the endocannabinoids show significant preference towards arrestin recruitment, relative to the available synthetic cannabinoids and phytocannabinoids. Our data also imply that it is possible to derive artificial ligands with specific biased signaling properties both useful for the understanding of the biology of endocannabinoid system, as well as for therapeutic use. In addition, by introducing mutations within the ligand binding pocket, we showed that disturbing receptor-ligand interactions can result in altered signaling profile and receptors biased towards G protein or β-arrestin signaling. Taken together, our data suggests that CB2 signaling can be readily influenced by the respective interacting ligands of diverse structures, providing a possible explanation for the complex physiological effects described for CB2 activation. It further confirms the importance of ligand structure and the interactions they form with the receptor in signaling, making it possible to design ligands with desired signaling outcome.

More research is needed to understand which signaling pathways downstream of cannabinoid receptors are pathologically important, and which could be pharmacologically exploited. At the same time, more structural data are needed to link the ligand binding pose and the interactions it makes to the receptor to its signaling properties – but this link definitely exists. Novel artificial ligands reported here can be used as powerful tools to study the effects of biased signaling in inflammatory cell or animal models, due to their favourable physicochemical properties and metabolic stability. Eventually, the data collected in this study can be utilized to design therapeutics that boost signaling only via the required pathway, leading to health improvement with less side effects.

## Supporting information

SI concentration-response curves

## Nonstandard abbreviations

BRET: bioluminescence resonance energy transfer
cAMP: cyclic 3’-5’ adenosine monophosphate
CB1: cannabinoid 1 receptor
CB2: cannabinoid 2 receptor
DBC: 1-bisdeoxycoelenterazine, di-dehydrocoelenterazine, Coelenterazine 400a, DeepBlueCTM
pEC50: negative logarithm of half-maximal response concentration
E_max_: maximal ligand-induced response
ESI-TOF: electrospray ionization time-of-flight mass spectrometry
GPCRs: G protein-coupled receptors
hCB1: human CB1 receptor
hCB2: human CB2 receptor
HRMS: high-resolution mass spectra
mCB2: mouse CB2 receptor
NBD: 4-nitro-1,2,3-benzoxadiazole fluorophore
THC: Δ^9^-THC, (-)-trans-Δ9-tetrahydrocannabinol
TLC: thin-layer chromatography

## Acknowledgements

The technical assistance of Christian Bartelmus, Markus Binder, Sophie Brogly, Markus Bürkler, Virginie Micallef, Isabelle Parrilla and Björn Wagner is greatly acknowledged. This work was supported by the Swiss National Science Foundation grants 135754 and 159748 to DBV; A Foundation grant (# 148431) from the Canadian Instatute of Health Research (CIHR) to MB; EMBO Short-term Fellowship (420-2016) and Swiss National Science Foundation Doc.Mobility supplement to TM. MB holds the Canada Research Chair in Signal Transduction and Molecular Pharmacology.

## Authorship Contributions

Participated in research design: TM, FMH, WG, UG, MB, DBV

Conducted experiments: TM, EZ, AO

Contributed new reagents or analytic tools: UG, AK, MRE, CU, M Nettekoven, M Nazaré, TG, CLG, MB

Performed data analysis: TM, DBV

Wrote or contributed to the writing of the manuscript: TM, AR, UG, JF, MB, DBV

## Materials and Methods

### Ligands

2-AG, anandamide, Δ^9^-THC, nabilone and HU210 were purchased from Toronto Research Chemicals, Canada. Cannabinol, (+)-WIN55,212-2 (mesylate), (-)-CP55,940, (*R*)-AM1241, (*S*)-AM1241 and (*rac*)-AM1241 were purchased from Cayman Chemical, USA. JWH133, HU308, AM630, SR144508 were purchased from Cedarlane-Tocris, Canada. Structures, IUPAC names and CAS numbers of the screened compounds are given in Table S1.

### General procedure

All reactions were performed using oven or flame-dried glassware and dry solvents. Reagents were purchased from Sigma Aldrich, Acros, Alfa Aesar, Apollo Scientific, TCI, or Merck, and used without further purification unless noted otherwise. All moisture-sensitive reactions were performed under a nitrogen atmosphere. ^1^H-NMR and ^13^C-NMR spectra were recorded either on AV 300 MHz, on AV 600 MHz or on AV 750 (for cryoprobes) from Bruker. Chemical shifts are recorded in parts per million (ppm). Spin multiplicities are described as s (singlet), d (dublet), t (triplet), q (quartet), dq (doublet of quartets) and m (multiplet). Coupling constant (*J*) are recorded in Hz. NMR data were analyzed with MestReNova software. All ^13^C-NMR-spectra were recorded with ^1^H-broad-band decoupling. All chemical shifts are reported in ppm relative to tetramethylsilane (δ = 0.00 ppm) and were calibrated with respect to their respective deuterated solvents. High-resolution mass spectra (HRMS) were recorded on an Agilent Technologies 6220 Accurate Mass TOF LC-MS apparatus linked to Agilent Technologies HPLC 1200 Series, using Thermo Accuore RP-MS column. Liquid chromatography–mass spectrometry (LC-MS) analysis were performed on an Agilent Technologies 6120 Quadrupole LC-MS apparatus linked to Agilent Technologies HPLC 1290 Infinity, using Thermo Accuore RP-MS column. Flash chromatography was performed using SiliCycle silica gel type SiliaFlash P60 (230 – 400 mesh). TLC analysis was performed on Merck silica gel 60/Kieselguhr F254, 0.25 mm.

### Synthesis of RO6844112, methyl N-{6-[(4-fluorophenyl)methyl]pyridine-2-carbonyl}-L-leucinate

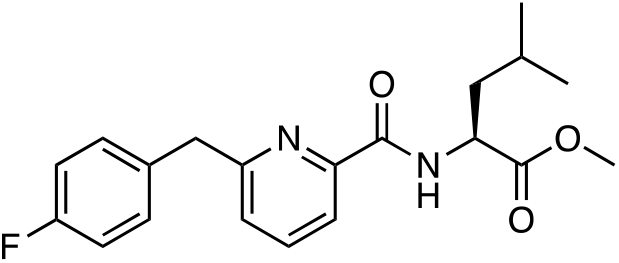

A mixture of 6-(4-fluorobenzyl)picolinic acid (CAS number 1415899-72-7; 20 mg, 86.5 μmol), (*S*)-methyl 2-amino-4-methylpentanoate hydrochloride (CAS number 7517-19-3; 18.9 mg, 104 μmol), 1-hydroxybenzotriazoloe hydrate (26.5 mg, 173 μmol) and *N*-ethyl-*N*-isopropylpropan-2-amine (44.7 mg, 60.4 μL, 346 μmol) in *N,N-*dimethyl formamide (400 μL) was stirred at ambient temperature for 16 h. The reaction mixture was poured onto 1 M HCl / ice water 1:1 (1 x 20 mL) and extracted with ethyl acetate (2 x 25 mL). The combined extracts were washed with ice water (2 x 25 mL), dried over Na_2_SO_4_ and filtered off. The solvent was removed under reduced pressure to give the crude which was purified by preparative thin layer chromatography (silica gel, 2.0 mm, heptane / ethyl acetate 2:1, elution with ethyl acetate) to give the title compound (23 mg, 74%) as colorless liquid. ^1^H NMR (500 MHz, DMSO-*d*6) δ 8.71 (br d, *J*=8.2 Hz, 1H), 7.89 - 7.95 (m, 1H), 7.82 - 7.87 (m, 1H), 7.48 (dd, *J*=7.6, 0.8 Hz, 1H), 7.37 - 7.43 (m, 2H), 7.08 - 7.17 (m, 2H), 4.56 (ddd, *J*=10.0, 8.3, 4.6 Hz, 1H), 4.20 (s, 2 H), 3.66 (s, 3H), 1.83 (ddd, *J*=13.4, 10.0, 4.9 Hz, 1H), 1.62 - 1.70 (m, 1H), 1.55 - 1.63 (m, 1H), 0.90 (dd, *J*=17.5, 6.4 Hz, 6H). ^13^C NMR (500 MHz, DMSO-*d*6) δ 172.8, 164.0, 161.2, 160.0, 148.9, 138.9, 135.7, 131.3, 126.4, 120.0, 115.5, 52.4, 50.7, 42.6, 40.1, 24.8, 23.1, 21.7. HRMS (pos. ESI): m/z calculated for C_20_H_23_FN_2_O_3_ [M+H]^+^ 359.1771, found 359.1771.

### Synthesis of RO5135445, methyl N-[6-(cyclobutylmethoxy)-5-(pyrrolidin-1-yl)pyrazine-2-carbonyl]-L-valinate

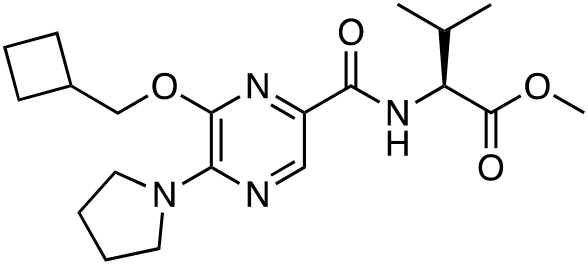

A solution of bromo-tris-pyrrolidino phosphoniumhexafluorophosphate (PyBOP) (156 mg, 0.3 mmol) in *N,N*-dimethylformamide (0.5 mL) was added to a solution of 6-cyclobutylmethoxy-5-pyrrolidin-1-yl-pyrazine-2-carboxylic acid (CAS number 1017603-71-2; 62 mg, 0.2 mmol) in *N,N*-dimethylformamide (1 mL). The mixture was shaken for 20 minutes. L-Valine methyl ester (CAS number 4070-48-8; 52 mg, 0.4 mmol) and *N*-ethyldiisopropylamine (DIPEA, 103 mg, 0.8 mmol) were added portion-wise and the reaction mixture was shaken for 16 hours. The solvent was removed *in vacuo* and the crude product was purified by preparative HPLC to afford methyl *N*-[6-(cyclobutylmethoxy)-5-(pyrrolidin-1-yl)pyrazine-2-carbonyl]-L-valinate (5.7 mg, 7%) as white solid. ^1^H NMR (500 MHz, DMSO-*d*6) δ 8.16 (s, 1H), 7.82 (d, *J*=8.5 Hz, 1H), 4.36 (dd, *J*=8.5, 5.9 Hz, 1H), 4.28 - 4.34 (m, 2H), 3.66 - 3.70 (m, 3H), 3.66 - 3.75 (m, 4H), 2.75 - 2.85 (m, 1H), 2.21 (dq, *J*=13.2, 6.7 Hz, 1H), 2.02 - 2.11 (m, 2H), 1.85 - 1.93 (m, 3H), 1.85 - 1.93 (m, 2H), 1.85 - 1.94 (m, 3H), 0.90 - 0.95 (m, 3H), 0.92 (d, *J*=6.9 Hz, 3H). ^13^C NMR (500 MHz, DMSO-*d*6) δ 172.6, 164.1, 147.2, 146.9, 134.8, 127.4, 70.1, 57.3, 52.3, 49.5, 34.0, 30.5, 25.2, 24.8, 19.3, 18.4, 18.2. HRMS (pos. ESI): m/z calculated for C_20_H_30_N_4_O_4_ [M+H]^+^ 391.2343, found 391.2344.

### Synthesis of RO6892033, 6-(3-*tert-*Butyl-1,2,4-oxadiazol-5-yl)-3-cyclopropyl-2-[(4-fluorophenyl)methyl]pyridine

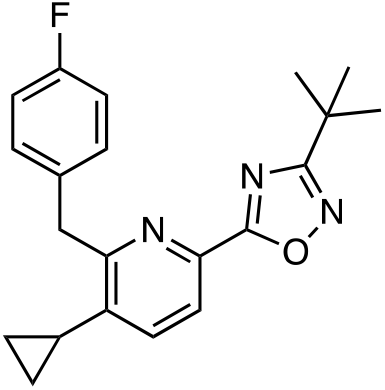

1,1′-Carbonyldiimidazole (13.4 mg, 83 μmol) was added to a solution of 5-cyclopropyl-6-(4-fluorobenzyl)picolinic acid (CAS number 1415899-48-7; 15 mg, 55.3 μmol) in *N,N*-dimethylformamide (643 μL). The mixture was stirred for 30 minutes at ambient temperature. *N*’-Hydroxypivalimidamide (CAS number 42956-75-2; 9.63 mg, 82.9 μmol) was added and the reaction mixture was stirred for 1 h at ambient temperature. The temperature was increased to 100 °C and stirring was continued for 72 h. The reaction mixture was purified by preparative HPLC and subsequent preparative thin layer chromatography (silica gel, 1.0 mm, heptane / ethyl acetate 3:1, elution with ethyl acetate) to give the title compound (6 mg, 31%) as colorless liquid. ^1^H NMR (500 MHz, DMSO-*d*6) δ 8.01 (d, *J*=8.0 Hz, 1H), 7.58 (d, *J*=8.1 Hz, 1H), 7.27 (dd, *J*=8.6, 5.6 Hz, 2H), 7.11 (t, *J*=8.9 Hz, 2H), 4.40 (s, 2H), 2.02 - 2.12 (m, 1H), 1.38 (s, 9H), 0.94 - 1.05 (m, 2H), 0.68 - 0.76 (m, 2H). ^13^C NMR (500 MHz, DMSO-*d*6) δ 178.1, 174.2, 161.1, 160.4, 141.7, 139.8, 135.3, 134.5, 130.7, 123.0, 115.5, 40.4, 32.6, 28.6, 12.6, 9.0. HRMS (pos. ESI): m/z calculated for C_21_H_22_FN_3_O [M+H]^+^ 352.1826, found 352.1825.

### Synthesis of FMP7234690, FMP7234691, FMP7234694, FMP7234698 and FMP7234699

The synthesis of the 2,5,6-trisubstituted pyridine series (compounds FMP7234690, FMP7234691, FMP7234694, FMP7234698 and FMP7234699) commences with the synthesis of the key intermediates ethyl 2-amino-2-ethylpent-4-enoate (**3**) and 5-cyclopropyl-6-(4-fluorobenzyl)picolinic acid (**4**) (Scheme 1). Starting from commercially available racemic amino butyric acid (**1**), classical Fischer esterification conditions, followed by imine formation using benzaldehyde under basic conditions, gave imine (**2**) in good 93% yield, over steps a and b. Lithium-assisted deprotonation/alkylation of imine (**2**) using LDA (lithium di-isopropyl amide) and allylbromide, followed by imine deprotection under acidic conditions yielded key intermediate (**3**) in overall 96% yield. The synthesis of picolinic acid (**4**) have been described by Bissantz and collaborators^56^ and was synthesized as reported. Amide coupling conditions using key intermediates (**3**) and (**4**) gave amide (**5**) in moderate 56% yield. Brown-hydroboration reaction in nearly quantitative yield (99%), followed by Mitsunobu reaction using thioacetic acid provided thioester (**6**) in 63% yield. Cleavage of the acetylthio group, followed by one-pot alkylation with the corresponding linker was achieved under strong basic conditions. Subsequent deprotection of the *boc*-group afforded amines (**7**– **11**) in moderate yields (46 – 56%). Final fluorophore coupling using 4-chloro-7-nitrobenzofurazan (NBD-chloride) and cesium carbonate as base provided compounds FMP7234690, FMP7234691, FMP7234694, FMP7234698 and FMP7234699.

**Scheme 1.**
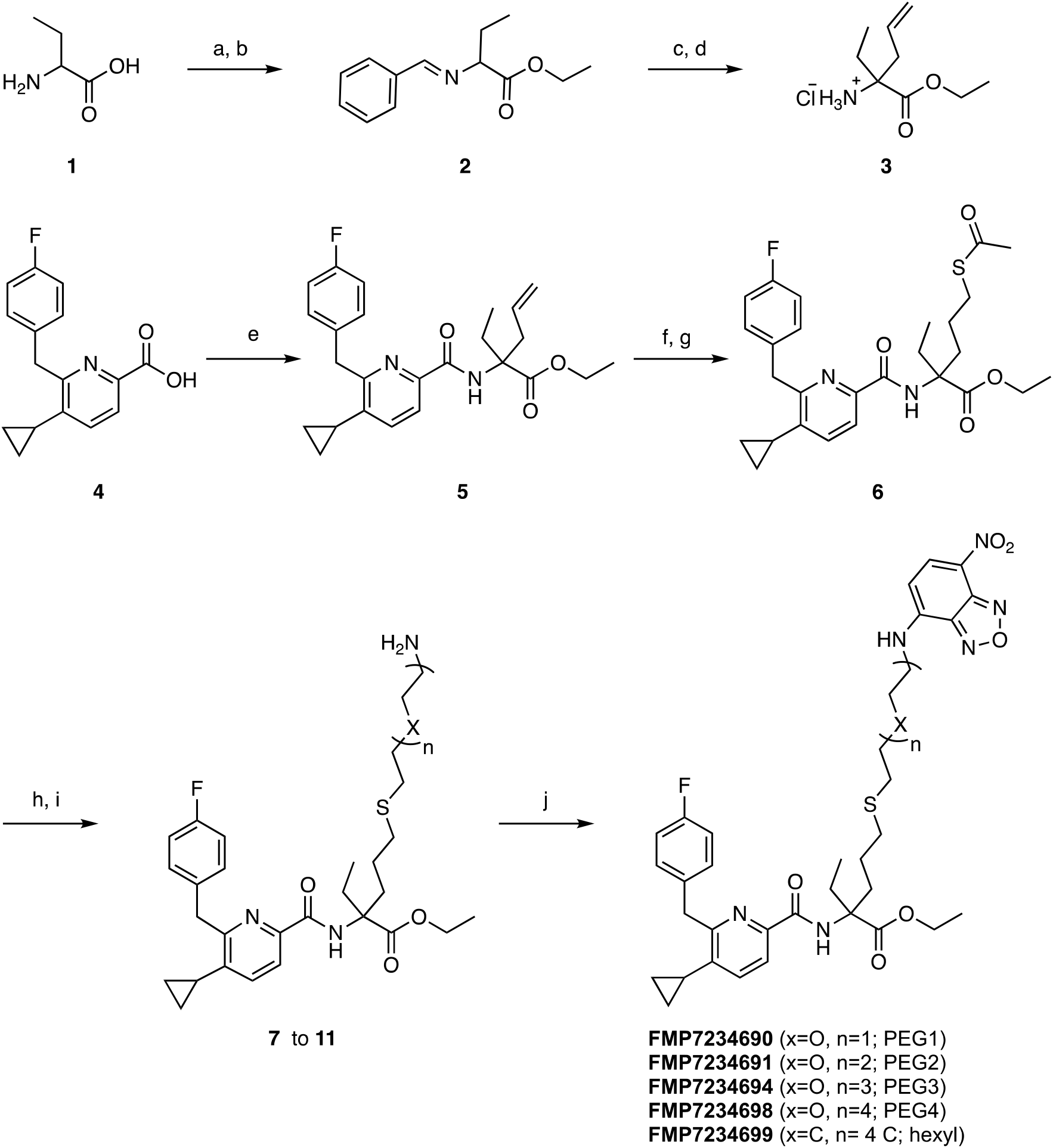
Reagents and conditions: a) thionyl chloride, EtOH, 0 to 78 °C, 5 h; b) Benzaldehyde, Et_3_N, MgSO_4_, DCM, room temperature, 30 h (93%, over 2 steps); c) Allyl Bromide, LDA, THF, −78 °C to room temperature, 24 h; d) HCl aq., H_2_O, Et_2_O, 0 °C to room temperature, 15 h (96%, over 2 steps); e) **14**, BOP-Cl, DIPEA, DCM, room temperature, 30 h (54%); f) (*i*) 9-BBN, THF, room temperature, 36 h; (*ii*) EtOH, room temperature, 30 min; then NaOH, H_2_O_2_, 0 °C, 1 h (99%, over 2 steps); g) thioacetic acid, DIAD, PPh_3_, THF, 0 °C, 2 h (64%); h) linker, EtONa, EtOH, −20 °C to room temperature, 18 h (46 – 56%); i) TFA:DCM (9:1), 0 °C to room temperature, 2 h; j) 4-Chloro-7-nitrobenzofurazan, Cs_2_CO_3_, DMF, 0 °C to room temperature, 24 h (18 – 49%, over 2 steps).

#### Ethyl 2-(benzylideneamino)butanoate(2)

Thionyl chloride (3.5 mL, 47.9 mmol) was added dropwise to a solution of amino butyric acid **1** (CAS number 2835-81-6; 3.8 g, 36.9 mmol) in ethanol (40 mL), over a period of 5 min at 0 °C. The reaction mixture was stirred at 0 °C for 1 additional hour. Afterwards, the resulting solution was refluxed (80 °C) for 4 h. The reaction mixture was cooled to room temperature and concentrated under reduced pressure to yield ethyl 2-aminobutanoate as a colorless oil (4.8 g, quant.), used without further purification for the next step. Ethyl 2-aminobutanoate (4.8 g, 36.9 mmol) and dried magnesium sulfate (4.4 g, 36.9 mmol) were stirred in anhydrous dichloromethane (DCM; 30 mL) at ambient temperature for 20 min. Afterwards, benzaldehyde (3.8 mL, 36.9 mmol) and triethylamine (9.5 mL, 68.2 mmol) were added sequentially and dropwise. The resulting mixture was stirred for 30 h at the same temperature then filtered and concentrated. The residue was dissolved in ether (8 mL) and water (8 mL) and the separated aqueous layer was extracted with ether. The combined ether solutions were washed with brine (8 mL), dried (MgSO_4_), filtered and concentrated under reduced pressure to give the desired imine **2** as a clear oil (7.53 g, 93%). ^1^H NMR (600 MHz, Chloroform-*d*) δ ppm 8.28 (s, 1H), 7.80 – 7.76 (m, 2H), 7.44 – 7.39 (m, 3H), 4.25 – 4.17 (m, 2H), 3.87 (dd, *J* = 8.2, 5.5 Hz, 1H), 2.09 – 2.00 (m, 1H), 1.94 – 1.88 (m, 1H), 1.27 (t, *J* = 7.1 Hz, 3H), 0.92 (t, *J* = 7.4 Hz, 3H). ^13^C NMR (150 MHz, Chloroform-*d*) δ ppm 172.3, 163.2, 135.9, 131.1, 128.7, 75.1, 61.0, 26.78, 14.34, 10.6. HRMS (pos. ESI-TOF): m/z calculated for C_13_H_17_NO_2_ [M+H]^+^ 220.1259, found 220.1351.

#### Ethyl 2-amino-2-ethylpent-4-enoate(3)

To a solution of lithium diisopropylamine (LDA; 4.2 mL, 8.4 mmol) in tetrahydrofuran (4 mL) cooled to –78 °C was added ethyl 2-(benzylideneamino) butyrate **2** (1.2 g, 5.61 mmol) in tetrahydrofuran (4 mL), followed by the dropwise addition of allyl bromide (0.73 mL, 8.41 mmol). The mixture was stirred at room temperature for 24 h, and then concentrated under reduced pressure. The residue was then partitioned between ethyl acetate and water. The organic layer was separated, and the aqueous phase was extracted with ethyl acetate (4x). The combined organic extracts were washed with brine, dried (MgSO_4_), filtered and concentrated under reduced pressure. The crude residue was dissolved in diethyl ether (15 mL) and treated with hydrochloric acid (1 M, 3.5 equiv., 21.9 mL) at 0 °C. The reaction mixture was allowed to warm to room temperature and stirred for additional 15 h. The ether layer was separated, and the water phase extracted with dichloromethane (2x). The dichloromethane extracts were extracted with hydrochloric acid (0.2 M, 2x). The aqueous layers were combined and lyophilized to yield the corresponding amino ester hydrochloric salt (1.1 g, 96%), without the need of further purification steps. ^1^H NMR (300 MHz, Chloroform-*d*) δ ppm 5.74 – 5.60 (m, 1H), 5.21 – 5.05 (m, 2H), 4.15 (q, *J* = 7.1 Hz, 2H), 2.53 (dd, *J* = 13.5, 6.4 Hz, 1H), 2.21 (dd, *J* = 13.5, 8.4 Hz, 1H), 1.84 – 1.66 (m, 3H), 1.60 – 1.48 (m, 1H), 1.25 (t, *J* = 7.1 Hz, 3H), 0.83 (t, *J* = 7.5 Hz, 3H). ^13^C NMR (75 MHz, Chloroform-*d*) δ ppm 176.6, 132.8, 119.3, 119.2, 60.8, 43.8, 32.7, 14.2, 8.1. HRMS (pos. ESI-TOF): m/z calculated for C_9_H_17_NO_2_ [M+H]^+^ 172.1259, found 172.1337.

#### Ethyl 2-(5-cyclopropyl-6-(4-fluorobenzyl)picolinamido)-2-ethylpent-4-enoate(5)

To a solution of 5-cyclopropyl-6-(4-fluorobenzyl)picolinic acid **4**^56^ (300.0 mg, 1.01 mmol) in dichloromethane (7 mL) at room temperature were added DIPEA (0.87 mL, 5.05 mmol), bis-(2-oxo-3-oxazolidinyl)phosphinic chloride (BOP-Cl; 310.6 mg, 1.22 mmol). The mixture was stirred at room temperature for 1 h before amino ester **3** (172.9 mg, 1.01 mmol) in dichloromethane (3 mL) was added. The mixture was stirred at room temperature for 24 h, diluted with dichloromethane (10 mL) and washed with hydrochloric acid (0.2 M, 3x) and brine (1x). The organic layer was dried (MgSO_4_) and concentrated under reduced pressure. Purification was performed by flash chromatography (silica gel, 4 g, 10% ethyl acetate in cyclohexane) to yield amide **5** as a yellow oil (238.8 mg, 56%). ^1^H NMR (300 MHz, Chloroform-*d*) δ ppm 8.95 (s, 1H), 7.92 (d, *J* = 7.9 Hz, 1H), 7.40 (d, *J* = 8.0 Hz, 1H), 7.31 – 7.20 (m, 2H), 6.97 (t, *J* = 8.6 Hz, 2H), 5.72 – 5.58 (m, 1H), 5.16 – 4.96 (m, 2H), 4.43 – 4.22 (m, 4H), 3.30 (dd, *J* = 14.0, 6.9 Hz, 1H), 2.69 – 2.49 (m, 2H), 2.00 – 1.88 (m, 2H), 1.33 (t, *J* = 7.1 Hz, 3H), 1.05 – 0.96 (m, 2H), 0.81 (t, *J* = 7.4 Hz, 3H), 0.83 – 0.65 (m, 2H). ^13^C NMR (75 MHz, Chloroform-*d*) δ ppm 173.1, 164.3, 163.0, 159.8, 158.3, 146.7, 139.7, 134.6, 134.5, 132.6, 130.4, 130.3, 119.6, 118.4, 115.2, 114.9, 64.8, 61.5, 40.6, 39.2, 28.1, 14.2, 12.6, 8.3, 7.7. HRMS (pos. ESI-TOF): m/z calculated for C_25_H_29_FN_2_O_3_ [M+Na]^+^ 447.2162, found 447.2083.

### Ethyl 5-(acetylthio)-2-(5-cyclopropyl-6-(4-fluorobenzyl)picolinamido)-2-ethylpentano-ate(6)

To a solution of compound **5** (238.8 mg, 0.56 mmol) in tetrahydrofuran (2.0 mL) at room temperature and nitrogen atmosphere were added 9-borabicyclo[3.3.1]nonane (9-BBN; 0.5 M in tetrahydrofuran, 1.7 mL, 0.84 mmol). The mixture was stirred at room temperature for 36 h, followed by quenching the excess of 9-BBN with ethanol (0.11 mL, 1.96 mmol). The mixture was stirred at room temperature for 30 min. followed by the concurrent dropwise addition of sodium hydroxide aq. sol. (2 M, 2.2 mL) and hydrogen peroxide aq. sol. (30%, 2.2 mL) at 0 °C. After complete addition, stirring was continued at the same temperature for additional 1 h. The solution was extracted with ether (3x). Subsequent purification by flash chromatography (silica gel, 15 g, 50% ethyl acetate in cyclohexane) was performed to give intermediate ethyl 2-(5-cyclopropyl-6-(4-fluorobenzyl)picolinamido)-2-ethyl-5-hydroxypentanoate as a colorless solid (245.6 mg, 99%). ^1^H NMR (600 MHz, Chloroform-*d*) δ ppm 9.02 (s, 1H), 7.89 (d, *J* = 8.0 Hz, 1H), 7.27 – 7.24 (m, 3H), 6.99 – 6.94 (m, 2H), 4.41 (s, 2H), 4.36 (s, 2H), 4.33 – 4.27 (m, 2H), 3.62 – 3.55 (m, 1H), 2.65 – 2.60 (m, 1H), 2.56 – 2.52 (m, 1H), 2.05 – 1.95 (m, 1H), 1.95 – 1.85 (m, 2H), 1.55 – 1.49 (m, 1H), 1.43 – 1.35 (m, 1H), 1.33 (td, *J* = 7.1, 1.8 Hz, 3H), 1.02 – 0.98 (m, 2H), 0.77 (td, *J* = 7.4, 2.6 Hz, 3H), 0.67 – 0.65 (m, 2H). ^13^C NMR (151 MHz, CDCl_3_) δ ppm 173.8, 163.6, 160.9, 158.6, 146.9, 140.0, 134.8, 130.5, 130.5, 119.9, 115.3, 115.2, 65.1, 62.8, 61.9, 40.8, 31.5, 28.9, 27.8, 14.4, 14.4, 12.9, 8.6, 7.9. HRMS (pos. ESI-TOF): m/z calculated for C_25_H_31_FN_2_O_4_ [M+Na]^+^ 443.2268, found 443.2337.

Diisopropyl azodicarboxylate (DIAD; 0.045 mL, 0.23 mmol) in anhydrous tetrahydrofuran (0.2 mL) was added dropwise to a stirred solution of triphenylphosphine (60.3 mg, 0.23 mmol) in anhydrous tetrahydrofuran (1.0 mL) at 0 °C under nitrogen atmosphere. The mixture was stirred at this temperature for 30 min, until formation of a white precipitate of Mitsunobu betaine, and a solution of thioacetic acid (0.02 mL, 0.23 mmol) and ethyl 2-(5-cyclopropyl-6-(4-fluorobenzyl)picolinamido)-2-ethyl-5-hydroxypentanoate (51.0 mg, 0.11 mmol) in anhydrous tetrahydrofuran (1.5 mL) was added slowly. The reaction was stirred at 0 °C for 1 h, allowed to reach room temperature, and stirred for additional 1 h. Afterwards, the reaction mixture was concentrated, and the residue taken up into a mixture of diethyl ether and cyclohexane (1:1) and triturated at 0 °C. The resulting white solid was filtered off and washed with diethyl ether and cyclohexane (1:1) mixture. The filtrate was evaporated under reduced pressure, and the residue purified by flash chromatography (silica gel, 4 g, 10% ethyl acetate in cyclohexane) to yield compound **6** as a colorless oil (35.6 mg, 64%). ^1^H NMR (300 MHz, Chloroform-*d*) δ ppm 8.99 (s, 1H), 7.90 (d, J = 7.9 Hz, 1H), 7.38 (d, J = 8.0 Hz, 1H), 7.30 – 7.19 (m, 3H), 7.02 – 6.91 (m, 2H), 4.39 – 4.23 (m, 4H), 2.88 – 2.76 (m, 2H), 2.69 – 2.45 (m, 2H), 2.28 (s, 3H), 2.01 – 1.81 (m, 3H), 1.64 – 1.49 (m, 1H), 1.43 – 1.23 (m, 5H), 1.04 – 0.95 (m, 2H), 0.76 (t, J = 7.4 Hz, 3H), 0.70 – 0.62 (m, 2H). ^13^C NMR (75 MHz, CDCl_3_) δ ppm 195.8, 173.6, 163.5, 160.0, 158.5, 146.8, 140.0, 134.8, 134.8, 130.6, 130.5, 119.8, 115.4, 115.1, 65.0, 61.9, 40.8, 34.4, 30.7, 29.1, 28.6, 24.7, 24.7, 14.4, 12.8, 8.5, 7.9, 7.9. HRMS (pos. ESI-TOF): m/z calculated for C_27_H_33_FN_2_O_4_S [M+Na]^+^ 523.2058, found 523.2169.

#### General Procedure for the linker synthesis

*N-Boc* PEG 2–5, or *N-Boc* hydroxyhexyl (1.0 equiv.) was dissolved in dichloromethane (1.5 mL) and treated with *p*-toluenesulfonyl chloride (3.0 equiv.) and pyridine (5.0 equiv.) at 0 °C. The reaction was allowed to warm to room temperature, and then stirred at 40 °C for 12 h. The mixture was diluted with dichloromethane, and washed with hydrochloric acid (0.2 M, 5 mL), water (5 mL), and brine (5 mL), dried (MgSO_4_) and concentrated under reduced pressure. The crude product was purified by flash chromatography (silica gel, 4 g, 15 to 30% ethyl acetate in cyclohexane), giving a colorless oil.

**2-(2-((*tert*-Butoxycarbonyl)amino)ethoxy)ethyl 4-methylbenzenesulfonate** was obtained in quantitative yield (86.2 mg) starting from *tert*-butyl (2-(2-hydroxyethoxy)ethyl)carbamate (*Boc*NH-PEG2, 50.0 mg, 0.24 mmol). ^1^H NMR (300 MHz, Chloroform-*d*) δ ppm 7.83 (d, *J* = 8.2 Hz, 2H), 7.38 (d, *J* = 8.1 Hz, 2H), 4.82 (s, 1H), 4.18 (dd, *J* = 5.7, 3.6 Hz, 2H), 3.65 (dd, *J* = 5.5, 3.8 Hz, 2H), 3.47 (t, *J* = 5.1 Hz, 2H), 3.26 (q, *J* = 5.2 Hz, 2H), 2.47 (s, 3H), 1.47 (s, 9H). ^13^C NMR (75 MHz, Chloroform-*d*) δ ppm 155.9, 144.9, 133.0, 129.8, 128.0, 79.4, 70.4, 69.1, 68.4, 40.2, 28.4, 21.7. HRMS (pos. ESI-TOF): m/z calculated for C_16_H_25_NO_6_S [M+Na]^+^ 382.1413, found 382.1303.

**2,2-Dimethyl-4-oxo-3,8,11-trioxa-5-azatridecan-13-yl 4-methylbenzenesulfonate** was obtained in quantitative yield (114.1 mg) starting from *tert*-butyl (2-(2-(2-hydroxyethoxy)ethoxy)ethyl)carbamate (*Boc*NH-PEG3, 60.0 mg, 0.24 mmol). ^1^H NMR (300 MHz, Chloroform-*d*) δ ppm 7.81 (d, *J* = 8.1 Hz, 2H), 7.36 (d, *J* = 8.0 Hz, 2H), 4.96 (bs, 1H), 4.18 (t, *J* = 4.8 Hz, 2H), 3.71 (t, *J* = 4.8 Hz, 2H), 3.62 – 3.47 (m, 6H), 3.33 – 3.28 (m, 2H), 2.46 (s, 3H), 1.45 (s, 9H). ^13^C NMR (75 MHz, Chloroform-*d*) δ ppm 156.0, 144.9, 133.0, 129.8, 128.0, 79.3, 70.71, 70.3, 70.2, 69.2, 68.7, 40.3, 28.4, 21.7. HRMS (pos. ESI-TOF): m/z calculated for C_18_H_29_NO_7_S [M+Na]^+^ 426.1658, found 426.1548.

**2,2-Dimethyl-4-oxo-3,8,11,14-tetraoxa-5-azahexadecan-16-yl 4-methylbenzenesulfonate** was obtained in 66% yield (71.0 mg) starting from *tert*-butyl (2-(2-(2-(2-hydroxyethoxy)ethoxy)ethoxy)ethyl)carbamate (*Boc*NH-PEG4, 70.0 mg, 0.24 mmol). ^1^H NMR (300 MHz, Chloroform-*d*) δ ppm 7.82 (d, *J* = 8.2 Hz, 2H), 7.36 (d, *J* = 8.1 Hz, 2H), 5.02 (bs, 1H), 4.23 – 4.14 (m, 2H), 3.75 – 3.67 (m, 2H), 3.62 (s, 8H), 3.56 – 3.53 (m, 2H), 3.34 – 3.29 (m, 2H), 2.47 (s, 3H), 1.45 (s, 9H). ^13^C NMR (75 MHz, Chloroform-*d*) δ ppm 156.0, 144.8, 133.0, 129.8, 128.0, 79.2, 70.8, 70.6, 70.5, 70.2, 69.2, 68.7, 40.5, 28.4, 21.7. HRMS (pos. ESI-TOF): m/z calculated for C_20_H_33_NO_8_S [M+Na]^+^ 465.1941, found 465.2275.

**2,2-Dimethyl-4-oxo-3,8,11,14,17-pentaoxa-5-azanonadecan-19-yl 4-methylbenzenesulfonate** was obtained in 94% yield (111.0 mg) starting from *tert*-butyl (14-hydroxy-3,6,9,12-tetraoxatetradecyl)carbamate (*Boc*NH-PEG5, 81.0 mg, 0.24 mmol). ^1^H NMR (300 MHz, Chloroform-*d*) δ ppm 7.80 (d, *J* = 8.0 Hz, 2H), 7.35 (d, *J* = 7.9 Hz, 2H), 5.04 (bs, 1H), 4.21 – 4.12 (m, 2H), 3.71 – 3.67 (m, 2H), 3.59 – 3.66 (m, 12H), 3.57 – 3.51 (m, 2H), 3.32 – 3.29 (m, 2H), 2.45 (s, 3H), 1.44 (s, 9H). ^13^C NMR (75 MHz, Chloroform-*d*) δ ppm 156.0, 144.8, 133.0, 129.8, 128.0, 79.2, 70.8,70.6, 70.5, 70.2, 69.2, 68.7, 40.4, 28.4, 21.7. HRMS (pos. ESI-TOF): m/z calculated for C_22_H_37_NO_9_S [M+Na]^+^ 530.2215, found 530.1841.

**6-((*tert*-butoxycarbonyl)amino)hexyl 4-methylbenzenesulfonate** was obtained in 42% yield (37.8 mg) starting from *tert*-butyl (6-hydroxyhexyl)carbamate (*Boc*NH-hydroxyhexyl, 52.0 mg, 0.24 mmol). ^1^H NMR (300 MHz, Chloroform-*d*) δ ppm 7.80 (d, *J* = 8.2 Hz, 2H), 7.37 (d, *J* = 8.1 Hz, 2H), 4.52 (s, 1H), 4.03 (t, *J* = 6.4 Hz, 2H), 3.11 – 3.05 (m, 2H), 2.47 (s, 3H), 1.72 – 1.58 (m, 2H), 1.45 (s, 11H), 1.37 – 1.23 (m, 4H). 13C NMR (75 MHz, Chloroform-d) δ ppm 156.0, 144.7, 133.1, 129.8, 127.9, 79.1, 70.5, 40.4, 29.9, 28.7, 28.4, 26.1, 25.1, 22.0. HRMS (pos. ESI-TOF): m/z calculated for C_18_H_29_NO_5_S [M+Na]^+^ 394.1794, found 394.1685.

#### General procedure for synthesis of intermediates 7 – 11

Thioderivative **6** (1.0 equiv.) and the corresponding tosyl linker (1.8 equiv.) were added to oxygen-free absol. ethanol (2 mL) under nitrogen atmosphere. The suspension was cooled to −20 °C, sodium ethoxyde (3.0 equiv.) was added along with catalytic amounts of potassium iodide (0.3 equiv.), and the mixture was allowed to warm up slowly to room temperature. The resulting solution was stirred for 20 h. The reaction mixture was concentrated under reduced pressure, and the resulting residue was dissolved in acetonitrile and water (1:1) and purified on reverse-phase preparative HPLC (30-90% acetonitrile in water with 0.1% trifluoroacetic acid (TFA), 32 min). The fractions containing the product were combined and lyophilized to yield the desired product.

**Ethyl 15-(5-cyclopropyl-6-(4-fluorobenzyl)picolinamido)-15-ethyl-2,2-dimethyl-4-oxo-3,8-dioxa-11-thia-5-azahexadecan-16-oate(7)** was obtained in 50% yield (16.2 mg) starting from 2-(2-((*tert*-butoxycarbonyl)amino)ethoxy)ethyl 4-methylbenzenesulfonate (32.3 mg, 0.09 mmol). ^1^H NMR (300 MHz, Chloroform-*d*) δ ppm 9.02 (s, 1H), 7.89 (d, *J* = 7.9 Hz, 1H), 7.39 (d, *J* = 8.0 Hz, 1H), 7.30 – 7.21 (m, 2H), 6.97 (t, *J* = 8.6 Hz, 2H), 4.95 (bs, 1H), 4.46 – 4.19 (m, 4H), 3.56 – 3.43 (m, 4H), 3.29 – 3.24 (m, 2H), 2.67 – 2.47 (m, 6H), 2.06 – 1.82 (m, 4H), 1.64 – 1.56 (m, 1H), 1.43 (s, 9H), 1.33 (t, *J* = 7.1 Hz, 3H), 1.05 – 0.94 (m, 2H), 0.76 (t, *J* = 7.4 Hz, 3H), 0.79 – 0.64 (m, 2H). ^13^C NMR (75 MHz, CDCl_3_) δ ppm 173.7, 163.5, 163.2, 160.0, 158.5, 156.1, 146.8, 140.1, 134.8, 130.6, 130.5, 119.8, 115.4, 115.1, 79.4, 70.7, 70.0, 65.1, 61.9, 40.8, 40.5, 34.5, 32.5, 31.6, 28.6, 28.6, 24.8, 14.5, 12.8, 8.6, 7.9. HRMS (pos. ESI-TOF): m/z calculated for C_34_H_48_FN_3_O_6_S [M+Na]^+^ 668.3278, found 668.3154.

**Ethyl 18-(5-cyclopropyl-6-(4-fluorobenzyl)picolinamido)-18-ethyl-2,2-dimethyl-4-oxo-3,8,11-trioxa-14-thia-5-azanonadecan-19-oate(8)** was obtained in 49% yield (13.4 mg) starting from 2,2-dimethyl-4-oxo-3,8,11-trioxa-5-azatridecan-13-yl 4-methylbenzenesulfonate (29.1 mg, 0.07 mmol). ^1^H NMR (300 MHz, Chloroform-*d*) δ ppm 9.01 (s, 1H), 7.89 (d, *J* = 7.9 Hz, 1H), 7.38 (d, *J* = 8.0 Hz, 1H), 7.30 – 7.20 (m, 2H), 7.03 – 6.92 (m, 2H), 5.02 (bs, 1H), 4.40 – 4.23 (m, 4H), 3.62 – 3.45 (m, 8H), 3.34 – 3.30 (m, 2H), 2.68 – 2.44 (m, 6H), 2.05 – 1.82 (m, 3H), 1.64 – 1.55 (m, 1H), 1.43 (s, 9H), 1.35 – 1.18 (m, 4H), 1.03 – 0.97 (m, 2H), 0.76 (t, *J* = 7.4 Hz, 3H), 0.72 – 0.62 (m, 2H). ^13^C NMR (75 MHz, CDCl_3_) δ ppm 173.7, 163.5, 163.2, 158.5, 156.1, 146.8, 140.0, 134.8, 130.6, 130.5, 119.8, 115.4, 115.1, 79.3, 71.0, 70.4, 70.3, 65.1, 61.9, 40.9, 40.4, 34.5, 32.5, 31.5, 28.6, 24.9, 14.5, 12.9, 8.6, 7.9. HRMS (pos. ESI-TOF): m/z calculated for C_36_H_52_FN_3_O_7_S [M+Na]^+^ 712.3462, found 712.3346.

**Ethyl 21-(5-cyclopropyl-6-(4-fluorobenzyl)picolinamido)-21-ethyl-2,2-dimethyl-4-oxo-3,8,11,14-tetraoxa-17-thia-5-azadocosan-22-oate(9)** was obtained in 52% yield (19.0 mg) starting from 2,2-dimethyl-4-oxo-3,8,11,14-tetraoxa-5-azahexadecan-16-yl 4-methylbenzenesulfonate (40.3 mg, 0.09 mmol). ^1^H NMR (300 MHz, Chloroform-*d*) δ ppm 9.01 (s, 1H), 7.89 (d, *J* = 7.9 Hz, 1H), 7.38 (d, *J* = 8.0 Hz, 1H), 7.30 – 7.21 (m, 2H), 6.97 (t, *J* = 8.6 Hz, 2H), 5.03 (bs, 1H), 4.43 – 4.21 (m, 4H), 3.66 – 3.68 (m, 12H), 3.33 – 3.27 (m, 2H), 2.69 – 2.44 (m, 6H), 2.08 – 1.76 (m, 4H), 1.64 – 1.55 (m, 1H), 1.43 (s, 9H), 1.33 (t, *J* = 7.1 Hz, 3H), 1.03 – 0.96 (m, 2H), 0.76 (t, *J* = 7.4 Hz, 3H), 0.78 – 0.64 (m, 2H). ^13^C NMR (75 MHz, CDCl_3_) δ ppm 173.7, 163.5, 163.2, 160.0, 158.5, 156.1, 146.8, 134.8, 130.6, 130.5, 119.8, 115.4, 115.1, 79.3, 71.0, 70.7, 70.4, 70.4, 65.1, 61.9, 40.8, 34.5, 32.5, 31.5, 28.6, 24.9, 14.5, 12.8, 8.6, 7.9. HRMS (pos. ESI-TOF): m/z calculated for C_38_H_56_FN_3_O_8_S [M+Na]^+^ 756.3792, found 756.3682.

**Ethyl 24-(5-cyclopropyl-6-(4-fluorobenzyl)picolinamido)-24-ethyl-2,2-dimethyl-4-oxo-3,8,11,14,17-pentaoxa-20-thia-5-azapentacosan-25-oate(10)** was obtained in 56% yield (21.8 mg) starting from 2,2-dimethyl-4-oxo-3,8,11,14,17-pentaoxa-5-azanonadecan-19-yl 4-methylbenzenesulfonate (44.2 mg, 0.09 mmol). ^1^H NMR (300 MHz, Chloroform-*d*) δ ppm 9.01 (s, 1H), 7.89 (d, *J* = 7.9 Hz, 1H), 7.38(d, *J* = 8.0 Hz, 1H), 7.29 – 7.21 (m, 2H), 6.97 (t, *J* = 8.6 Hz, 2H), 5.04 (bs, 1H), 4.39 – 4.24 (m, 4H), 3.68 – 3.47 (m, 16H), 3.35 – 3.30 (m, 2H), 2.69 – 2.42 (m, 6H), 2.08 – 1.80 (m, 4H), 1.65 – 1.51 (m, 1H), 1.43 (s, 9H), 1.33 (t, *J* = 7.1 Hz, 3H), 1.04 – 0.95 (m, 2H), 0.76 (t, *J* = 7.4 Hz, 3H), 0.78 – 0.64 (m, 2H). ^13^C NMR (75 MHz, CDCl_3_) δ ppm 173.7, 163.4, 160.0, 158.5, 156.1, 146.8, 140.1, 134.9, 130.6, 130.5, 119.9, 115.4, 115.1, 79.3, 70.7, 70.7, 70.7, 70.4, 70.3, 65.1, 61.9, 40.8, 40.5, 34.5, 32.5, 31.5, 28.6, 24.8, 14.5, 12.8, 8.6, 7.9. HRMS (pos. ESI-TOF): m/z calculated for C_40_H_60_FN_3_O_9_S [M+Na]^+^ 800.4078, found 800.3942.

**Ethyl 5-((6-((*tert*-butoxycarbonyl)amino)hexyl)thio)-2-(5-cyclopropyl-6-(4-fluorobenzyl) picolinamido)-2-ethylpentanoate(11)** was obtained in 51% yield (16.9 mg) starting from 6-((*tert*-butoxycarbonyl)amino)hexyl 4-methylbenzenesulfonate (33.4 mg, 0.09 mmol). ^1^H NMR (300 MHz, Chloroform-*d*) δ ppm 9.02 (s, 1H), 7.89 (d, *J* = 7.9 Hz, 1H), 7.38 (d, *J* = 8.0 Hz, 1H), 7.32 – 7.19 (m, 2H), 6.96 (t, *J* = 8.6 Hz, 2H), 4.53 (s, 1H), 4.38 – 4.24 (m, 4H), 3.11 – 3.04 (m, 2H), 2.67 – 2.37 (m, 6H), 2.07 – 1.84 (m, 3H), 1.60 – 1.24 (m, 22H), 1.06 – 0.94 (m, 2H), 0.76 (t, *J* = 7.4 Hz, 3H), 0.79 – 0.63 (m, 2H). ^13^C NMR (75 MHz, CDCl_3_) δ ppm 173.8, 163.5, 160.0, 158.3, 156.1, 146.8, 140.0, 134.8, 130.6, 130.5, 119.8, 115.4, 115.1, 79.2, 65.1, 61.9, 40.8, 34.6, 32.1, 32.0, 30.1, 29.6, 28.6, 28.6, 26.5, 24.7, 14.4, 12.8, 8.6, 7.9, 7.9. HRMS (pos. ESI-TOF): m/z calculated for C_36_H_52_FN_3_O_5_S [M+Na]^+^ 680.3664, found 680.3516.

#### General procedure for synthesis of FMP7234690, FMP7234691, FMP7234694, FMP7234698 and FMP7234699

TFA (0.25 mL) was added dropwise to a stirring solution of the corresponding *Boc*-protected intermediate **7**– **11** (1.0 equiv.) in dichloromethane (2.25 mL) at 0 °C. The reaction mixture was stirred at 0 °C for 1 h, and at room temperature for additional 1 h. Afterwards, the mixture was concentrated under reduced pressure, and the residue re-suspended in ethyl acetate (5 mL). The solvent was removed by rotary evaporation. This process was repeated 3 times to remove TFA traces. The removal of the *tert-* butyloxycarbonyl group was quantitative as observed by TLC (50% ethyl acetate in cyclohexane). The product was obtained as its corresponding TFA salt and was used for the next step without further purification. To a solution of free amino intermediate (1.0 equiv.) and Cs_2_CO_3_ (5.0 equiv.) in dimethylformamide (2 mL) at room temperature 4-chloro-7-nitrobenzofurazan (NBD-Cl, CAS number 10199-89-0; 1.0 equiv.) was added. The reaction was stirred at room temperature for 24 h. The reaction mixture was concentrated under reduced pressure, and the obtained residue was diluted with a mixture of acetonitrile and water (1:1) and purified on reverse-phase preparative HPLC (30 to 90% acetonitrile in water with 0.1% TFA, 32 min). The fractions containing the product were combined and lyophilized to yield the desired product.

**Ethyl 2-(5-cyclopropyl-6-(4-fluorobenzyl)picolinamido)-2-ethyl-5-((2-(2-((7-nitrobenzo-[c][1,2,5]oxadiazol-4-yl)amino)ethoxy)ethyl)thio)pentanoate(FMP7234690)** was obtained in 49% yield (4.5 mg) starting from *Boc*-protected **7** (10.0 mg, 0.02 mmol). ^1^H NMR (*cryo* 600 MHz, DMSO-d_6_) δ ppm 9.41 (s, 1H), 8.69 (s, 1H), 8.50 (d, *J* = 8.9 Hz, 1H), 7.74 (d, *J* = 8.0 Hz, 1H), 7.51 (d, *J* = 8.1 Hz, 1H), 7.34 – 7.29 (m, 2H), 7.11 – 7.06 (m, 2H), 6.44 (d, *J* = 9.0 Hz, 1H), 4.34 (s, 2H), 4.20 (q, *J* = 7.1 Hz, 2H), 3.70 – 3.63 (m, 4H), 3.51 (t, *J* = 6.7 Hz, 2H), 2.54 (t, *J* = 6.7 Hz, 2H), 2.46 – 2.41 (m, 2H), 2.34 – 2.24 (m, 2H), 2.09 – 2.05 (m, 1H), 1.88 – 1.75 (m, 2H), 1.38 – 1.32 (m, 2H), 1.20 (t, *J* = 7.1 Hz, 4H), 1.10 – 0.96 (m, 2H), 0.70 – 0.64 (m, 4H). ^13^C NMR (*cryo* 151 MHz, DMSO-d_6_) δ ppm 172.8, 162.3, 161.6, 160.0, 158.2, 158.0, 146.1, 145.3, 144.4, 144.1, 140.3, 137.8, 135.0, 134.4, 130.7, 120.9, 119.3, 114.9, 99.5, 70.2, 67.6, 63.6, 61.4, 43.3, 33.4, 31.3, 30.5, 26.6, 23.9, 14.4, 12.1, 8.2, 8.1, 8.0. HRMS (pos. ESI-TOF): m/z calculated for C_35_H_41_FN_6_O_7_S [M+Na]^+^ 731.2760, found 731.2646.

**Ethyl 2-(5-cyclopropyl-6-(4-fluorobenzyl)picolinamido)-2-ethyl-5-((2-(2-(2-((7-nitrobenzo[c][1,2,5]oxadiazol-4-yl)amino)ethoxy)ethoxy)ethyl)thio)pentanoate(FMP7234691)** was obtained in 26% yield (2.5 mg) starting from *Boc*-protected **8** (10.0 mg, 0.01 mmol). ^1^H NMR (*cryo* 600 MHz, DMSO-d_6_) δ ppm 9.43 (s, 1H), 8.70 (s, 1H), 8.49 (d, *J* = 9.0 Hz, 1H), 7.74 (d, *J* = 8.0 Hz, 1H), 7.51 (d, *J* = 8.0 Hz, 1H), 7.34 – 7.30 (m, 2H), 7.11 – 7.06 (m, 2H), 6.54 (d, *J* = 9.0 Hz, 1H), 4.34 (s, 2H), 4.31 (q, *J* = 7.1 Hz, 2H), 3.69 (t, *J* = 5.3 Hz, 2H), 3.49 – 3.51 (m, 3H), 3.43 – 3.39 (m, 3H), 2.48 (t, *J* = 6.7 Hz, 2H), 2.40 – 2.45 (m, 2H), 2.35 – 2.26 (m, 3H), 2.10 – 2.04 (m, 2H), 1.92 – 1.77 (m, 3H), 1.38 – 1.33 (m, 2H), 1.21 (t, *J* = 7.1 Hz, 3H), 1.01 – 0.96 (m, 2H), 0.70 – 0.65 (m, 4H). ^13^C NMR (*cryo* 151 MHz, DMSO-d_6_) δ ppm 172.8, 162.3, 161.6, 160.0, 158.0, 145.8, 145.3, 144.4, 144.1, 140.3, 137.8, 135.0, 134.4, 130.7, 130.6, 120.8, 119.3, 115.1, 114.9, 99.5, 70.2, 69.8, 69.4, 63.6, 61.4, 43.4, 33.4, 31.3, 30.4, 27.6, 23.9, 14.1, 12.1, 8.1, 8.0. HRMS (pos. ESI-TOF): m/z calculated for C_37_H_45_FN_6_O_8_S [M+Na]^+^ 775.3027, found 775.2911.

**Ethyl 16-(5-cyclopropyl-6-(4-fluorobenzyl)picolinamido)-16-ethyl-1-((7-nitrobenzo[c]-[1,2,5]oxadiazol-4-yl)amino)-3,6,9-trioxa-12-thiaheptadecan-17-oate(FMP7234694)** was obtained in 49% yield (3.1 mg) starting from *Boc*-protected **9** (10.0 mg, 0.01 mmol). ^1^H NMR (*cryo* 600 MHz, DMSO-d_6_) δ ppm 9.44 (s, 1H), 8.70 (s, 1H), 8.50 (d, *J* = 8.8 Hz, 1H), 7.75 (d, *J* = 8.0 Hz, 1H), 7.51 (d, *J* = 8.0 Hz, 1H), 7.34 – 7.30 (m, 2H), 7.12 – 7.06 (m, 2H), 6.47 (d, *J* = 9.0 Hz, 1H), 4.34 (s, 2H), 4.21 (q, *J* = 7.1 Hz, 2H), 3.70 (t, *J* = 5.3 Hz, 2H), 3.70 – 3.62 (m, 2H), 3.55 – 3.53 (m, 2H), 3.49 – 3.47 (m, 3H), 3.41 – 3.39 (m, 5H), 3.37 – 3.34 (m, 2H), 2.45 (td, *J* = 7.1, 3.0 Hz, 2H), 2.37 – 2.26 (m, 2H), 2.10 – 2.05 (m, 2H), 1.92 – 1.87 (m, 1H), 1.84 – 1.79 (m, 1H), 1.41 – 1.33 (m, 2H), 1.21 (t, *J* = 7.1 Hz, 3H), 1.01 – 0.97 (m, 2H), 0.70 – 0.65 (m, 4H). ^13^C NMR (*cryo* 151 MHz, DMSO-d_6_) δ ppm 172.8, 162.3, 161.6, 160.0, 158.0, 145.8, 145.3, 144.1, 140.3, 137.8, 135.0, 134.4, 130.7, 130.6, 120.8, 119.3, 115.1, 114.9, 99.5, 70.1, 69.8, 69.7, 69.4, 68.0, 63.6, 61.4, 43.4, 33.4, 31.3, 30.4, 27.6, 23.9, 14.1, 12.1, 8.1, 8.0. HRMS (pos. ESI-TOF): m/z calculated for C_39_H_49_FN_6_O_9_S [M+Na]^+^ 819.3277, found 819.3161.

**Ethyl 19-(5-cyclopropyl-6-(4-fluorobenzyl)picolinamido)-19-ethyl-1-((7-nitrobenzo[c]-[1,2,5]oxadiazol-4-yl)amino)-3,6,9,12-tetraoxa-15-thiaicosan-20-oate(FMP7234698)** was obtained in 23% yield (2.7 mg) starting from *Boc*-protected **10** (14.0 mg, 0.02 mmol). ^1^H NMR (*cryo* 600 MHz, DMSO-d_6_) δ ppm 9.45 (s, 1H), 8.71 (s, 1H), 8.50 (d, *J* = 9.0 Hz, 1H), 7.75 (d, *J* = 8.0 Hz, 1H), 7.52 (d, *J* = 7.9 Hz, 1H), 7.35 – 7.30 (m, 2H), 7.12 – 7.07 (m, 2H), 6.47 (d, *J* = 9.0 Hz, 1H), 4.34 (s, 2H), 4.21 (q, *J* = 7.1 Hz, 2H), 3.70 (t, *J* = 5.3 Hz, 1H), 3.56 – 3.54 (m, 2H), 3.50 – 3.48 (m, 2H), 3.46 – 3.40 (m, 16H), 3.38 – 3.36 (m, 2H), 2.47 – 2.44 (m, 1H), 2.37 – 2.27 (m, 2H), 2.10 – 2.06 (m, 1H), 1.93 – 1.97 (m, 1H), 1.85 – 1.79 (m, 1H), 1.41 – 1.33 (m, 1H), 122 (t, *J* = 7.1 Hz, 3H), 1.03 – 0.95 (m, 2H), 0.72 – 0.64 (m, 4H). ^13^C NMR (*cryo* 151 MHz, DMSO-d_6_) δ ppm 172.8, 162.3, 161.6, 160.0, 158.0, 145.8, 145.3, 144.4, 144.1, 140.3, 137.8, 135.0, 134.4, 130.7, 130.6, 120.8, 119.3, 115.0, 114.9, 99.5, 70.2, 69.8, 69.7, 69.7, 69.4, 68.0, 63.6, 61.4, 60.2, 43.4, 33.4, 31.3, 30.4, 27.6, 23.9, 14.1, 12.1, 8.2, 8.0. HRMS (pos. ESI-TOF): m/z calculated for C_41_H_53_FN_6_O_10_S [M+Na]^+^ 841.3595, found 841.3658.

**Ethyl 2-(5-cyclopropyl-6-(4-fluorobenzyl)picolinamido)-2-ethyl-5-((6-((7-nitrobenzo[c]-[1,2,5]oxadiazol-4-yl)amino)hexyl)thio)pentanoate(FMP7234699)** was obtained in 18% yield (1.6 mg) starting from *Boc*-protected **11** (11.0 mg, 0.02 mmol). ^1^H NMR (*cryo* 600 MHz, DMSO-d_6_) δ ppm 9.52 (t, *J* = 5.7 Hz, 1H), 8.72 (s, 1H), 8.50 (d, *J* = 8.9 Hz, 1H), 7.75 (d, *J* = 8.0 Hz, 1H), 7.51 (d, *J* = 8.0 Hz, 1H), 7.35 – 7.31 (m, 2H), 7.11 – 7.06 (m, 2H), 6.38 (d, *J* = 9.0 Hz, 1H), 4.34 (d, *J* = 3.5 Hz, 2H), 4.21 (q, *J* = 7.1 Hz, 2H), 3.42 (q, *J* = 6.8, 6.3 Hz, 2H), 2.40 (t, *J* = 7.1 Hz, 2H), 2.37 – 2.27 (m, 4H), 2.10 – 2.06 (m, 1H), 1.93 – 1.78 (m, 3H), 1.59 – 1.64 (m, 2H), 1.44 – 1.20 (m, 11H), 1.02 – 0.96 (m, 2H), 0.70 – 0.65 (m, 4H). ^13^C NMR (*cryo* 151 MHz, DMSO-d_6_) δ ppm 172.8, 162.3, 161.6, 160.0, 158.0, 145.8, 145.2, 144.4, 144.1, 140.3, 138.0, 135.0, 134.4, 130.7, 130.6, 120.5, 119.3, 115.0, 114.9, 99.1, 63.6, 61.4, 43.3, 33.4, 30.8, 30.7, 28.9, 27.8, 27.7, 27.5, 25.9, 23.7, 14.1, 12.1, 8.2, 8.1. HRMS (pos. ESI-TOF): m/z calculated for C_37_H_45_FN_6_O_6_S [M+Na]^+^ 743.3244, found 743.3021.

### *In vitro* pharmacology

#### Human CB2, mouse CB2 and human CB1 ligand binding

Radioligand binding assays were performed with membranes prepared from cells expressing human CB2 or CB1 or mouse CB2 receptor using [^3^H]-CP55940 (Perkin Elmer) as a radioligand. K_i_ values were calculated from a single experiment using triplicates of ten different compound concentrations^73^.

#### Human CB2, mouse CB2 and human CB1 cAMP assay

cAMP assays were performed using CHO cells stably expressing human CB2 or CB1 or mouse CB2, as previously described^73^. Efficacies are expressed as percentage relative to 10 μM CP55940. EC50 values are the average of determinations (n=1) performed in triplicate.

#### Human CB2, mouse CB2 and human CB1 β-arrestin assay

PathHunter β-arrestin recruitment assay was performed using the PathHunter hCB1_bgal, hCB2_bgal or mCB2_bgal CHOK1 β-arrestin recruitment assay kit according to the manufacturer’s protocol as previously described^38^.

### Early ADME profile

#### Kinetic solubility (LYSA – lyophilisation solubility assay)

The solubility of a test compound was measured in phosphate buffer at pH 6.5 from evaporated DMSO compound stock solution as previously described^62^.

#### Passive membrane permeability (PAMPA)

PAMPA (Parallel Artificial Membrane Permeability Assay) is a method which determines the permeability of substances from a donor compartment, through a lipid-infused artificial membrane, into an acceptor compartment. Read-out is a permeation coefficient P_eff_ well as test compound concentrations in donor, membrane and acceptor compartments^63^.

#### Microsomal clearance

For human or mice, pooled commercially available microsome preparations from liver tissues are used^74^. For human, ultra-pooled (150 mixed gender donors) liver microsomes are purchased to account for the biological variance *in vivo*. For the microsome incubations, 96 deep well plates are applied, which are incubated at 37 °C on a TECAN (Tecan Group Ltd, Switzerland) equipped with Te-Shake shakers and a warming device (Tecan Group Ltd, Switzerland). The incubation buffer is 0.1 M phosphate buffer at pH 7.4. The NADPH regenerating system consists of 30 mM glucose-6-phosphate disodium salt hydrate; 10 mM NADP; 30 mM mgCl_2_ x 6 H_2_O and 5 mg/mL glucose-6-phosphate dehydrogenase (Roche Diagnostics) in 0.1 M potassium phosphate buffer pH 7.4. Log peak area ratios (test compound peak area / internal standard peak area) are plotted against incubation time using a linear fit. The calculated slope is used to determine the intrinsic clearance: Clint (μL/min/mg protein) = -slope (min-1) * 1000 / [protein concentration (mg/mL)].

#### Hepatocyte clearance

For animals, hepatocyte suspension cultures are either freshly prepared by liver perfusion studies or prepared from cryopreserved hepatocyte batches. For human, commercially available, pooled (5-20 donors), cryopreserved human hepatocytes from non-transplantable liver tissues are mainly used^75^. For the suspension cultures, Nunc U96 PP-0.5 mL (Nunc Natural, 267245) plates are used, which are incubated in a Thermo Forma incubator from Fischer Scientific (Wohlen, Switzerland) equipped with shakers from Variomag® Teleshake shakers (Sterico, Wangen, Switzerland) for maintaining cell dispersion. The cell culture medium is William’s media supplemented with Glutamine, antibiotics, insulin, dexamethasone and 10% FCS. Incubations of a test compound at 1 μM test concentration in suspension cultures of 1×10^6^ cells/mL (~1 mg/mL protein concentration) are performed in 96 well plates and shaken at 900 rpm for up to 2 h in a 5% CO_2_ atmosphere and 37 °C. After 3, 6, 10, 20, 40, 60 and 120 minutes 100 μL cell suspension in each well is quenched with 200 μL methanol containing an internal standard. Samples are then cooled and centrifuged before analysis by LC-MS/MS. Log peak area ratios (test compound peak area / internal standard peak area) or concentrations are plotted against incubation time and a linear fit made to the data with emphasis upon the initial rate of compound disappearance. The slope of the fit is then used to calculate the intrinsic clearance: Clint (μL/min/1×106 cells) = -slope (min-1) * 1000 / [1×106 cells].

#### Plasma protein binding

Pooled and frozen plasma from selected species were obtained from commercial suppliers ^76,77^. The Teflon equilibrium dialysis plate (96-well, 150 μL, half-cell capacity) and cellulose membranes (12–14 kDa molecular weight cut-off) were purchased from HT-Dialysis (Gales Ferry, Connecticut). Both biological matrix and phosphate buffer pH are adjusted to 7.4 on the day of the experiment. The reference substance is diazepam. The determination of unbound compound is performed using a 96-well format equilibrium dialysis device with a molecular weight cut-off membrane of 12 to 14 kDa. The equilibrium dialysis device itself is made of Teflon to minimize non-specific binding of the test substance. Compounds are tested in cassettes of 2-5 with an initial total concentration of 1000 nM, one of the cassette compounds being the positive control diazepam. Equal volumes of matrix samples containing substances and blank dialysis buffer (Soerensen buffer at pH 7.4) are loaded into the opposite compartments of each well. The dialysis block is sealed and kept for 5 h at a temperature of 37 °C and 5% CO_2_ environment in an incubator. After this time, equilibrium will have been reached for most small molecule compounds with a molecular weight of < 600. The seal is then removed and matrix and buffer from each dialysis is prepared for analysis by LC-MS/MS. All protein binding determinations are performed in triplicates. The integrity of membranes is tested in the HTDialysis device by determining the unbound fraction values for the positive control diazepam in each well. At equilibrium, the unbound drug concentration in the biological matrix compartment of the equilibrium dialysis apparatus is the same as the concentration of the compound in the buffer compartment. Thus, the percent unbound fraction (fu) can be calculated by determining the compound concentrations in the buffer and matrix compartments after dialysis as follows: %fu = 100 * buffer conc after dialysis / matrix conc after dialysis. The device recovery is checked by measuring the compound concentrations in the matrix before dialysis and calculating the percent recovery (mass balance). The recovery must be within 80% to 120% for data acceptance.

#### P-glycoprotein assay

P-glycoprotein (permeability-glycoprotein, abbreviated as “P-gp” also known as multidrug resistance protein 1 (MDR1) is the most studied and best characterized drug transporter^78^. The P-gp assay evaluates the ability of test compounds to serve as a P-gp substrate.

### Plasmids

Synthesis of CB2, vasopressin V2 receptor (V2R), TwinStrep and 1D4 tags was ordered (Genewiz). CB1 gene was from Addgene. SNAP-tag comes from pSNAPf-ADRβ_2_ control plasmid (NEB, #N9184). SNAP-CB2-TwinStrep-1D4, TwinStrep-SNAP-CB1 and TwinStrep-SNAP-V2R constructs (sequences in SI) were cloned into the multiple cloning site of the pcDNA4/TO vector from Invitrogen by PCR using the respective plasmids as templates, followed by fragment assembly using home-made Gibson assembly mix^79^. CB2 single-residue mutants F87A, V113A, F183A and W194A were generated from wild-type SNAP-CB2-TwinStrep-1D4 construct using two-fragment PCR approach with mutation-containing primers, followed by Gibson assembly, as described earlier^80^. Plasmids encoding β-arrestin1-RlucII^81^, β-arrestin2-RlucII^82^, human GRK2^83^, rGFP-CAAX^65^, rGFP-FYVE^65^, G_s_-RlucII^67^, Gβ_1_ ^64^, Gγ_2_-GFP10^64^, wild-type G proteins (G_i1_, G_i2_, G_i3_, G_oA_, G_oB_, G_z_, G_q_, G_11_, G_14_, G_15_, G_12_, G_13_) and effector proteins fused to RlucII (RabGAP-RlucII, p63-RlucII, p115-RlucII, PDZ-RhoGEF-RlucII)^66^ were previously described. For G protein activation assay, receptor and wild-type Gα subunit were combined with an appropriate RlucII-fused signaling effector (depending on Gα subunit) and rGFP-fused CAAX domain (embedded in plasma membrane), monitoring increase in BRET signal upon recruitment of signaling effector to the plasma membrane as a result of G protein activation. An exemption is activation of G_s_ protein where receptor was combined with RlucII-fused Gα subunit, wild-type Gβ_1_, and GFP10-fused Gγ_2_ subunit, monitoring decrease in BRET signal upon G protein heterotrimer dissociation. In β-arrestin recruitment assay, receptor was combined with RlucII-fused β-arrestin, wild-type GRK2, and rGFP-fused CAAX domain (embedded in plasma membrane), monitoring increase of BRET signal upon β-arrestin recruitment to the receptor in plasma membrane. In order to detect β-arrestin trafficking to the endosomes upon receptor internalized, rGFP-FYVE (embedded in endosomal membrane) was transfected instead of rGFP-CAAX.

### Transfection

For all experiments human embryonic kidney (HEK) 293SL cells^84^ were used and regularly tested for mycoplasma contamination (PCR Mycoplasma Detection kit, ABM, Canada). HEK293SL cells were maintained in Dulbecco’s Modified Eagle’s Medium (DMEM), 4.5 g/L glucose, with L-glutamine and phenol red, without sodium pyruvate (Wisent Inc., Canada), supplemented with 10% Newborn Calf Serum (heat-inactivated; Wisent Inc., Canada) and 1x Penicillin-Streptomycin (Wisent Inc., Canada) in adherent culture at 37°C and 5% CO_2_. Cells were dissociated using 0.05% Trypsin with 0.53 mM EDTA (Wisent Inc., Canada) and transiently co-transfected with receptor and biosensor DNA using 25 kDa linear polyethylenimin (PEI; Polysciences Inc., USA). PEI was dissolved in water as 1 mg/mL stock solution with pH of 6.5–7.5 and kept at −20°C. Once thawed, aliquots were not refrozen. For the transfection, PEI was diluted to 0.03 mg/mL with sterile PBS: 30 μg PEI in 100 μL total volume for a condition. It was vortexed and incubated at RT for 10 min. DNA was mixed and diluted to 0.01 mg/mL with PBS: 1 μg DNA per 100 μL total volume for a condition. If less than 1 μg receptor/biosensor DNA was needed per condition, it was topped up by sheared salmon sperm DNA (Invitrogen – Life Technologies Inc., Canada). Detailed amounts of DNA used for each biosensor can be found in Table S4, Table S5 and Table S6. Diluted DNA and PEI were combined 1:1 (100 μL + 100 μL), vortexed, and incubated at RT for 20 min. The mixture was added to a suspension of 240’000 HEK293SL cells in 1.2 mL of the cell growth medium, followed by the distribution into Cellstar® PS 96-well cell culture plates (Greiner Bio-One, Germany) at a density of 20’000 cells per well and incubation at 37°C, 5% CO_2_. The transfection protocol is illustrated in Figure S5. For experiments with CB2 mutants, HEK293SL cells were transfected in Dulbecco’s Modified Eagle’s Medium (DMEM), 4.5 g/L glucose, with L-glutamine, without phenol red and sodium pyruvate (Wisent Inc., Canada), supplemented with 10% Newborn Calf Serum (heat-inactivated; Wisent Inc., Canada) and 1x Penicillin-Streptomycin (Wisent Inc., Canada).

### BRET experiments

Coelenterazine 400a (Prolume Ltd., USA) was used as the luciferase substrate. It was solubilized as 1 mM stock solution in anhydrous ethanol and kept at −20°C. Ligands were solubilized in DMSO as 1, 10 or 50 mM stock solutions and kept at −20°C. BRET experiments are performed 2 days post-transfection. Transfection medium was aspirated from wells, cells were washed with 200 μL PBS (Wisent Inc., Canada) and 100 μL Tyrode’s buffer was added (137 mM NaCl, 1 mM mgCl_2_, 1 mM CaCl_2_, 0.9 mM KCl, 3.6 mM NaH_2_PO_4_, 11.9 mM NaHCO_3_, 25 mM HEPES, 5.55 mM D-glucose, pH = 7.4). Plates were kept at 37°C for at least 30 min before the measurement. Just before the experiment, ligand dilutions in DMSO and DBC working solution were prepared (25 μM DBC and 1% Pluronic F127 (Sigma, Canada) in Tyrode’s buffer). For the experiment, one plate at a time was taken out of the incubator, 1 μL of appropriate drug dilution in DMSO was added using an electronic multichannel pipette and mixed by moving the plate up-down-left-right for 5 s before returning the plate in the incubator. Five minutes before a BRET measurement, 10 μL of DBC working solution was added (final 2.5 μM) and the plate was returned to the incubator. BRET measurements were performed on a Biotek Synergy Neo plate reader at room temperature 10 minutes (G protein) or 20 minutes (β-arrestin) after ligand stimulation. RlucII signal was recorded using 410/80 nm filter, while rGFP signal was recorded using 515/30 nm filter, with the integration time of 0.4 s and a gain of 150 for both channels, read height being 4.5 mm. The BRET measurement protocol is illustrated in Figure S5. For experiments with CB2 mutants, cells were not washed with PBS, but transfection medium was immediately replaced with 100 μL Tyrode’s buffer. Ligand was added in 0.4 μL of appropriate dilution in DMSO and 10 μL of DBC working solution was added using a Multidrop Combi (Thermo Scientific) and shaken for 10 seconds (final 5 μM), followed by incubation at RT prior to BRET measurement.

### Cell surface ELISA measurements

Cell surface expression levels of CB2 mutants relative to the wild-type receptor were measured by ELISA as follows. HEK293SL cells were diluted to 200’000 cells/ml. For each receptor variant, 200 ng receptor plasmid were combined with 800 ng salmon sperm DNA and PEI at a 1:3 ratio of DNA to PEI, followed by vortexing. After 20 min incubation, 1.2 ml cells were added to each sample. The transfection mixture was plated into poly-D-lysine coated 96-well plates using 100 μl/well and four wells per condition. The cells were kept at 37°C 5% CO_2_ for 2 days. Each well was washed with 200 μl PBS and the cells were fixed with 50 μl 3% paraformaldehyde (PFA) per well for 10 min. The wells were washed three times with 100 μl/well washing buffer (PBS + 0.5% BSA), including last wash step with 10 min incubation period at RT. Per well, 50 μl of 0.25 μg/ml primary rabbit anti-SNAP antibody (GenScript, USA) was added and incubated for 1 h at RT, followed by washing as above. Then, 50 μl of a 1:1000 dilution of anti-rabbit HRP antibody (GE Healthcare) was added per well, followed by incubation at RT for 1 h. The cells were washed with washing buffer as described above, followed by three 10 min wash steps with PBS. SigmaFast solution (100 μl) was added to cells, followed by incubation at RT protected from light until color change was observed. To stop the reaction, 25 μl of 3M HCl were added to each well and 100 μl of the solution were transferred to a new transparent 96-well plate for the measurement. The absorbance at 492 nm was measured using a Tecan GENios Plus microplate reader, reporting on the amount of receptor expressed. Cell-surface ELISAs were carried out in three biological replicates with technical quadruplicates.

### Effect of the receptor expression level on maximal ligand-induced response

Receptor titrations were performed to test how E_max_ for each pathway/ligand changes with the amount of expressed wild-type CB2. Various amounts of CB2 DNA were transfected (0, 0.1, 0.5, 1, 2.5, 5, 10, 25, 50, 100, 150, 200% of the amount used for each biosensor), together with other biosensor components and the appropriate amount of ssDNA (to have 1 μg DNA in total) in duplicates. Two days post-transfection one replicate was used to perform ELISA, while the other replicate was used to measure BRET signal upon stimulation with 1 μM HU210 ligand, as described above. Change in BRET (ΔBRET) vs. amount of CB2 transfected (percentage of the amount used for each biosensor), and cell-surface expression level (determined by ELISA) vs. amount of CB2 transfected were measured. Correlation of ΔBRET and ELISA-measured expression was determined, and data fit equation parameters (linear regression or non-linear hyperbolic fit) were used to correct ΔBRET output of each mutant for its previously measured expression level.

### Data analysis

Ligand screen experiments were performed as three independent experiments in technical triplicates (CB1) or quadruplicates (CB2). Experiments with CB2 mutants were performed as three independent experiments, but without technical replicates. Nonlinear regression analysis of the concentration-response curves was done with GraphPad Prism 7.03 for Windows (GraphPad Software, USA). Concentration-response data were fitted into “Sigmoidal dose-response” (Hill Slope was constrained to 1) and are reported as mean ± standard error of mean (SE). For each pathway ligand-induced responses were normalized to WIN55212-2 for the given pathway, to allow comparison of ligand effects across different biosensors. In order to calculate bias factors, data were fitted into the equation for operational model of biased agonism and analyzed as described in van der Westhuizen *et al*.^68^, using WIN55252-2 as a reference compound in all the pathways, and results are reported as mean ± standard error of mean (SE). Bar charts and concentration-response charts were plotted using GraphPad Prism.

### Selection of CB2 mutants using Arpeggio

A CB2 homology model with HU210 ligand was generated with MOE (Molecular Operating Environment (MOE) 2019.01; Chemical Computing Group ULC, Montreal, Canada) applying default settings. The template was the active CB1 structure with bound THC (5XR8). Arpeggio server^69^ (http://biosig.unimelb.edu.au/arpeggioweb/) was used to analyze contacts formed between the ligand and receptor, selecting the residues that make the largest number of “specific” types of atom-atom interactions (F87, F94, F183, W194 and F281, Figure S2) such as π‒π stacking, hydrophobic‒van der Waals and weak polar‒van der Waals interactions. The selected residues were mutated into alanines one at a time, as described above.

