## Supplementary material for "Diverse chemotypes drive biased signaling by cannabinoid receptors": SI

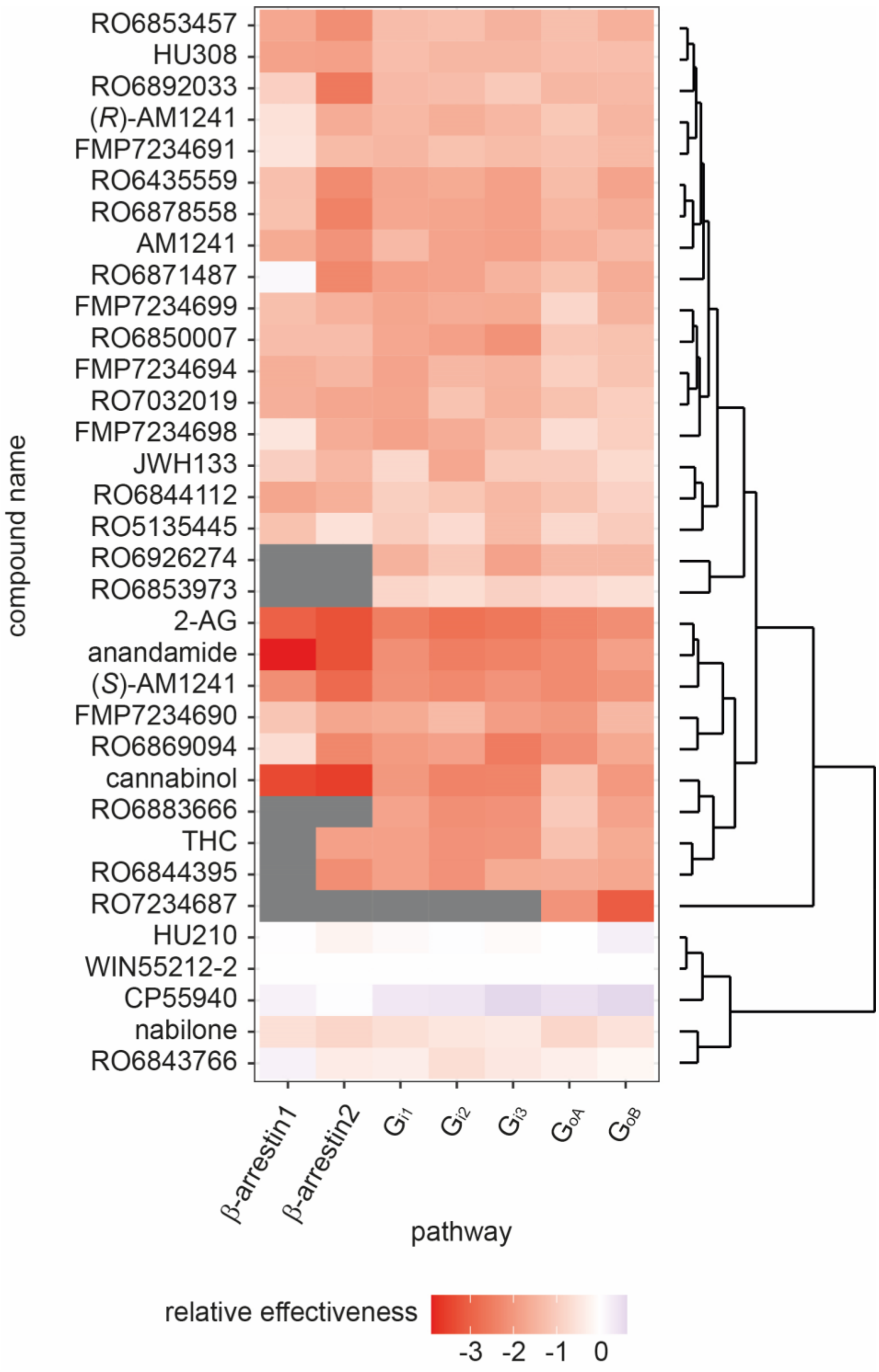

**Figure S1.** Ligand relative effectiveness (RE) on CB2 signaling represented as a heat map. Data from Table S16. For several ligands logR did not converge and RE could not be determined (grey). Three main clusters are observed, one having similar signaling properties as the reference ligand WIN55212-2, another with signaling reduced more or less uniform across the observed signaling pathways, and the third group where β-arrestin signaling was affected more strongly compared to G protein signaling.

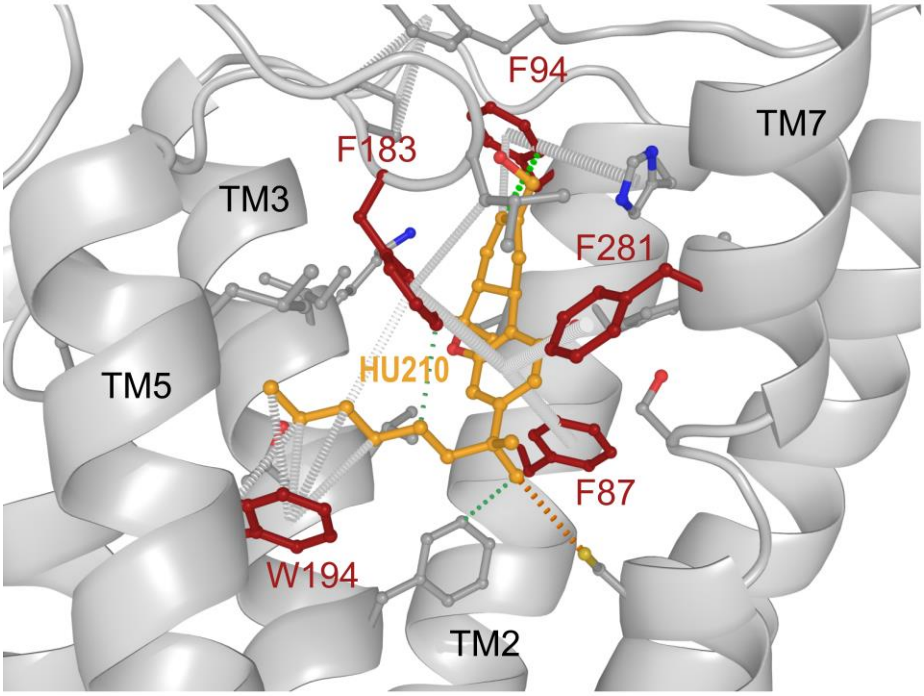

| residue | no. of interactions* |
|---|---|
| 25 | 0 |
| 87 | 16 |
| 90 | 0 |
| 91 | 0 |
| 94 | 9 |
| 95 | 1 |
| 109 | 0 |
| 110 | 3 |
| 113 | 5 |
| 114 | 2 |
| 117 | 6 |
| 182 | 1 |
| 183 | 22 |
| 184 | 1 |
| 190 | 1 |
| 191 | 1 |
| 194 | 16 |
| 261 | 1 |
| 265 | 5 |
| 281 | 7 |
| 282 | 0 |
| 285 | 1 |
| 288 | 2 |

**Figure S2.** Interactions between CB2 receptor and HU210 ligand, as calculated using Arpeggio server. HU210 is shown in orange sticks and CB2 residues forming the highest number of specific interactions with ligand in red. Specific interactions include pi-pi stacking (grey), hydrophobic-van der Waals (green) and weak polar-van der Waals (orange). Table shows all residues found to contact HU210, while in red are highlighted residues forming the highest number of specific interactions.

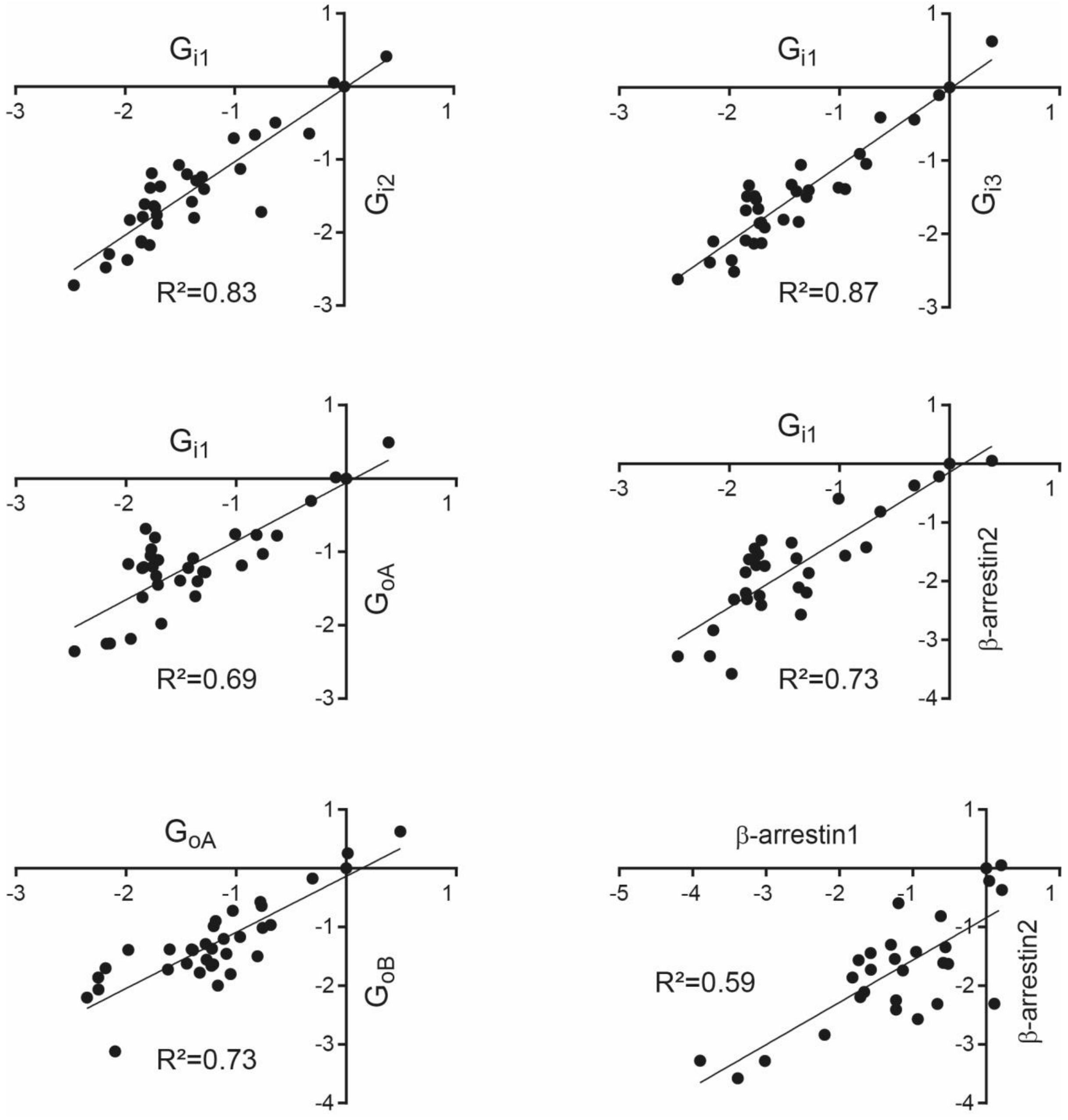

**Figure S3.** Correlation of ligand relative effectiveness (RE) data on CB2 signaling across the pathways. Data from Table S16. While strong correlation was observed among $G_i$ family members, somewhat reduced correlation was observed between $G_i$ vs. $G_o$, $G_i$ vs. β-arrestin and among $G_o$ family responses. The weakest correlation was observed between the two β-arrestin responses.

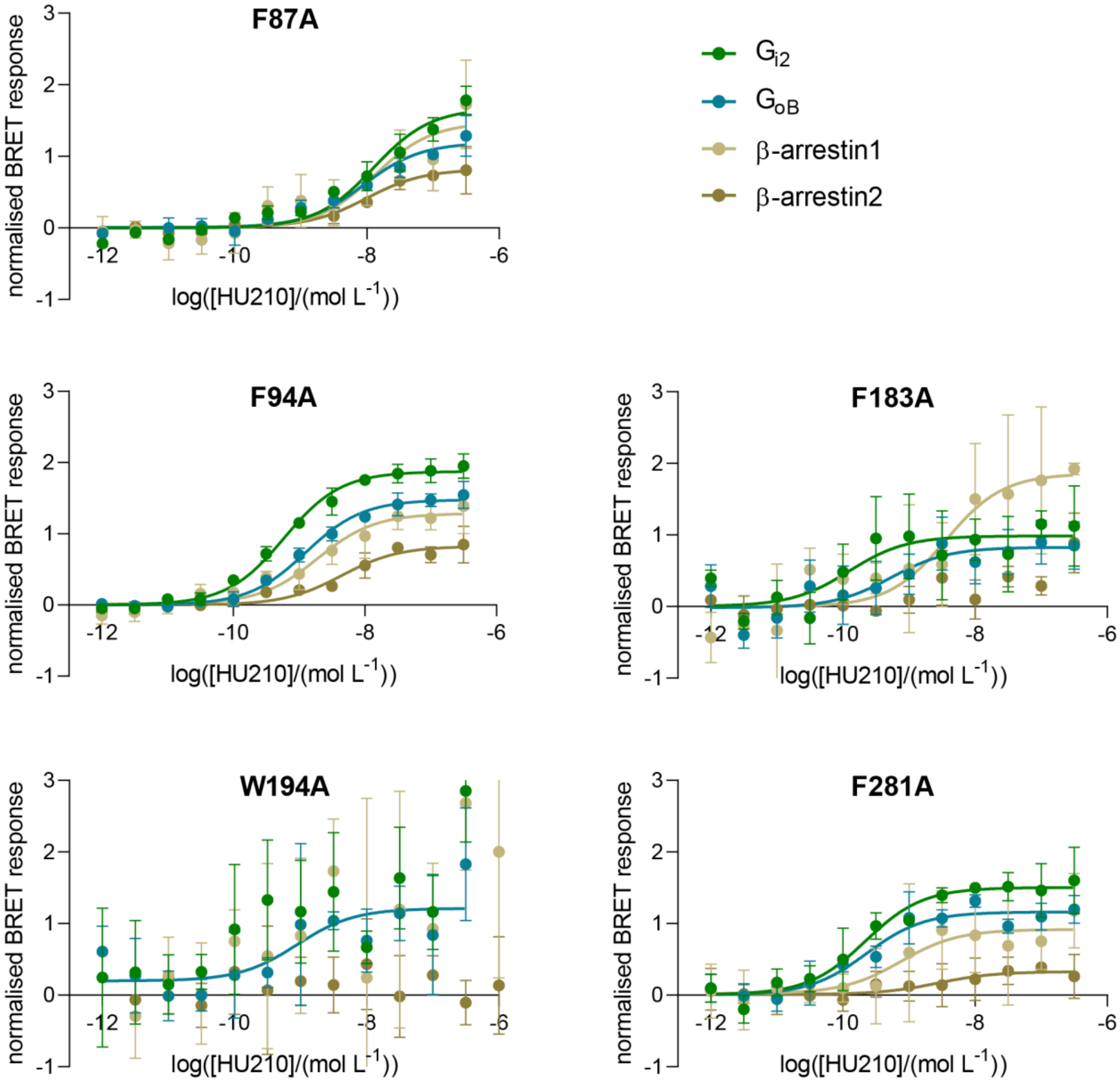

**Figure S4.** Concentration-dependent activation of G proteins and β-arrestins mediated by single amino acid CB2 mutants F87A, F94A, F183A, W194A and F281A upon stimulation with agonist HU210. Signal response was corrected for expression levels of the mutants and normalized to the wild type receptor response.

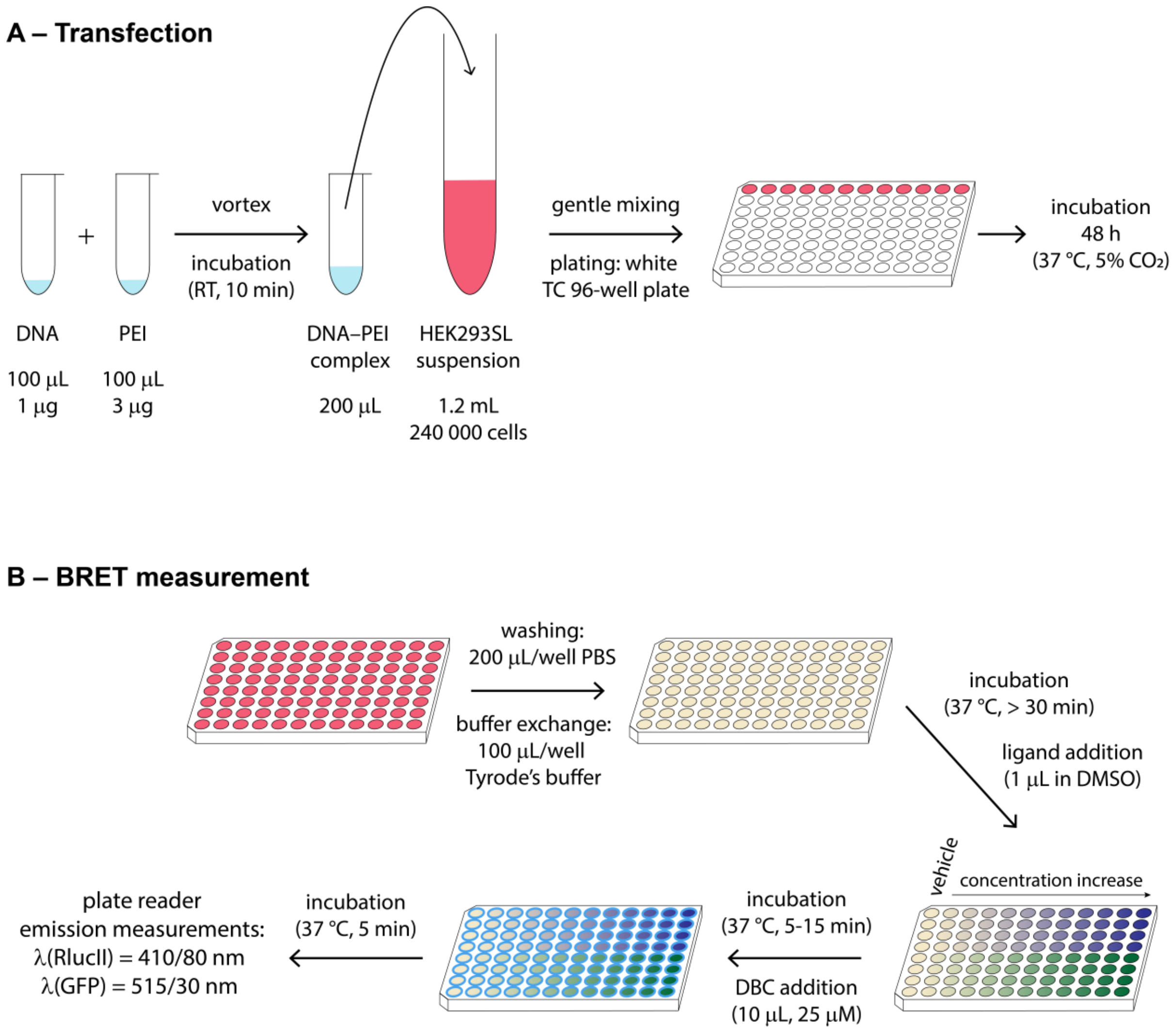

**Figure S5.** Schematic representation of transfection (A) and BRET measurement (B) protocols as described in Materials and Methods.

**Table S1.** Chemical structures, IUPAC names and CAS numbers of the cannabinoid receptor ligands used in the biased signaling studies.

| Compound | Chemical structure | IUPAC name | CAS number |
|---|---|---|---|
| SR144528 | 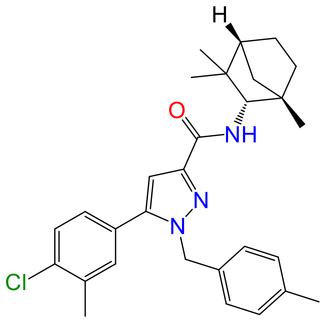 | 5-(4-Chloro-3-methylphenyl)-1-[(4-methylphenyl)methyl]-N-[(1S,2S,4R)-1,3,3-trimethylbicyclo[2.2.1]heptan-2-yl]-1H-pyrazole-3-carboxamide | 192703-06-3 |
| AM630 | 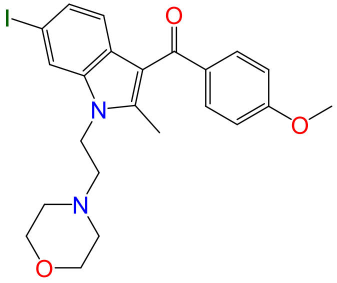 | {6-Iodo-2-methyl-1-[2-(morpholin-4-yl)ethyl]-1H-indol-3-yl}(4-methoxyphenyl)methanone | 164178-33-0 |
| THC | 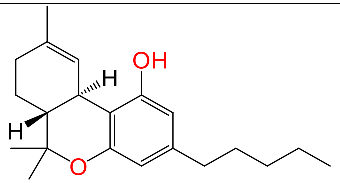 | (-)-(6aR,10aR)-6,6,9-Trimethyl-3-pentyl-6a,7,8,10a-tetrahydro-6H-dibenzo[b,d]pyran-1-ol | 1972-08-3 |

| | | | |
|---|---|---|---|
| cannabinol | 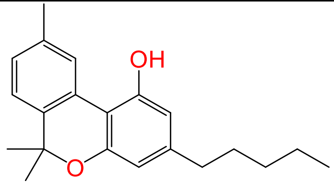 | 6,6,9-Trimethyl-3-pentyl-6H-dibenzo[b,d]pyran-1-ol | 521-35-7 |
| (*rac*)-AM1241 | 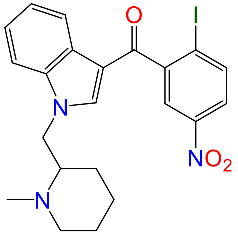 | (2-Iodo-5-nitrophenyl){1-[(1-methylpiperidin-2-yl)methyl]-1H-indol-3-yl}methanone | 444912-48-5 |
| (*R*)-AM1241 | 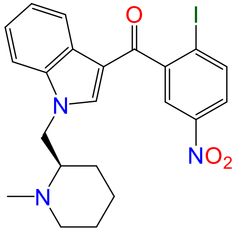 | (2-Iodo-5-nitrophenyl)(1-{[(2R)-1-methylpiperidin-2-yl]methyl}-1H-indol-3-yl)methanone | 444912-51-0 |
| (*S*)-AM1241 | 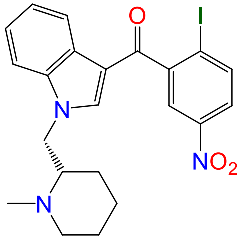 | (2-Iodo-5-nitrophenyl)(1-{[(2S)-1-methylpiperidin-2-yl]methyl}-1H-indol-3-yl)methanone | 444912-53-2 |

| | | | |
|---|---|---|---|
| anandamide | 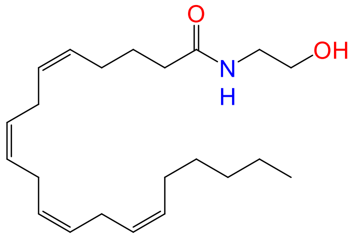 | (5Z,8Z,11Z,14Z)-N-(2-Hydroxyethyl)eicosa-5,8,11,14-tetraenamide | 94421-68-8 |
| HU210 | 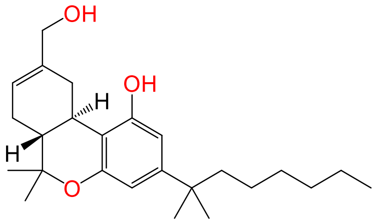 | (6aR,10aR)-9-(Hydroxymethyl)-6,6-dimethyl-3-(2-methyloctan-2-yl)-6a,7,10,10a-tetrahydro-6H-dibenzo[b,d]pyran-1-ol | 112830-95-2 |
| nabilone | 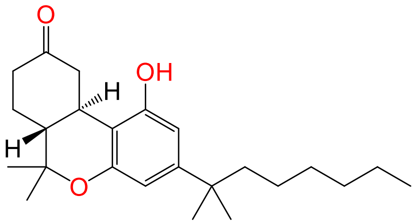 | (6aR,10aR)-1-Hydroxy-6,6-dimethyl-3-(2-methyloctan-2-yl)-6,6a,7,8,10,10a-hexahydro-9H-dibenzo[b,d]pyran-9-one | 51022-71-0 |
| HU308 | 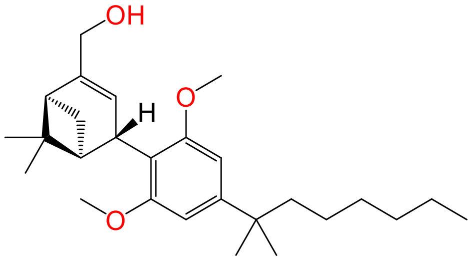 | {(1S,4S,5S)-4-[2,6-Dimethoxy-4-(2-methyloctan-2-yl)phenyl]-6,6-dimethylbicyclo[3.1.1]hept-2-en-2-yl}methanol | 256934-39-1 |

| | | | |
|---|---|---|---|
| WIN55212-2 | 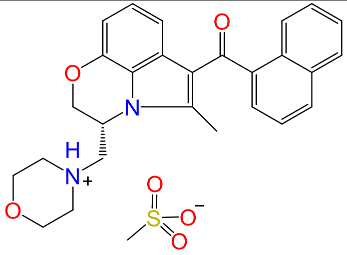 | Methanesulfonic acid--{(3R)-5-methyl-3-[(morpholin-4-yl)methyl]-2,3-dihydro[1,4]oxazino[2,3,4-hi]indol-6-yl}(naphthalen-1-yl)methanone (1/1) | 131543-23-2 |
| JWH133 | 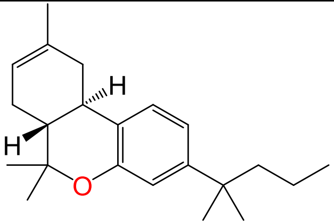 | (6aR,10aR)-6,6,9-Trimethyl-3-(2-methylpentan-2-yl)-6a,7,10,10a-tetrahydro-6H-dibenzo[b,d]pyran | 259869-55-1 |
| CP55940 | 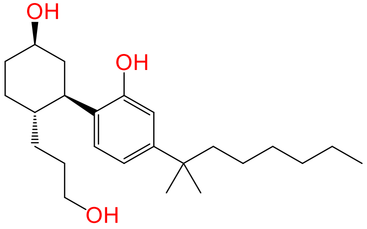 | 2-[(1R,2R,5R)-5-Hydroxy-2-(3-hydroxypropyl)cyclohexyl]-5-(2-methyloctan-2-yl)phenol | 83002-04-4 |
| 2-AG | 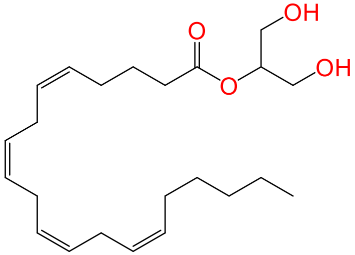 | 1,3-Dihydroxypropan-2-yl (5Z,8Z,11Z,14Z)-eicosa-5,8,11,14-tetraenoate | 53847-30-6 |

| | | | |
|---|---|---|---|
| RO6435559 | 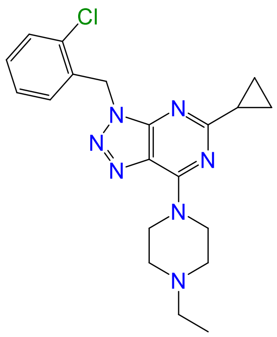 | 3-[(2-Chlorophenyl)methyl]-5-cyclopropyl-7-(4-ethylpiperazin-1-yl)-3H-[1,2,3]triazolo[4,5-d]pyrimidine | 841215-25-6 |
| RO6843766 | 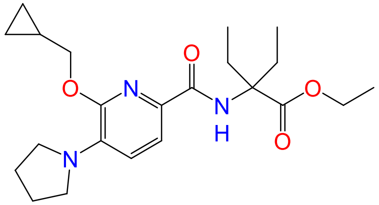 | Ethyl 2-{[6-(cyclopropylmethoxy)-5-(pyrrolidin-1-yl)pyridine-2-carbonyl]amino}-2-ethylbutanoate | 1415897-32-3 |
| RO6844112 | 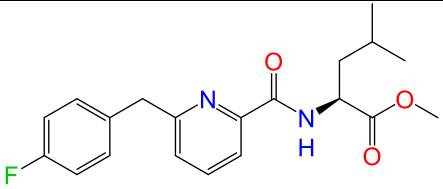 | Methyl N-{6-[(4-fluorophenyl)methyl]pyridine-2-carbonyl}-L-leucinate | Not existent |

| | | | |
|---|---|---|---|
| RO6844395 | 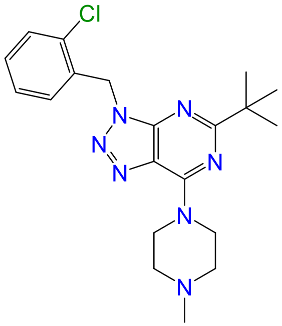 | 5-tert-Butyl-3-[(2-chlorophenyl)methyl]-7-(4-methylpiperazin-1-yl)-3H-[1,2,3]triazolo[4,5-d]pyrimidine | 1433356-80-9 |
| RO6850007 | 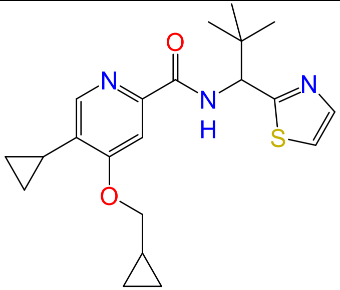 | 5-Cyclopropyl-4-(cyclopropylmethoxy)-N-[2,2-dimethyl-1-(1,3-thiazol-2-yl)propyl]pyridine-2-carboxamide | 1613235-74-7 |
| RO6853457 | 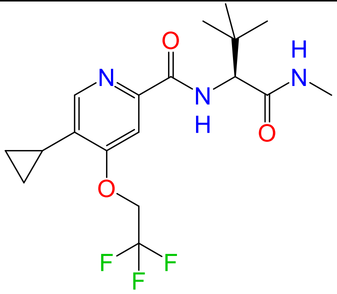 | 5-Cyclopropyl-N-[(2S)-3,3-dimethyl-1-(methylamino)-1-oxobutan-2-yl]-4-(2,2,2-trifluoroethoxy)pyridine-2-carboxamide | 1613235-79-2 |

| | | | |
|---|---|---|---|
| RO6853973 | 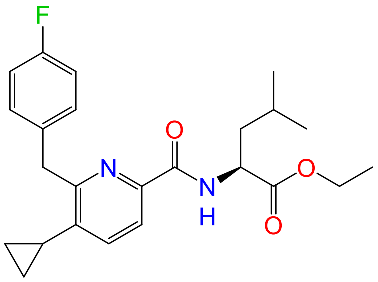 | Ethyl N-{5-cyclopropyl-6-[(4-fluorophenyl)methyl]pyridine-2-carbonyl}-L-leucinate | 1415900-27-4 |
| RO6869094 | 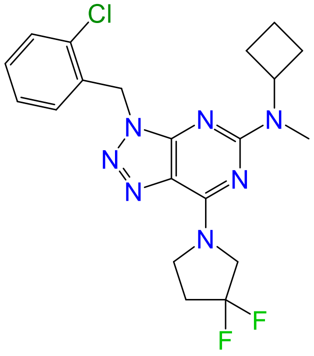 | 3-[(2-Chlorophenyl)methyl]-N-cyclobutyl-7-(3,3-difluoropyrrolidin-1-yl)-N-methyl-3H-[1,2,3]triazolo[4,5-d]pyrimidin-5-amine | 1672656-05-1 |
| RO6871487 | 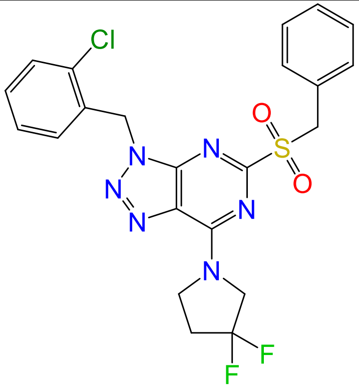 | 3-[(2-Chlorophenyl)methyl]-7-(3,3-difluoropyrrolidin-1-yl)-5-(phenylmethanesulfonyl)-3H-[1,2,3]triazolo[4,5-d]pyrimidine | 1672656-74-4 |

| | | | |
|---|---|---|---|
| RO6878558 | 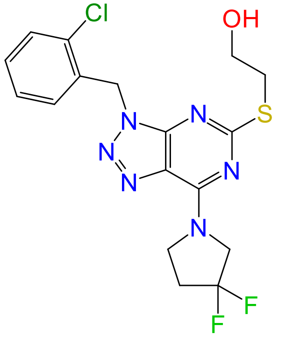 | 2-({3-[(2-Chlorophenyl)methyl]-7-(3,3-difluoropyrrolidin-1-yl)-3H-[1,2,3]triazolo[4,5-d]pyrimidin-5-yl}sulfanyl)ethan-1-ol | 1672656-76-6 |
| RO5135445 | 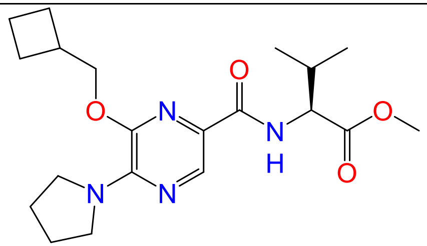 | Methyl N-[6-(cyclobutylmethoxy)-5-(pyrrolidin-1-yl)pyrazine-2-carbonyl]-L-valinate | Not existent |
| RO6883666 | 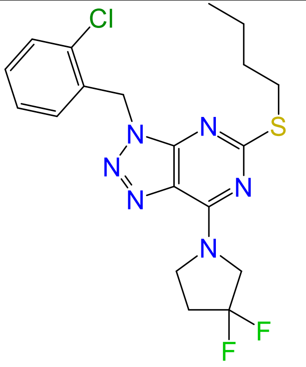 | 5-(Butylsulfanyl)-3-[(2-chlorophenyl)methyl]-7-(3,3-difluoropyrrolidin-1-yl)-3H-[1,2,3]triazolo[4,5-d]pyrimidine | 1672656-78-8 |

| | | | |
|---|---|---|---|
| RO6892033 | 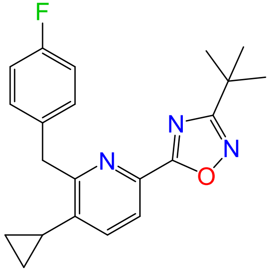 | 6-(3-tert-Butyl-1,2,4-oxadiazol-5-yl)-3-cyclopropyl-2-[(4-fluorophenyl)methyl]pyridine | Not existent |
| RO6926274 | 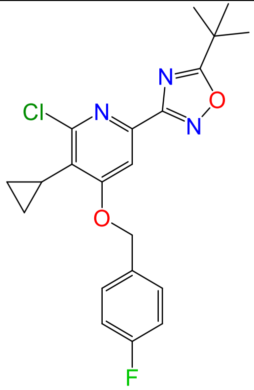 | 6-(5-tert-Butyl-1,2,4-oxadiazol-3-yl)-2-chloro-3-cyclopropyl-4-[(4-fluorophenyl)methoxy]pyridine | 1629991-47-4 |
| RO7032019 | 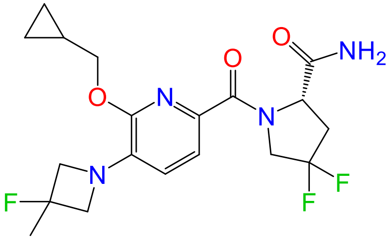 | 1-[6-(Cyclopropylmethoxy)-5-(3-fluoro-3-methylazetidin-1-yl)pyridine-2-carbonyl]-4,4-difluoro-L-prolinamide | 1812888-93-9 |

| | | | |
|---|---|---|---|
| FMP7234690 | 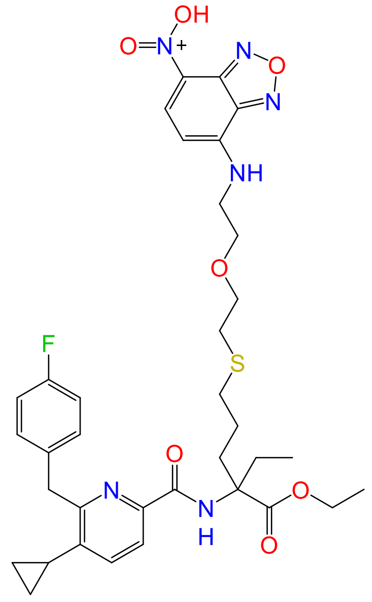 | Ethyl N-{5-cyclopropyl-6-[(4-fluorophenyl)methyl]pyridine-2-carbonyl}-2-ethyl-5-[(2-{2-[(7-nitro-2,1,3-benzoxadiazol-4-yl)amino]ethoxy}ethyl)sulfanyl]norvalinate | Not existent |

| | | | |
|---|---|---|---|
| FMP7234691 | 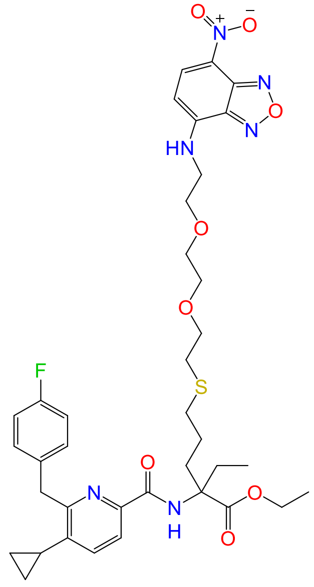 | Ethyl N-{5-cyclopropyl-6-[(4-fluorophenyl)methyl]pyridine-2-carbonyl}-2-ethyl-5-{[2-(2-{2-[(7-nitro-2,1,3-benzoxadiazol-4-yl)amino]ethoxy}ethoxy)ethyl]sulfanyl}norvalinate | Not existent |

| | | | |
|---|---|---|---|
| FMP7234694 | 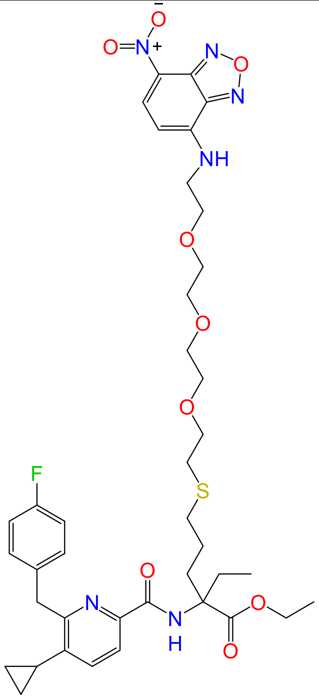 | Ethyl N-{5-cyclopropyl-6-[(4-fluorophenyl)methyl]pyridine-2-carbonyl}-2-ethyl-5-({2-[2-(2-{2-[(7-nitro-2,1,3-benzoxadiazol-4-yl)amino]ethoxy}ethoxy)ethoxy]ethyl}sulfanyl)norvalinate | Not existent |

| | | | |
|---|---|---|---|
| FMP7234698 | 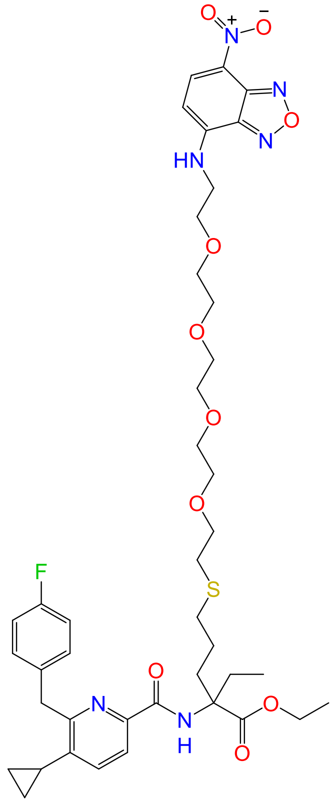 | Ethyl N-{5-cyclopropyl-6-[(4-fluorophenyl)methyl]pyridine-2-carbonyl}-2-ethyl-5-({14-[(7-nitro-2,1,3-benzoxadiazol-4-yl)amino]-3,6,9,12-tetraoxatetradecan-1-yl}sulfanyl)norvalinate | Not existent |
| FMP7234699 | | Ethyl N-{5-cyclopropyl-6-[(4-fluorophenyl)methyl]pyridine-2-carbonyl}-2-ethyl-5-({6-[(7-nitro-2,1,3-benzoxadiazol-4-yl)amino]hexyl}sulfanyl)norvalinate | Not existent |

| | |
|---|---|
| | 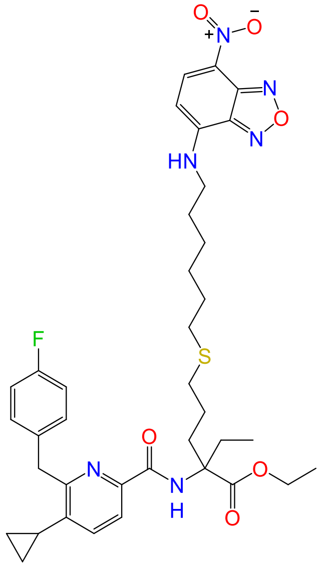 |

**Table S2.** In vitro pharmacology data for cannabinoid receptor ligands. CB2 selectivity calculated as $10^{pKi(CB2)-pKi(CB1)}$.

| Compound | $pK_i$ ± SE (hCB2) | $pK_i$ ± SE (hCB1) | $K_i$ (hCB1) / $K_i$ (hCB2) | $pK_i$ ± SE (mCB2) | cAMP pEC50 ± SE (hCB2) | cAMP $E_{max}$ ± SE (hCB2) | cAMP pEC50 ± SE (hCB1) | cAMP $E_{max}$ ± SE (hCB1) | cAMP pEC50 (hCB1) / cAMP pEC50 (hCB2) | cAMP pEC50 ± SE (mCB2) | cAMP $E_{max}$ ± SE (mCB2) | β-arrestin pEC50 ± SE (hCB2) | β-arrestin $E_{max}$ ± SE (hCB2) | β-arrestin pEC50 ± SE (hCB1) | β-arrestin $E_{max}$ ± SE (hCB1) | β-arrestin pEC50 (hCB1) / β-arrestin pEC50 (hCB2) | β-arrestin pEC50 ± SE (mCB2) | β-arrestin $E_{max}$ ± SE (mCB2) |
|---|---|---|---|---|---|---|---|---|---|---|---|---|---|---|---|---|---|---|
| SR144528 | *7.88 ± 0.06* | *5.77 ± 0.09* | *129* | *10.7 ± 0.16* | *7.67 ± 0.18* | *-151 ± 10* | *< 5* | *n.a.* | *> 468* | *8.25 ± 0.29* | *-100* | *7.47 ± 0.27* | *-111 ± 3* | *4.97 ± 0.13* | *-82 ± 5* | *316* | < 4.5 | -2 |
| AM630 | *7.39 ± 0.02* | *6.51 ± 0.25* | *8* | *7.66 ± 0.23* | *7.56 ± 0.05* | *-152 ± 13* | *< 5* | *28 ± 5* | *> 363* | *7.93 ± 0.3* | *-65* | *6.15 ± 0.08* | *-124 ± 6* | *5.56 ± 0.1* | *-120 ± 25* | *4* | < 4.5 | -1 |
| THC | *8.16 ± 0.17* | *8.48 ± 0.08* | *0.5* | *ND* | *ND* | *ND* | *ND* | *ND* | *ND* | *ND* | *-* | *-* | *31 ± 3§* | *6.29 ± 0.03* | *35 ± 3* | *-* | ND | ND |

| | | | | | | | | | | | | | | | | | | |
|---|---|---|---|---|---|---|---|---|---|---|---|---|---|---|---|---|---|---|
| cannabinol | 5.49 ± 0.01 | 6 ± 0.10 | 0.4 | 6.00 ± 0.05## | 7.09 ± 0.07## | 70 | < 5 | n.a. | > 123 | 7.59 ± 0.13 | 83 | ND | ND | ND | ND | ND | ND | ND |
| (rac)-AM1241 | *8.39 ± 0.10* | *6.89 ± 0.31* | *32* | *7.73 ± 0.25* | *8.49± 0.08* | *67 ± 3* | *< 5* | *n.a.* | *3103* | *7.03 ± 0.26* | *-6* | *< 5* | *14 ± 2* | *< 5* | *6 ± 5* | *1* | ND | ND |
| (*R*)-AM1241 | *8.44 ± 0.25* | *6.86 ± 0.46* | *38* | *7.59 ± 0.11* | *8.76 ± 0.18* | *65 ± 2* | *6.17 ± 0.12* | *54 ± 7* | *391* | *6.17 ± 0.23* | *-22* | *7.52 ± 0.43* | *29 ± 4* | *5.59 ± 0.13* | *26 ± 2* | *86* | ND | ND |
| (*S*)-AM1241 | *7.02 ± 0.15* | *< 5* | *105* | *6.74 ± 0.2* | *7.40 ± 0.11* | *78 ± 1* | *< 5* | *16 ± 2* | *> 250* | *7.6 ± 0.3* | *35* | *6.54 ± 0.44* | *29 ± 8* | *< 5* | *2 ± 1* | *35* | ND | ND |
| anandamide | *6.91 ± 0.28* | *7.04 ± 0.28#* | *1* | *6.46 ± 0.18* | *5.93 ± 0.29* | *87± 7* | *< 5* | *39 ± 3* | *> 8* | *6.82 ± 0.51* | *48* | *6.21 ± 0.63* | *38 ± 3* | - | *36 ± 13§* | - | ND | ND |
| HU210 | *9.78 ± 0.04* | *9.55 ± 0.06* | *2* | *9.27 ± 0.38* | *9.45 ± 0.27* | *94 ± 0.1* | *9.58 ± 0.24* | *104 ± 3* | *1* | *ND* | *ND* | *ND* | *ND* | *ND* | *ND* | *ND* | ND | ND |
| nabilone | 8.09 ± 0.04## | 8.75 ± 0.08## | 0.2 | 7.72 ± 0.04## | 9.05 ± 0.24## | 100 | 8.24 ± 0.11## | 107 | 6 | 8.55 ± 0.07## | 103 | ND | ND | ND | ND | ND | ND | ND |

| | | | | | | | | | | | | | | | | | | |
|---|---|---|---|---|---|---|---|---|---|---|---|---|---|---|---|---|---|---|
| HU308 | *7.44 ± 0.12* | *< 5* | *278* | *7.15 ± 0.21* | *8.53 ± 0.06* | *98 ± 1* | *< 5* | *18 ± 4* | *> 3388* | *8.43 ± 0.47* | *56* | *7.45 ± 0.07* | *57 ± 10* | *< 5* | *5.0 ± 1* | *282* | *6.47 ± 0.13* | *69 ± 3* |
| WIN55212-2 | *8.57 ± 0.16* | *8.72 ± 0.24* | *1* | *7.28 ± 0.17* | *9.50 ± 0.09* | *98 ± 1* | *7.86 ± 0.23* | *107 ± 1* | *44* | *8.37 ± 0.23* | *97* | *7.93 ± 0.42* | *55 ± 5* | *6.74 ± 0.21* | *89 ± 2* | *15* | < 5 | -0.3 |
| JWH133 | *7.18 ± 0.34* | *< 5* | *153* | *7.69 ± 0.23* | *8.38 ± 0.09* | *98 ± 1* | *< 5* | *37 ± 6* | *> 2399* | *7.75 ± 0.33* | *95* | *7.81 ± 0.22* | *62 ± 11* | *5.37 ± 0.05* | *40 ± 7* | *275* | *6.80 ± 0.17* | *63 ± 7* |
| CP55940 | *8.44 ± 0.18* | *9.26 ± 0.12* | *0.2* | *9.22 ± 0.26* | *10.33 ± 0.09* | *98 ± 1* | *9.73 ± 0.10* | *100 ± 3* | *4* | *10.17 ± 0.3* | *100* | *8.39 ± 0.21* | *80 ± 5* | *7.96 ± 0.2* | *94 ± 2* | *3* | *8.02 ± 0.27* | *96 ± 1* |
| 2-AG | *6.94 ± 0.43* | *7.15 ± 0.47#* | *1* | *7.53 ± 0.22* | *6.82 ± 0.05* | *94 ± 1* | *< 5* | *n.a.* | *> 67* | *7.15 ± 0.4* | *53* | *5.70 ± 0.21* | *80 ± 21* | - | *51 ± 21§* | - | ND | ND |
| RO6435559 | 8.05 ± 0.09## | 6.08 ± 0.09## | 93 | 7.48 ± 0.09## | 8.27 ± 0.10 | 77 ± 9 | 5.81 ± 0.28## | 79 | 288 | <5 | -13 | ND | ND | ND | ND | - | ND | ND |
| RO6843766 | 8.83 ± 0.05## | 6.21 ± 0.03## | 416 | 8.55 ± 0.03## | 8.85 ± 0.15## | 91 | 6.63 ± 0.16## | 59 | 165 | 7.92 ± 0.05## | 61 | ND | ND | ND | ND | - | ND | ND |

| | | | | | | | | | | | | | | | | | | |
|---|---|---|---|---|---|---|---|---|---|---|---|---|---|---|---|---|---|---|
| RO6844112 | 7.82 ± 0.06$^{\#\#}$ | 7.31 ± 0.08$^{\#\#}$ | 3 | 7.16 ± 0.03$^{\#\#}$ | 7.99 ± 0.02$^{\#\#}$ | 92 | 7.14 ± 0.07$^{\#\#}$ | 101 | 7 | 8.03 ± 0.16$^{\#\#}$ | 58 | ND | ND | ND | ND | - | ND | ND |
| RO6844395 | 7.67 ± 0.04$^{\#\#}$ | 5.76 ± 0.05$^{\#\#}$ | 81 | 6.87 ± 0.05$^{\#\#}$ | 8.85 ± 0.22$^{\#\#}$ | 99 | 6.72 ± 0.17$^{\#\#}$ | 57 | 134 | 7.30 ± 0.05$^{\#\#}$ | 80 | ND | ND | ND | ND | - | ND | ND |
| RO6850007 | 7.65 ± 0.21 | 5.83 ± 0.07$^{\#\#}$ | 66 | 9.21 ± 0.09$^{\#\#}$ | 8.49 ± 0.06$^{\#\#}$ | 63 | <5 | 42 | > 3090 | 8.23 ± 0.03$^{\#\#}$ | -48 | ND | ND | ND | ND | - | ND | ND |
| RO6853457 | 8.12 ± 0.22 | 5.99 ± 0.23 | 134 | 8.46 ± 0.27 | 8.81 ± 0.12 | 74 ± 2.1 | 6.18 ± 0.13$^{\#\#}$ | 75 | 426 | 8.42 ± 0.18 | 48 ± 8.3 | ND | ND | 5.47 ± 0.05$^{\#\#}$ | 3 | >0.3 | <4.5 | -1 |
| RO6853973 | 9.45 ± 0.16$^{\#\#}$ | 8.21 ± 0.09 | 17 | 9.17 ± 0.18 | 8.73 ± 0.12$^{\#\#}$ | 45 | 8.12 ± 0.05$^{\#\#}$ | 83 | 4 | <5 | 23 | ND | ND | ND | ND | - | ND | ND |
| RO6869094 | 7.71 ± 0.08$^{\#\#}$ | 5.96 ± 0.04$^{\#\#}$ | 56 | 8.39 ± 0.08$^{\#\#}$ | 8.09 ± 0.12$^{\#\#}$ | 48 | 6.01 ± 0.11$^{\#\#}$ | 60 | 120 | <5 | 28 | ND | ND | ND | ND | - | ND | ND |
| RO6871487 | 7.98 ± 0.11$^{\#\#}$ | 5.34 ± 0.10$^{\#\#}$ | 436 | 8.16 ± 0.08$^{\#\#}$ | 7.08 ± 0.13$^{\#\#}$ | 52 | <5 | 9 | > 120 | <5 | 24 | ND | ND | ND | ND | - | ND | ND |

| | | | | | | | | | | | | | | | | | | |
|---|---|---|---|---|---|---|---|---|---|---|---|---|---|---|---|---|---|---|
| RO6878558 | 6.94 ± 0.04[##] | <5 | > 87 | 6.69 ± 0.04[##] | 8.13 ± 0.03[##] | 87 | <5 | 20 | > 1348 | 8.40 ± 0.04[##] | 92 | ND | ND | ND | ND | - | ND | ND |
| RO5135445 | 8.97 ± 0.35 | 6.93 ± 0.08 | 109 | 7.06 ± 0.05[##] | 8.63 ± 0.08 | 75 ± 9 | 6.36 | 74 | 186 | 7.89 ± 0.11[##] | 64 | ND | ND | ND | ND | - | ND | ND |
| RO6883666 | 7.41 ± 0.06[##] | 6.00 ± 0.44[##] | 25 | 7.77 ± 0.03[##] | 8.16 ± 0.09[##] | 67 | <5 | 44 | > 1445 | <5 | 2 | ND | ND | ND | ND | - | ND | ND |
| RO6892033 | 7.59 ± 0.06[##] | 6.20 ± 0.06[##] | 24 | 8.12 ± 0.14[##] | 8.33 ± 0.10[##] | 66 | 6.56 ± 0.08[##] | 70 | 58 | 8.39 ± 0.32[##] | 45 | ND | ND | ND | ND | - | ND | ND |
| RO6926274 | 8.15 ± 0.12[##] | 6.68 ± 0.05 | 29 | 7.40 ± 0.06[##] | 8.32 ± 0.28[##] | 38 | 7.14 ± 0.04[##] | 73 | 15 | 6.29 ± 0.11 | -160 | ND | ND | ND | ND | - | ND | ND |
| RO7032019 | 7.38 ± 0.20[##] | <5 | > 239 | 6.86 ± 0.07[##] | 7.96 ± 0.08[##] | -92 | <5 | 27 | > 912 | 7.55 ± 0.09[##] | -87 | ND | ND | ND | ND | - | ND | ND |
| FMP7234690 | 7.66 ± 0.09 | 6.38 ± 0.07 | 19 | 7.23 ± 0.08[##] | 6.77 ± 0.19[##] | 121 | 5.69 ± 0.35[##] | 152 | 12 | 6.74 ± 0.20[##] | 118 | ND | ND | ND | ND | - | ND | ND |

| | | | | | | | | | | | | | | | | | | |
|---|---|---|---|---|---|---|---|---|---|---|---|---|---|---|---|---|---|---|
| FMP7234691 | 7.65 ± 0.13 | 6.42 ± 0.14 | 16 | 7.14 ± 0.02[##] | 8.33 ± 1.40 | 100 ± 1 | 6.22 ± 0.01 | 126 ± 5 | 128 | 6.99 ± 0.27[##] | 118 | ND | ND | ND | ND | - | ND | ND |
| FMP7234694 | 7.57 ± 0.13 | 6.34 ± 0.06 | 16 | 7.07 ± 0.05[##] | 8.04 ± 0.28[##] | 100 | 6.10 ± 0.14[##] | 96 | 87 | 7.13 ± 0.16[##] | 113 | ND | ND | ND | ND | - | ND | ND |
| FMP7234698 | 7.42 ± 0.07 | 6.21 ± 0.06 | 16 | 6.93 ± 0.03[##] | 7.69 ± 0.13[##] | 102 | 5.95 ± 0.21[##] | 110 | 54 | 6.72 ± 0.15[##] | 117 | ND | ND | ND | ND | - | ND | ND |
| FMP7234699 | 7.26 ± 0.03 | 5.97 ± 0.07 | 19 | 6.84 ± 0.05[##] | 8.00 ± 0.14[##] | 100 | 6.01 ± 0.10[##] | 122 | 97 | 6.85 ± 0.16[##] | 122 | ND | ND | ND | ND | - | ND | ND |

cAMP assays: the effect of 10 µM agonists is normalized to the effect of 10 µM CP55940, the potency of antagonists/inverse agonists is determined in presence of the $EC_{80}$ of CP55940.
β-arrestin recruitment assays: the effect of 10 µM agonists is normalized to the effect of 10 µM CP55940, the potency of antagonists/inverse agonists is determined in presence of the $EC_{80}$ of CP55940.
Negative values represent inhibition of the $EC_{80}$ of CP55940 (> -100, indication of inverse agonism); n.a.: not active; ND = not determined. Effect at 10 µM.
[#] Mean ± SEM of 4 independent experiments.
[##] pKi values and ΔpKi errors were calculated from archived $IC_{50}$ and standard error of the fit data using the Cheng-Prusoff equation for competitive binding and error propagation. $pEC_{50}$ values and $\Delta pEC_{50}$ errors were calculated from archived $EC_{50}$ and standard error of the fit data using error propagation.
§ $E_{max}$ at 10 µM treatment, no plateau observed.
*Italicized values* described in Soethoudt *et al. Nat. Commun.* (2017) **8**: 13958.

**Table S3.** Calculated and measured physicochemical and early ADME properties with relevance for good *in vivo* performance of CB2 ligands. Green values from [1].

| compound | MW (g/mol) | PSA[a] (Å²) | HBD | $K_{ow}$ clogP[b] | Log*D*[c] (at pH 7.4) | p$K_a$ | kinetic solubility[d] (µg/mL) | PAMPA[e] $P_{eff}$ ($10^{-6}$ cm/s), %Acc./%Mem./%Don. | micros. CL human/mouse/rat (µL/min/kg) | hepat. CL human/mouse/rat (µL/min/million cells) | plasma protein binding: free fraction (%) human/mouse/rat | P-gp mediated efflux extraction ratio human/mouse |
|---|---|---|---|---|---|---|---|---|---|---|---|---|
| SR144528 | 476.06 | 37.5 | 1 | 9.2 | > 3 | - | <0.2 | 0, 0 / 54 / 47 | 129 / 446 / 520 | ND / ND / 8 | <0.2 / ND / < 0.2 | 4.1 / 5.6 |
| AM630 | 504.36 | 39.8 | 0 | 4.9 | 3.8 | 5.8 (B, calc.) | < 0.3 | 0.2, 0.4 / 64 / 35 | 82 / 468 / ND | 18 / 101 / ND | <0.1 / 0.4 / ND | 1.3 / 1.3 |
| THC | 314.47 | 22.2 | 1 | 7.6 | ND | - | ND | ND | ND | ND | ND | ND |
| cannabinol | 310.43 | 22.6 | 1 | 7.2 | ND | 9.8 (A, calc.) | ND | ND | ND | ND | ND | ND |
| (*rac*)-AM1241 | 503.33 | 62.5 | 0 | 5.7 | 3.66 ± 0.03 | 7.7 (B, calc.) | 25 ± 2 | 1.86 ± 0.62, 3 / 56 / 41 | 111 / 254 / ND | ND | ND | ND |
| (*R*)-AM1241 | 503.33 | 62.6 | 0 | 5.7 | 3.66 | 7.8 (B) | 12 | 1.22, 2 / 60 / 38 | 304 / 314 / ND | ND | ND | ND |
| (*S*)-AM1241 | 503.33 | 62.6 | 0 | 5.7 | 3.64 | 7.8 (B) | 6 | 1.41, 2 / 64 / 34 | 152 / 640 / ND | 34 / 251 / ND | 0.5 / 1.2 / ND | 2.7 / 1.4 |
| anandamide | 347.54 | 41.9 | 2 | 6.3 | out of range | - | <1.1 | 0.26, 1 / 3 / 96 | 47 / 222 / ND | ND | ND | ND |
| HU210 | 386.57 | 38.0 | 2 | 8.0 | prec. | - | <0.2 | 0.22, 0 / 76 / 23 | 25 / 33 / ND | ND | ND | ND |
| nabilone | 372.55 | 37.9 | 1 | 7.1 | ND | - | ND | ND | ND | ND | ND | ND |
| HU308 | 414.63 | 27.6 | 1 | 9.0 | 4.29 | - | <0.8 | 2.53, 3 / 74 / 23 | ND | 8 / 5 / ND | ND | 6.2 / 7.3 |
| WIN55912-2 | 426.51 | 36.3 | 0 | 4.7 | 3.67 ± 0.03 | 5.5 (B, calc.) | <0.3 | 0.25 ± 0.22, 1 / 49 / 51 | 194 / 745 / ND | 42 / 568 / ND | <0.3 / 1.0 / ND | 1.5 / 1.4 |
| JWH133 | 312.49 | 7.9 | 0 | 8.5 | > 3 prec. | - | < 0.4 | 1.86, 2 / 62 / 36 | 38 / 63 / ND | 6 / 10 / ND | <0.1 / <0.1 / <3.5 | ND / 1.7 |
| CP55940 | 376.58 | 49.9 | 3 | 7.5 | ND | 9.7 (A, calc.) | 1.4 | 0, 0 / 47 / 53 | 80 / 745 / ND | ND | ND | ND |
| 2-AG | 378.55 | 51.3 | 2 | 6.7 | ND | - | < 0.5 | ND | 13 / 12 / ND | ND | ND | ND |
| RO6435559 | 397.91 | 55.0 | 0 | 4.0 | ND | 8.0 (B, calc.) | ND | ND | ND | ND | ND | ND |
| RO6843766 | 403.52 | 64.1 | 1 | 5.4 | ND | 2.4 (B, calc.) | ND | ND | ND | ND | ND | ND |
| RO6844112 | 358.41 | 55.6 | 1 | 4.5 | ND | 2.6 (B, calc.) | ND | ND | ND | ND | ND | ND |
| RO6844395 | 399.93 | 54.2 | 0 | 4.1 | 3.22 | 7.9 (B, calc.) | 8 | 0.4, 1 / 45 / 54 | 27 / 491 / ND | ND | ND | ND |
| RO6850007 | 385.53 | 49.5 | 1 | 5.9 | ND | 11.1 (A, calc.), 3.7 and 2.7 (B, calc.) | ND | ND | ND | ND | ND | ND |

| | | | | | | | | | | | | |
|---|---|---|---|---|---|---|---|---|---|---|---|---|
| RO6853457 | 387.40 | 64.9 | 2 | 3.2 | 3.89 | 2.6 (B, calc.) | 92 | 3.9, 12 / 41 / 47 | 14 / 28 / 28 | ND / 85 / ND | ND | ND |
| RO6853973 | 412.50 | 53.2 | 1 | 6.3 | ND | 2.2 (B, calc.) | ND | ND | ND | ND | ND | ND |
| RO6869094 | 433.89 | 52.7 | 0 | 5.6 | ND | 2.4 (B, calc.) | ND | ND | ND | ND | ND | ND |
| RO6871487 | 504.95 | 84.9 | 0 | 4.6 | ND | - | ND | ND | ND | ND | ND | ND |
| RO6878558 | 426.88 | 69.8 | 1 | 3.4 | ND | - | 1 | ND | 56 / 174 | ND | ND | ND |
| RO5135445 | 390.48 | 74.4 | 1 | 5.5 | ND | 3.6 (B, calc.) | ND | ND | ND | ND | ND | ND |
| RO6883666 | 438.93 | 53.0 | 0 | 5.9 | ND | - | ND | ND | ND | ND | ND | ND |
| RO6892033 | 351.42 | 44.3 | 0 | 6.0 | ND | < 2.0 (B) | 1 | 0, 0 / 80 / 20 | 24 / 41 / ND | ND | ND | ND |
| RO6926274 | 401.87 | 49.7 | 0 | 6.2 | ND | - | ND | ND | ND | ND | ND | ND |
| RO7032019 | 412.41 | 70.4 | 1 | 1.5 | 2.52 | 2.2 (B, calc.) | 10 | 6.4, 12 / 43 / 45 | 10 / 10 / ND | ND | ND | ND |
| FMP7234690 | 708.80 | 151.3 | 2 | 9.0 | ND | 2.6 (B, calc.) | 2 | ND | 59 / 46 / ND | ND | ND | ND |
| FMP7234691 | 752.85 | 160.0 | 2 | 8.8 | 2.84 | 2.6 and 2.0 (B, calc.) | 9 | ND | 50 / 47 / ND | ND | ND | ND |
| FMP7234694 | 796.91 | 168.6 | 2 | 8.5 | ND | 2.6 and 2.0 (B, calc.) | 14 | 0.7, 2 / 38 / 61 | 130 / 132 / ND | ND | ND | ND |
| FMP7234698 | 840.96 | 176.9 | 2 | 8.2 | ND | 2.6 and 2.0 (B, calc.) | 6 | 0.7, 2 / 45 / 53 | 67 / 83 / ND | ND | ND | ND |
| FMP7234699 | 720.85 | 143.0 | 2 | 11.3 | ND | 2.6 (B, calc.) | 14 | 0.5, 2 / 12 / 87 | 22 / 11 / ND | ND | ND | ND |

MW – molecular weight; PSA – polar surface area; HBD – number of hydrogen-bond donors; B – basic; A – acidic; calc. – calculated; prec. – precipitated; micros. CL – microsomal clearance; hepat. CL – hepatocyte clearance; P-gp – P-glycoprotein; ND – not determined.
[a] Surface sum of all polar atoms in the molecule; [b] Calculated partition coefficient values (cLogP) from experimentally determined octanol/water partition coefficient values ($K_{ow}$); [c] Distribution coefficient values; [d]Solubility of the compound in aqueous buffer (pH 6.5) after lyophilization from DMSO superstock; [e] Parallel artificial membrane permeability assay (PAMPA) was used to determine membrane permeation coefficient values ($P_{eff}$), percentage of molecule that permeates into acceptor compartment, percentage of molecule found in membrane and percentage of molecule that stays in donor compartment.
*Italicised values* described in Soethoudt *et al. Nat. Commun.* (2017) **8**: 13958.

**Table S4.** Receptor and biosensor DNA amount used for transfection, optimized for the maximal ligand-induced response prior to ligand screen, given in nanograms of plasmid (pcDNA3.1 or pcDNA4) encoding for each biosensor component.

| condition | receptor | β-arrestin-RlucII | GRK2 | Gα | effector-RlucII | CAAX-rGFP | ssDNA |
|---|---|---|---|---|---|---|---|
| CB1, β-arrestin1 | 75 | 10 | 100 | - | - | 600 | 215 |
| CB1, β-arrestin2 | 125 | 10 | 100 | - | - | 600 | 165 |
| CB1, $G_{i1}$ | 25 | - | - | 100 | 10 | 500 | 365 |
| CB1, $G_{i2}$ | 25 | - | - | 100 | 10 | 500 | 365 |
| CB1, $G_{i3}$ | 25 | - | - | 100 | 10 | 600 | 265 |
| CB1, $G_{oA}$ | 25 | - | - | 100 | 10 | 500 | 365 |
| CB1, $G_{oB}$ | 25 | - | - | 100 | 10 | 500 | 365 |
| CB1, $G_z$ | 25 | - | - | 100 | 10 | 600 | 265 |
| CB1, $G_{12}$ | 25 | - | - | 100 | 10 | 500 | 365 |
| CB1, $G_{13}$ | 25 | - | - | 100 | 5 | 500 | 370 |
| CB1, G15 | 25 | - | - | 100 | 10 | 500 | 365 |
| CB2, β-arrestin1 | 75 | 25 | 100 | - | - | 600 | 200 |
| CB2, β-arrestin2 | 125 | 10 | 100 | - | - | 600 | 165 |
| CB2, $G_{i1}$ | 25 | - | - | 150 | 10 | 400 | 415 |
| CB2, $G_{i2}$ | 10 | - | - | 150 | 10 | 400 | 430 |
| CB2, $G_{i3}$ | 10 | - | - | 150 | 10 | 600 | 230 |
| CB2, $G_{oA}$ | 25 | - | - | 150 | 10 | 600 | 215 |
| CB2, $G_{oB}$ | 25 | - | - | 150 | 10 | 600 | 215 |

**Table S5.** Receptor and biosensor DNA amount used for transfection, optimized for the maximal ligand-induced response prior to ligand screen, given in nanograms of plasmid (pcDNA3.1 or pcDNA4) encoding for each biosensor component.

| condition | receptor | β-arrestin-RlucII | GRK2 | FYVE-rGFP | ssDNA |
|---|---|---|---|---|---|
| V2R, β-arrestin1 | 5 | 10 | - | 600 | 385 |
| V2R, β-arrestin2 | 5 | 10 | - | 600 | 385 |
| CB1, β-arrestin1 | 75 | 10 | 100 | 600 | 215 |
| CB1, β-arrestin2 | 125 | 10 | 100 | 600 | 165 |
| CB2, β-arrestin1 | 75 | 25 | 100 | 600 | 200 |
| CB2, β-arrestin2 | 125 | 10 | 100 | 600 | 165 |

**Table S6.** Receptor and biosensor DNA amount used for transfection of Gα–Gβγ dissociation-based G protein biosensor, given in nanograms of plasmid (pcDNA3.1 or pcDNA4) encoding for each biosensor component.

| **condition** | **receptor** | **Gα-RlucII** | **$G\beta_1$** | **$G\gamma_2$-GFP10** | **ssDNA** |
|---|---|---|---|---|---|
| CB1, $G_s$ | 200 | 100 | 100 | 400 | 200 |
| CB2, $G_s$ | 100 | 400 | 100 | 400 | 0 |

**Table S7.** Maximal ligand-induced response ($E_{max}$) indicates which pathways are activated by CB1 and CB2. Mean and standard error or mean (SE) from four technical replicates are shown.

| pathway | CB1 | | CB2 | |
|---|---|---|---|---|
| | **mean** | **SE** | **mean** | **SE** |
| $G_{i1}$ | 0.173 | 0.014 | 0.264 | 0.034 |
| $G_{i2}$ | 0.357 | 0.092 | 0.613 | 0.023 |
| $G_{i3}$ | 0.158 | 0.018 | 0.198 | 0.026 |
| $G_{oA}$ | 0.296 | 0.019 | 0.220 | 0.053 |
| $G_{oB}$ | 0.429 | 0.058 | 0.898 | 0.068 |
| $G_z$ | 0.299 | 0.061 | 0.191 | 0.119 |
| $G_q$ | 0.025 | 0.015 | 0.010 | 0.008 |
| $G_{11}$ | 0.015 | 0.010 | 0.007 | 0.009 |
| $G_{14}$ | -0.006 | 0.010 | 0.005 | 0.022 |
| $G_{15}$ | 0.214 | 0.012 | -0.004 | 0.015 |
| $G_{12}$ | 0.125 | 0.021 | 0.011 | 0.012 |
| $G_{13}$ | 0.150 | 0.015 | 0.008 | 0.013 |
| $G_s$ | -0.006 | 0.005 | 0.010 | 0.009 |
| β-arrestin1 | 0.044 | 0.015 | 0.110 | 0.031 |
| β-arrestin2 | 0.261 | 0.013 | 0.300 | 0.027 |

**Table S8.** Maximal ligand-induced response ($E_{max}$) and negative logarithm of half-maximal response concentration (pEC50) values extracted from CB1 ligand screen data, reported as mean of three biological replicates and the corresponding standard errors (SE).

| pathway | | HU210 | WIN55212-2 | CP55940 | nabilone | THC | cannabinol | anandamide | 2-AG |
|---|---|---|---|---|---|---|---|---|---|
| $G_{i1}$ | pEC50 | 9.419 | 8.413 | 9.172 | 8.282 | 8.595 | 8.387 | 6.978 | 6.822 |
| | SE (pEC50) | 0.121 | 0.086 | 0.119 | 0.108 | 0.120 | 0.231 | 0.096 | 0.194 |
| | $E_{max}$ | 0.358 | 0.647 | 0.326 | 0.321 | 0.532 | 0.244 | 0.164 | 0.317 |
| | SE ($E_{max}$) | 0.029 | 0.038 | 0.026 | 0.023 | 0.043 | 0.038 | 0.011 | 0.042 |
| $G_{i2}$ | pEC50 | 9.413 | 8.115 | 9.604 | 8.982 | 8.607 | 7.807 | 7.421 | 7.168 |
| | SE (pEC50) | 0.121 | 0.086 | 0.119 | 0.108 | 0.120 | 0.231 | 0.096 | 0.194 |
| | $E_{max}$ | 0.647 | 0.618 | 0.617 | 0.574 | 0.350 | 0.240 | 0.551 | 0.551 |
| | SE ($E_{max}$) | 0.029 | 0.038 | 0.026 | 0.023 | 0.043 | 0.038 | 0.011 | 0.042 |
| $G_{i3}$ | pEC50 | 9.172 | 7.484 | 9.426 | 9.102 | 8.466 | 7.885 | 7.364 | 6.830 |
| | SE (pEC50) | 0.119 | 0.114 | 0.108 | 0.133 | 0.166 | 0.333 | 0.107 | 0.096 |
| | $E_{max}$ | 0.326 | 0.348 | 0.334 | 0.292 | 0.180 | 0.080 | 0.243 | 0.296 |
| | SE ($E_{max}$) | 0.026 | 0.028 | 0.024 | 0.027 | 0.020 | 0.018 | 0.018 | 0.020 |
| $G_{oA}$ | pEC50 | 9.282 | 8.090 | 9.847 | 9.246 | 8.731 | 7.834 | 7.588 | 6.828 |
| | SE (pEC50) | 0.108 | 0.101 | 0.126 | 0.192 | 0.196 | 0.322 | 0.128 | 0.198 |
| | $E_{max}$ | 0.321 | 0.419 | 0.440 | 0.408 | 0.284 | 0.131 | 0.365 | 0.384 |
| | SE ($E_{max}$) | 0.023 | 0.029 | 0.038 | 0.056 | 0.038 | 0.029 | 0.032 | 0.052 |
| $G_{oB}$ | pEC50 | 9.595 | 8.189 | 9.612 | 9.348 | 8.524 | 7.591 | 7.694 | 6.978 |
| | SE (pEC50) | 0.120 | 0.075 | 0.114 | 0.142 | 0.230 | 0.188 | 0.099 | 0.123 |
| | $E_{max}$ | 0.532 | 0.713 | 0.665 | 0.625 | 0.379 | 0.303 | 0.525 | 0.621 |
| | SE ($E_{max}$) | 0.043 | 0.036 | 0.052 | 0.064 | 0.059 | 0.039 | 0.035 | 0.053 |
| $G_z$ | pEC50 | 9.387 | 8.321 | 9.857 | 9.016 | 8.470 | 6.940 | 7.712 | 7.000 |
| | SE (pEC50) | 0.231 | 0.142 | 0.173 | 0.277 | 0.263 | 0.405 | 0.304 | 0.186 |
| | $E_{max}$ | 0.244 | 0.399 | 0.316 | 0.251 | 0.189 | 0.188 | 0.157 | 0.291 |
| | SE ($E_{max}$) | 0.038 | 0.039 | 0.038 | 0.049 | 0.034 | 0.057 | 0.033 | 0.038 |
| $G_{12}$ | pEC50 | 8.978 | 7.413 | 8.932 | 8.442 | 7.961 | 8.843 | 6.790 | 6.069 |
| | SE (pEC50) | 0.096 | 0.071 | 0.090 | 0.112 | 0.231 | 0.525 | 0.101 | 0.051 |
| | $E_{max}$ | 0.164 | 0.291 | 0.230 | 0.196 | 0.055 | 0.025 | 0.134 | 0.270 |
| | SE ($E_{max}$) | 0.011 | 0.014 | 0.014 | 0.015 | 0.009 | 0.009 | 0.009 | 0.010 |
| $G_{13}$ | pEC50 | 8.750 | 7.497 | 8.866 | 8.310 | 7.923 | 7.300 | 7.047 | 6.148 |
| | SE (pEC50) | 0.053 | 0.055 | 0.058 | 0.080 | 0.180 | 0.155 | 0.072 | 0.045 |
| | $E_{max}$ | 0.300 | 0.385 | 0.345 | 0.341 | 0.151 | 0.121 | 0.276 | 0.416 |
| | SE ($E_{max}$) | 0.011 | 0.015 | 0.014 | 0.018 | 0.018 | 0.013 | 0.014 | 0.013 |
| $G_{15}$ | pEC50 | 8.913 | 7.285 | 9.043 | 8.576 | 8.572 | 6.116 | 6.921 | 6.083 |
| | SE (pEC50) | 0.247 | 0.111 | 0.173 | 0.120 | 0.523 | 0.464 | 0.191 | 0.088 |
| | $E_{max}$ | 0.140 | 0.316 | 0.257 | 0.218 | 0.048 | -0.117 | 0.143 | 0.375 |

| | | | | | | | | | |
|---|---|---|---|---|---|---|---|---|---|
| | SE ($E_{max}$) | 0.024 | 0.025 | 0.030 | 0.018 | 0.017 | 0.051 | 0.019 | 0.024 |
| β-arrestin1 | pEC50 | 8.313 | 7.044 | 8.886 | 8.192 | 7.660 | 0.000 | 6.826 | 4.867 |
| | SE (pEC50) | 0.131 | 0.125 | 0.155 | 0.122 | 0.428 | 0.000 | 0.294 | 0.101 |
| | $E_{max}$ | 0.074 | 0.117 | 0.058 | 0.070 | 0.028 | 0.000 | 0.045 | 0.214 |
| | SE ($E_{max}$) | 0.007 | 0.011 | 0.006 | 0.006 | 0.008 | 0.000 | 0.009 | 0.018 |
| β-arrestin2 | pEC50 | 8.353 | 7.137 | 8.680 | 8.102 | 7.927 | 6.713 | 6.646 | 5.321 |
| | SE (pEC50) | 0.038 | 0.041 | 0.038 | 0.047 | 0.108 | 0.273 | 0.063 | 0.027 |
| | $E_{max}$ | 0.318 | 0.445 | 0.271 | 0.317 | 0.085 | 0.040 | 0.214 | 0.587 |
| | SE ($E_{max}$) | 0.009 | 0.013 | 0.007 | 0.010 | 0.006 | 0.009 | 0.009 | 0.012 |

**Table S9.** Maximal ligand-induced response ($E_{max}$) and negative logarithm of half-maximal response concentration (pEC50) values extracted from CB2 ligand screen data, reported as mean of three independent experiments and the corresponding standard errors (SE).

| ligand | | $G_{i1}$ | $G_{i2}$ | $G_{i3}$ | $G_{oA}$ | $G_{oB}$ | β-arrestin1 | β-arrestin2 |
|---|---|---|---|---|---|---|---|---|
| JWH133 | pEC50 | 8.791 | 7.811 | 8.298 | 7.979 | 8.535 | 8.237 | 7.973 |
| | SE (pEC50) | 0.122 | 0.108 | 0.078 | 0.082 | 0.059 | 0.151 | 0.123 |
| | $E_{max}$ | 0.294 | 0.692 | 0.358 | 0.416 | 0.862 | 0.101 | 0.229 |
| | SE ($E_{max}$) | 0.025 | 0.051 | 0.019 | 0.023 | 0.035 | 0.010 | 0.019 |
| HU308 | pEC50 | 8.167 | 8.008 | 7.793 | 7.490 | 7.914 | 7.560 | 7.613 |
| | SE (pEC50) | 0.075 | 0.073 | 0.085 | 0.175 | 0.078 | 0.267 | 0.155 |
| | $E_{max}$ | 0.372 | 0.899 | 0.431 | 0.583 | 0.974 | 0.066 | 0.194 |
| | SE ($E_{max}$) | 0.026 | 0.064 | 0.041 | 0.137 | 0.078 | 0.012 | 0.020 |
| (*rac*)-AM1241 | pEC50 | 8.358 | 7.994 | 7.946 | 7.980 | 8.171 | 8.031 | 8.063 |
| | SE (pEC50) | 0.193 | 0.160 | 0.171 | 0.212 | 0.112 | 0.376 | 0.313 |
| | $E_{max}$ | 0.193 | 0.375 | 0.130 | 0.120 | 0.437 | 0.032 | 0.039 |
| | SE ($E_{max}$) | 0.025 | 0.041 | 0.015 | 0.017 | 0.033 | 0.008 | 0.008 |
| AM630 | pEC50 | 8.198 | 8.023 | 8.369 | 7.643 | 8.396 | 0.000 | 7.771 |
| | SE (pEC50) | 0.116 | 0.091 | 0.182 | 0.267 | 0.107 | 0.000 | 0.333 |
| | $E_{max}$ | -0.211 | -0.381 | -0.067 | -0.096 | -0.315 | 0.000 | -0.037 |
| | SE ($E_{max}$) | 0.017 | 0.024 | 0.008 | 0.018 | 0.023 | 0.000 | 0.008 |
| (*R*)-AM1241 | pEC50 | 8.299 | 8.227 | 8.339 | 8.555 | 8.120 | 0.000 | 8.399 |
| | SE (pEC50) | 0.164 | 0.128 | 0.161 | 0.219 | 0.095 | 0.000 | 0.287 |
| | $E_{max}$ | 0.212 | 0.364 | 0.138 | 0.105 | 0.409 | 0.000 | 0.056 |
| | SE ($E_{max}$) | 0.024 | 0.032 | 0.015 | 0.014 | 0.026 | 0.000 | 0.011 |
| (*S*)-AM1241 | pEC50 | 7.434 | 7.308 | 7.338 | 6.975 | 7.329 | 7.323 | 6.734 |
| | SE (pEC50) | 0.143 | 0.086 | 0.098 | 0.119 | 0.079 | 0.347 | 0.131 |
| | $E_{max}$ | 0.273 | 0.582 | 0.285 | 0.275 | 0.627 | 0.047 | 0.155 |
| | SE ($E_{max}$) | 0.027 | 0.035 | 0.020 | 0.024 | 0.035 | 0.012 | 0.015 |
| HU210 | pEC50 | 9.414 | 9.594 | 9.273 | 9.138 | 9.591 | 9.309 | 9.361 |
| | SE (pEC50) | 0.136 | 0.077 | 0.136 | 0.082 | 0.071 | 0.223 | 0.150 |
| | $E_{max}$ | 0.321 | 0.670 | 0.327 | 0.347 | 0.716 | 0.085 | 0.152 |
| | SE ($E_{max}$) | 0.030 | 0.035 | 0.030 | 0.019 | 0.035 | 0.012 | 0.016 |
| WIN55212-2 | pEC50 | 9.409 | 9.520 | 9.326 | 9.095 | 9.271 | 9.284 | 9.468 |
| | SE (pEC50) | 0.047 | 0.045 | 0.042 | 0.064 | 0.049 | 0.197 | 0.132 |
| | $E_{max}$ | 0.404 | 0.703 | 0.376 | 0.388 | 0.830 | 0.081 | 0.196 |
| | SE ($E_{max}$) | 0.013 | 0.022 | 0.011 | 0.018 | 0.029 | 0.011 | 0.019 |
| CP55940 | pEC50 | 10.010 | 9.972 | 9.985 | 9.574 | 9.908 | 9.393 | 9.383 |
| | SE (pEC50) | 0.137 | 0.087 | 0.096 | 0.099 | 0.075 | 0.189 | 0.125 |
| | $E_{max}$ | 0.244 | 0.640 | 0.345 | 0.394 | 0.813 | 0.102 | 0.269 |
| | SE ($E_{max}$) | 0.023 | 0.039 | 0.023 | 0.027 | 0.042 | 0.013 | 0.022 |
| nabilone | pEC50 | 8.885 | 9.165 | 8.977 | 8.297 | 8.737 | 8.640 | 8.658 |
| | SE (pEC50) | 0.186 | 0.106 | 0.123 | 0.068 | 0.085 | 0.267 | 0.151 |
| | $E_{max}$ | 0.319 | 0.511 | 0.322 | 0.396 | 0.759 | 0.087 | 0.193 |
| | SE ($E_{max}$) | 0.041 | 0.038 | 0.027 | 0.018 | 0.044 | 0.016 | 0.020 |
| THC | pEC50 | 8.123 | 7.886 | 7.791 | 8.597 | 8.116 | 0.000 | 8.458 |
| | SE (pEC50) | 0.249 | 0.172 | 0.216 | 0.419 | 0.142 | 0.000 | 0.577 |
| | $E_{max}$ | 0.110 | 0.231 | 0.104 | 0.070 | 0.259 | 0.000 | 0.028 |
| | SE ($E_{max}$) | 0.018 | 0.027 | 0.015 | 0.020 | 0.025 | 0.000 | 0.011 |

| | | | | | | | | |
|---|---|---|---|---|---|---|---|---|
| cannabinol | pEC50 | 7.820 | 7.506 | 7.356 | 8.350 | 7.662 | 0.000 | 6.664 |
| | SE (pEC50) | 0.246 | 0.165 | 0.122 | 0.220 | 0.147 | 0.000 | 0.480 |
| | $E_{max}$ | 0.164 | 0.307 | 0.152 | 0.140 | 0.338 | 0.000 | 0.033 |
| | SE ($E_{max}$) | 0.028 | 0.035 | 0.013 | 0.021 | 0.034 | 0.000 | 0.012 |
| anandamide | pEC50 | 7.415 | 7.173 | 7.165 | 7.095 | 7.563 | 5.448 | 6.509 |
| | SE (pEC50) | 0.126 | 0.086 | 0.092 | 0.140 | 0.105 | 0.175 | 0.224 |
| | $E_{max}$ | 0.266 | 0.520 | 0.220 | 0.207 | 0.587 | 0.109 | 0.094 |
| | SE ($E_{max}$) | 0.023 | 0.030 | 0.014 | 0.020 | 0.042 | 0.016 | 0.015 |
| 2-AG | pEC50 | 7.027 | 6.827 | 6.742 | 6.797 | 7.140 | 5.997 | 5.979 |
| | SE (pEC50) | 0.156 | 0.088 | 0.097 | 0.115 | 0.078 | 0.121 | 0.090 |
| | $E_{max}$ | 0.331 | 0.659 | 0.343 | 0.325 | 0.711 | 0.141 | 0.316 |
| | SE ($E_{max}$) | 0.037 | 0.042 | 0.024 | 0.026 | 0.040 | 0.012 | 0.021 |
| SR144528 | pEC50 | 8.624 | 8.353 | 8.825 | 9.318 | 9.004 | 8.523 | 9.173 |
| | SE (pEC50) | 0.099 | 0.099 | 0.152 | 0.175 | 0.114 | 0.288 | 0.168 |
| | $E_{max}$ | -0.264 | -0.410 | -0.112 | -0.084 | -0.355 | -0.033 | -0.060 |
| | SE ($E_{max}$) | 0.018 | 0.029 | 0.012 | 0.010 | 0.028 | 0.007 | 0.007 |
| RO6843766 | pEC50 | 9.341 | 9.096 | 9.127 | 9.110 | 9.314 | 9.967 | 9.557 |
| | SE (pEC50) | 0.181 | 0.101 | 0.115 | 0.152 | 0.079 | 0.443 | 0.234 |
| | $E_{max}$ | 0.226 | 0.422 | 0.213 | 0.177 | 0.504 | 0.028 | 0.068 |
| | SE ($E_{max}$) | 0.028 | 0.029 | 0.017 | 0.018 | 0.027 | 0.009 | 0.011 |
| RO6853457 | pEC50 | 8.431 | 8.535 | 8.344 | 8.290 | 8.045 | 7.960 | 7.655 |
| | SE (pEC50) | 0.182 | 0.087 | 0.161 | 0.212 | 0.096 | 0.471 | 0.251 |
| | $E_{max}$ | 0.193 | 0.392 | 0.114 | 0.127 | 0.389 | 0.033 | 0.081 |
| | SE ($E_{max}$) | 0.025 | 0.024 | 0.013 | 0.019 | 0.027 | 0.011 | 0.016 |
| RO7032019 | pEC50 | 8.588 | 8.561 | 8.103 | 8.137 | 8.423 | 8.130 | 8.226 |
| | SE (pEC50) | 0.299 | 0.145 | 0.149 | 0.159 | 0.100 | 0.368 | 0.213 |
| | $E_{max}$ | 0.077 | 0.416 | 0.184 | 0.210 | 0.606 | 0.031 | 0.064 |
| | SE ($E_{max}$) | 0.016 | 0.042 | 0.020 | 0.024 | 0.042 | 0.008 | 0.010 |
| RO6871487 | pEC50 | 8.016 | 8.085 | 8.540 | 8.430 | 8.160 | 0.000 | 8.037 |
| | SE (pEC50) | 0.193 | 0.105 | 0.216 | 0.193 | 0.156 | 0.000 | 0.674 |
| | $E_{max}$ | 0.144 | 0.313 | 0.074 | 0.106 | 0.249 | 0.000 | 0.026 |
| | SE ($E_{max}$) | 0.020 | 0.024 | 0.011 | 0.014 | 0.028 | 0.000 | 0.013 |
| RO6878558 | pEC50 | 7.945 | 7.986 | 7.893 | 8.028 | 7.824 | 8.512 | 7.624 |
| | SE (pEC50) | 0.152 | 0.123 | 0.192 | 0.169 | 0.079 | 0.370 | 0.487 |
| | $E_{max}$ | 0.230 | 0.419 | 0.144 | 0.154 | 0.555 | 0.029 | 0.054 |
| | SE ($E_{max}$) | 0.026 | 0.038 | 0.021 | 0.019 | 0.033 | 0.007 | 0.021 |
| RO6883666 | pEC50 | 8.021 | 7.652 | 7.986 | 8.613 | 7.903 | 0.000 | 0.000 |
| | SE (pEC50) | 0.148 | 0.146 | 0.417 | 0.259 | 0.158 | 0.000 | 0.000 |
| | $E_{max}$ | 0.165 | 0.351 | 0.060 | 0.100 | 0.305 | 0.000 | 0.000 |
| | SE ($E_{max}$) | 0.018 | 0.041 | 0.018 | 0.018 | 0.036 | 0.000 | 0.000 |
| RO6853973 | pEC50 | 9.052 | 9.450 | 9.296 | 9.017 | 9.329 | 0.000 | 0.000 |
| | SE (pEC50) | 0.183 | 0.163 | 0.394 | 0.262 | 0.215 | 0.000 | 0.000 |
| | $E_{max}$ | 0.141 | 0.181 | 0.049 | 0.076 | 0.167 | 0.000 | 0.000 |
| | SE ($E_{max}$) | 0.018 | 0.020 | 0.013 | 0.014 | 0.024 | 0.000 | 0.000 |
| RO6850007 | pEC50 | 8.267 | 8.237 | 7.925 | 8.319 | 8.709 | 8.371 | 8.806 |

| | | | | | | | | |
|---|---|---|---|---|---|---|---|---|
| | SE (pEC50) | 0.201 | 0.188 | 0.321 | 0.322 | 0.277 | 0.747 | 0.553 |
| | $E_{max}$ | 0.110 | 0.179 | 0.070 | 0.119 | 0.191 | 0.034 | 0.045 |
| | SE ($E_{max}$) | 0.016 | 0.024 | 0.017 | 0.027 | 0.036 | 0.018 | 0.017 |
| RO6844395 | pEC50 | 7.855 | 7.499 | 7.994 | 7.607 | 7.665 | 0.000 | 7.671 |
| | SE (pEC50) | 0.137 | 0.106 | 0.127 | 0.167 | 0.103 | 0.000 | 0.274 |
| | $E_{max}$ | 0.205 | 0.537 | 0.167 | 0.274 | 0.637 | 0.000 | 0.077 |
| | SE ($E_{max}$) | 0.021 | 0.046 | 0.016 | 0.036 | 0.051 | 0.000 | 0.016 |
| RO6844112 | pEC50 | 8.632 | 8.496 | 8.160 | 8.159 | 8.497 | 7.714 | 8.242 |
| | SE (pEC50) | 0.109 | 0.083 | 0.121 | 0.165 | 0.080 | 0.189 | 0.269 |
| | $E_{max}$ | 0.273 | 0.554 | 0.223 | 0.208 | 0.628 | 0.056 | 0.090 |
| | SE ($E_{max}$) | 0.021 | 0.032 | 0.019 | 0.025 | 0.035 | 0.008 | 0.017 |
| RO5135445 | pEC50 | 8.748 | 9.121 | 8.563 | 8.718 | 8.504 | 8.438 | 9.551 |
| | SE (pEC50) | 0.164 | 0.108 | 0.161 | 0.218 | 0.096 | 0.431 | 0.659 |
| | $E_{max}$ | 0.182 | 0.345 | 0.119 | 0.153 | 0.470 | 0.037 | 0.041 |
| | SE ($E_{max}$) | 0.020 | 0.025 | 0.013 | 0.023 | 0.031 | 0.011 | 0.019 |
| RO6926274 | pEC50 | 8.574 | 9.040 | 8.202 | 8.250 | 8.765 | 0.000 | 0.000 |
| | SE (pEC50) | 0.436 | 0.216 | 0.239 | 0.330 | 0.417 | 0.000 | 0.000 |
| | $E_{max}$ | 0.086 | 0.178 | 0.078 | 0.105 | 0.108 | 0.000 | 0.000 |
| | SE ($E_{max}$) | 0.026 | 0.026 | 0.015 | 0.025 | 0.031 | 0.000 | 0.000 |
| RO6892033 | pEC50 | 8.536 | 8.599 | 8.856 | 8.018 | 8.328 | 8.566 | 0.000 |
| | SE (pEC50) | 0.219 | 0.114 | 0.225 | 0.225 | 0.148 | 0.161 | 0.000 |
| | $E_{max}$ | 0.135 | 0.301 | 0.095 | 0.173 | 0.305 | 0.050 | 0.000 |
| | SE ($E_{max}$) | 0.020 | 0.024 | 0.015 | 0.029 | 0.032 | 0.006 | 0.000 |
| RO6435559 | pEC50 | 7.984 | 8.154 | 7.941 | 7.998 | 7.794 | 8.518 | 8.131 |
| | SE (pEC50) | 0.152 | 0.131 | 0.246 | 0.192 | 0.110 | 0.235 | 0.326 |
| | $E_{max}$ | 0.202 | 0.359 | 0.125 | 0.217 | 0.415 | 0.028 | 0.024 |
| | SE ($E_{max}$) | 0.023 | 0.034 | 0.023 | 0.031 | 0.035 | 0.005 | 0.006 |
| RO6869094 | pEC50 | 8.214 | 8.341 | 7.645 | 0.000 | 8.321 | 9.084 | 8.016 |
| | SE (pEC50) | 0.340 | 0.198 | 0.561 | 0.000 | 0.217 | 0.251 | 0.310 |
| | $E_{max}$ | 0.070 | 0.158 | 0.055 | 0.000 | 0.148 | 0.028 | 0.027 |
| | SE ($E_{max}$) | 0.017 | 0.022 | 0.024 | 0.000 | 0.023 | 0.005 | 0.006 |
| FMP7234690 | pEC50 | 7.991 | 8.358 | 7.571 | 7.302 | 8.018 | 8.536 | 8.042 |
| | SE (pEC50) | 0.133 | 0.097 | 0.121 | 0.207 | 0.088 | 0.604 | 0.297 |
| | $E_{max}$ | 0.222 | 0.438 | 0.259 | 0.240 | 0.606 | 0.033 | 0.094 |
| | SE ($E_{max}$) | 0.022 | 0.030 | 0.025 | 0.043 | 0.039 | 0.014 | 0.021 |
| FMP7234691 | pEC50 | 8.148 | 8.416 | 8.173 | 7.954 | 7.997 | 8.981 | 8.224 |
| | SE (pEC50) | 0.132 | 0.075 | 0.095 | 0.161 | 0.076 | 0.503 | 0.181 |
| | $E_{max}$ | 0.271 | 0.561 | 0.249 | 0.309 | 0.669 | 0.046 | 0.156 |
| | SE ($E_{max}$) | 0.026 | 0.030 | 0.017 | 0.037 | 0.038 | 0.016 | 0.020 |
| FMP7234694 | pEC50 | 7.846 | 8.210 | 7.982 | 8.199 | 8.243 | 8.120 | 8.109 |
| | SE (pEC50) | 0.124 | 0.062 | 0.103 | 0.144 | 0.084 | 0.616 | 0.108 |
| | $E_{max}$ | 0.251 | 0.585 | 0.267 | 0.316 | 0.601 | 0.032 | 0.161 |
| | SE ($E_{max}$) | 0.023 | 0.026 | 0.020 | 0.033 | 0.036 | 0.014 | 0.013 |
| FMP7234698 | pEC50 | 7.781 | 8.041 | 8.165 | 8.429 | 8.449 | 8.851 | 7.858 |
| | SE (pEC50) | 0.115 | 0.113 | 0.161 | 0.125 | 0.080 | 0.207 | 0.096 |

| | | | | | | | | |
|---|---|---|---|---|---|---|---|---|
| | $E_{max}$ | 0.258 | 0.519 | 0.247 | 0.354 | 0.600 | 0.067 | 0.188 |
| | SE ($E_{max}$) | 0.023 | 0.043 | 0.029 | 0.031 | 0.034 | 0.010 | 0.014 |
| FMP7234699 | pEC50 | 7.933 | 7.960 | 7.886 | 8.330 | 7.866 | 8.094 | 8.058 |
| | SE (pEC50) | 0.141 | 0.126 | 0.136 | 0.158 | 0.093 | 0.175 | 0.153 |
| | $E_{max}$ | 0.222 | 0.590 | 0.224 | 0.337 | 0.671 | 0.072 | 0.144 |
| | SE ($E_{max}$) | 0.023 | 0.055 | 0.023 | 0.038 | 0.047 | 0.009 | 0.016 |

**Table S10.** Concentration-response CB1-mediated signaling by anandamide. Mean and standard error of mean (SE) from three biological replicates are shown.

| $\log\left(\frac{[\text{ligand}]}{\text{mol L}^{-1}}\right)$ | **$G_{i2}$** | | **$G_{oB}$** | | **$G_z$** | | **$G_{12}$** | | **$G_{13}$** | | **$G_{15}$** | | **β-arrestin1** | | **β-arrestin2** | |
|---|---|---|---|---|---|---|---|---|---|---|---|---|---|---|---|---|
| | mean | SE | mean | SE | mean | SE | mean | SE | mean | SE | mean | SE | mean | SE | mean | SE |
| -10 | 0.037 | 0.074 | 0.032 | 0.057 | 0.055 | 0.061 | 0.008 | 0.020 | -0.045 | 0.025 | 0.038 | 0.054 | 0.037 | 0.050 | 0.015 | 0.014 |
| -9.5 | -0.076 | 0.047 | 0.017 | 0.034 | 0.043 | 0.089 | -0.011 | 0.022 | 0.003 | 0.030 | 0.037 | 0.045 | -0.037 | 0.069 | -0.001 | 0.013 |
| -9 | 0.031 | 0.054 | -0.003 | 0.080 | -0.007 | 0.076 | 0.019 | 0.033 | 0.012 | 0.035 | -0.015 | 0.056 | 0.010 | 0.064 | 0.007 | 0.010 |
| -8.5 | 0.120 | 0.051 | 0.127 | 0.042 | -0.033 | 0.064 | -0.018 | 0.014 | 0.051 | 0.040 | -0.045 | 0.054 | 0.030 | 0.087 | 0.011 | 0.015 |
| -8 | 0.184 | 0.058 | 0.205 | 0.047 | 0.118 | 0.059 | 0.043 | 0.024 | 0.086 | 0.038 | 0.014 | 0.075 | -0.005 | 0.050 | 0.012 | 0.013 |
| -7.5 | 0.413 | 0.060 | 0.426 | 0.042 | 0.256 | 0.072 | 0.058 | 0.028 | 0.196 | 0.038 | 0.114 | 0.057 | 0.061 | 0.057 | 0.042 | 0.013 |
| -7 | 0.626 | 0.048 | 0.685 | 0.030 | 0.420 | 0.059 | 0.211 | 0.034 | 0.379 | 0.049 | 0.220 | 0.036 | 0.174 | 0.088 | 0.140 | 0.015 |
| -6.5 | 0.792 | 0.087 | 0.751 | 0.023 | 0.370 | 0.060 | 0.280 | 0.030 | 0.536 | 0.040 | 0.340 | 0.044 | 0.207 | 0.083 | 0.293 | 0.009 |
| -6 | 0.888 | 0.085 | 0.685 | 0.051 | 0.341 | 0.066 | 0.387 | 0.022 | 0.646 | 0.036 | 0.350 | 0.056 | 0.413 | 0.089 | 0.399 | 0.024 |
| -5.5 | 0.831 | 0.076 | 0.698 | 0.037 | 0.352 | 0.096 | 0.424 | 0.026 | 0.704 | 0.033 | 0.378 | 0.061 | 0.362 | 0.082 | 0.443 | 0.033 |
| -5 | 0.827 | 0.072 | 0.712 | 0.047 | 0.432 | 0.085 | 0.487 | 0.059 | 0.702 | 0.027 | 0.484 | 0.083 | 0.378 | 0.095 | 0.459 | 0.029 |
| -4.5 | 0.989 | 0.125 | 0.739 | 0.057 | 0.369 | 0.136 | 0.448 | 0.033 | 0.733 | 0.018 | 0.499 | 0.054 | 0.345 | 0.079 | 0.483 | 0.031 |

**Table S11.** Concentration-response CB1-mediated signaling by 2-AG. Mean and standard error of mean (SE) from three biological replicates are shown.

| $\log\left(\frac{[\text{ligand}]}{\text{mol L}^{-1}}\right)$ | $G_{i2}$ | | $G_{oB}$ | | $G_z$ | | $G_{12}$ | | $G_{13}$ | | $G_{15}$ | | β-arrestin1 | | β-arrestin2 | |
|---|---|---|---|---|---|---|---|---|---|---|---|---|---|---|---|---|
| | mean | SE | mean | SE | mean | SE | mean | SE | mean | SE | mean | SE | mean | SE | mean | SE |
| -9.3 | -0.082 | 0.059 | -0.031 | 0.048 | 0.008 | 0.088 | -0.004 | 0.032 | -0.021 | 0.026 | 0.039 | 0.050 | -0.099 | 0.055 | -0.007 | 0.010 |
| -8.8 | 0.068 | 0.050 | 0.034 | 0.053 | -0.040 | 0.064 | 0.013 | 0.018 | -0.032 | 0.021 | -0.124 | 0.070 | -0.041 | 0.091 | -0.010 | 0.007 |
| -8.3 | 0.063 | 0.070 | 0.057 | 0.056 | 0.059 | 0.045 | 0.002 | 0.028 | 0.004 | 0.025 | -0.048 | 0.060 | -0.039 | 0.083 | -0.016 | 0.011 |
| -7.8 | 0.241 | 0.079 | 0.123 | 0.062 | 0.142 | 0.072 | 0.018 | 0.024 | 0.052 | 0.023 | 0.037 | 0.051 | -0.044 | 0.081 | -0.020 | 0.015 |
| -7.3 | 0.374 | 0.081 | 0.260 | 0.074 | 0.231 | 0.113 | 0.051 | 0.024 | 0.113 | 0.029 | 0.161 | 0.073 | 0.003 | 0.056 | 0.005 | 0.010 |
| -6.8 | 0.586 | 0.063 | 0.538 | 0.075 | 0.439 | 0.070 | 0.133 | 0.027 | 0.177 | 0.024 | 0.250 | 0.067 | 0.055 | 0.047 | 0.039 | 0.026 |
| -6.3 | 0.760 | 0.083 | 0.718 | 0.083 | 0.629 | 0.116 | 0.353 | 0.039 | 0.477 | 0.038 | 0.474 | 0.035 | 0.154 | 0.070 | 0.207 | 0.029 |
| -5.8 | 0.859 | 0.112 | 0.806 | 0.078 | 0.610 | 0.083 | 0.593 | 0.032 | 0.737 | 0.039 | 0.759 | 0.058 | 0.330 | 0.071 | 0.337 | 0.019 |
| -5.3 | 0.890 | 0.085 | 0.861 | 0.080 | 0.735 | 0.097 | 0.809 | 0.039 | 0.896 | 0.031 | 0.917 | 0.048 | 0.502 | 0.060 | 0.680 | 0.018 |
| -4.8 | 0.899 | 0.091 | 0.898 | 0.073 | 0.770 | 0.096 | 0.862 | 0.018 | 1.011 | 0.029 | 1.054 | 0.054 | 1.027 | 0.155 | 0.960 | 0.018 |
| -4.3 | 0.904 | 0.123 | 0.814 | 0.091 | 0.653 | 0.103 | 0.917 | 0.036 | 1.126 | 0.023 | 1.321 | 0.081 | 1.184 | 0.103 | 1.187 | 0.023 |
| -3.8 | 0.888 | 0.170 | 0.923 | 0.123 | 0.801 | 0.132 | - | - | - | - | - | - | 1.864 | 0.144 | 1.332 | 0.017 |

**Table S12.** Concentration-response CB2-mediated signaling by anandamide. Mean and standard error or mean (SE) from three independent experiments are shown.

| $\log\left(\frac{[\text{ligand}]}{\text{mol L}^{-1}}\right)$ | **$G_{i2}$** | | **$G_{oB}$** | | **β-arrestin1** | | **β-arrestin2** | |
|---|---|---|---|---|---|---|---|---|
| | mean | SE | mean | SE | mean | SE | mean | SE |
| -10 | 0.027 | 0.033 | 0.037 | 0.033 | -0.125 | 0.082 | -0.047 | 0.051 |
| -9.5 | 0.000 | 0.038 | -0.022 | 0.042 | -0.021 | 0.097 | 0.071 | 0.079 |
| -9 | -0.013 | 0.048 | 0.019 | 0.054 | 0.014 | 0.139 | 0.105 | 0.088 |
| -8.5 | 0.044 | 0.027 | 0.040 | 0.041 | -0.015 | 0.089 | 0.009 | 0.055 |
| -8 | 0.063 | 0.027 | 0.241 | 0.051 | -0.097 | 0.088 | -0.054 | 0.044 |
| -7.5 | 0.262 | 0.037 | 0.379 | 0.076 | 0.012 | 0.113 | -0.028 | 0.047 |
| -7 | 0.479 | 0.039 | 0.533 | 0.053 | 0.231 | 0.109 | 0.107 | 0.051 |
| -6.5 | 0.559 | 0.056 | 0.607 | 0.049 | 0.175 | 0.096 | 0.238 | 0.061 |
| -6 | 0.635 | 0.041 | 0.685 | 0.060 | 0.327 | 0.097 | 0.397 | 0.050 |
| -5.5 | 0.712 | 0.037 | 0.727 | 0.050 | 0.607 | 0.101 | 0.502 | 0.102 |
| -5 | 0.760 | 0.045 | 0.713 | 0.037 | 0.909 | 0.154 | 0.473 | 0.095 |
| -4.5 | 0.792 | 0.065 | 0.722 | 0.041 | 1.259 | 0.181 | 0.391 | 0.066 |

**Table S13.** Concentration-response CB2-mediated signaling by 2-AG. Mean and standard error or mean (SE) from three independent experiments are shown.

| $\log\left(\frac{[\text{ligand}]}{\text{mol L}^{-1}}\right)$ | **$G_{i2}$** | | **$G_{oB}$** | | **β-arrestin1** | | **β-arrestin2** | |
|---|---|---|---|---|---|---|---|---|
| | mean | SE | mean | SE | mean | SE | mean | SE |
| -9.3 | -0.016 | 0.030 | -0.052 | 0.036 | -0.248 | 0.117 | -0.053 | 0.044 |
| -8.8 | -0.014 | 0.034 | 0.034 | 0.040 | -0.150 | 0.083 | 0.020 | 0.032 |
| -8.3 | 0.024 | 0.041 | 0.107 | 0.037 | 0.012 | 0.082 | 0.029 | 0.038 |
| -7.8 | 0.115 | 0.033 | 0.148 | 0.050 | 0.179 | 0.100 | 0.043 | 0.047 |
| -7.3 | 0.297 | 0.022 | 0.360 | 0.048 | 0.227 | 0.086 | 0.061 | 0.062 |
| -6.8 | 0.461 | 0.045 | 0.546 | 0.051 | 0.397 | 0.123 | 0.196 | 0.085 |
| -6.3 | 0.704 | 0.059 | 0.767 | 0.030 | 0.630 | 0.133 | 0.579 | 0.125 |
| -5.8 | 0.810 | 0.078 | 0.852 | 0.029 | 0.884 | 0.194 | 0.952 | 0.128 |
| -5.3 | 0.838 | 0.085 | 0.894 | 0.037 | 1.395 | 0.156 | 1.250 | 0.126 |
| -4.8 | 1.061 | 0.038 | 0.818 | 0.056 | 1.783 | 0.138 | 1.667 | 0.079 |
| -4.3 | 0.980 | 0.063 | 0.791 | 0.036 | 1.643 | 0.096 | 1.510 | 0.116 |
| -3.8 | - | - | - | - | 1.548 | 0.236 | 1.622 | 0.192 |

**Table S14.** Calculation of CB1 relative effectiveness (RE = $10^{\Delta logR}$) as described in van der Westhuizen *et al.*, using WIN55212-2 as a reference ligand. NC = not converged, ND = not determined, IA = inverse agonist. ΔlogR = logR(ligand) - logR(reference).

| pathway | | WIN55,212-2 | HU210 | CP55940 | nabilone | anandamide | 2-AG | THC | cannabinol |
|---|---|---|---|---|---|---|---|---|---|
| $G_{i1}$ | logR | 8.414 | 9.162 | 8.874 | 7.978 | 6.382 | 6.512 | 8.512 | 7.965 |
| | SE (logR) | 0.064 | 0.108 | 0.120 | 0.121 | 0.239 | 0.123 | 0.073 | 0.159 |
| | ΔlogR | 0.000 | 0.748 | 0.460 | -0.436 | -2.032 | -1.902 | 0.098 | -0.449 |
| | SE (ΔlogR) | 0.091 | 0.126 | 0.136 | 0.137 | 0.248 | 0.138 | 0.097 | 0.171 |
| | RE | 1.000 | 5.598 | 2.884 | 0.366 | 0.009 | 0.013 | 1.253 | 0.356 |
| $G_{i2}$ | logR | 8.094 | 9.414 | 9.583 | 8.931 | 7.353 | 7.099 | 8.519 | 7.378 |
| | SE (logR) | 0.085 | 0.086 | 0.084 | 0.090 | 0.095 | 0.094 | 0.143 | 0.220 |
| | ΔlogR | 0.000 | 1.320 | 1.489 | 0.837 | -0.741 | -0.995 | 0.425 | -0.716 |
| | SE (ΔlogR) | 0.121 | 0.121 | 0.120 | 0.124 | 0.127 | 0.127 | 0.166 | 0.236 |
| | RE | 1.000 | 20.893 | 30.832 | 6.871 | 0.182 | 0.101 | 2.661 | 0.192 |
| $G_{i3}$ | logR | 7.484 | 9.143 | 9.408 | 9.026 | 7.207 | 6.759 | 8.161 | 7.243 |
| | SE (logR) | 0.086 | 0.082 | 0.079 | 0.089 | 0.108 | 0.110 | 0.142 | 0.332 |
| | ΔlogR | 0.000 | 1.659 | 1.924 | 1.542 | -0.277 | -0.725 | 0.677 | -0.241 |
| | SE (ΔlogR) | 0.122 | 0.119 | 0.117 | 0.124 | 0.138 | 0.139 | 0.166 | 0.343 |
| | RE | 1.000 | 45.604 | 83.946 | 34.834 | 0.528 | 0.188 | 4.753 | 0.574 |
| $G_{oA}$ | logR | 8.068 | 9.145 | 9.847 | 9.213 | 7.506 | 6.768 | 8.603 | 7.308 |
| | SE (logR) | 0.105 | 0.135 | 0.102 | 0.104 | 0.118 | 0.113 | 0.147 | 0.334 |
| | ΔlogR | 0.000 | 1.077 | 1.779 | 1.145 | -0.562 | -1.300 | 0.535 | -0.760 |
| | SE (ΔlogR) | 0.149 | 0.171 | 0.147 | 0.148 | 0.158 | 0.154 | 0.181 | 0.351 |
| | RE | 1.000 | 11.940 | 60.117 | 13.964 | 0.274 | 0.050 | 3.428 | 0.174 |
| $G_{oB}$ | logR | 8.189 | 9.470 | 9.581 | 9.291 | 7.562 | 6.919 | 8.613 | 7.221 |
| | SE (logR) | 0.078 | 0.097 | 0.078 | 0.082 | 0.098 | 0.084 | 0.126 | 0.176 |
| | ΔlogR | 0.000 | 1.281 | 1.392 | 1.102 | -0.627 | -1.270 | 0.424 | -0.968 |
| | SE (ΔlogR) | 0.111 | 0.125 | 0.111 | 0.113 | 0.126 | 0.115 | 0.148 | 0.193 |
| | RE | 1.000 | 19.099 | 24.660 | 12.647 | 0.236 | 0.054 | 2.655 | 0.108 |
| $G_z$ | logR | 8.322 | 9.174 | 9.753 | 8.816 | 7.304 | 6.863 | 8.032 | 6.613 |
| | SE (logR) | 0.118 | 0.181 | 0.139 | 0.173 | 0.278 | 0.151 | 0.220 | 0.315 |
| | ΔlogR | 0.000 | 0.852 | 1.431 | 0.494 | -1.018 | -1.459 | -0.290 | -1.709 |
| | SE (ΔlogR) | 0.167 | 0.216 | 0.182 | 0.210 | 0.302 | 0.192 | 0.249 | 0.336 |
| | RE | 1.000 | 7.112 | 26.977 | 3.119 | 0.096 | 0.035 | 0.513 | 0.020 |
| $G_{12}$ | logR | 7.413 | 8.729 | 8.830 | 8.270 | 6.452 | 6.036 | 7.045 | 7.785 |
| | SE (logR) | 0.054 | 0.083 | 0.060 | 0.069 | 0.103 | 0.054 | 0.243 | 0.519 |

| | | | | | | | | | |
|---|---|---|---|---|---|---|---|---|---|
| | ΔlogR | 0.000 | 1.316 | 1.417 | 0.857 | -0.961 | -1.377 | -0.368 | 0.372 |
| | SE (ΔlogR) | 0.076 | 0.099 | 0.081 | 0.088 | 0.116 | 0.076 | 0.249 | 0.521 |
| | RE | 1.000 | 20.701 | 26.122 | 7.194 | 0.109 | 0.042 | 0.429 | 2.355 |
| $G_{13}$ | logR | 7.438 | 8.624 | 8.800 | 8.239 | 6.885 | 6.198 | 7.497 | 6.781 |
| | SE (logR) | 0.044 | 0.057 | 0.050 | 0.049 | 0.062 | 0.044 | 0.120 | 0.146 |
| | ΔlogR | 0.000 | 1.186 | 1.362 | 0.801 | -0.553 | -1.240 | 0.059 | -0.657 |
| | SE (ΔlogR) | 0.062 | 0.072 | 0.067 | 0.066 | 0.076 | 0.062 | 0.127 | 0.152 |
| | RE | 1.000 | 15.346 | 23.014 | 6.324 | 0.280 | 0.058 | 1.146 | 0.220 |
| $G_{15}$ | logR | 7.210 | 8.485 | 8.879 | 8.340 | 6.501 | 6.083 | 7.766 | IA |
| | SE (logR) | 0.089 | 0.189 | 0.103 | 0.119 | 0.185 | 0.083 | 0.521 | IA |
| | ΔlogR | 0.000 | 1.275 | 1.669 | 1.130 | -0.709 | -1.127 | 0.556 | ND |
| | SE (ΔlogR) | 0.125 | 0.208 | 0.136 | 0.149 | 0.205 | 0.121 | 0.529 | ND |
| | RE | 1.000 | 18.836 | 46.666 | 13.490 | 0.195 | 0.075 | 3.597 | ND |
| β-arrestin1 | logR | 6.783 | 7.854 | 8.318 | 7.707 | 6.149 | 4.868 | 6.776 | NC |
| | SE (logR) | 0.088 | 0.134 | 0.165 | 0.135 | 0.212 | 0.069 | 0.348 | NC |
| | ΔlogR | 0.000 | 1.071 | 1.535 | 0.924 | -0.634 | -1.915 | -0.007 | ND |
| | SE (ΔlogR) | 0.125 | 0.161 | 0.187 | 0.162 | 0.230 | 0.112 | 0.359 | ND |
| | RE | 1.000 | 11.776 | 34.277 | 8.395 | 0.232 | 0.012 | 0.984 | ND |
| β-arrestin2 | logR | 7.017 | 8.086 | 8.344 | 7.834 | 6.208 | 5.321 | 7.086 | 5.547 |
| | SE (logR) | 0.025 | 0.034 | 0.038 | 0.032 | 0.049 | 0.022 | 0.121 | 0.299 |
| | ΔlogR | 0.000 | 1.069 | 1.327 | 0.817 | -0.809 | -1.696 | 0.069 | -1.470 |
| | SE (ΔlogR) | 0.035 | 0.042 | 0.046 | 0.041 | 0.055 | 0.033 | 0.123 | 0.300 |
| | RE | 1.000 | 11.722 | 21.232 | 6.561 | 0.155 | 0.020 | 1.172 | 0.034 |

**Table S15.** CB1 bias factor (BF = $10^{\Delta\Delta \log R}$) calculation as described in van der Weshuizen *et al.*, using WIN55212-2 as a reference ligand. ΔΔlogR=ΔlogR(pathway1) - ΔlogR(pathway2). ND = not determined.

| pathways | | WIN55,212-2 | HU210 | CP55940 | nabilone | anandamide | 2-AG | THC | cannabinol |
|---|---|---|---|---|---|---|---|---|---|
| $G_{i2}$ - $G_{i1}$ | ΔΔlogR | 0.000 | 0.572 | 1.029 | 1.273 | 1.291 | 0.907 | 0.327 | -0.267 |
| | SE (ΔΔlogR) | 0.151 | 0.175 | 0.181 | 0.185 | 0.278 | 0.188 | 0.192 | 0.292 |
| | BF | 1.000 | 3.733 | **10.691** | **18.750** | **19.543** | 8.072 | 2.123 | 0.541 |
| $G_{i2}$ - $G_{i3}$ | ΔΔlogR | 0.000 | -0.339 | -0.435 | -0.705 | -0.464 | -0.270 | -0.252 | -0.475 |
| | SE (ΔΔlogR) | 0.172 | 0.169 | 0.168 | 0.175 | 0.188 | 0.189 | 0.235 | 0.416 |
| | BF | 1.000 | 0.458 | 0.367 | 0.197 | 0.344 | 0.537 | 0.560 | 0.335 |
| $G_{i2}$ - $G_{oA}$ | ΔΔlogR | 0.000 | 0.243 | -0.290 | -0.308 | -0.179 | 0.305 | -0.110 | 0.044 |
| | SE (ΔΔlogR) | 0.192 | 0.209 | 0.190 | 0.193 | 0.203 | 0.200 | 0.245 | 0.423 |
| | BF | 1.000 | 1.750 | 0.513 | 0.492 | 0.662 | 2.018 | 0.776 | 1.107 |
| $G_{i2}$ - $G_{oB}$ | ΔΔlogR | 0.000 | 0.039 | 0.097 | -0.265 | -0.114 | 0.275 | 0.001 | 0.252 |
| | SE (ΔΔlogR) | 0.164 | 0.174 | 0.163 | 0.168 | 0.179 | 0.171 | 0.223 | 0.305 |
| | BF | 1.000 | 1.094 | 1.250 | 0.543 | 0.769 | 1.884 | 1.002 | 1.786 |
| $G_{i2}$ - $G_z$ | ΔΔlogR | 0.000 | 0.468 | 0.058 | 0.343 | 0.277 | 0.464 | 0.715 | 0.993 |
| | SE (ΔΔlogR) | 0.206 | 0.248 | 0.218 | 0.244 | 0.328 | 0.230 | 0.300 | 0.411 |
| | BF | 1.000 | 2.938 | 1.143 | 2.203 | 1.892 | 2.911 | 5.188 | **9.840** |
| $G_{i2}$ - $G_{12}$ | ΔΔlogR | 0.000 | 0.004 | 0.072 | -0.020 | 0.220 | 0.382 | 0.793 | -1.088 |
| | SE (ΔΔlogR) | 0.143 | 0.156 | 0.145 | 0.152 | 0.172 | 0.148 | 0.299 | 0.573 |
| | BF | 1.000 | 1.009 | 1.180 | 0.955 | 1.660 | 2.410 | 6.209 | **0.082** |
| $G_{i2}$ - $G_{13}$ | ΔΔlogR | 0.000 | 0.134 | 0.127 | 0.036 | -0.188 | 0.245 | 0.366 | -0.059 |
| | SE (ΔΔlogR) | 0.136 | 0.141 | 0.137 | 0.141 | 0.148 | 0.141 | 0.209 | 0.281 |
| | BF | 1.000 | 1.361 | 1.340 | 1.086 | 0.649 | 1.758 | 2.323 | 0.873 |
| $G_{i2}$ - $G_{15}$ | ΔΔlogR | 0.000 | 0.045 | -0.180 | -0.293 | -0.032 | 0.132 | -0.131 | ND |
| | SE (ΔΔlogR) | 0.174 | 0.241 | 0.181 | 0.193 | 0.242 | 0.176 | 0.554 | ND |
| | BF | 1.000 | 1.109 | 0.661 | 0.509 | 0.929 | 1.355 | 0.740 | ND |
| $G_{i2}$ - β-arrestin1 | ΔΔlogR | 0.000 | 0.249 | -0.046 | -0.087 | -0.107 | 0.920 | 0.432 | ND |
| | SE (ΔΔlogR) | 0.174 | 0.201 | 0.223 | 0.204 | 0.263 | 0.170 | 0.395 | ND |
| | BF | 1.000 | 1.774 | 0.899 | 0.818 | 0.782 | 8.318 | 2.704 | ND |
| $G_{i2}$ - β-arrestin2 | ΔΔlogR | 0.000 | 0.251 | 0.162 | 0.020 | 0.068 | 0.701 | 0.356 | 0.754 |
| | SE (ΔΔlogR) | 0.126 | 0.128 | 0.128 | 0.130 | 0.139 | 0.131 | 0.207 | 0.382 |
| | BF | 1.000 | 1.782 | 1.452 | 1.047 | 1.169 | 5.023 | 2.270 | 5.675 |
| $G_{oA}$ - $G_{oB}$ | ΔΔlogR | 0.000 | -0.204 | 0.387 | 0.043 | 0.065 | -0.030 | 0.111 | 0.208 |
| | SE (ΔΔlogR) | 0.186 | 0.212 | 0.184 | 0.187 | 0.202 | 0.192 | 0.233 | 0.400 |
| | BF | 1.000 | 0.625 | 2.438 | 1.104 | 1.161 | 0.933 | 1.291 | 1.614 |
| $G_{12}$ - $G_{13}$ | ΔΔlogR | 0.000 | 0.130 | 0.055 | 0.056 | -0.408 | -0.137 | -0.427 | 1.029 |
| | SE (ΔΔlogR) | 0.099 | 0.123 | 0.105 | 0.110 | 0.138 | 0.098 | 0.279 | 0.543 |

| | | | | | | | | | |
|---|---|---|---|---|---|---|---|---|---|
| | BF | 1.000 | 1.349 | 1.135 | 1.138 | 0.391 | 0.729 | 0.374 | **10.691** |
| β-arrestin1 – β-arrestin2 | ΔΔlogR | 0.000 | 0.002 | 0.208 | 0.107 | 0.175 | -0.219 | -0.076 | ND |
| | SE (ΔΔlogR) | 0.130 | 0.166 | 0.193 | 0.167 | 0.236 | 0.117 | 0.379 | ND |
| | BF | 1.000 | 1.005 | 1.614 | 1.279 | 1.496 | 0.604 | 0.839 | ND |

**Table S16.** Calculation of CB2 relative effectiveness (RE = $10^{\Delta\log R}$) as described in van der Westhuizen *et al.*, using WIN55212-2 as a reference ligand. NC = not converged, ND = not determined. ΔlogR = logR(ligand) - logR(reference).

| ligand | | $G_{i1}$ | $G_{i2}$ | $G_{i3}$ | $G_{oA}$ | $G_{oB}$ | β-arrestin1 | β-arrestin2 |
|---|---|---|---|---|---|---|---|---|
| WIN55,212-2 | logR | 9.409 | 9.414 | 9.318 | 9.061 | 9.202 | 9.062 | 9.259 |
| | SE (logR) | 0.075 | 0.063 | 0.062 | 0.059 | 0.060 | 0.152 | 0.081 |
| | ΔlogR | 0.000 | 0.000 | 0.000 | 0.000 | 0.000 | 0.000 | 0.000 |
| | SE (ΔlogR) | 0.106 | 0.089 | 0.088 | 0.083 | 0.084 | 0.215 | 0.115 |
| | RE | 1.000 | 1.000 | 1.000 | 1.000 | 1.000 | 1.000 | 1.000 |
| HU308 | logR | 8.131 | 8.008 | 7.912 | 7.784 | 7.914 | 7.242 | 7.400 |
| | SE (logR) | 0.095 | 0.088 | 0.058 | 0.061 | 0.092 | 0.199 | 0.089 |
| | ΔlogR | -1.278 | -1.406 | -1.406 | -1.277 | -1.288 | -1.820 | -1.859 |
| | SE (ΔlogR) | 0.121 | 0.109 | 0.085 | 0.085 | 0.109 | 0.250 | 0.121 |
| | RE | 0.053 | 0.039 | 0.039 | 0.053 | 0.052 | 0.015 | 0.014 |
| HU210 | logR | 9.314 | 9.467 | 9.210 | 9.076 | 9.457 | 9.103 | 9.043 |
| | SE (logR) | 0.081 | 0.063 | 0.062 | 0.070 | 0.063 | 0.160 | 0.107 |
| | ΔlogR | -0.095 | 0.053 | -0.108 | 0.015 | 0.255 | 0.041 | -0.216 |
| | SE (ΔlogR) | 0.110 | 0.089 | 0.088 | 0.092 | 0.087 | 0.221 | 0.134 |
| | RE | 0.804 | 1.130 | 0.780 | 1.035 | 1.799 | 1.099 | 0.608 |
| CP55940 | logR | 9.795 | 9.824 | 9.944 | 9.553 | 9.830 | 9.268 | 9.311 |
| | SE (logR) | 0.105 | 0.065 | 0.058 | 0.059 | 0.058 | 0.124 | 0.067 |
| | ΔlogR | 0.386 | 0.410 | 0.626 | 0.492 | 0.628 | 0.206 | 0.052 |
| | SE (ΔlogR) | 0.129 | 0.091 | 0.085 | 0.083 | 0.083 | 0.196 | 0.105 |
| | RE | 2.432 | 2.570 | 4.227 | 3.105 | 4.246 | 1.607 | 1.127 |
| RO6843766 | logR | 9.089 | 8.768 | 8.877 | 8.756 | 9.028 | 9.277 | 8.892 |
| | SE (logR) | 0.114 | 0.092 | 0.095 | 0.148 | 0.081 | 0.444 | 0.234 |
| | ΔlogR | -0.320 | -0.646 | -0.441 | -0.305 | -0.174 | 0.215 | -0.367 |
| | SE (ΔlogR) | 0.137 | 0.112 | 0.113 | 0.159 | 0.101 | 0.470 | 0.248 |
| | RE | 0.479 | 0.226 | 0.362 | 0.495 | 0.670 | 1.641 | 0.430 |
| RO6853457 | logR | 8.111 | 8.175 | 7.823 | 7.793 | 7.647 | 7.346 | 7.064 |
| | SE (logR) | 0.140 | 0.101 | 0.183 | 0.206 | 0.107 | 0.416 | 0.229 |
| | ΔlogR | -1.298 | -1.239 | -1.495 | -1.268 | -1.555 | -1.716 | -2.195 |
| | SE (ΔlogR) | 0.158 | 0.119 | 0.194 | 0.214 | 0.123 | 0.443 | 0.243 |
| | RE | 0.050 | 0.058 | 0.032 | 0.054 | 0.028 | 0.019 | 0.006 |
| RO7032019 | logR | 7.653 | 8.227 | 7.791 | 7.858 | 8.217 | 7.491 | 7.532 |
| | SE (logR) | 0.487 | 0.158 | 0.202 | 0.215 | 0.114 | 0.529 | 0.328 |
| | ΔlogR | -1.756 | -1.187 | -1.527 | -1.203 | -0.985 | -1.571 | -1.727 |
| | SE (ΔlogR) | 0.492 | 0.170 | 0.211 | 0.223 | 0.129 | 0.550 | 0.338 |
| | RE | 0.018 | 0.065 | 0.030 | 0.063 | 0.104 | 0.027 | 0.019 |
| RO6871487 | logR | 7.569 | 7.628 | 7.831 | 7.853 | 7.567 | 9.173 | 6.954 |
| | SE (logR) | 0.194 | 0.128 | 0.342 | 0.293 | 0.159 | 0.490 | 0.663 |
| | ΔlogR | -1.840 | -1.786 | -1.487 | -1.208 | -1.635 | 0.111 | -2.305 |

| | | | | | | | | |
|---|---|---|---|---|---|---|---|---|
| | SE (ΔlogR) | 0.208 | 0.143 | 0.347 | 0.298 | 0.170 | 0.513 | 0.668 |
| | RE | 0.014 | 0.016 | 0.033 | 0.062 | 0.023 | 1.291 | 0.005 |
| RO6878558 | logR | 7.700 | 7.654 | 7.473 | 7.613 | 7.579 | 7.834 | 6.853 |
| | SE (logR) | 0.124 | 0.100 | 0.154 | 0.172 | 0.082 | 0.553 | 0.349 |
| | ΔlogR | -1.709 | -1.760 | -1.845 | -1.448 | -1.623 | -1.228 | -2.406 |
| | SE (ΔlogR) | 0.144 | 0.118 | 0.166 | 0.182 | 0.102 | 0.573 | 0.358 |
| | RE | 0.020 | 0.017 | 0.014 | 0.036 | 0.024 | 0.059 | 0.004 |
| RO6883666 | logR | 7.632 | 7.242 | 7.186 | 8.010 | 7.399 | NC | NC |
| | SE (logR) | 0.170 | 0.124 | 0.363 | 0.441 | 0.136 | ND | ND |
| | ΔlogR | -1.777 | -2.172 | -2.132 | -1.051 | -1.803 | ND | ND |
| | SE (ΔlogR) | 0.186 | 0.139 | 0.368 | 0.445 | 0.149 | ND | ND |
| | RE | 0.017 | 0.007 | 0.007 | 0.089 | 0.016 | ND | ND |
| RO6853973 | logR | 8.595 | 8.754 | 8.409 | 8.294 | 8.564 | 10.010 | 8.202 |
| | SE (logR) | 0.186 | 0.201 | 0.486 | 0.401 | 0.218 | 0.621 | 1.082 |
| | ΔlogR | -0.814 | -0.660 | -0.909 | -0.767 | -0.638 | 0.948 | -1.057 |
| | SE (ΔlogR) | 0.200 | 0.211 | 0.490 | 0.405 | 0.226 | 0.640 | 1.085 |
| | RE | 0.153 | 0.219 | 0.123 | 0.171 | 0.230 | 8.872 | 0.088 |
| RO6850007 | logR | 7.702 | 7.536 | 7.192 | 7.949 | 8.000 | 7.764 | 7.958 |
| | SE (logR) | 0.248 | 0.216 | 0.314 | 0.419 | 0.198 | 0.476 | 0.445 |
| | ΔlogR | -1.707 | -1.878 | -2.126 | -1.112 | -1.202 | -1.298 | -1.301 |
| | SE (ΔlogR) | 0.259 | 0.225 | 0.320 | 0.423 | 0.206 | 0.500 | 0.453 |
| | RE | 0.020 | 0.013 | 0.007 | 0.077 | 0.063 | 0.050 | 0.050 |
| RO6844395 | logR | 7.560 | 7.275 | 7.639 | 7.443 | 7.481 | NC | 7.056 |
| | SE (logR) | 0.141 | 0.087 | 0.160 | 0.129 | 0.078 | ND | 0.242 |
| | ΔlogR | -1.849 | -2.139 | -1.679 | -1.618 | -1.721 | ND | -2.203 |
| | SE (ΔlogR) | 0.159 | 0.108 | 0.171 | 0.142 | 0.098 | ND | 0.255 |
| | RE | 0.014 | 0.007 | 0.021 | 0.024 | 0.019 | ND | 0.006 |
| RO6844112 | logR | 8.462 | 8.286 | 7.930 | 7.875 | 8.306 | 7.325 | 7.696 |
| | SE (logR) | 0.098 | 0.076 | 0.096 | 0.124 | 0.071 | 0.315 | 0.198 |
| | ΔlogR | -0.947 | -1.128 | -1.388 | -1.186 | -0.896 | -1.737 | -1.563 |
| | SE (ΔlogR) | 0.123 | 0.098 | 0.115 | 0.138 | 0.093 | 0.350 | 0.214 |
| | RE | 0.113 | 0.074 | 0.041 | 0.065 | 0.127 | 0.018 | 0.027 |
| RO5135445 | logR | 8.401 | 8.705 | 7.951 | 8.301 | 8.188 | 7.866 | 8.667 |
| | SE (logR) | 0.145 | 0.110 | 0.157 | 0.207 | 0.105 | 0.357 | 0.473 |
| | ΔlogR | -1.008 | -0.709 | -1.367 | -0.760 | -1.014 | -1.196 | -0.592 |
| | SE (ΔlogR) | 0.163 | 0.127 | 0.169 | 0.215 | 0.120 | 0.388 | 0.480 |
| | RE | 0.098 | 0.195 | 0.043 | 0.174 | 0.097 | 0.064 | 0.256 |
| RO6926274 | logR | 7.902 | 8.338 | 7.514 | 7.669 | 7.810 | 10.460 | NC |
| | SE (logR) | 0.309 | 0.206 | 0.294 | 0.430 | 0.345 | 0.398 | ND |
| | ΔlogR | -1.507 | -1.076 | -1.804 | -1.392 | -1.392 | 1.398 | ND |
| | SE (ΔlogR) | 0.318 | 0.215 | 0.301 | 0.434 | 0.350 | 0.426 | ND |
| | RE | 0.031 | 0.084 | 0.016 | 0.041 | 0.041 | 25.003 | ND |
| RO6892033 | logR | 8.058 | 8.124 | 8.258 | 7.658 | 7.822 | 8.129 | 6.689 |
| | SE (logR) | 0.199 | 0.128 | 0.214 | 0.195 | 0.131 | 0.451 | 1.076 |
| | ΔlogR | -1.351 | -1.290 | -1.060 | -1.403 | -1.380 | -0.933 | -2.570 |

| | | | | | | | | |
|---|---|---|---|---|---|---|---|---|
| | SE (ΔlogR) | 0.212 | 0.142 | 0.223 | 0.204 | 0.144 | 0.476 | 1.079 |
| | RE | 0.045 | 0.051 | 0.087 | 0.040 | 0.042 | 0.117 | 0.003 |
| RO6435559 | logR | 7.683 | 7.755 | 7.462 | 7.732 | 7.424 | 7.832 | 7.013 |
| | SE (logR) | 0.140 | 0.113 | 0.175 | 0.159 | 0.105 | 0.806 | 1.236 |
| | ΔlogR | -1.726 | -1.659 | -1.856 | -1.329 | -1.778 | -1.230 | -2.246 |
| | SE (ΔlogR) | 0.158 | 0.129 | 0.186 | 0.170 | 0.121 | 0.820 | 1.239 |
| | RE | 0.019 | 0.022 | 0.014 | 0.047 | 0.017 | 0.059 | 0.006 |
| RO6869094 | logR | 7.453 | 7.586 | 6.803 | 6.876 | 7.501 | 8.398 | 6.947 |
| | SE (logR) | 0.391 | 0.241 | 0.423 | 1.622 | 0.259 | 0.776 | 1.121 |
| | ΔlogR | -1.956 | -1.828 | -2.515 | -2.185 | -1.701 | -0.664 | -2.312 |
| | SE (ΔlogR) | 0.398 | 0.249 | 0.428 | 1.623 | 0.266 | 0.791 | 1.124 |
| | RE | 0.011 | 0.015 | 0.003 | 0.007 | 0.020 | 0.217 | 0.005 |
| FMP7234690 | logR | 7.731 | 8.046 | 7.406 | 7.081 | 7.811 | 7.926 | 7.518 |
| | SE (logR) | 0.127 | 0.111 | 0.092 | 0.130 | 0.084 | 0.388 | 0.185 |
| | ΔlogR | -1.678 | -1.368 | -1.912 | -1.980 | -1.391 | -1.136 | -1.741 |
| | SE (ΔlogR) | 0.147 | 0.128 | 0.111 | 0.143 | 0.103 | 0.417 | 0.202 |
| | RE | 0.021 | 0.043 | 0.012 | 0.010 | 0.041 | 0.073 | 0.018 |
| FMP7234691 | logR | 7.974 | 8.212 | 7.991 | 7.842 | 7.833 | 8.507 | 7.917 |
| | SE (logR) | 0.103 | 0.075 | 0.086 | 0.086 | 0.070 | 0.278 | 0.121 |
| | ΔlogR | -1.435 | -1.202 | -1.327 | -1.219 | -1.369 | -0.555 | -1.342 |
| | SE (ΔlogR) | 0.127 | 0.098 | 0.106 | 0.104 | 0.092 | 0.316 | 0.146 |
| | RE | 0.037 | 0.063 | 0.047 | 0.060 | 0.043 | 0.279 | 0.045 |
| FMP7234694 | logR | 7.639 | 8.024 | 7.832 | 8.097 | 8.033 | 7.486 | 7.815 |
| | SE (logR) | 0.115 | 0.074 | 0.082 | 0.082 | 0.074 | 0.425 | 0.132 |
| | ΔlogR | -1.770 | -1.390 | -1.486 | -0.964 | -1.169 | -1.576 | -1.444 |
| | SE (ΔlogR) | 0.137 | 0.097 | 0.103 | 0.101 | 0.095 | 0.451 | 0.155 |
| | RE | 0.017 | 0.041 | 0.033 | 0.109 | 0.068 | 0.027 | 0.036 |
| FMP7234698 | logR | 7.587 | 7.803 | 7.980 | 8.376 | 8.239 | 8.544 | 7.632 |
| | SE (logR) | 0.113 | 0.083 | 0.087 | 0.088 | 0.086 | 0.232 | 0.117 |
| | ΔlogR | -1.822 | -1.611 | -1.338 | -0.685 | -0.963 | -0.518 | -1.627 |
| | SE (ΔlogR) | 0.136 | 0.104 | 0.107 | 0.106 | 0.104 | 0.278 | 0.143 |
| | RE | 0.015 | 0.024 | 0.046 | 0.207 | 0.109 | 0.303 | 0.024 |
| FMP7234699 | logR | 7.672 | 7.777 | 7.658 | 8.256 | 7.704 | 7.818 | 7.717 |
| | SE (logR) | 0.129 | 0.075 | 0.099 | 0.093 | 0.071 | 0.230 | 0.148 |
| | ΔlogR | -1.737 | -1.637 | -1.660 | -0.805 | -1.498 | -1.244 | -1.542 |
| | SE (ΔlogR) | 0.149 | 0.098 | 0.117 | 0.110 | 0.093 | 0.276 | 0.169 |
| | RE | 0.018 | 0.023 | 0.022 | 0.157 | 0.032 | 0.057 | 0.029 |
| JWH133 | logR | 8.653 | 7.697 | 8.274 | 8.032 | 8.481 | 8.107 | 7.835 |
| | SE (logR) | 0.107 | 0.063 | 0.057 | 0.059 | 0.056 | 0.131 | 0.072 |
| | ΔlogR | -0.756 | -1.717 | -1.044 | -1.029 | -0.721 | -0.955 | -1.424 |
| | SE (ΔlogR) | 0.130 | 0.089 | 0.084 | 0.083 | 0.082 | 0.201 | 0.109 |
| | RE | 0.175 | 0.019 | 0.090 | 0.094 | 0.190 | 0.111 | 0.038 |
| (*rac*)-AM1241 | logR | 8.038 | 7.615 | 7.483 | 7.457 | 7.822 | 7.401 | 7.154 |
| | SE (logR) | 0.134 | 0.103 | 0.158 | 0.221 | 0.091 | 0.396 | 0.427 |
| | ΔlogR | -1.371 | -1.799 | -1.835 | -1.604 | -1.380 | -1.661 | -2.105 |

| | | | | | | | | |
|---|---|---|---|---|---|---|---|---|
| | SE (ΔlogR) | 0.153 | 0.120 | 0.169 | 0.229 | 0.109 | 0.424 | 0.435 |
| | RE | 0.043 | 0.016 | 0.015 | 0.025 | 0.042 | 0.022 | 0.008 |
| (*R*)-AM1241 | logR | 8.020 | 7.835 | 7.899 | 7.974 | 7.744 | 8.477 | 7.647 |
| | SE (logR) | 0.122 | 0.105 | 0.145 | 0.254 | 0.096 | 0.530 | 0.286 |
| | ΔlogR | -1.389 | -1.579 | -1.419 | -1.087 | -1.458 | -0.585 | -1.612 |
| | SE (ΔlogR) | 0.143 | 0.122 | 0.158 | 0.260 | 0.113 | 0.552 | 0.298 |
| | RE | 0.041 | 0.026 | 0.038 | 0.082 | 0.035 | 0.260 | 0.024 |
| (*S*)-AM1241 | logR | 7.264 | 7.120 | 7.215 | 6.812 | 7.138 | 6.862 | 6.425 |
| | SE (logR) | 0.100 | 0.073 | 0.075 | 0.096 | 0.071 | 0.280 | 0.123 |
| | ΔlogR | -2.145 | -2.294 | -2.103 | -2.249 | -2.064 | -2.200 | -2.834 |
| | SE (ΔlogR) | 0.125 | 0.097 | 0.097 | 0.113 | 0.093 | 0.318 | 0.147 |
| | RE | 0.007 | 0.005 | 0.008 | 0.006 | 0.009 | 0.006 | 0.001 |
| nabilone | logR | 8.782 | 8.920 | 8.907 | 8.283 | 8.629 | 8.445 | 8.444 |
| | SE (logR) | 0.081 | 0.077 | 0.062 | 0.059 | 0.060 | 0.176 | 0.102 |
| | ΔlogR | -0.627 | -0.494 | -0.411 | -0.778 | -0.573 | -0.617 | -0.815 |
| | SE (ΔlogR) | 0.110 | 0.100 | 0.088 | 0.083 | 0.085 | 0.232 | 0.130 |
| | RE | 0.236 | 0.321 | 0.388 | 0.167 | 0.267 | 0.242 | 0.153 |
| THC | logR | 7.556 | 7.296 | 7.229 | 7.838 | 7.541 | NC | 7.409 |
| | SE (logR) | 0.237 | 0.162 | 0.196 | 0.345 | 0.145 | ND | 0.564 |
| | ΔlogR | -1.853 | -2.118 | -2.089 | -1.223 | -1.661 | ND | -1.850 |
| | SE (ΔlogR) | 0.248 | 0.173 | 0.206 | 0.350 | 0.156 | ND | 0.570 |
| | RE | 0.014 | 0.008 | 0.008 | 0.060 | 0.022 | ND | 0.014 |
| cannabinol | logR | 7.430 | 7.039 | 6.960 | 7.895 | 7.202 | 5.678 | 5.681 |
| | SE (logR) | 0.160 | 0.126 | 0.138 | 0.191 | 0.116 | 0.975 | 0.562 |
| | ΔlogR | -1.979 | -2.375 | -2.358 | -1.166 | -2.000 | -3.384 | -3.578 |
| | SE (ΔlogR) | 0.177 | 0.141 | 0.151 | 0.200 | 0.130 | 0.987 | 0.568 |
| | RE | 0.010 | 0.004 | 0.004 | 0.068 | 0.010 | 0.000 | 0.000 |
| anandamide | logR | 7.233 | 6.935 | 6.930 | 6.808 | 7.343 | 5.164 | 5.980 |
| | SE (logR) | 0.098 | 0.077 | 0.092 | 0.118 | 0.080 | 0.103 | 0.179 |
| | ΔlogR | -2.176 | -2.479 | -2.388 | -2.253 | -1.859 | -3.898 | -3.279 |
| | SE (ΔlogR) | 0.123 | 0.100 | 0.111 | 0.132 | 0.100 | 0.183 | 0.196 |
| | RE | 0.007 | 0.003 | 0.004 | 0.006 | 0.014 | 0.000 | 0.001 |
| 2-AG | logR | 6.941 | 6.692 | 6.700 | 6.706 | 7.002 | 6.046 | 5.979 |
| | SE (logR) | 0.080 | 0.066 | 0.060 | 0.075 | 0.064 | 0.108 | 0.062 |
| | ΔlogR | -2.468 | -2.722 | -2.618 | -2.355 | -2.200 | -3.016 | -3.280 |
| | SE (ΔlogR) | 0.110 | 0.091 | 0.086 | 0.096 | 0.087 | 0.186 | 0.102 |
| | RE | 0.003 | 0.002 | 0.002 | 0.004 | 0.006 | 0.001 | 0.001 |

**Table S17.** CB2 bias factor (BF = $10^{\Delta\Delta\log R}$) calculation as described in van der Weshuizen *et al.*, using WIN55212-2 as a reference ligand. ΔΔlogR = ΔlogR(pathway1) - ΔlogR(pathway2). ND = not determined.

| ligand | | $G_{i2}$ – $G_{i1}$ | $G_{i2}$ – $G_{i3}$ | $G_{i2}$ – $G_{oA}$ | $G_{i2}$ – $G_{oB}$ | $G_{i2}$ – β-arrestin1 | $G_{i2}$ – β-arrestin2 | $G_{oB}$ – $G_{oA}$ | $G_{oB}$ – β-arrestin2 | β-arrestin1 – β-arrestin2 |
|---|---|---|---|---|---|---|---|---|---|---|
| WIN55,212-2 | ΔΔlogR | 0.000 | 0.000 | 0.000 | 0.000 | 0.000 | 0.000 | 0.000 | 0.000 | 0.000 |
| | SE (ΔΔlogR) | 0.138 | 0.125 | 0.122 | 0.123 | 0.232 | 0.146 | 0.119 | 0.143 | 0.244 |
| | BF | 1.000 | 1.000 | 1.000 | 1.000 | 1.000 | 1.000 | 1.000 | 1.000 | 1.000 |
| HU308 | ΔΔlogR | -0.128 | 0.000 | -0.129 | -0.118 | 0.414 | 0.453 | -0.011 | 0.571 | 0.039 |
| | SE (ΔΔlogR) | 0.163 | 0.138 | 0.138 | 0.154 | 0.273 | 0.163 | 0.139 | 0.163 | 0.278 |
| | BF | 0.745 | 1.000 | 0.743 | 0.762 | 2.594 | 2.838 | 0.975 | 3.724 | 1.094 |
| HU210 | ΔΔlogR | 0.148 | 0.161 | 0.038 | -0.202 | 0.012 | 0.269 | 0.240 | 0.471 | 0.257 |
| | SE (ΔΔlogR) | 0.142 | 0.125 | 0.128 | 0.124 | 0.238 | 0.161 | 0.126 | 0.160 | 0.258 |
| | BF | 1.406 | 1.449 | 1.091 | 0.628 | 1.028 | 1.858 | 1.738 | 2.958 | 1.807 |
| CP55940 | ΔΔlogR | 0.024 | -0.216 | -0.082 | -0.218 | 0.204 | 0.358 | 0.136 | 0.576 | 0.154 |
| | SE (ΔΔlogR) | 0.157 | 0.124 | 0.123 | 0.123 | 0.216 | 0.139 | 0.118 | 0.134 | 0.223 |
| | BF | 1.057 | 0.608 | 0.828 | 0.605 | 1.600 | 2.280 | 1.368 | 3.767 | 1.426 |
| RO6843766 | ΔΔlogR | -0.326 | -0.205 | -0.341 | -0.472 | -0.861 | -0.279 | 0.131 | 0.193 | 0.582 |
| | SE (ΔΔlogR) | 0.177 | 0.159 | 0.194 | 0.150 | 0.483 | 0.272 | 0.188 | 0.268 | 0.531 |
| | BF | 0.472 | 0.624 | 0.456 | 0.337 | 0.138 | 0.526 | 1.352 | 1.560 | 3.819 |
| RO6853457 | ΔΔlogR | 0.059 | 0.256 | 0.029 | 0.316 | 0.477 | 0.956 | -0.287 | 0.640 | 0.479 |
| | SE (ΔΔlogR) | 0.198 | 0.227 | 0.245 | 0.171 | 0.459 | 0.271 | 0.247 | 0.273 | 0.506 |
| | BF | 1.146 | 1.803 | 1.069 | 2.070 | 2.999 | **9.036** | 0.516 | 4.365 | 3.013 |
| RO7032019 | ΔΔlogR | 0.569 | 0.340 | 0.016 | -0.202 | 0.384 | 0.540 | 0.218 | 0.742 | 0.156 |
| | SE (ΔΔlogR) | 0.521 | 0.271 | 0.281 | 0.213 | 0.576 | 0.378 | 0.258 | 0.361 | 0.645 |
| | BF | 3.707 | 2.188 | 1.038 | 0.628 | 2.421 | 3.467 | 1.652 | 5.521 | 1.432 |
| RO6871487 | ΔΔlogR | 0.054 | -0.299 | -0.578 | -0.151 | -1.897 | 0.519 | -0.427 | 0.670 | 2.416 |
| | SE (ΔΔlogR) | 0.253 | 0.375 | 0.331 | 0.222 | 0.533 | 0.683 | 0.344 | 0.689 | 0.842 |
| | BF | 1.132 | 0.502 | 0.264 | 0.706 | **0.013** | 3.304 | 0.374 | 4.677 | **260.615** |
| RO6878558 | ΔΔlogR | -0.051 | 0.085 | -0.312 | -0.137 | -0.532 | 0.646 | -0.175 | 0.783 | 1.178 |
| | SE (ΔΔlogR) | 0.187 | 0.203 | 0.217 | 0.156 | 0.585 | 0.377 | 0.208 | 0.372 | 0.676 |
| | BF | 0.889 | 1.216 | 0.488 | 0.729 | 0.294 | 4.426 | 0.668 | 6.067 | 15.066 |
| RO[illegible] | ΔΔlogR | -0.395 | -0.040 | -1.121 | -0.369 | ND | ND | -0.752 | ND | ND |

| | | | | | | | | | | |
|---|---|---|---|---|---|---|---|---|---|---|
| | SE (ΔΔlogR) | 0.232 | 0.393 | 0.466 | 0.203 | ND | ND | 0.469 | ND | ND |
| | BF | 0.403 | 0.912 | **0.076** | 0.428 | ND | ND | 0.177 | ND | ND |
| RO6853973 | ΔΔlogR | 0.154 | 0.249 | 0.107 | -0.022 | -1.608 | 0.397 | 0.129 | 0.419 | 2.005 |
| | SE (ΔΔlogR) | 0.291 | 0.534 | 0.456 | 0.309 | 0.673 | 1.105 | 0.464 | 1.108 | 1.260 |
| | BF | 1.426 | 1.774 | 1.279 | 0.951 | **0.025** | 2.495 | 1.346 | 2.624 | 101.158 |
| RO6850007 | ΔΔlogR | -0.171 | 0.248 | -0.766 | -0.676 | -0.580 | -0.577 | -0.090 | 0.099 | 0.003 |
| | SE (ΔΔlogR) | 0.343 | 0.391 | 0.479 | 0.305 | 0.548 | 0.505 | 0.471 | 0.497 | 0.674 |
| | BF | 0.675 | 1.770 | 0.171 | 0.211 | 0.263 | 0.265 | 0.813 | 1.256 | 1.007 |
| RO6844395 | ΔΔlogR | -0.290 | -0.460 | -0.521 | -0.418 | ND | 0.064 | -0.103 | 0.482 | ND |
| | SE (ΔΔlogR) | 0.192 | 0.202 | 0.178 | 0.146 | ND | 0.277 | 0.172 | 0.273 | ND |
| | BF | 0.513 | 0.347 | 0.301 | 0.382 | ND | 1.159 | 0.789 | 3.034 | ND |
| RO6844112 | ΔΔlogR | -0.181 | 0.260 | 0.058 | -0.232 | 0.609 | 0.435 | 0.290 | 0.667 | -0.174 |
| | SE (ΔΔlogR) | 0.158 | 0.151 | 0.169 | 0.135 | 0.363 | 0.236 | 0.166 | 0.233 | 0.410 |
| | BF | 0.659 | 1.820 | 1.143 | 0.586 | 4.064 | 2.723 | 1.950 | 4.645 | 0.670 |
| RO5135445 | ΔΔlogR | 0.299 | 0.658 | 0.051 | 0.305 | 0.487 | -0.117 | -0.254 | -0.422 | -0.604 |
| | SE (ΔΔlogR) | 0.207 | 0.211 | 0.250 | 0.175 | 0.408 | 0.496 | 0.247 | 0.495 | 0.617 |
| | BF | 1.991 | 4.550 | 1.125 | 2.018 | 3.069 | 0.764 | 0.557 | 0.378 | 0.249 |
| RO6926274 | ΔΔlogR | 0.431 | 0.728 | 0.316 | 0.316 | -2.474 | ND | 0.000 | ND | ND |
| | SE (ΔΔlogR) | 0.384 | 0.370 | 0.484 | 0.411 | 0.477 | ND | 0.558 | ND | ND |
| | BF | 2.698 | 5.346 | 2.070 | 2.070 | **0.003** | ND | 1.000 | ND | ND |
| RO6892033 | ΔΔlogR | 0.061 | -0.230 | 0.113 | 0.090 | -0.357 | 1.280 | 0.023 | 1.190 | 1.637 |
| | SE (ΔΔlogR) | 0.255 | 0.264 | 0.248 | 0.202 | 0.496 | 1.088 | 0.249 | 1.089 | 1.179 |
| | BF | 1.151 | 0.589 | 1.297 | 1.230 | 0.440 | 19.055 | 1.054 | 15.488 | 43.351 |
| RO6435559 | ΔΔlogR | 0.067 | 0.197 | -0.330 | 0.119 | -0.429 | 0.587 | -0.449 | 0.468 | 1.016 |
| | SE (ΔΔlogR) | 0.204 | 0.226 | 0.213 | 0.177 | 0.830 | 1.245 | 0.209 | 1.245 | 1.485 |
| | BF | 1.167 | 1.574 | 0.468 | 1.315 | 0.372 | 3.864 | 0.356 | 2.938 | 10.375 |
| RO6869094 | ΔΔlogR | 0.128 | 0.687 | 0.357 | -0.127 | -1.164 | 0.484 | 0.484 | 0.611 | 1.648 |
| | SE (ΔΔlogR) | 0.470 | 0.495 | 1.642 | 0.365 | 0.829 | 1.151 | 1.645 | 1.155 | 1.374 |
| | BF | 1.343 | 4.864 | 2.275 | 0.746 | 0.069 | 3.048 | 3.048 | 4.083 | 44.463 |
| FMP7234690 | ΔΔlogR | 0.310 | 0.544 | 0.612 | 0.023 | -0.232 | 0.373 | 0.589 | 0.350 | 0.605 |
| | SE (ΔΔlogR) | 0.195 | 0.169 | 0.191 | 0.164 | 0.436 | 0.239 | 0.176 | 0.227 | 0.463 |
| | BF | 2.042 | 3.499 | 4.093 | 1.054 | 0.586 | 2.360 | 3.882 | 2.239 | 4.027 |
| FMP7234691 | ΔΔlogR | 0.233 | 0.125 | 0.017 | 0.167 | -0.647 | 0.140 | -0.150 | -0.027 | 0.787 |
| | SE (ΔΔlogR) | 0.161 | 0.145 | 0.143 | 0.135 | 0.331 | 0.176 | 0.139 | 0.172 | 0.348 |

| | | | | | | | | | | |
|---|---|---|---|---|---|---|---|---|---|---|
| | BF | 1.710 | 1.334 | 1.040 | 1.469 | 0.225 | 1.380 | 0.708 | 0.940 | 6.124 |
| FMP7234694 | ΔΔlogR | 0.380 | 0.096 | -0.426 | -0.221 | 0.186 | 0.054 | -0.205 | 0.275 | -0.132 |
| | SE (ΔΔlogR) | 0.168 | 0.142 | 0.140 | 0.136 | 0.462 | 0.183 | 0.139 | 0.182 | 0.477 |
| | BF | 2.399 | 1.247 | 0.375 | 0.601 | 1.535 | 1.132 | 0.624 | 1.884 | 0.738 |
| FMP7234698 | ΔΔlogR | 0.211 | -0.273 | -0.926 | -0.648 | -1.093 | 0.016 | -0.278 | 0.664 | 1.109 |
| | SE (ΔΔlogR) | 0.171 | 0.149 | 0.148 | 0.147 | 0.296 | 0.177 | 0.148 | 0.177 | 0.312 |
| | BF | 1.626 | 0.533 | 0.119 | 0.225 | **0.081** | 1.038 | 0.527 | 4.613 | **12.853** |
| FMP7234699 | ΔΔlogR | 0.100 | 0.023 | -0.832 | -0.139 | -0.393 | -0.095 | -0.693 | 0.044 | 0.298 |
| | SE (ΔΔlogR) | 0.178 | 0.153 | 0.147 | 0.135 | 0.293 | 0.195 | 0.144 | 0.193 | 0.323 |
| | BF | 1.259 | 1.054 | 0.147 | 0.726 | 0.405 | 0.804 | 0.203 | 1.107 | 1.986 |
| JWH133 | ΔΔlogR | -0.961 | -0.673 | -0.688 | -0.996 | -0.762 | -0.293 | 0.308 | 0.703 | 0.469 |
| | SE (ΔΔlogR) | 0.158 | 0.123 | 0.122 | 0.121 | 0.220 | 0.141 | 0.117 | 0.136 | 0.228 |
| | BF | **0.109** | 0.212 | 0.205 | **0.101** | **0.173** | 0.509 | 2.032 | 5.047 | 2.944 |
| (*rac*)-AM1241 | ΔΔlogR | -0.428 | 0.036 | -0.195 | -0.419 | -0.138 | 0.306 | 0.224 | 0.725 | 0.444 |
| | SE (ΔΔlogR) | 0.195 | 0.208 | 0.258 | 0.162 | 0.441 | 0.451 | 0.253 | 0.448 | 0.607 |
| | BF | 0.373 | 1.086 | 0.638 | 0.381 | 0.728 | 2.023 | 1.675 | 5.309 | 2.780 |
| (*R*)-AM1241 | ΔΔlogR | -0.190 | -0.160 | -0.492 | -0.121 | -0.994 | 0.033 | -0.371 | 0.154 | 1.027 |
| | SE (ΔΔlogR) | 0.188 | 0.199 | 0.288 | 0.167 | 0.565 | 0.322 | 0.284 | 0.318 | 0.627 |
| | BF | 0.646 | 0.692 | 0.322 | 0.757 | 0.101 | 1.079 | 0.426 | 1.426 | 10.641 |
| (*S*)-AM1241 | ΔΔlogR | -0.149 | -0.191 | -0.045 | -0.230 | -0.094 | 0.540 | 0.185 | 0.770 | 0.634 |
| | SE (ΔΔlogR) | 0.158 | 0.137 | 0.149 | 0.134 | 0.333 | 0.176 | 0.146 | 0.174 | 0.351 |
| | BF | 0.710 | 0.644 | 0.902 | 0.589 | 0.805 | 3.467 | 1.531 | 5.888 | 4.305 |
| nabilone | ΔΔlogR | 0.133 | -0.083 | 0.284 | 0.079 | 0.123 | 0.321 | 0.205 | 0.242 | 0.198 |
| | SE (ΔΔlogR) | 0.148 | 0.133 | 0.130 | 0.131 | 0.253 | 0.164 | 0.119 | 0.155 | 0.266 |
| | BF | 1.358 | 0.826 | 1.923 | 1.199 | 1.327 | 2.094 | 1.603 | 1.746 | 1.578 |
| THC | ΔΔlogR | -0.265 | -0.029 | -0.895 | -0.457 | ND | -0.268 | -0.438 | 0.189 | ND |
| | SE (ΔΔlogR) | 0.303 | 0.269 | 0.391 | 0.234 | ND | 0.596 | 0.384 | 0.591 | ND |
| | BF | 0.543 | 0.935 | 0.127 | 0.349 | ND | 0.540 | 0.365 | 1.545 | ND |
| cannabinol | ΔΔlogR | -0.396 | -0.017 | -1.209 | -0.375 | 1.009 | 1.203 | -0.834 | 1.578 | 0.194 |
| | SE (ΔΔlogR) | 0.226 | 0.207 | 0.245 | 0.192 | 0.997 | 0.585 | 0.239 | 0.582 | 1.138 |
| | BF | 0.402 | 0.962 | **0.062** | 0.422 | 10.209 | 15.959 | **0.147** | **37.844** | 1.563 |
| anandamide | ΔΔlogR | -0.303 | -0.091 | -0.226 | -0.620 | 1.419 | 0.800 | 0.394 | 1.420 | -0.619 |
| | SE (ΔΔlogR) | 0.158 | 0.149 | 0.165 | 0.141 | 0.209 | 0.220 | 0.165 | 0.220 | 0.269 |

| | | | | | | | | | | |
|---|---|---|---|---|---|---|---|---|---|---|
| | BF | 0.498 | 0.811 | 0.594 | 0.240 | **26.242** | 6.310 | 2.477 | **26.303** | 0.240 |
| 2-AG | ΔΔlogR | -0.254 | -0.104 | -0.367 | -0.522 | 0.294 | 0.558 | 0.155 | 1.080 | 0.264 |
| | SE (ΔΔlogR) | 0.142 | 0.126 | 0.132 | 0.126 | 0.207 | 0.137 | 0.129 | 0.134 | 0.212 |
| | BF | 0.557 | 0.787 | 0.430 | 0.301 | 1.968 | 3.614 | 1.429 | **12.023** | 1.837 |
