## Supplementary material for "Diverse chemotypes drive biased signaling by cannabinoid receptors": SI concentration-response curves

### CB1: THC

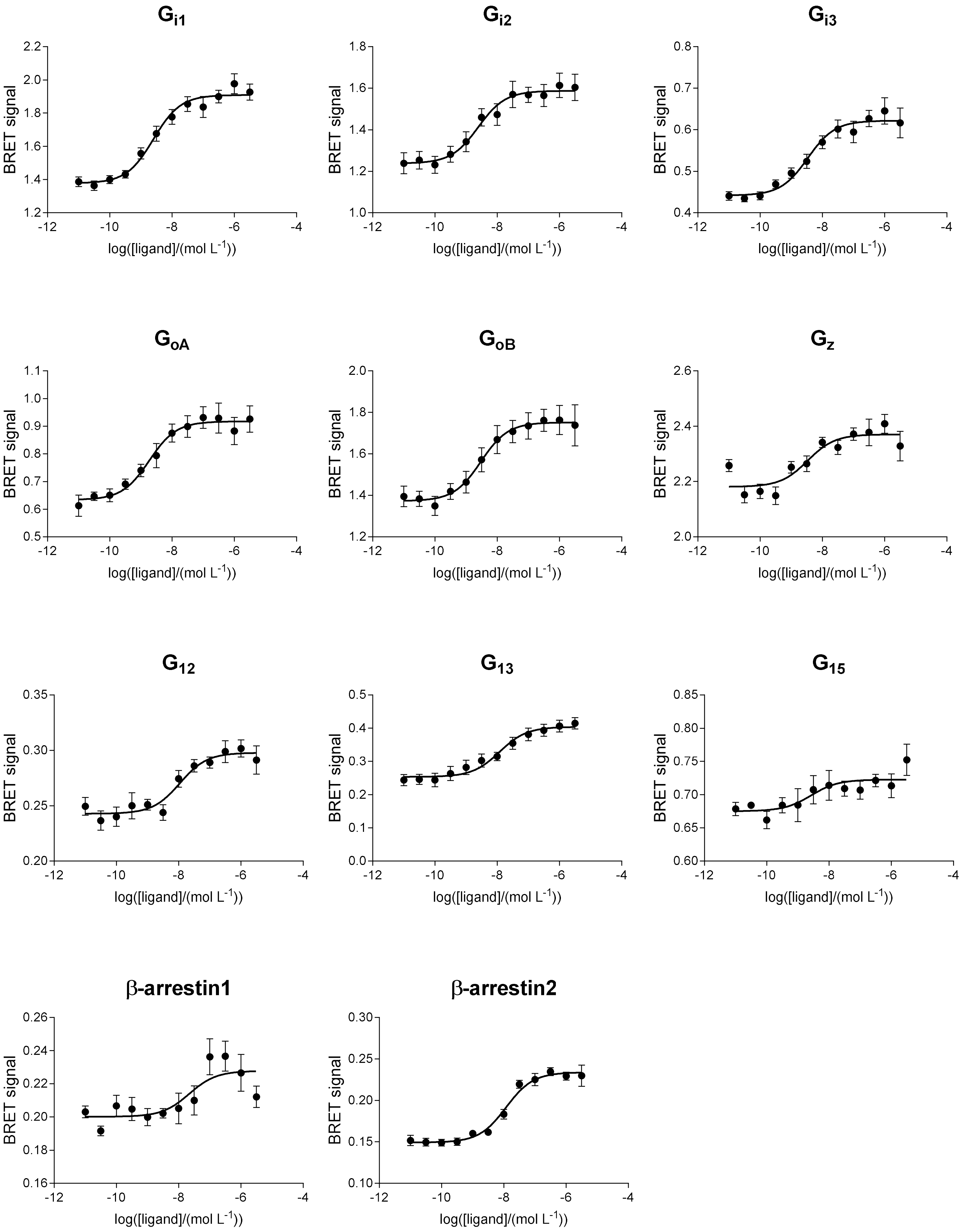

$G_{i1}$
$G_{i2}$
$G_{i3}$
$G_{oA}$
$G_{oB}$
$G_{z}$
$G_{12}$
$G_{13}$
$G_{15}$
β-arrestin1
β-arrestin2
BRET signal
log([ligand]/(mol L$^{-1}$))

### CB1: cannabinol

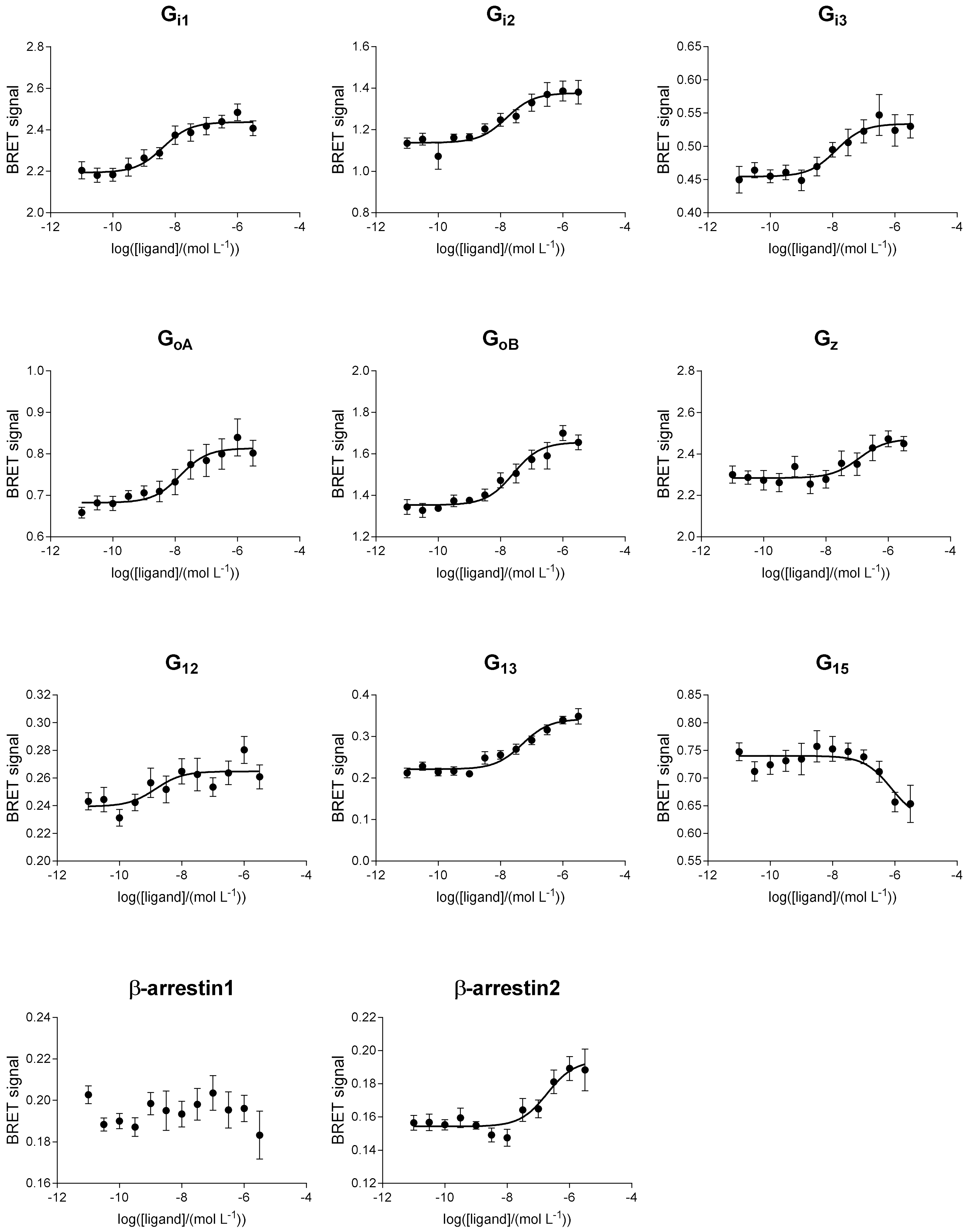

### CB1: anandamide

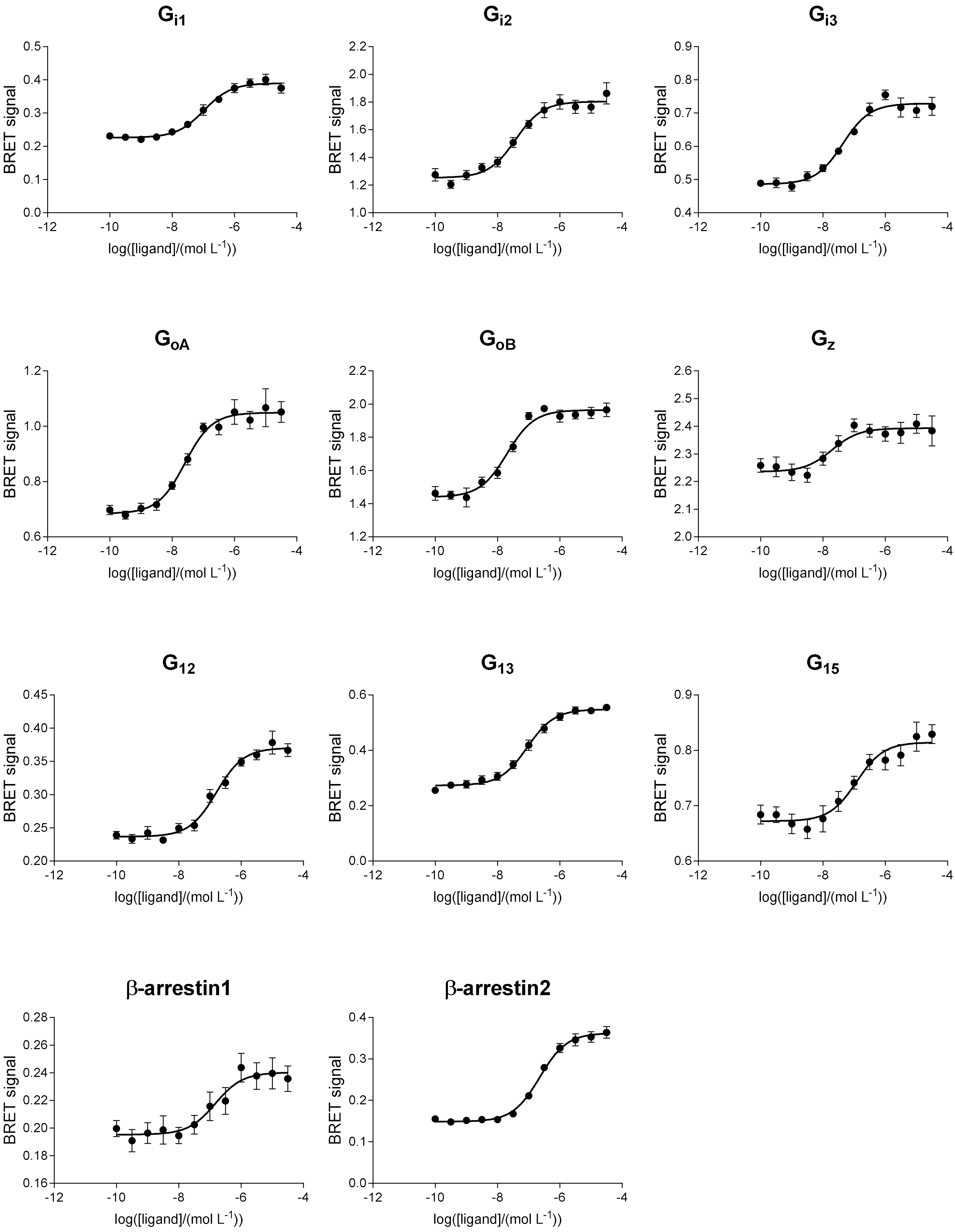

# CB1: HU210

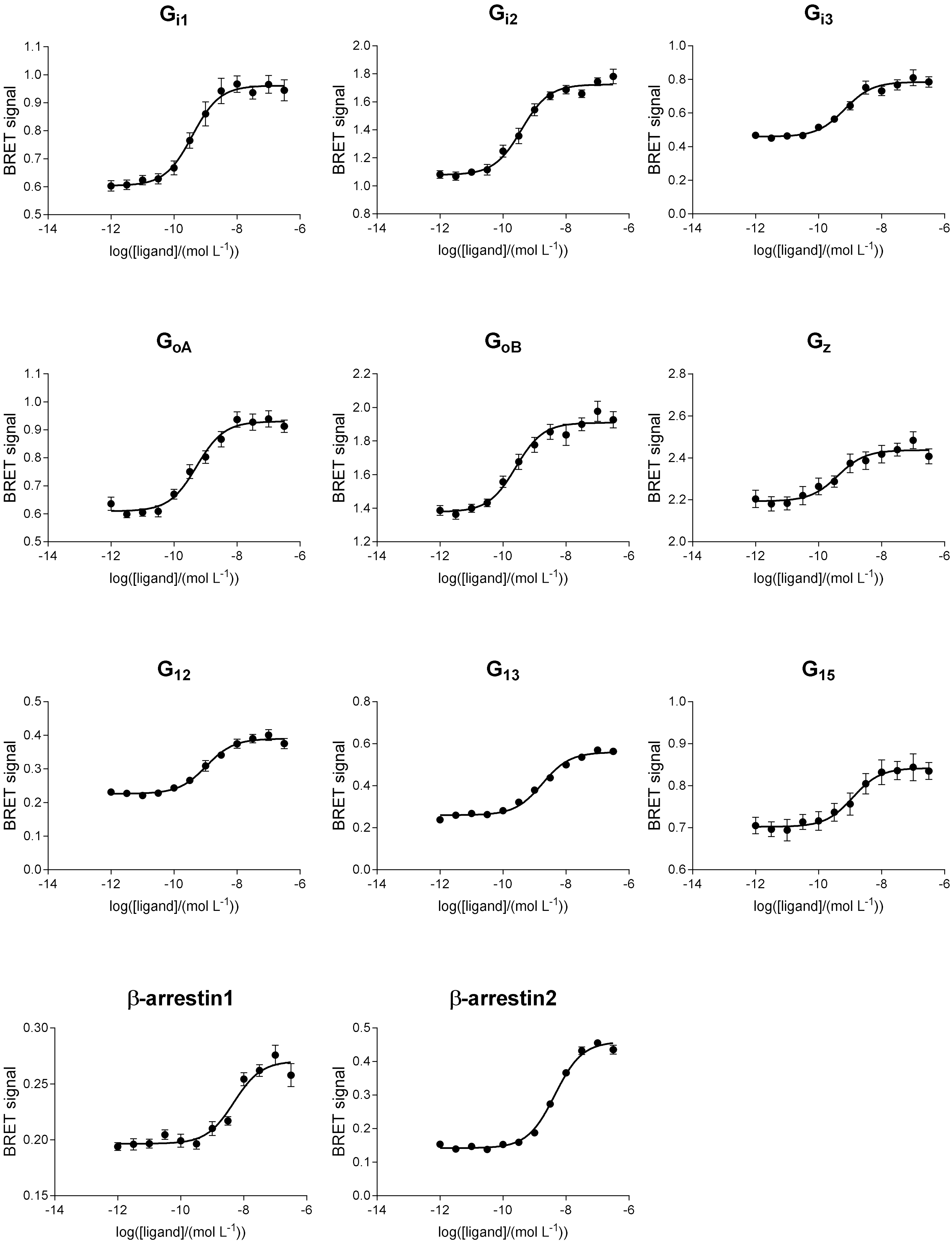

Gi1
Gi2
Gi3
GoA
GoB
Gz
G12
G13
G15
β-arrestin1
β-arrestin2
BRET signal
log([ligand]/(mol L-1))

### CB1: nabilone

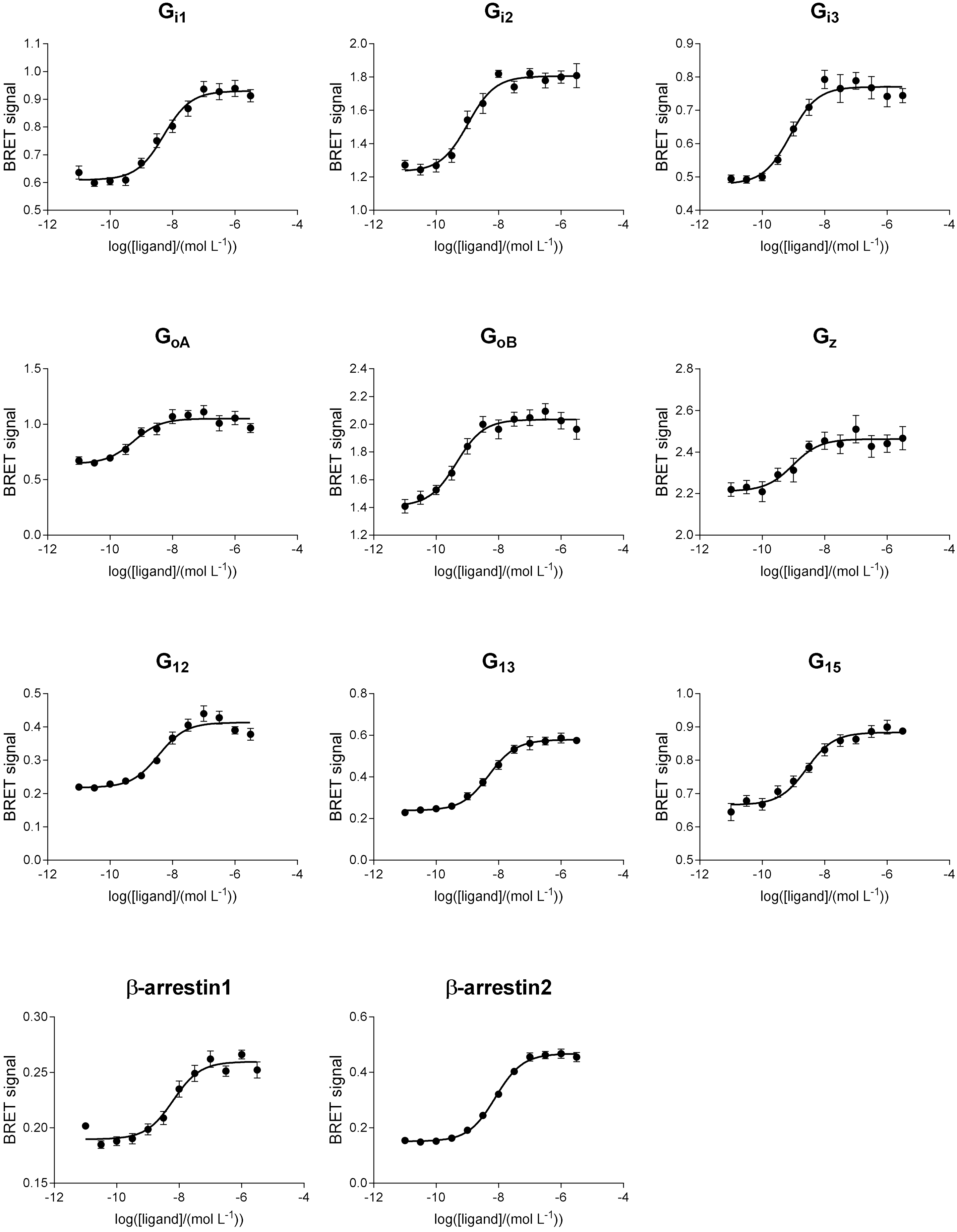

### CB1: WIN55212-2

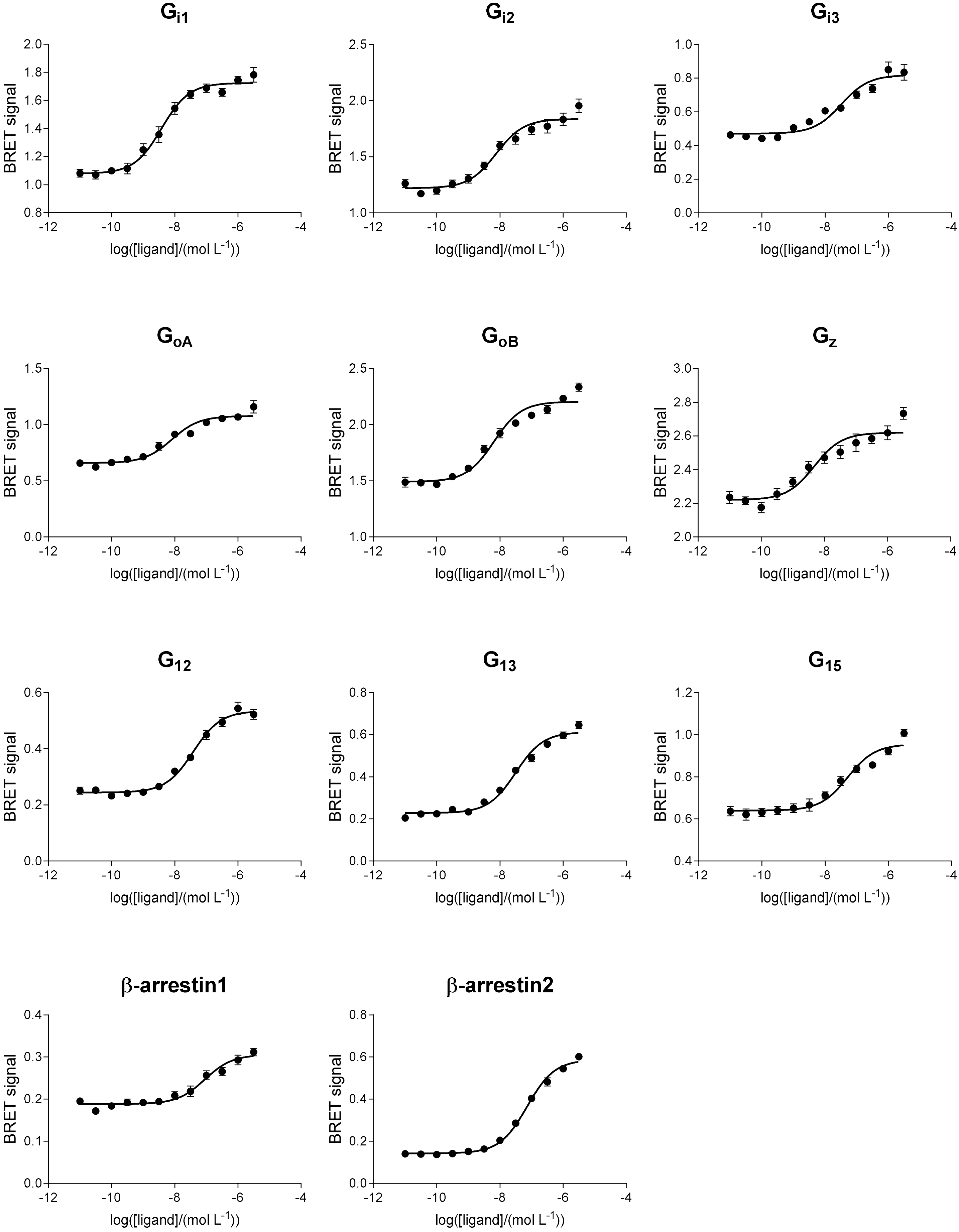

# CB1: CP55940

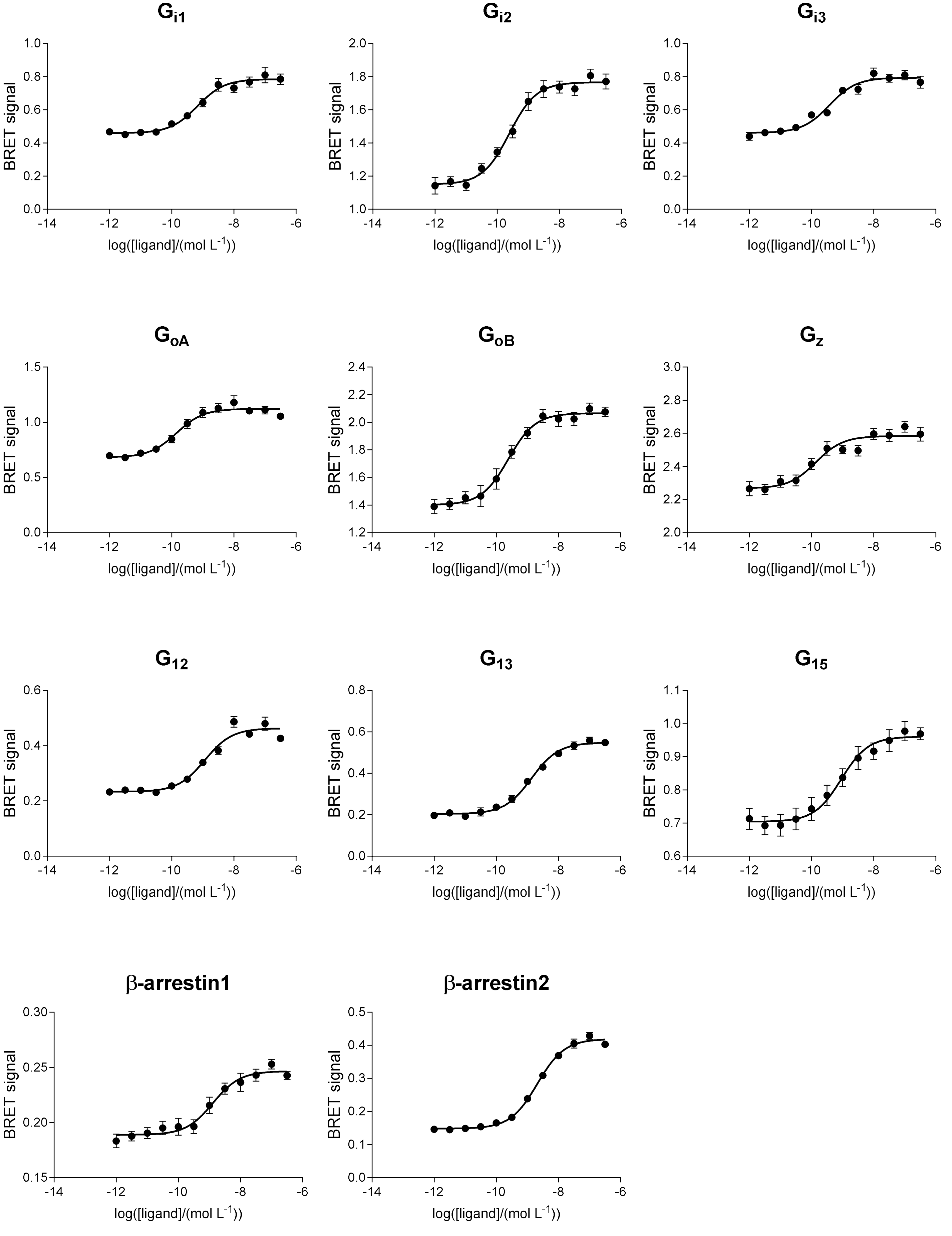

Gi1
Gi2
Gi3
GoA
GoB
Gz
G12
G13
G15
β-arrestin1
β-arrestin2
BRET signal
log([ligand]/(mol L-1))

# CB1: 2-AG

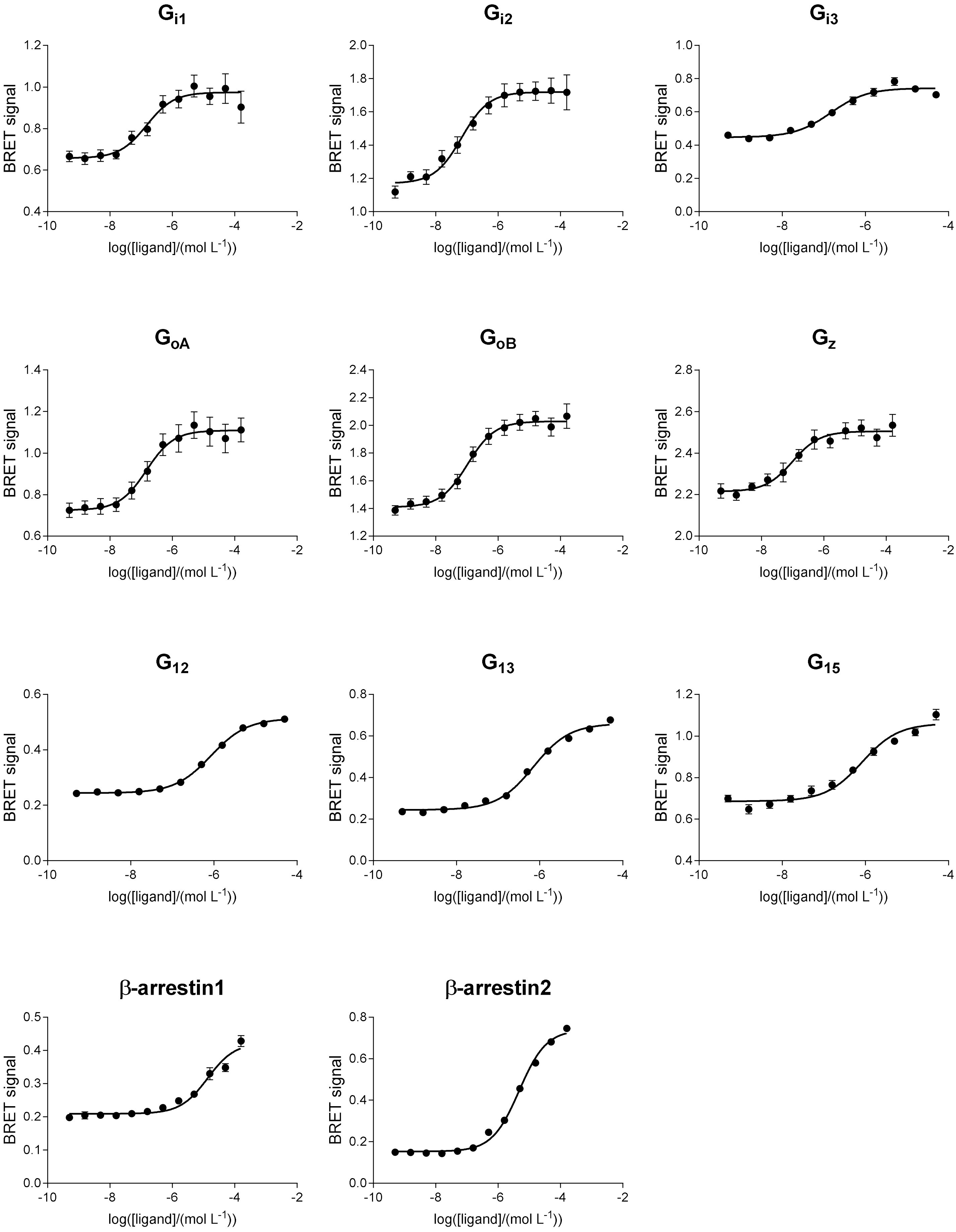

G$_{i1}$
G$_{i2}$
G$_{i3}$
G$_{oA}$
G$_{oB}$
G$_{z}$
G$_{12}$
G$_{13}$
G$_{15}$
β-arrestin1
β-arrestin2
BRET signal
log([ligand]/(mol L$^{-1}$))

# CB2: SR144528

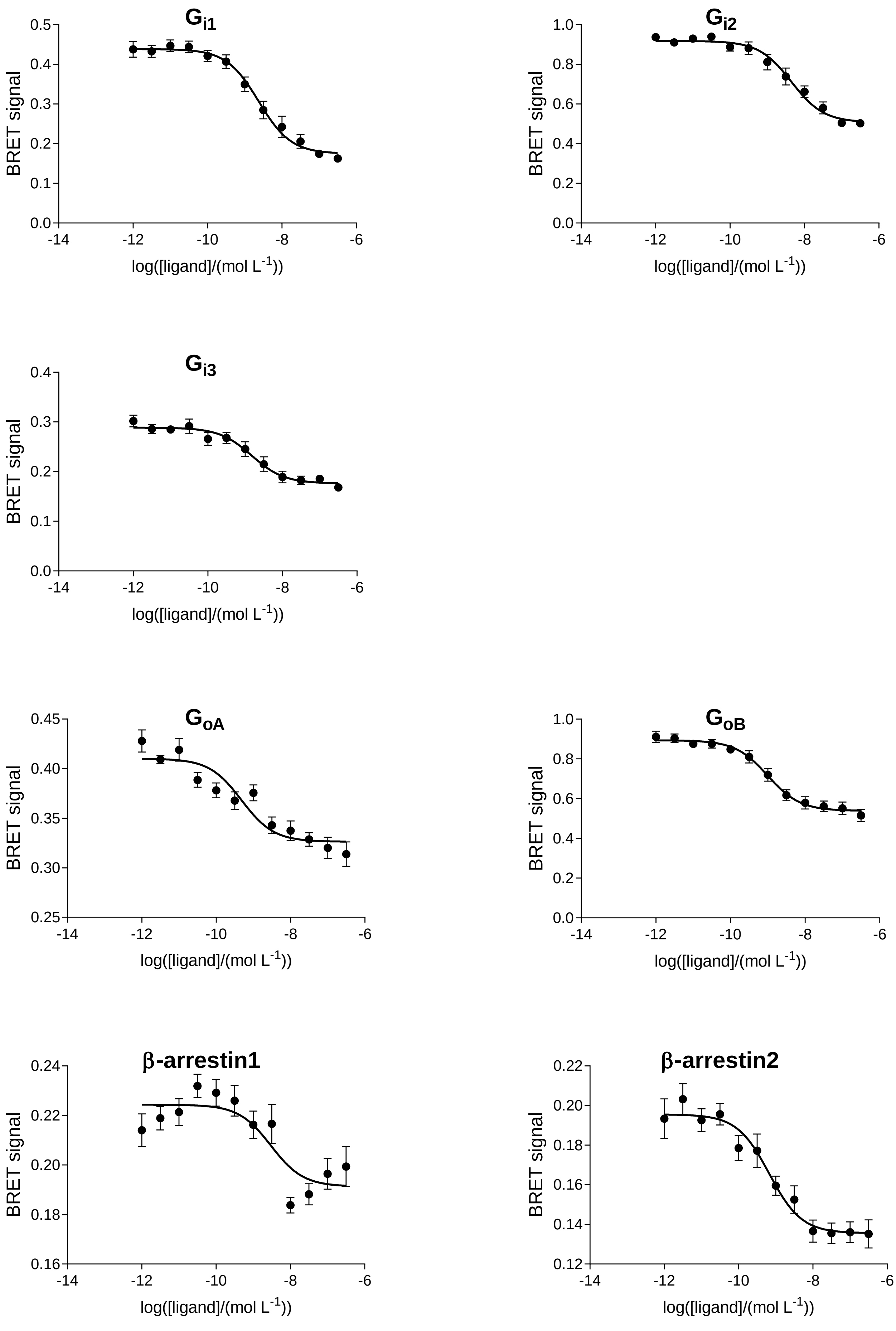

# CB2: AM630

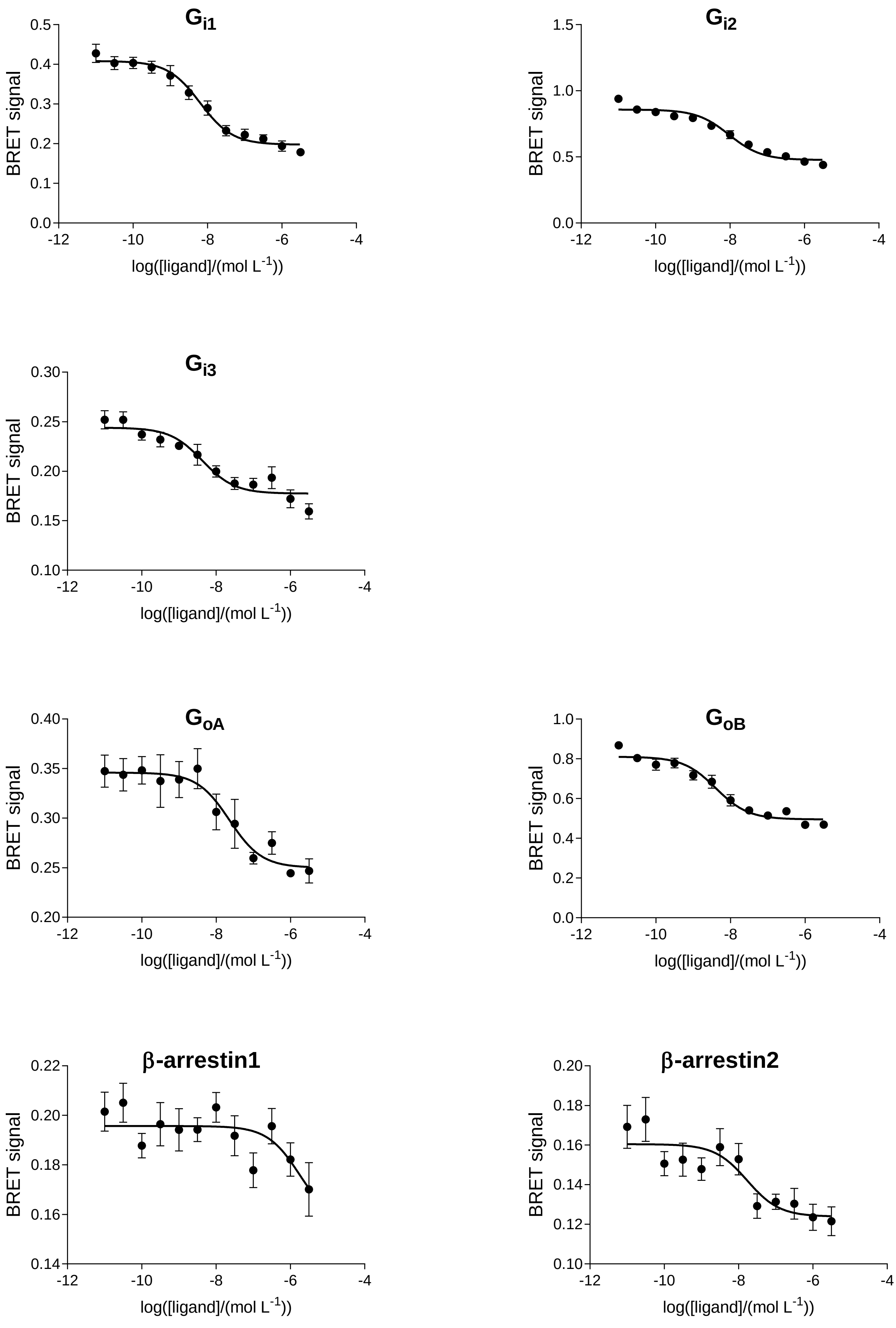

Gi1
Gi2
Gi3
GoA
GoB
β-arrestin1
β-arrestin2
BRET signal
log([ligand]/(mol L-1))

### CB2: THC

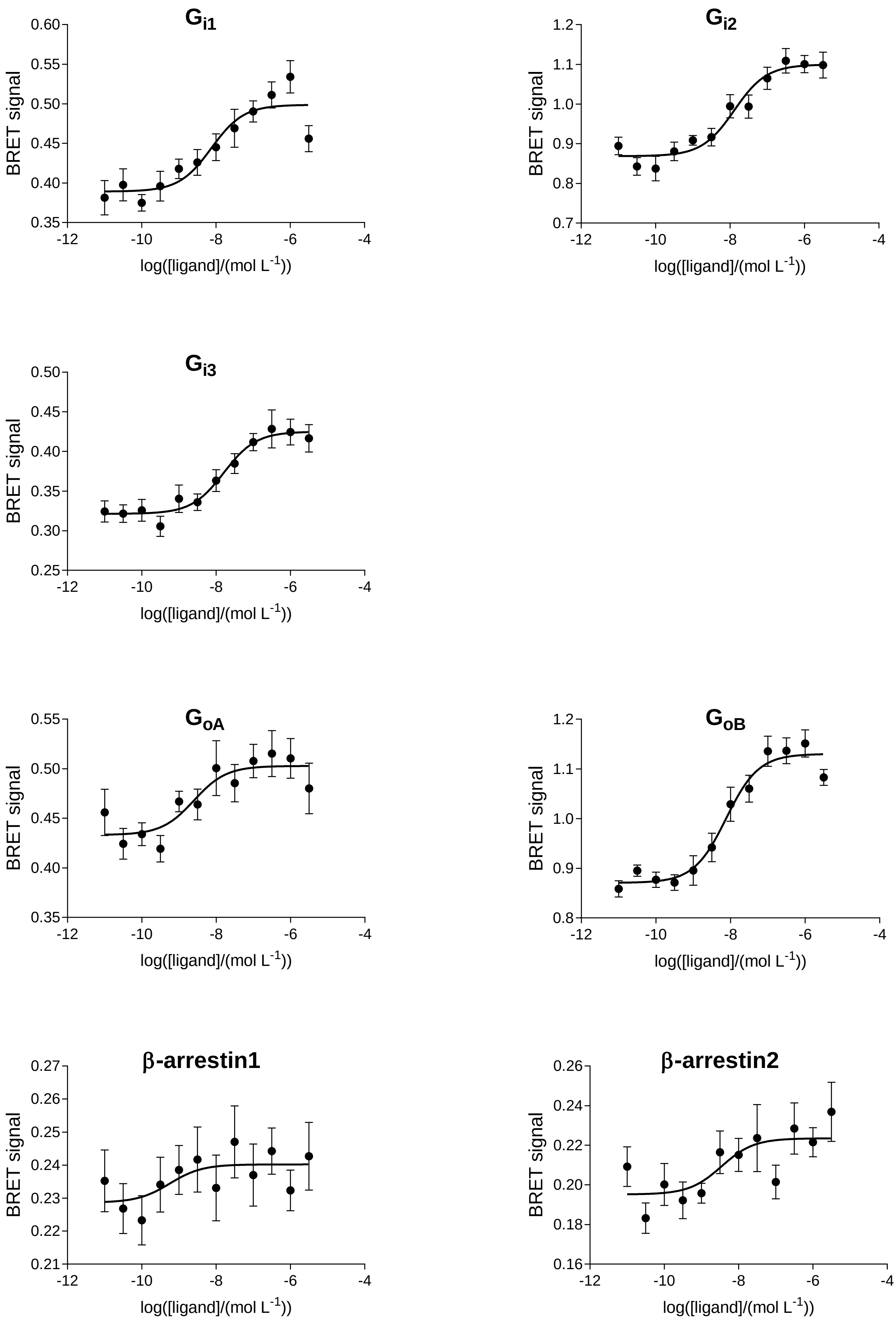

Gi1
Gi2
Gi3
GoA
GoB
β-arrestin1
β-arrestin2
BRET signal
log([ligand]/(mol L-1))

### CB2: cannabinol

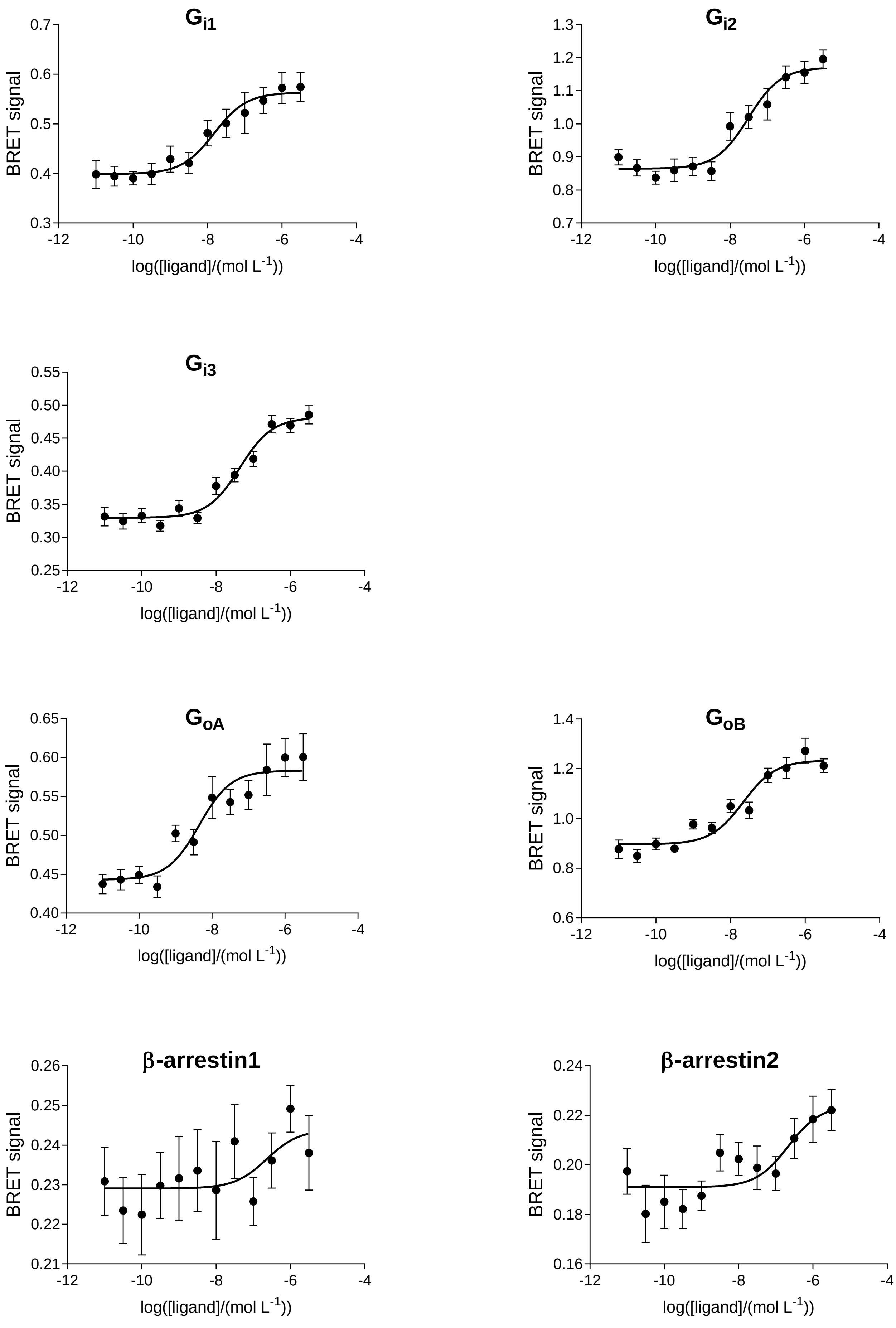

### CB2: (*rac*)-AM1241

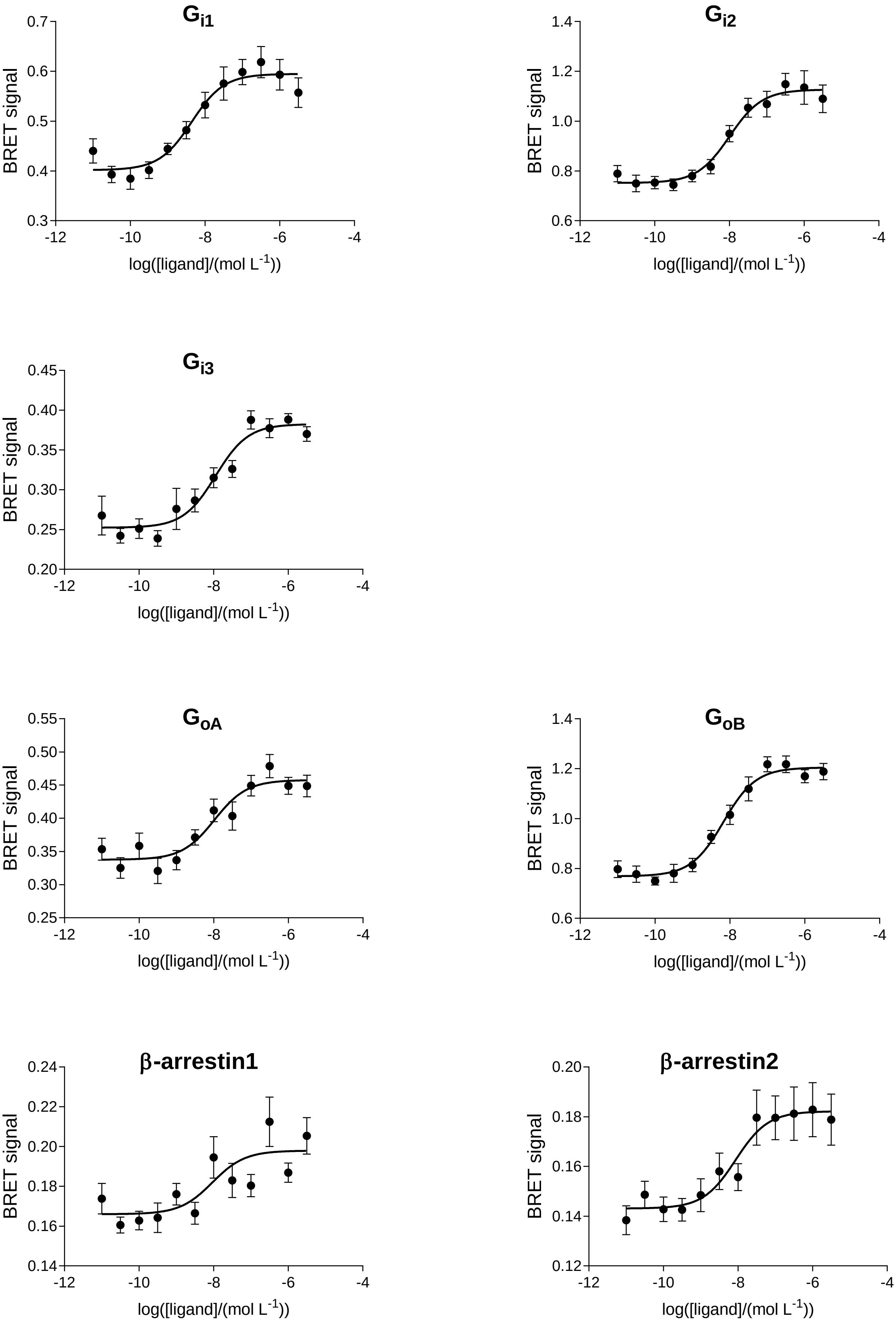

# CB2: (*R*)-AM1241

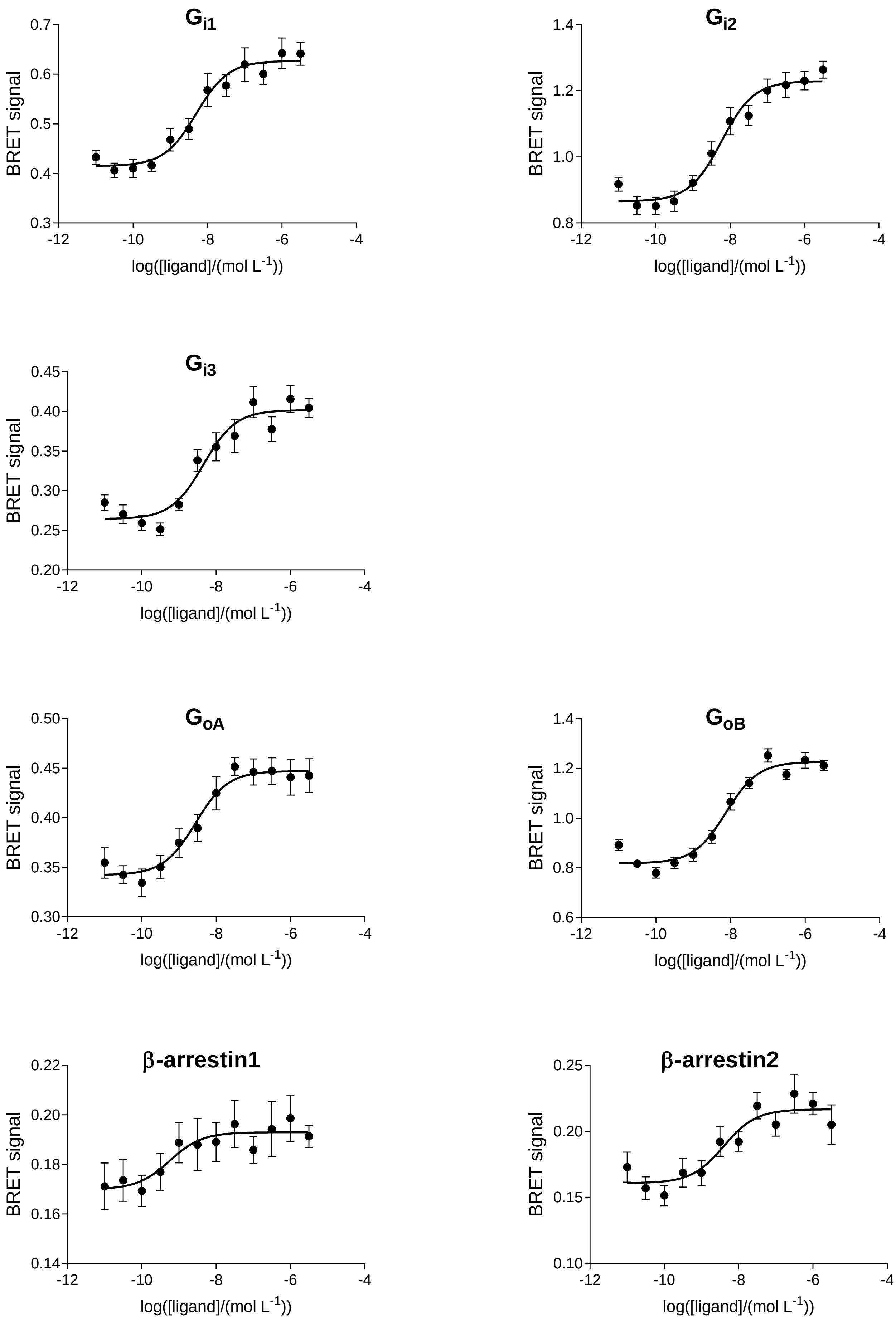

Gi1
Gi2
Gi3
GoA
GoB
β-arrestin1
β-arrestin2
BRET signal
log([ligand]/(mol L-1))
-12
-10
-8
-6
-4

# CB2: (*S*)-AM1241

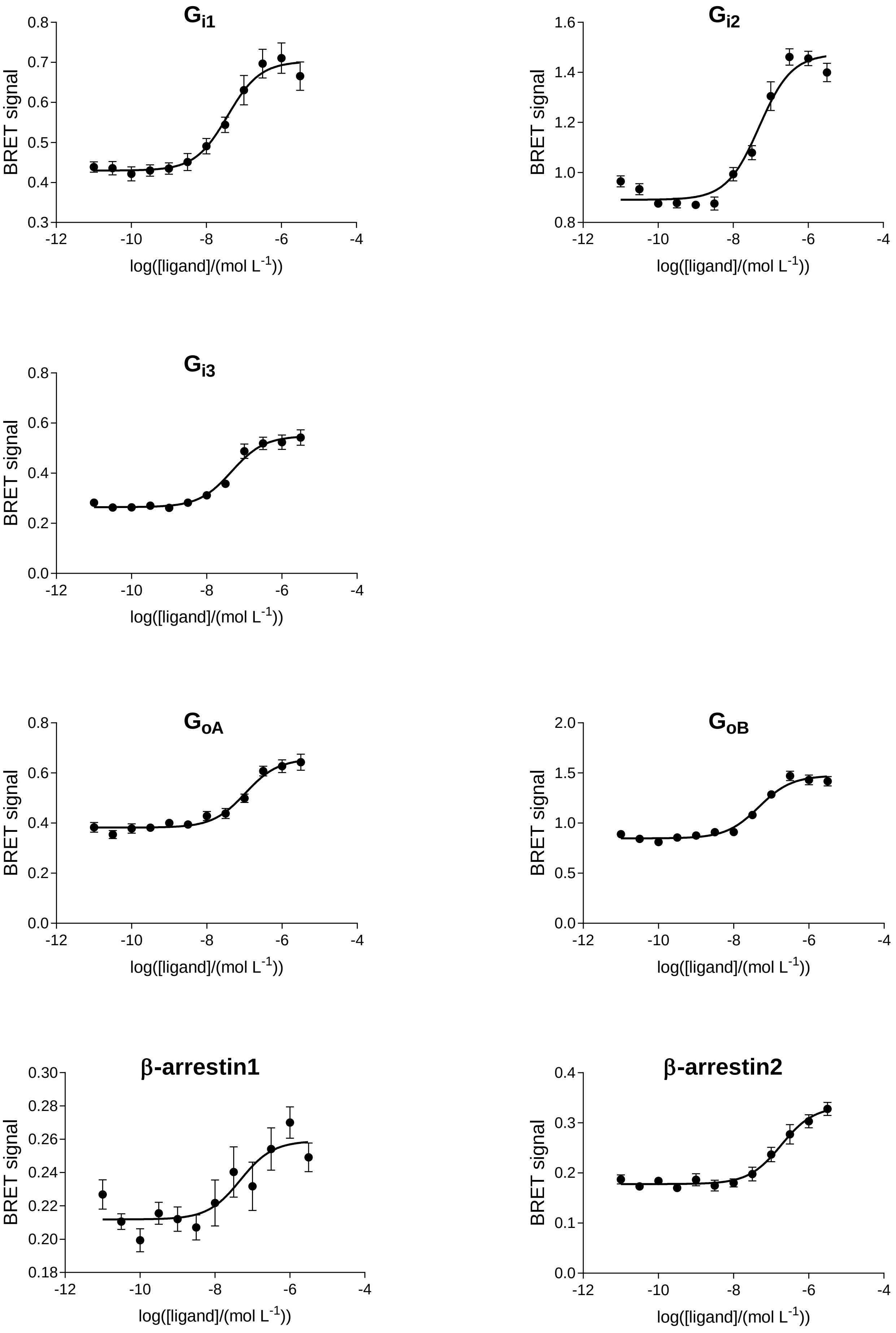

Gi1
Gi2
Gi3
GoA
GoB
β-arrestin1
β-arrestin2
BRET signal
log([ligand]/(mol L-1))
0.8
0.7
0.6
0.5
0.4
0.3
1.6
1.4
1.2
1.0
0.0
0.2
2.0
1.5
0.5
0.30
0.28
0.26
0.24
0.22
0.20
0.18
0.1
-12
-10
-8
-6
-4

### CB2: anandamide

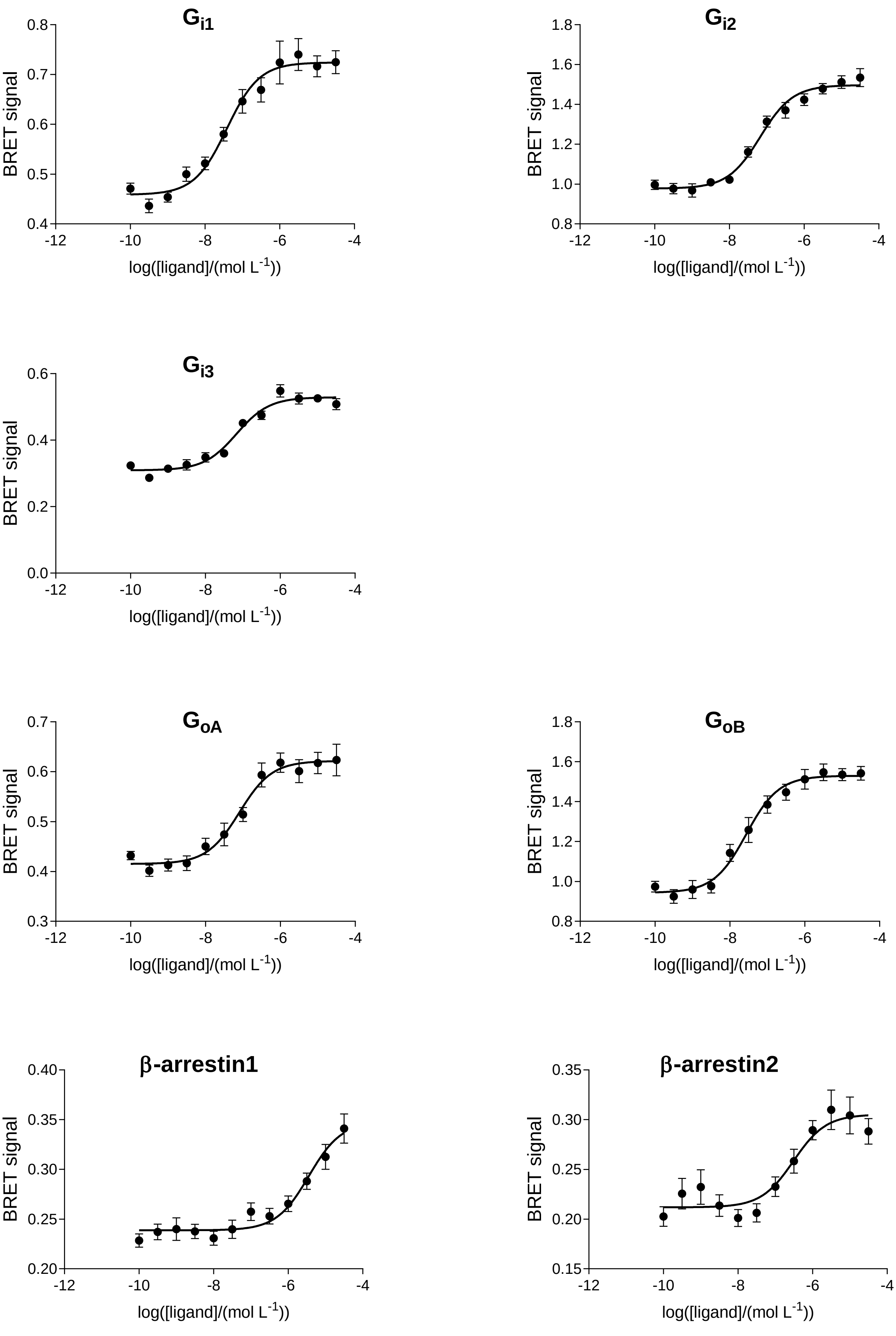

# CB2: HU210

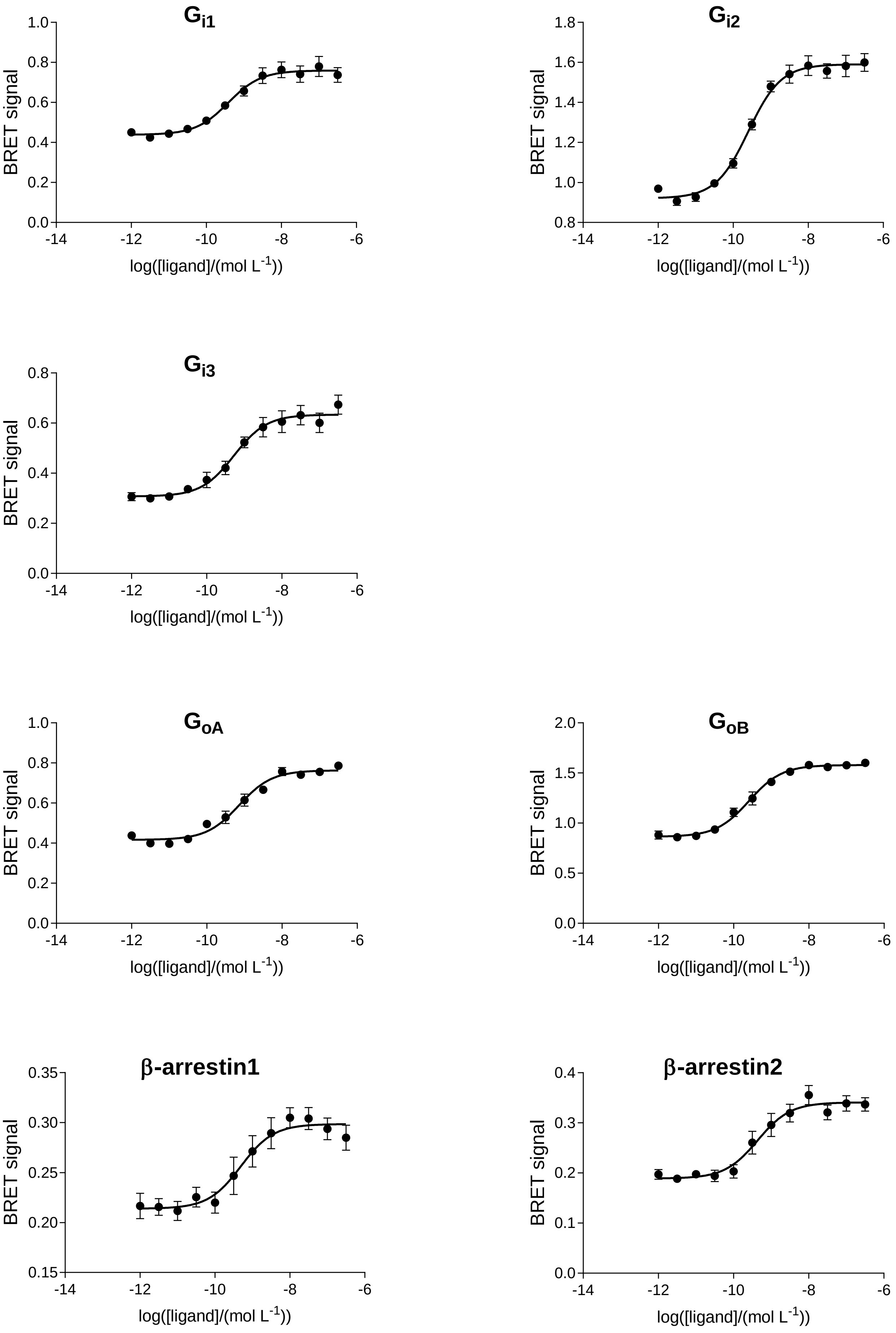

$G_{i1}$
$G_{i2}$
$G_{i3}$
$G_{oA}$
$G_{oB}$
β-arrestin1
β-arrestin2
BRET signal
log([ligand]/(mol $L^{-1}$))

### CB2: nabilone

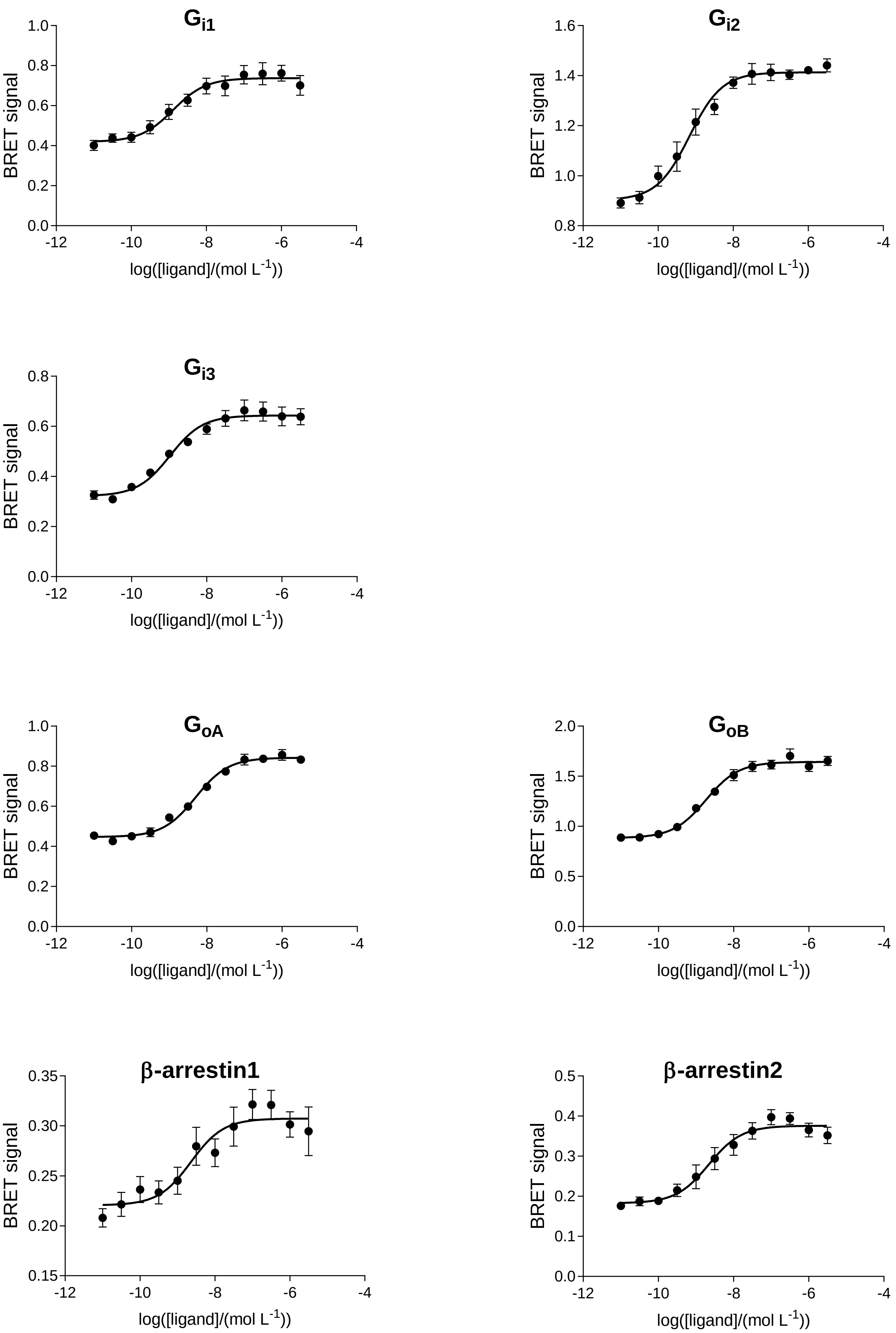

# CB2: HU308

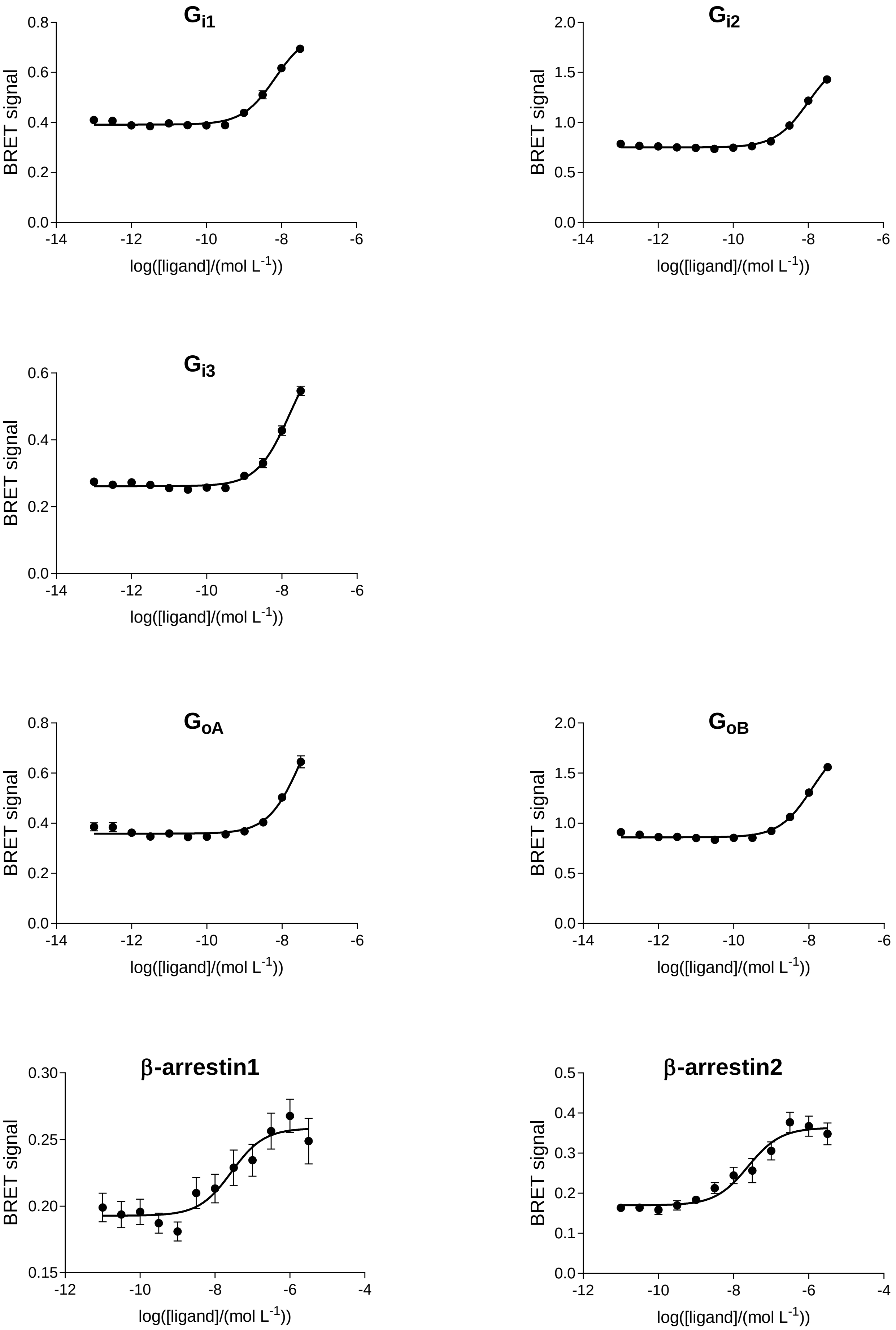

G_i1
G_i2
G_i3
G_oA
G_oB
β-arrestin1
β-arrestin2
BRET signal
log([ligand]/(mol L^-1))
0.0
0.2
0.4
0.6
0.8
0.5
1.0
1.5
2.0
0.15
0.20
0.25
0.30
0.1
0.3
0.5
-14
-12
-10
-8
-6
-4

### CB2: WIN55212-2

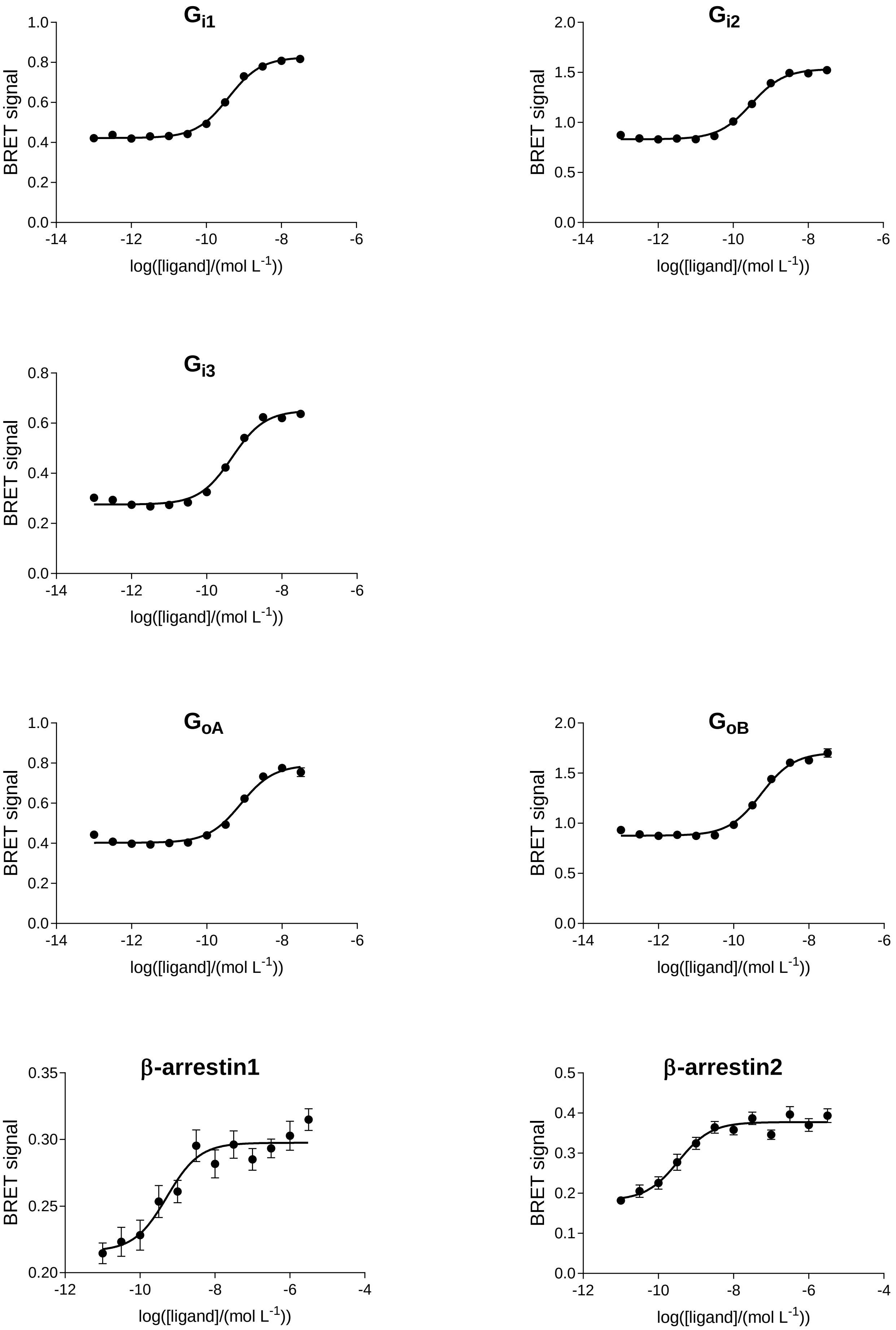

### CB2: JWH133

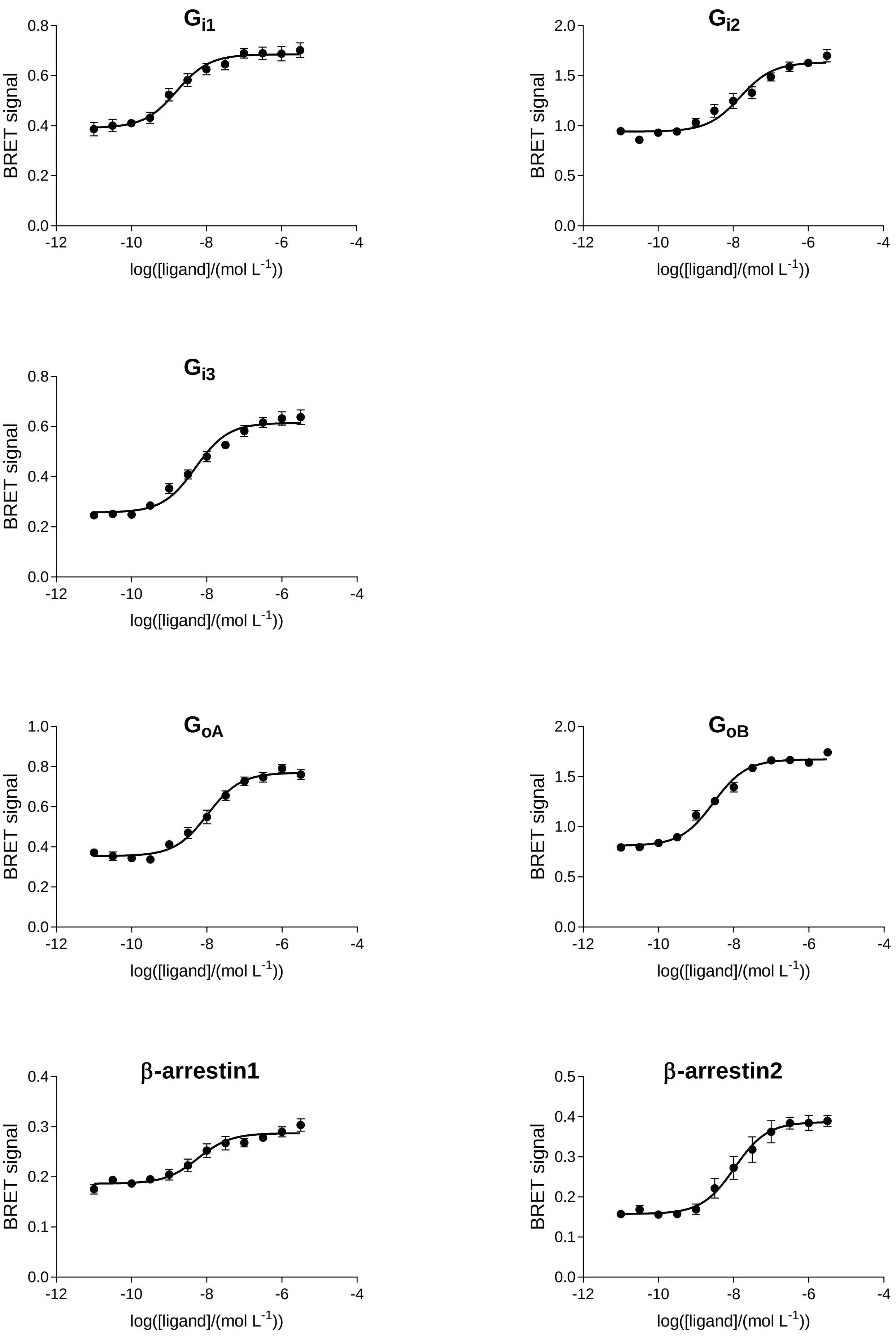

Gi1
Gi2
Gi3
GoA
GoB
β-arrestin1
β-arrestin2
BRET signal
log([ligand]/(mol L-1))
0.0
0.2
0.4
0.6
0.8
1.0
0.5
1.5
2.0
0.1
0.3
-12
-10
-8
-6
-4

# CB2: CP55940

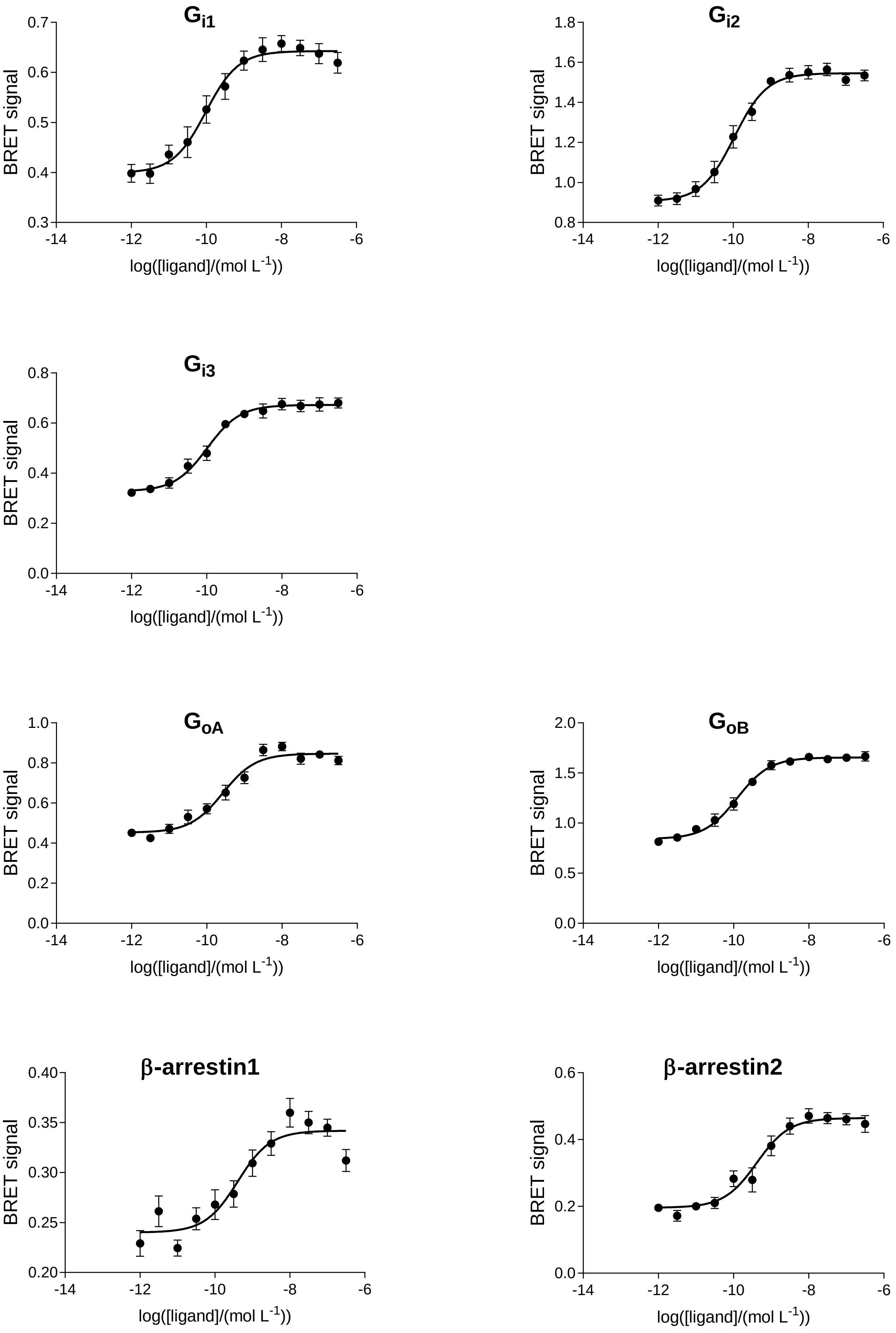
Gi1
Gi2
Gi3
GoA
GoB
β-arrestin1
β-arrestin2
BRET signal
log([ligand]/(mol L-1))
0.7
0.6
0.5
0.4
0.3
1.8
1.6
1.4
1.2
1.0
0.8
0.6
0.4
0.2
0.0
2.0
1.5
0.5
0.40
0.35
0.30
0.25
0.20
-14
-12
-10
-8
-6

# CB2: 2-AG

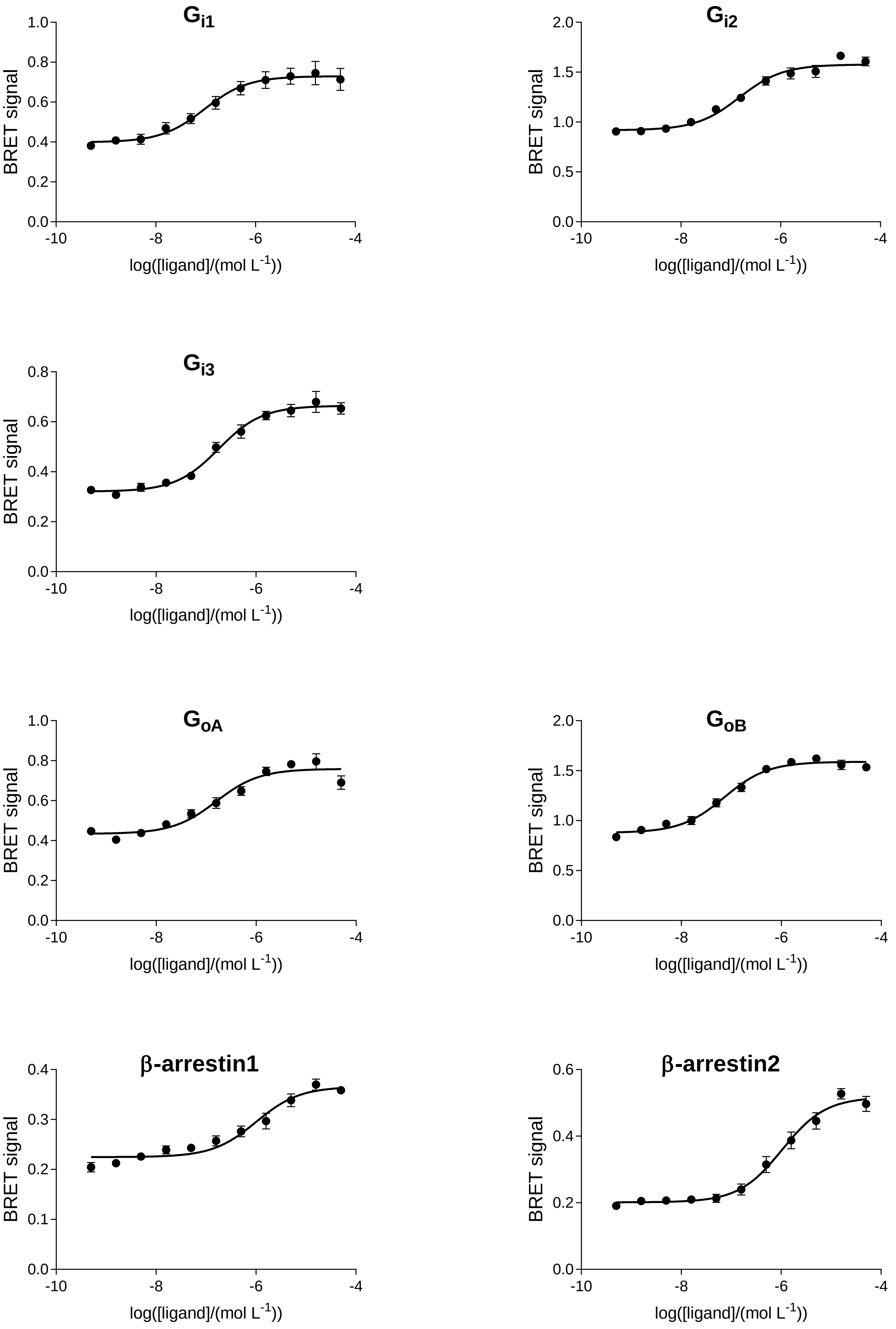

Gi1
Gi2
Gi3
GoA
GoB
β-arrestin1
β-arrestin2
BRET signal
log([ligand]/(mol L-1))

# CB2: RO6435559

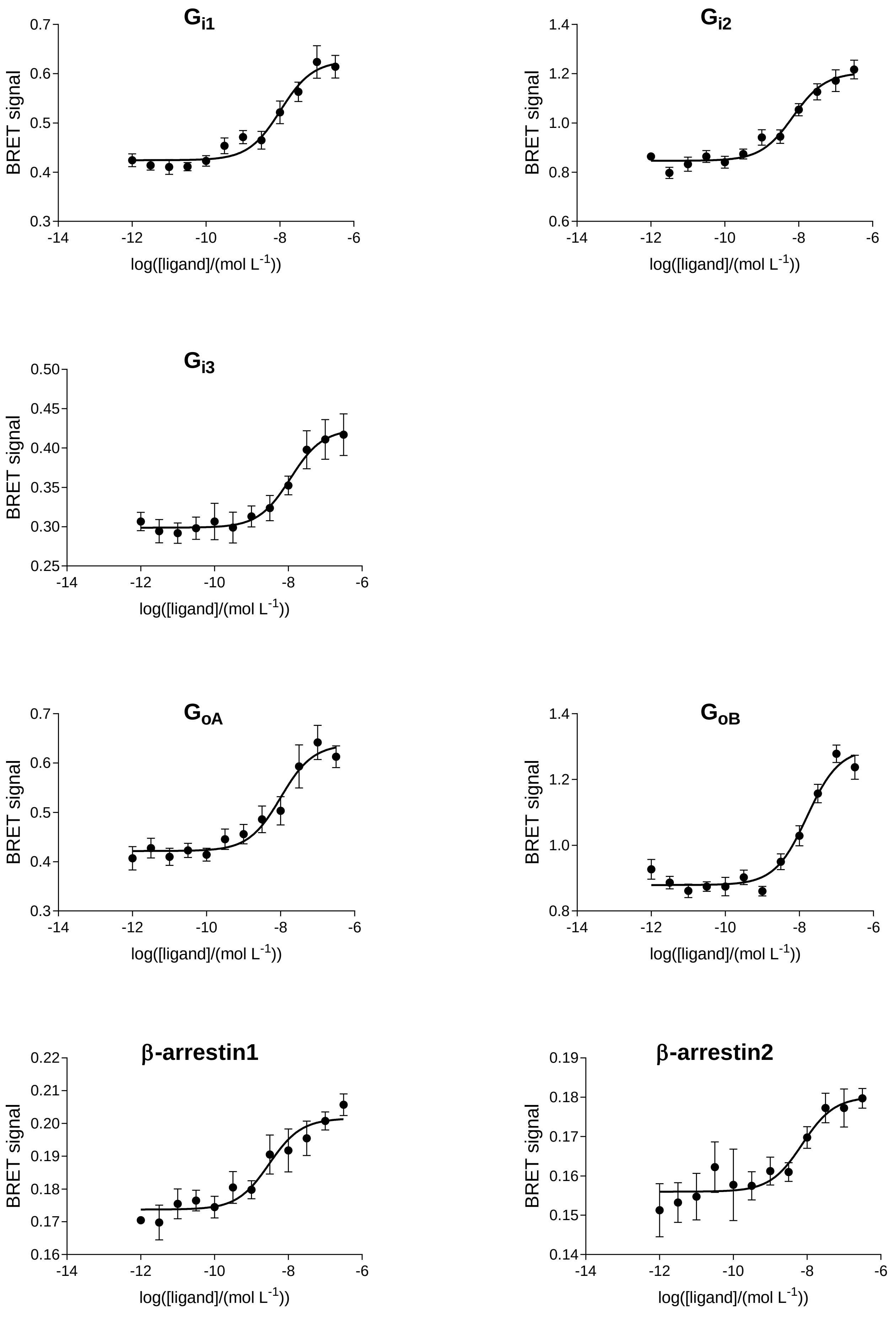

# CB2: RO6843766

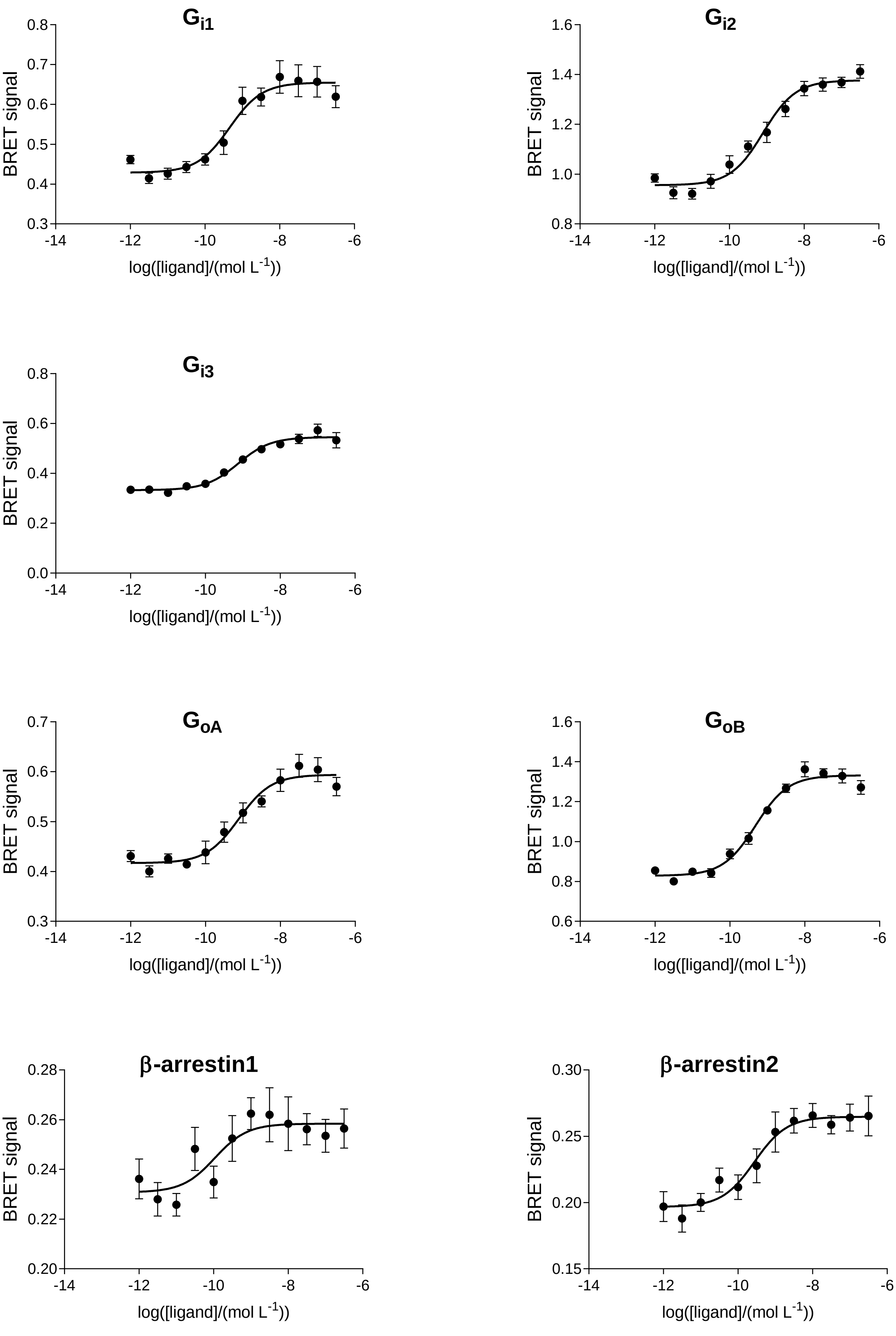

# CB2: RO6844112

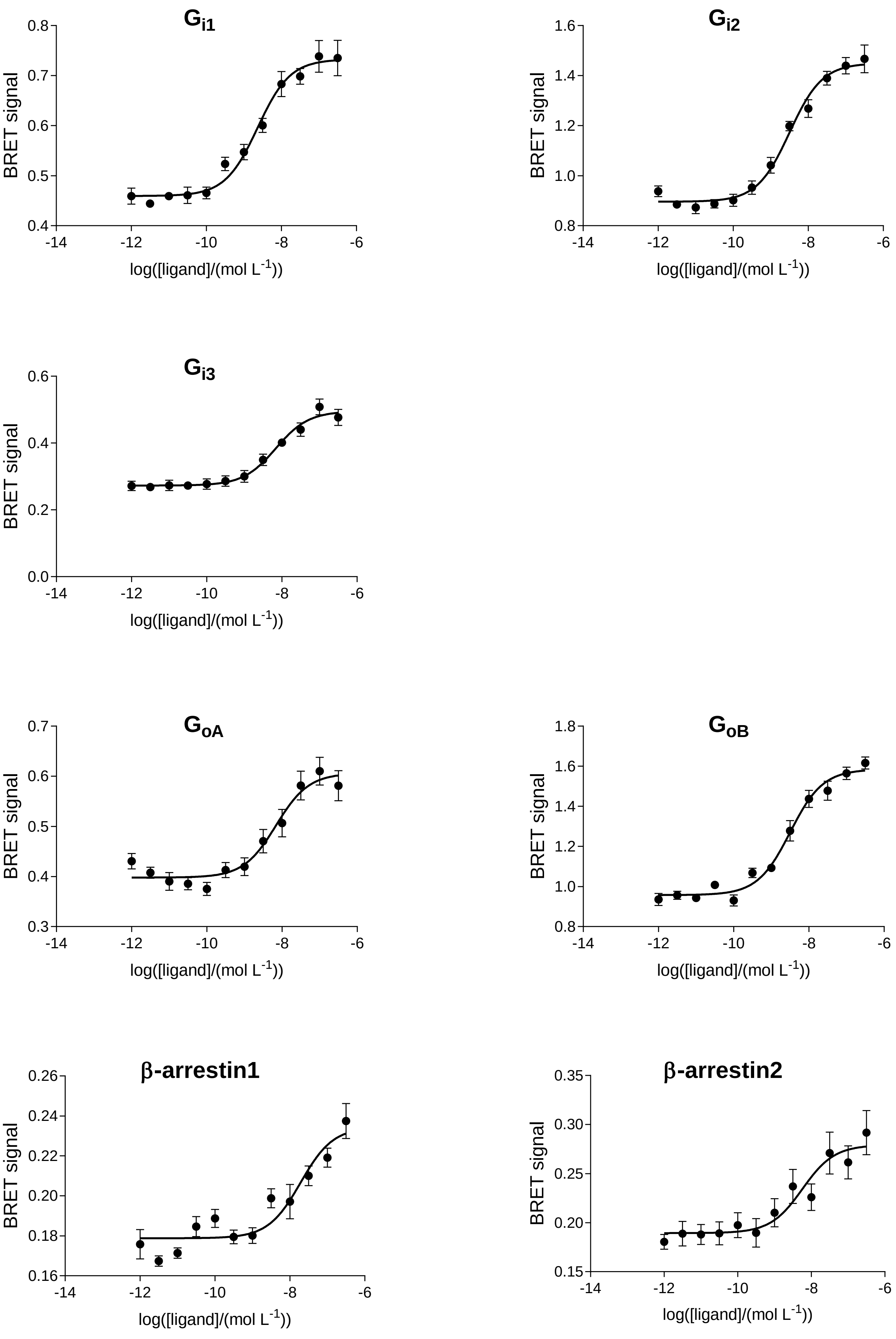

# CB2: RO6844395

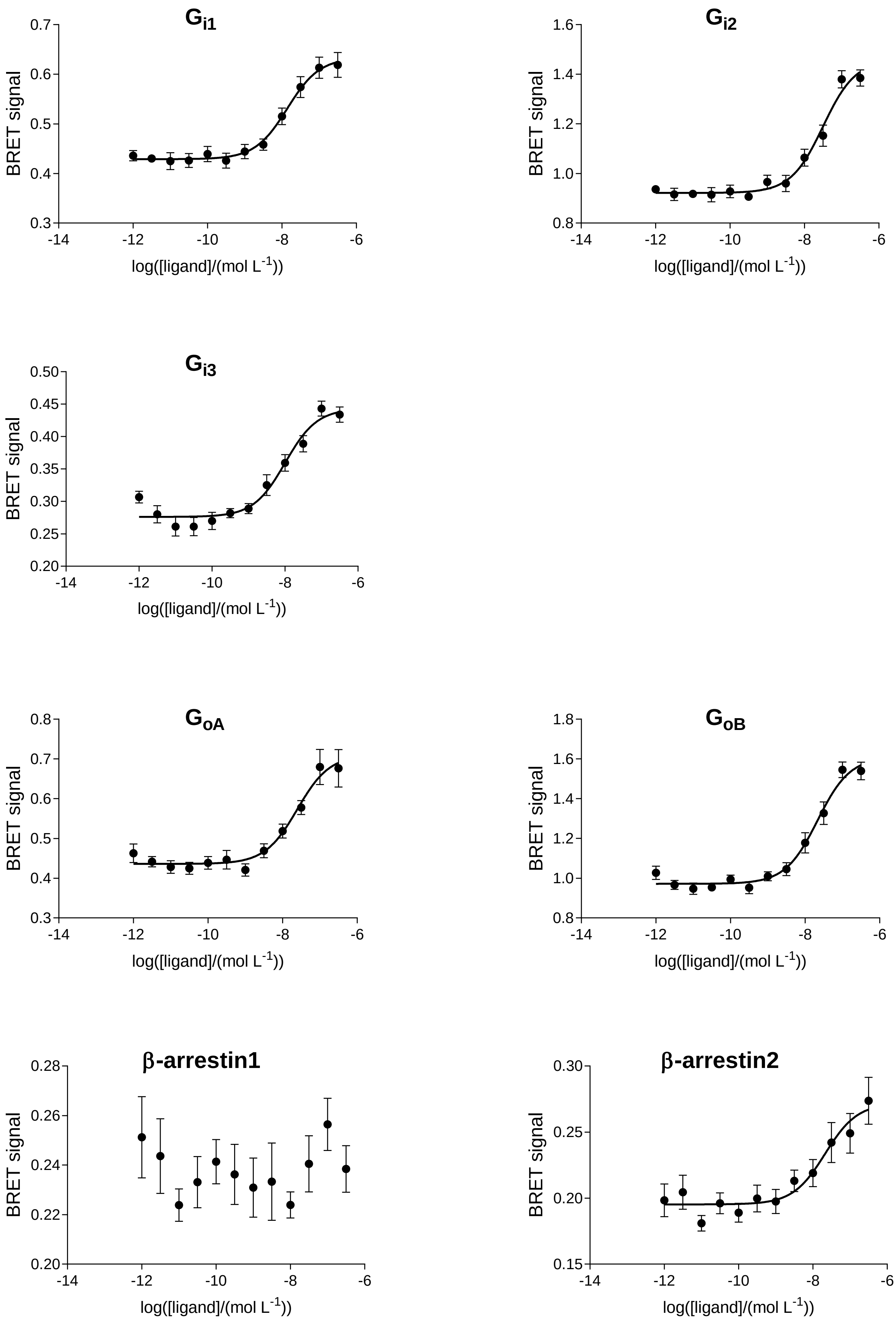

# CB2: RO6850007

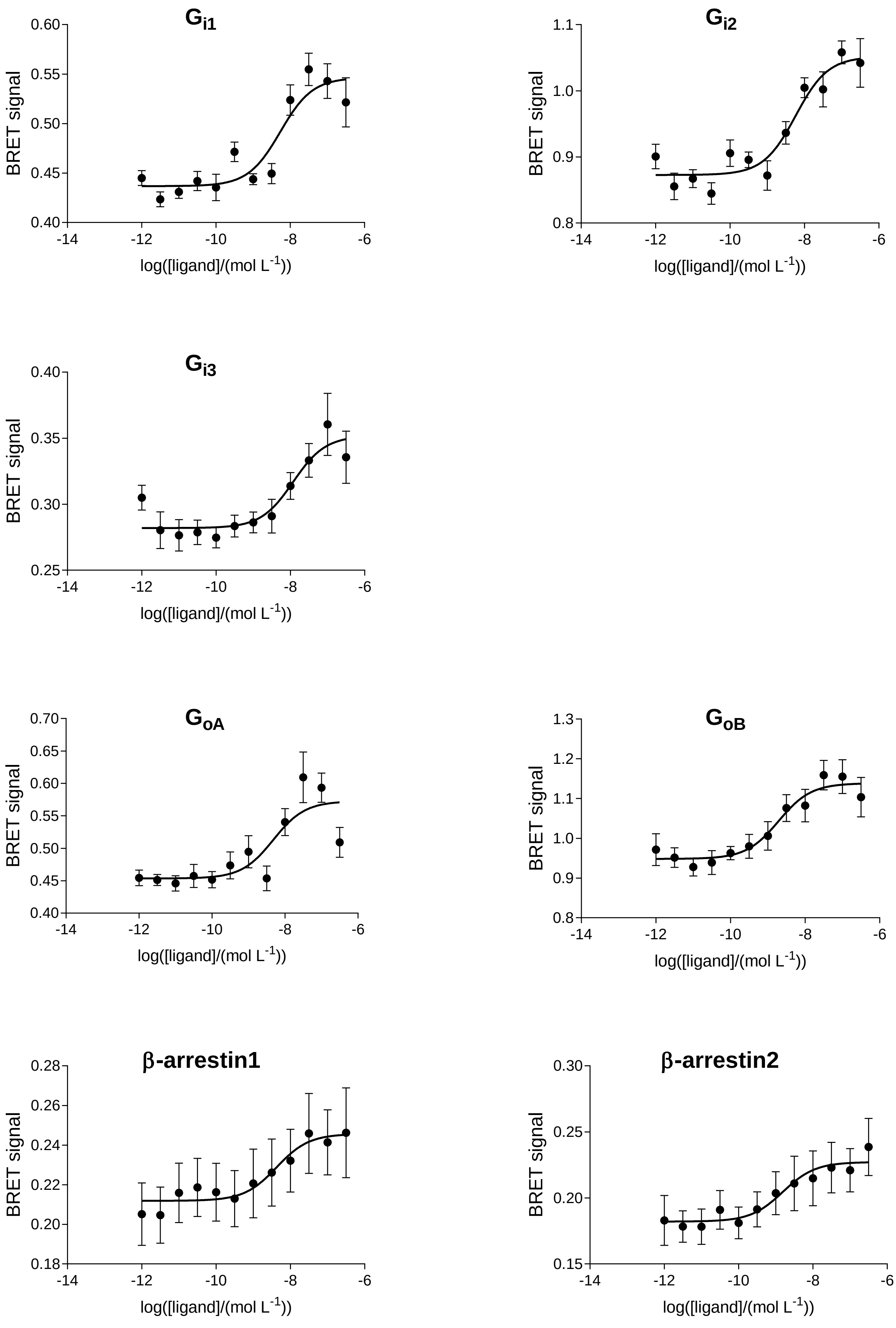

# CB2: RO6853457

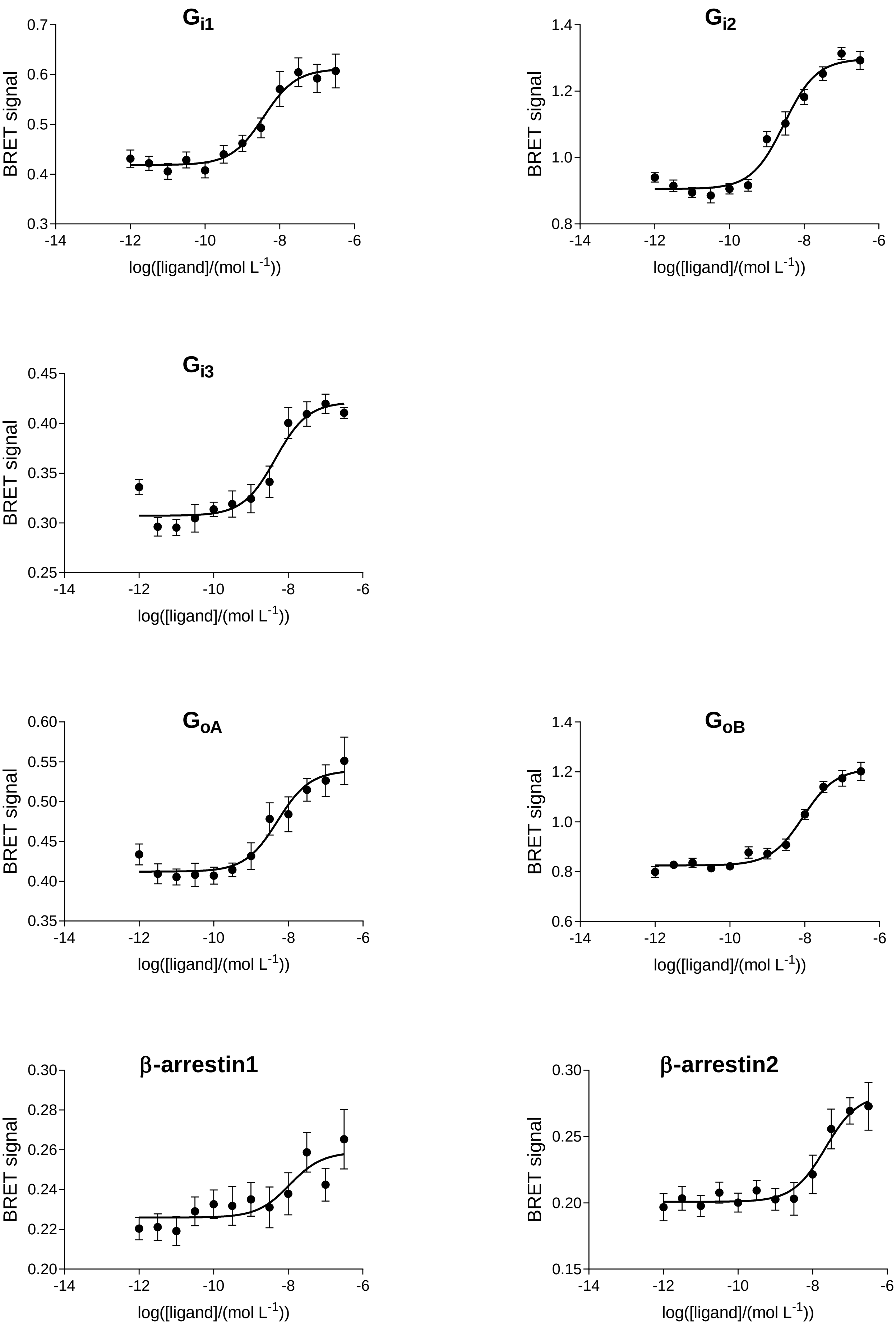

# CB2: RO6853973

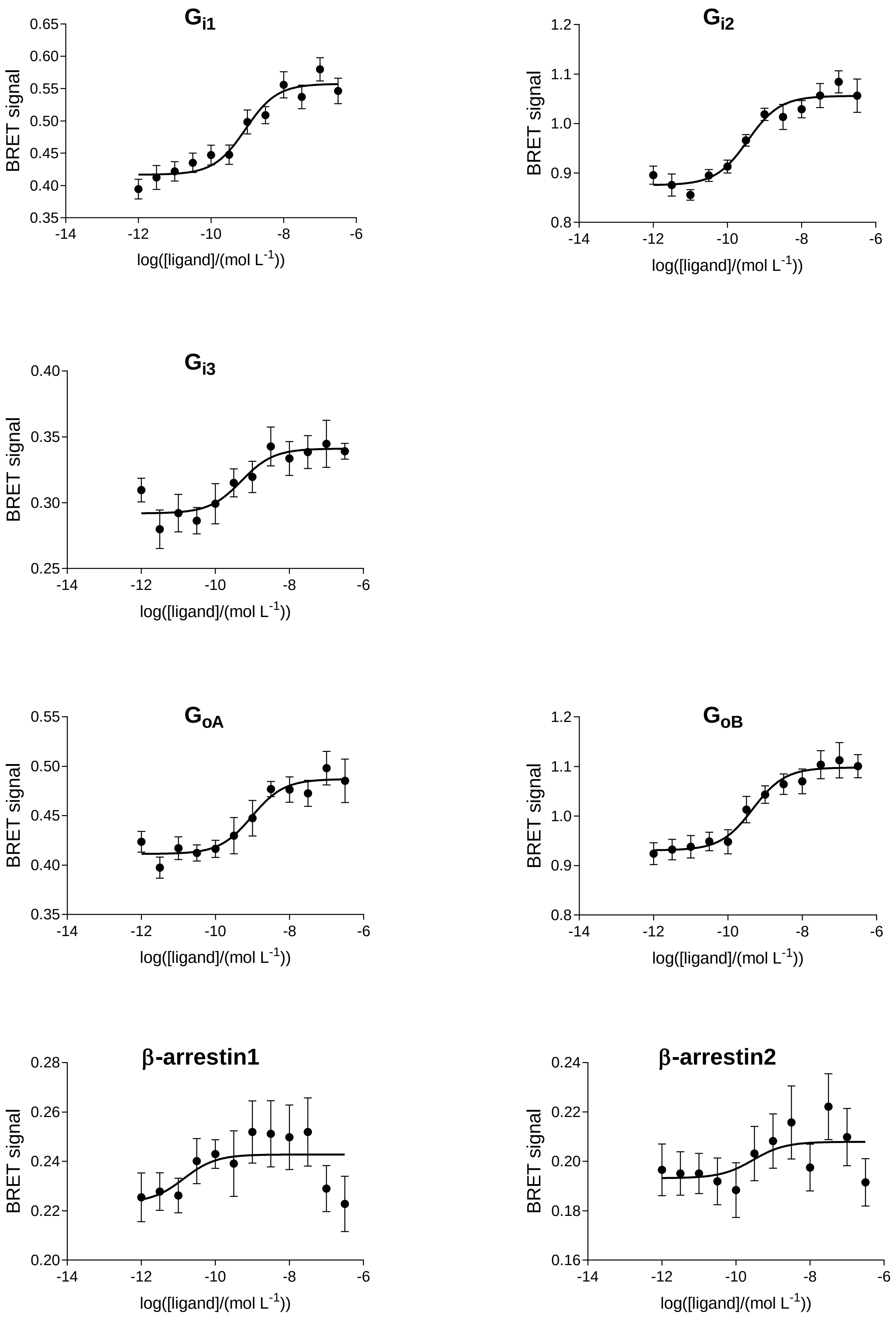

# CB2: RO6869094

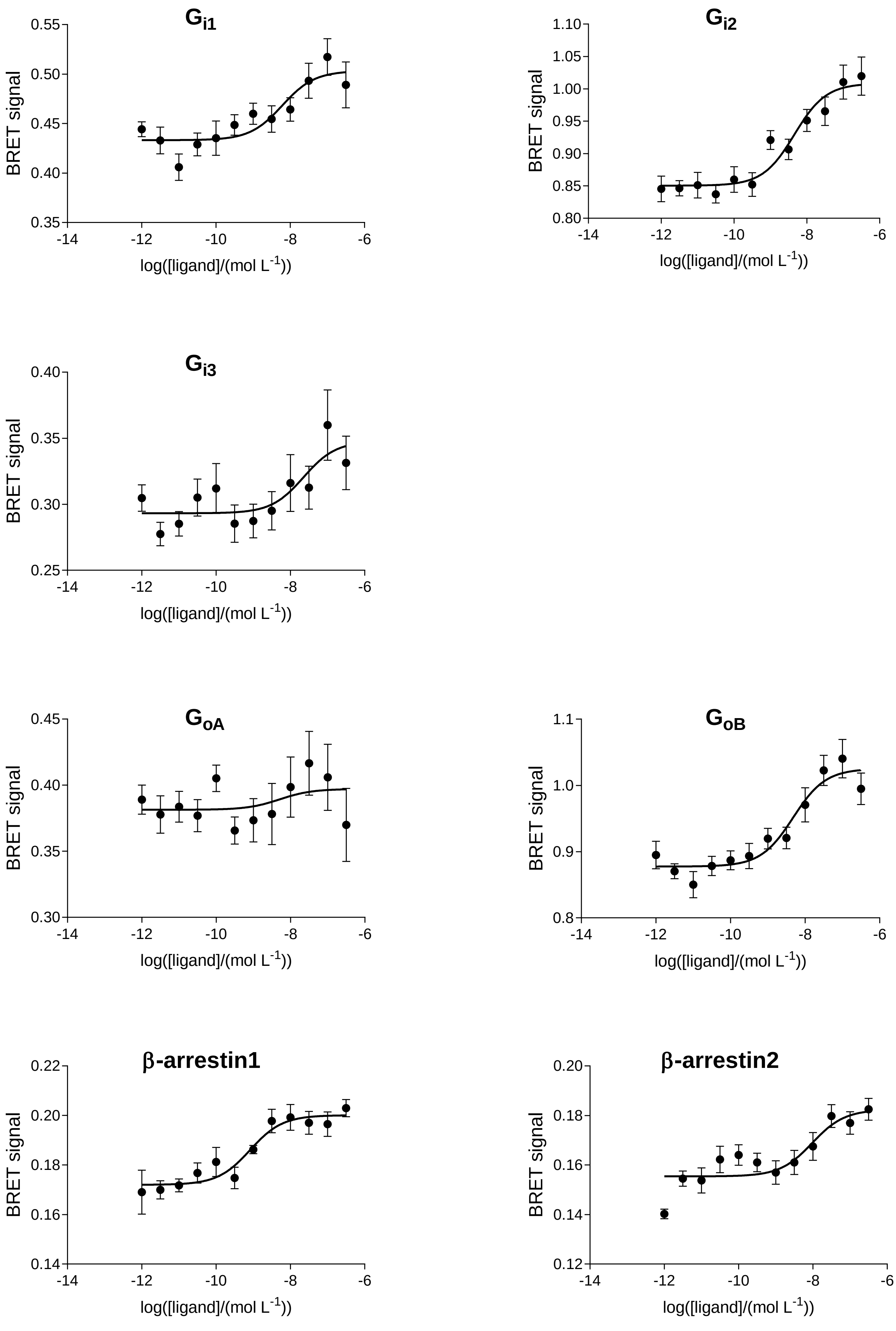

# CB2: RO6871487

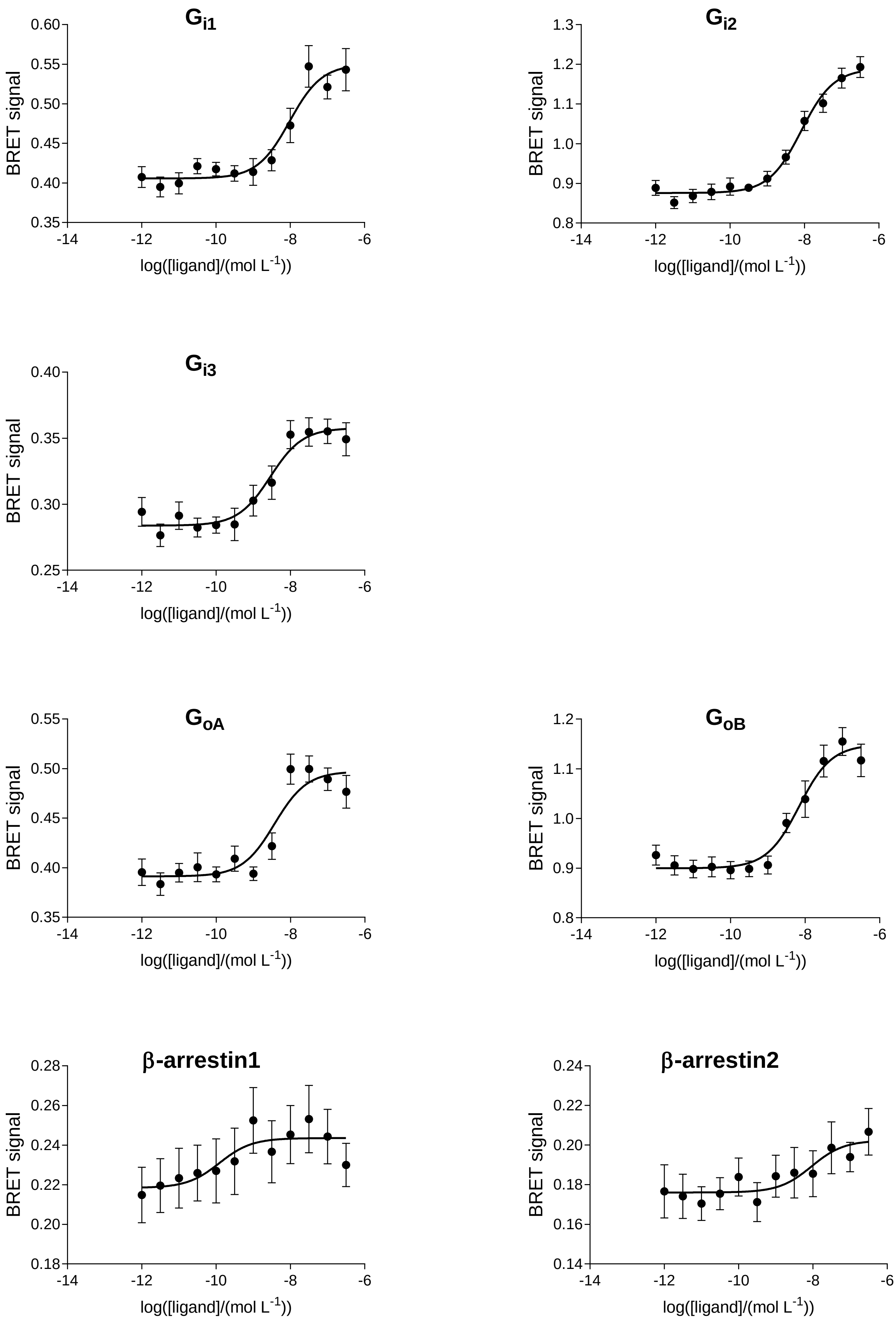

# CB2: RO6878558

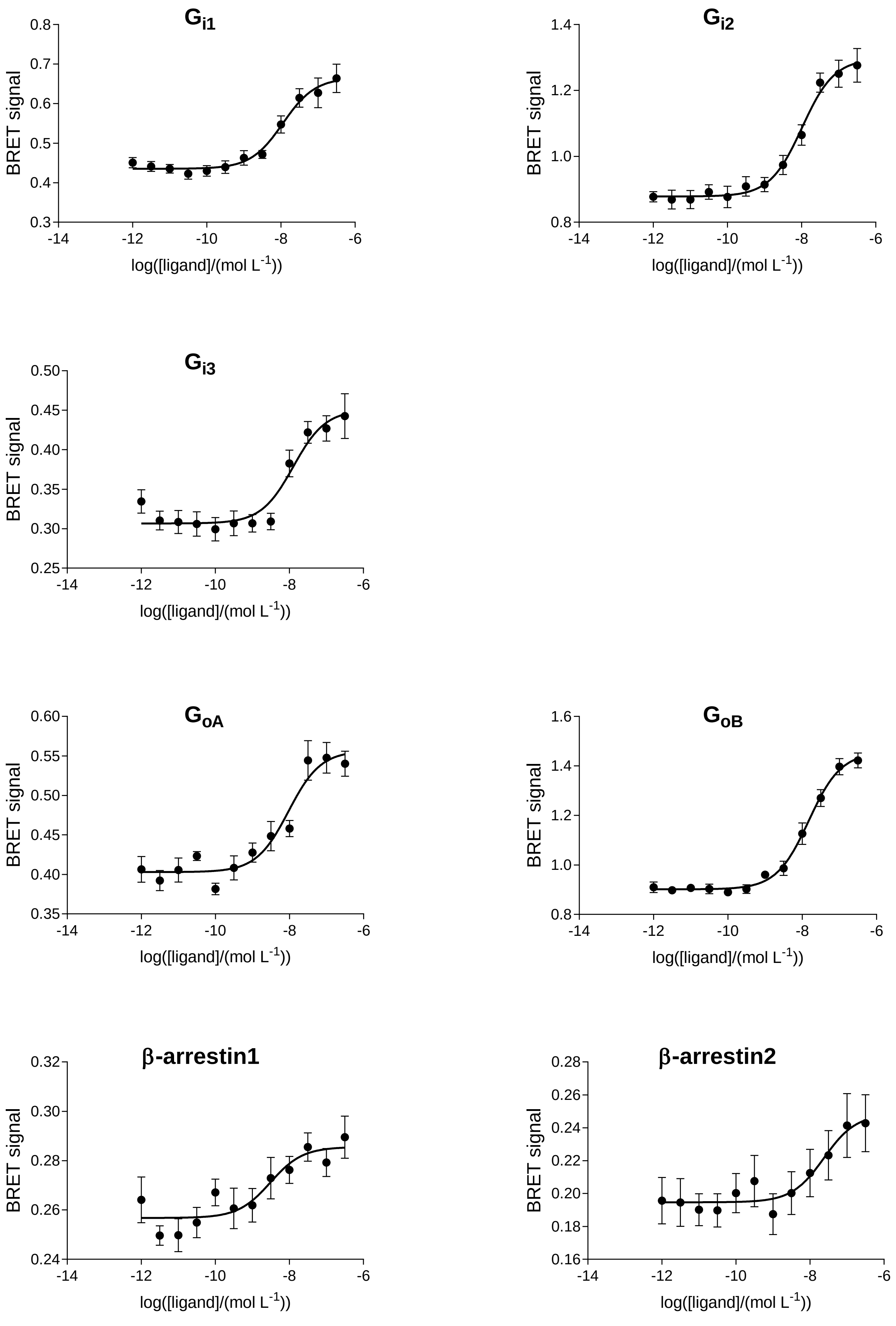

# CB2: RO5135445

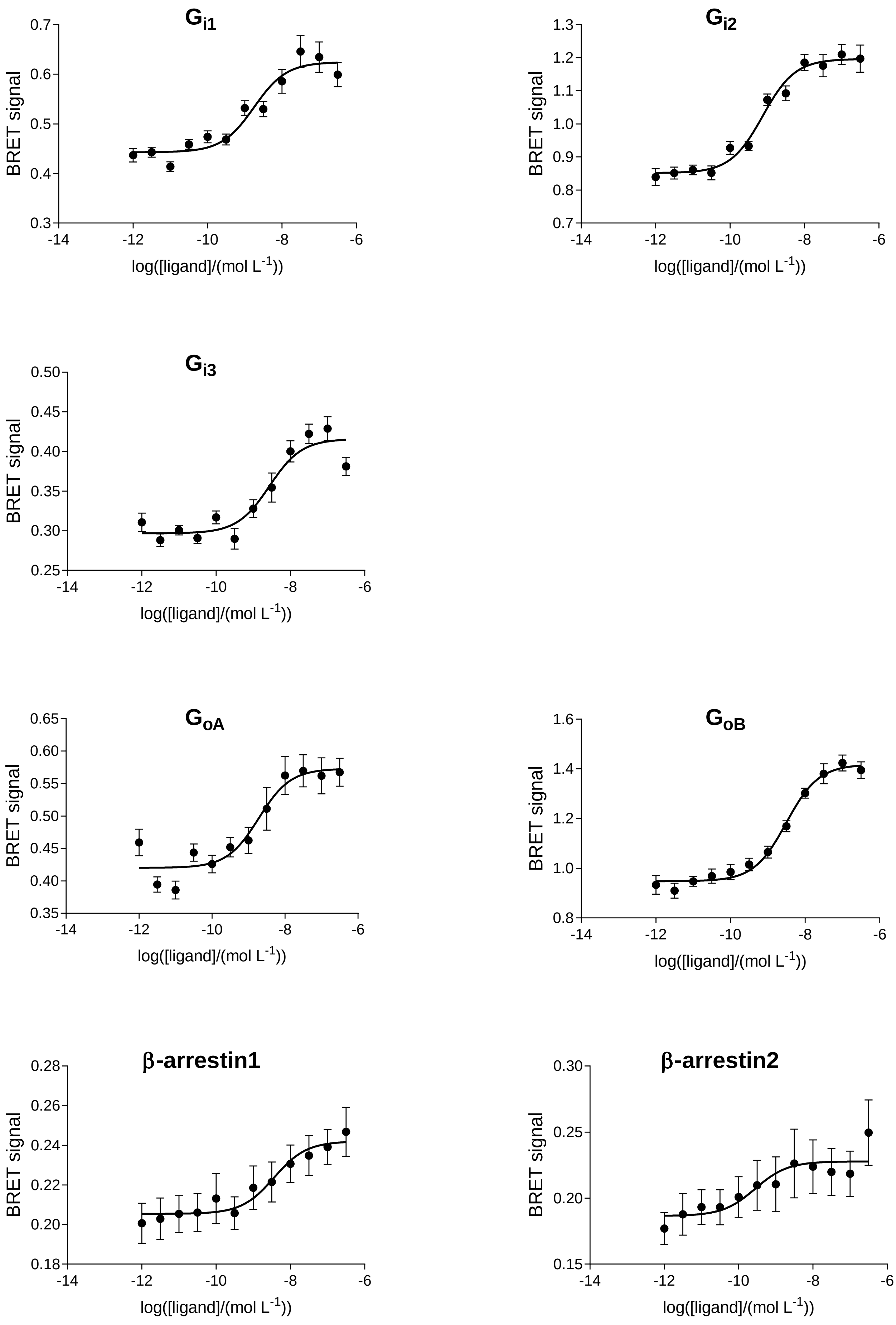

# CB2: RO6883666

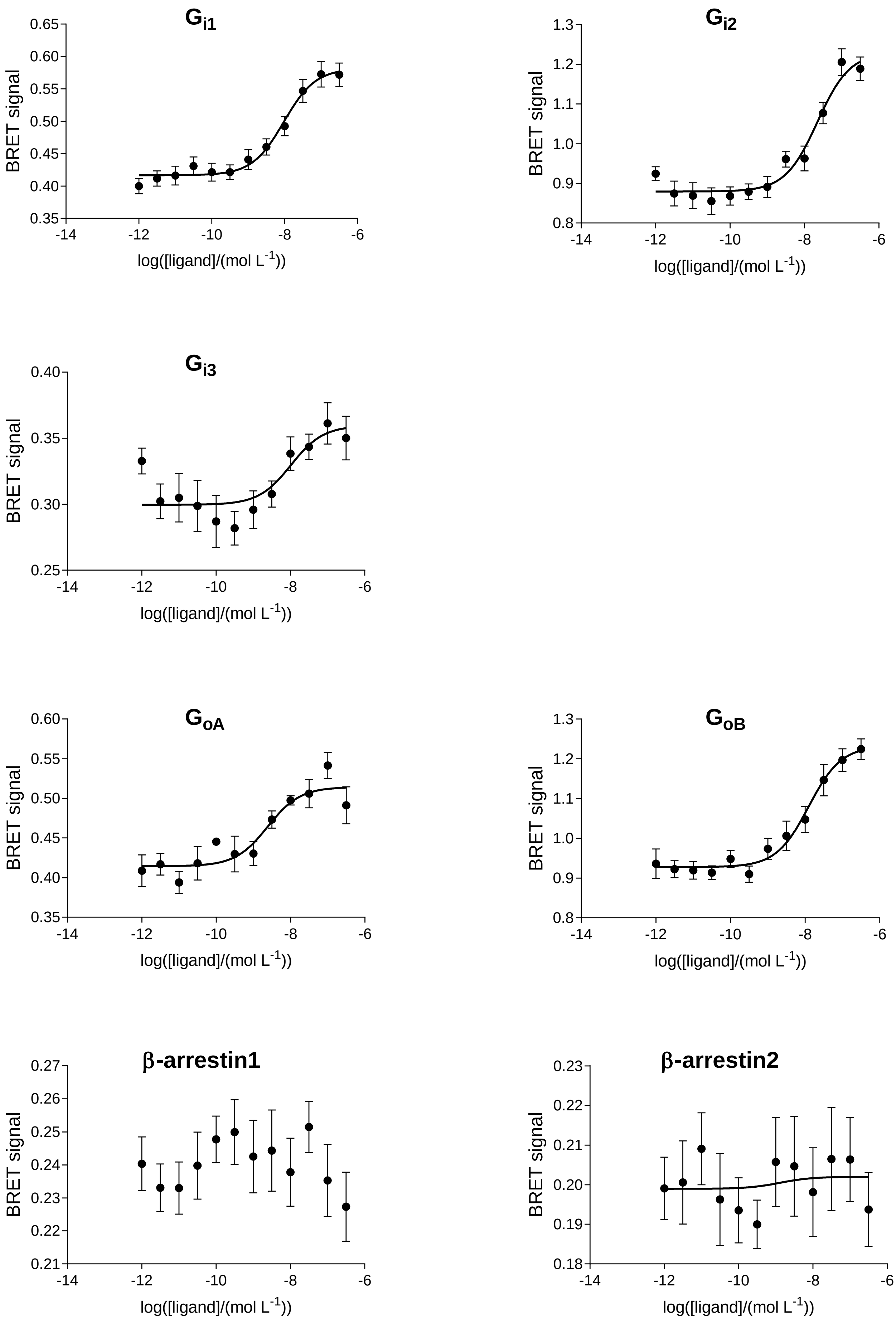

# CB2: RO6892033

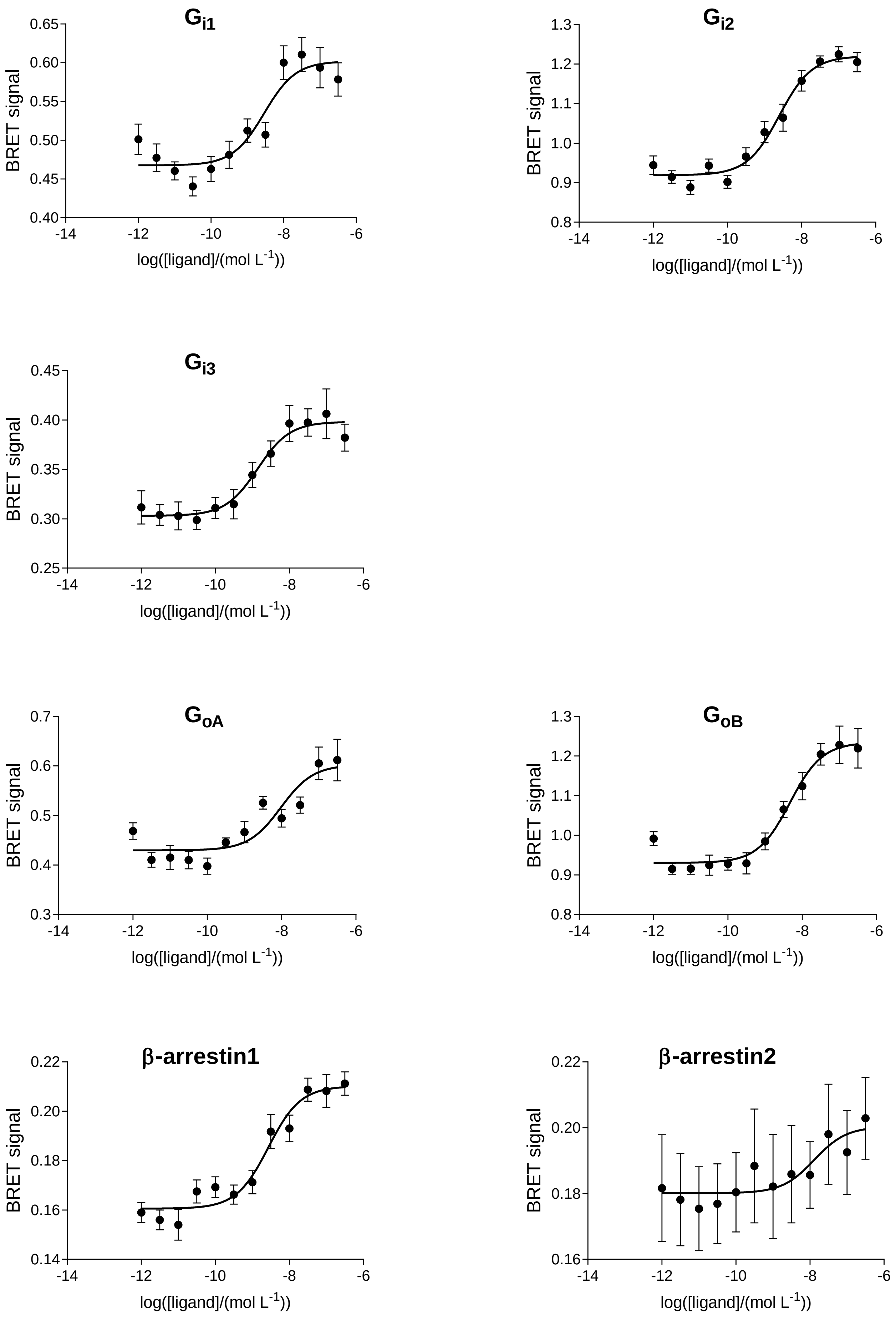

# CB2: RO6926274

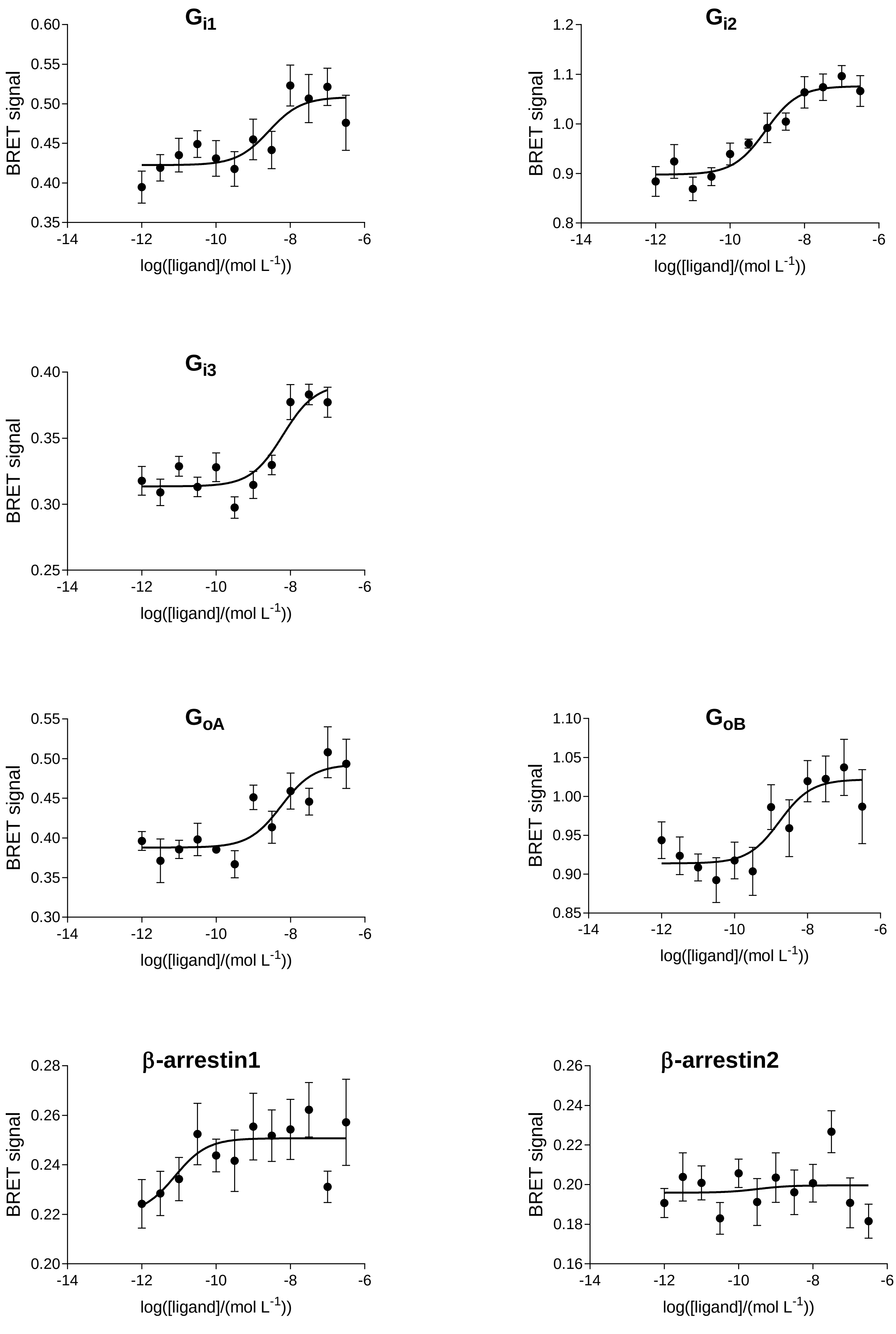

# CB2: RO7032019

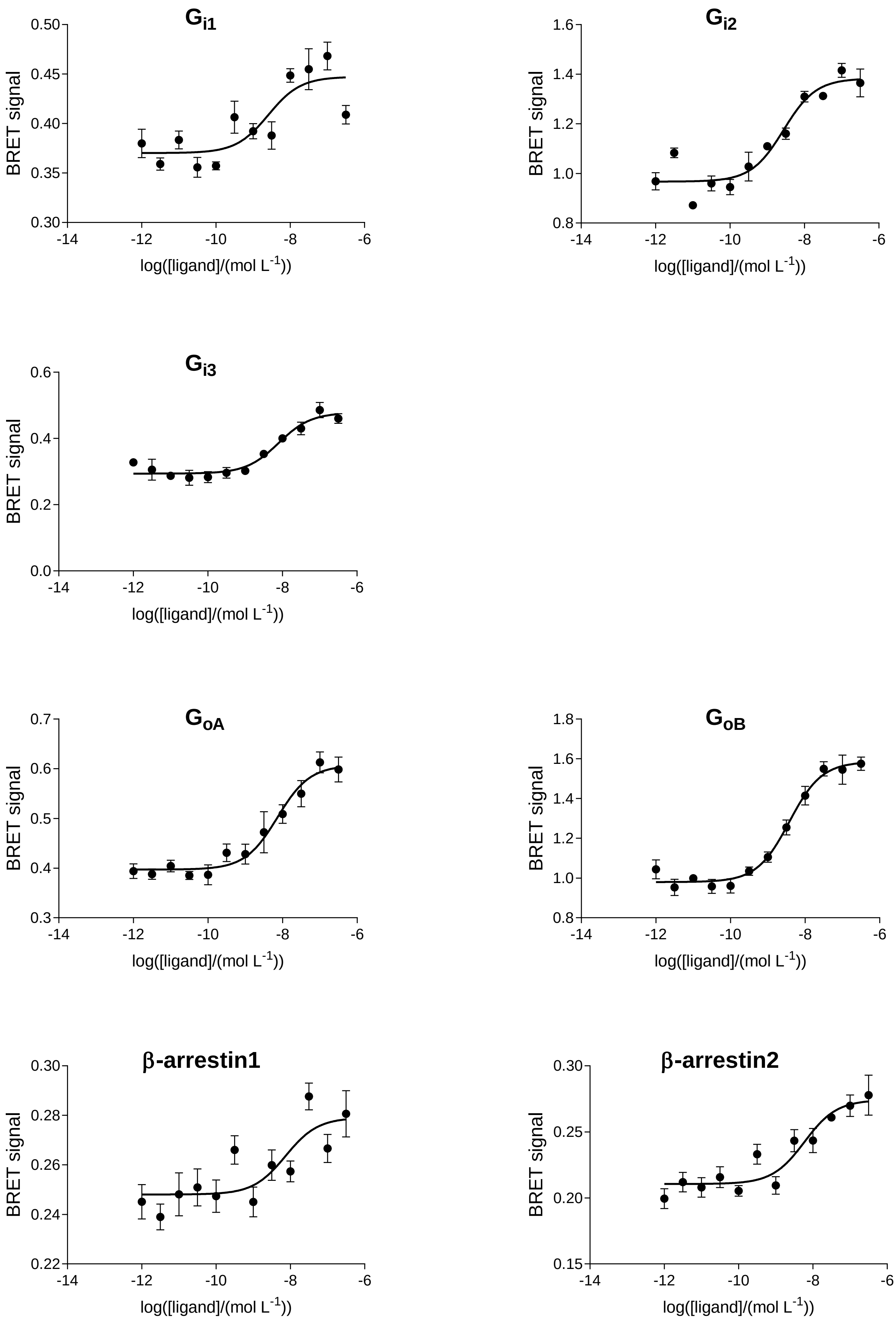

### CB2: FMP7234690

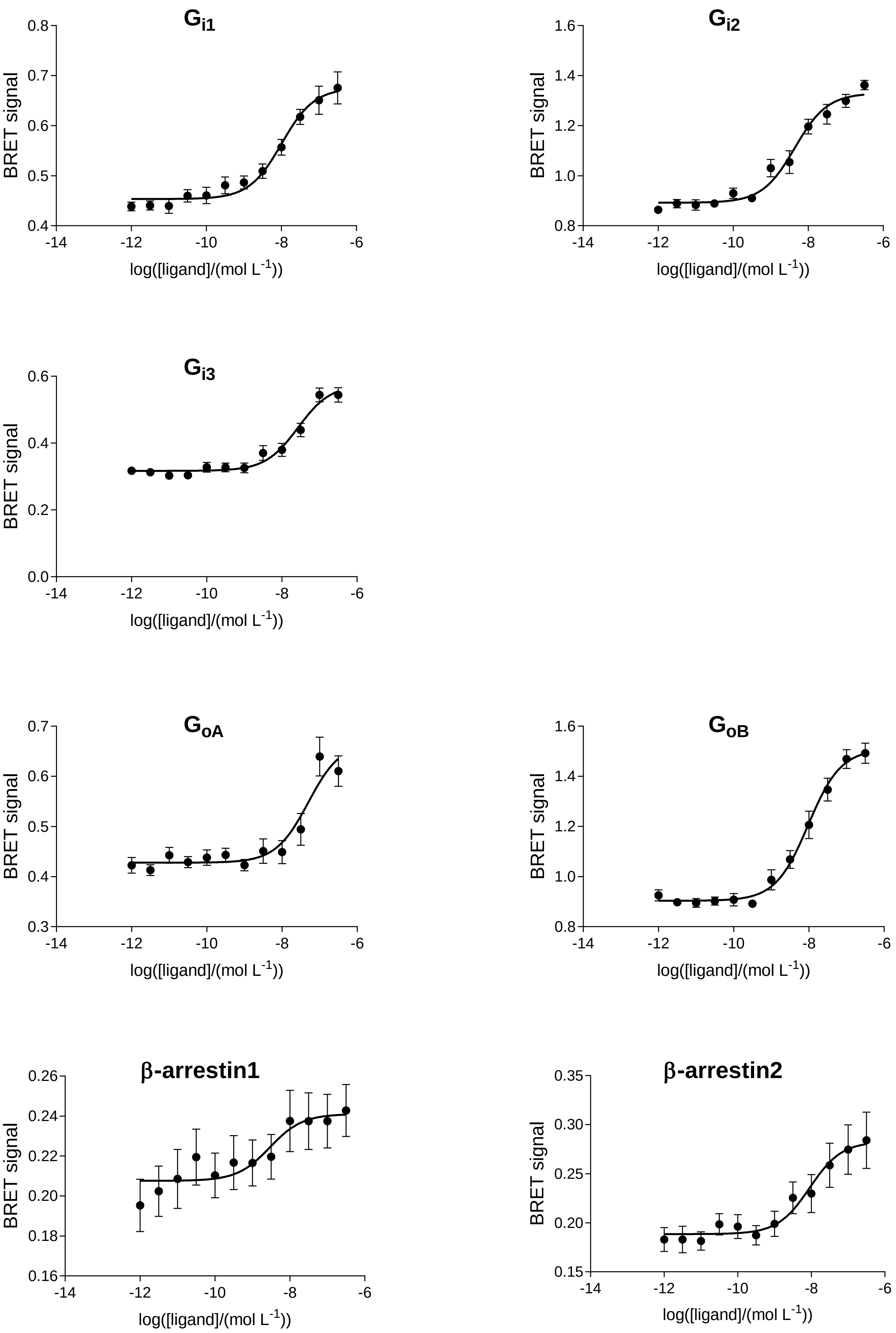

### CB2: FMP7234691

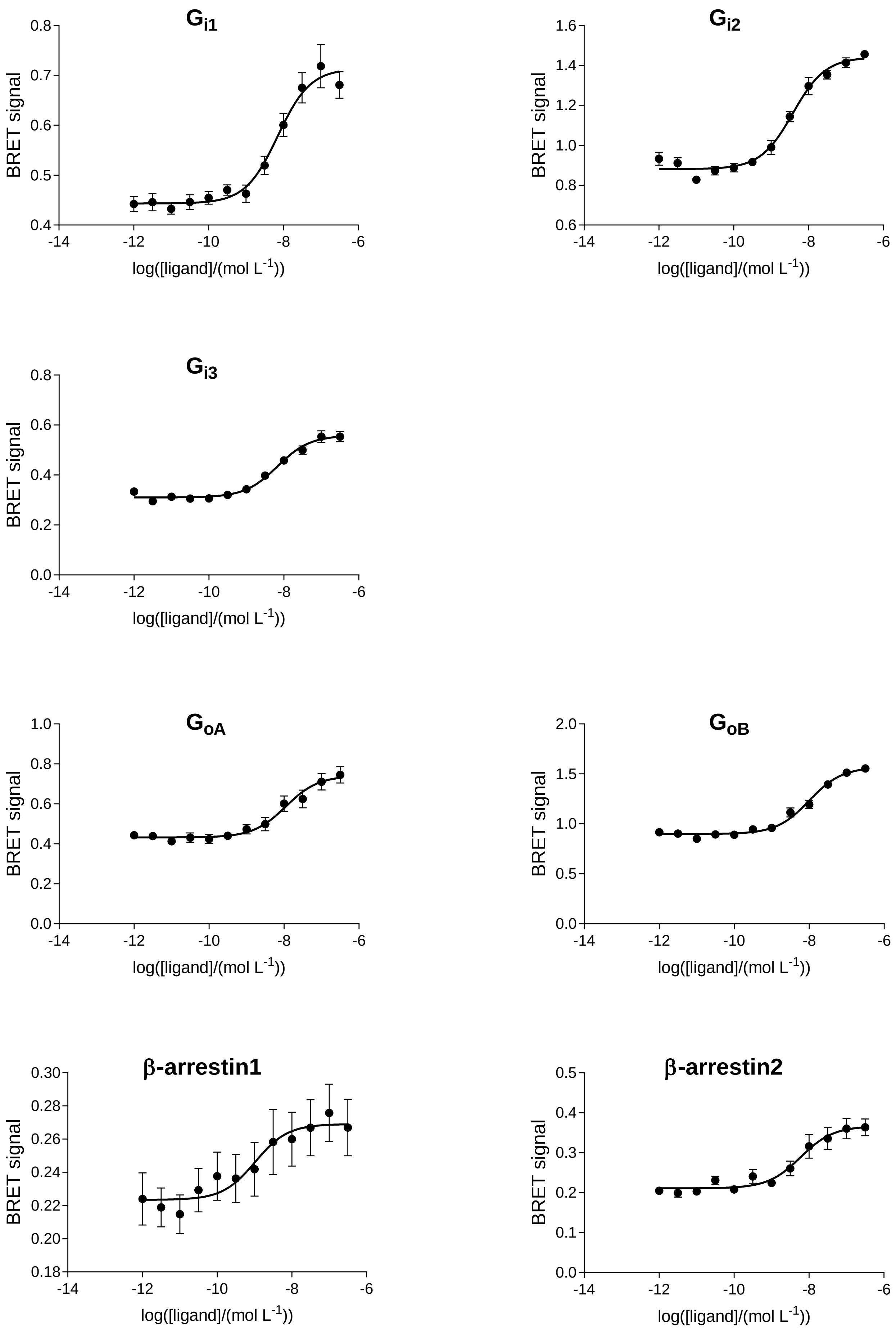

### CB2: FMP7234694

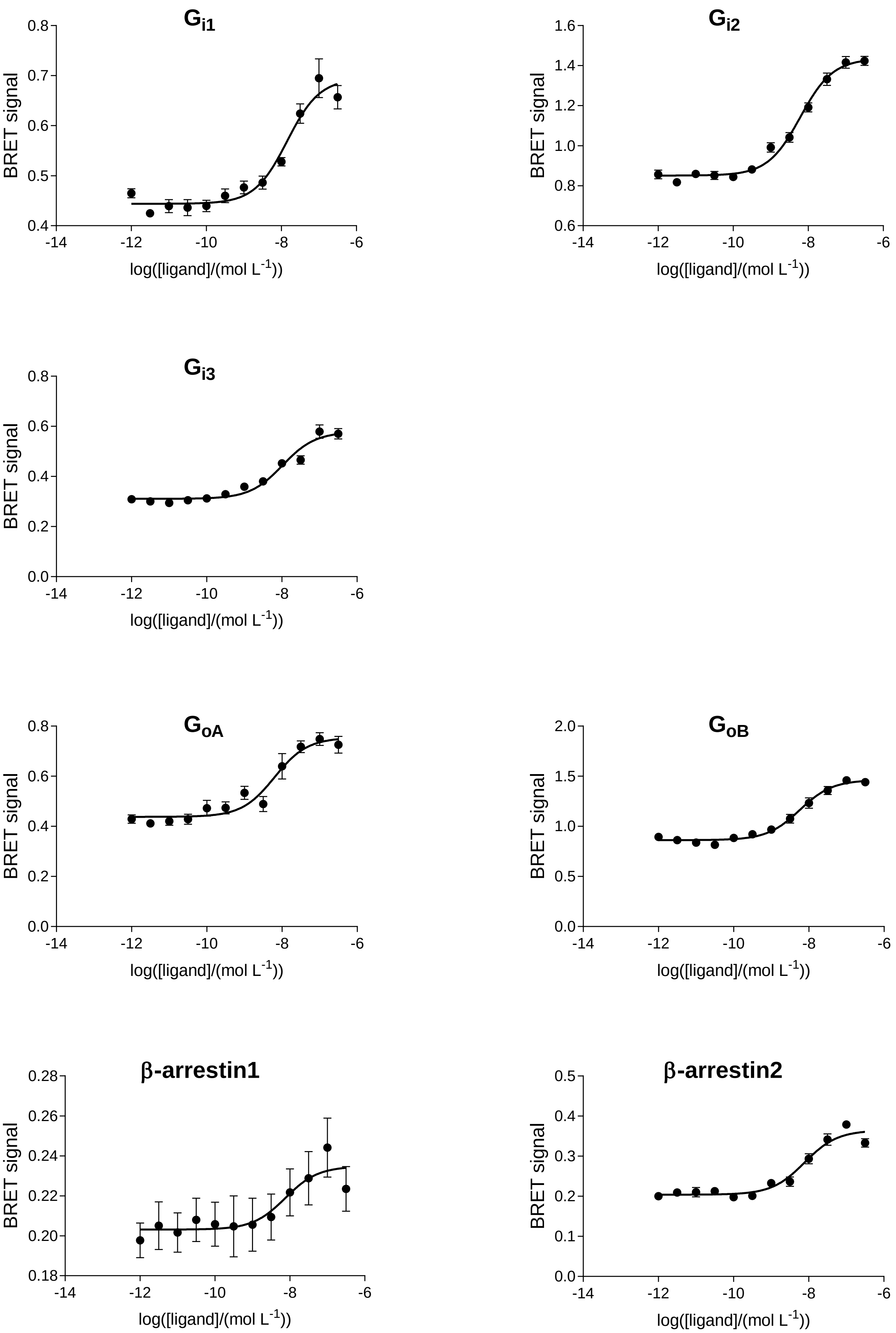

### CB2: FMP7234698

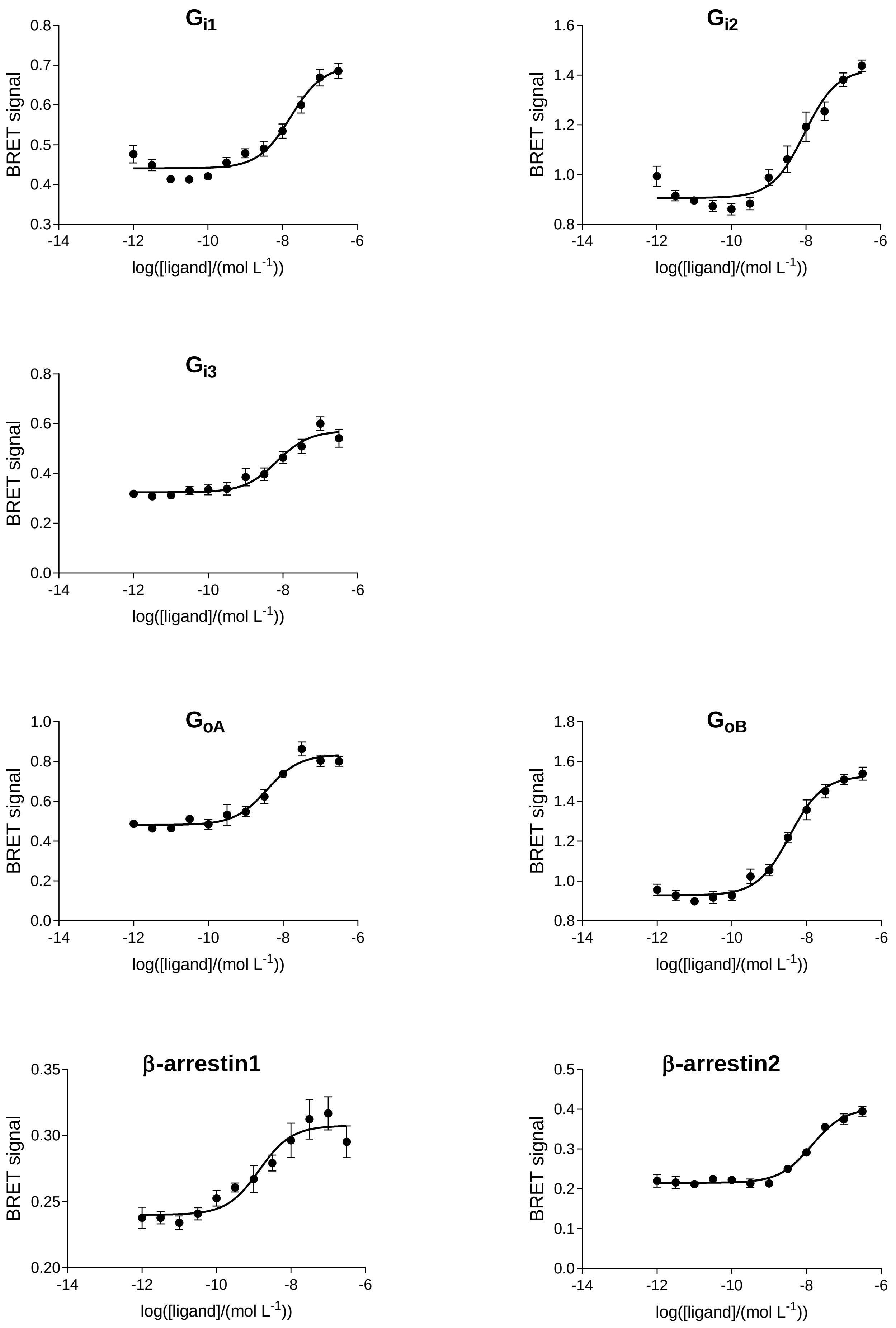

### CB2: FMP7234699

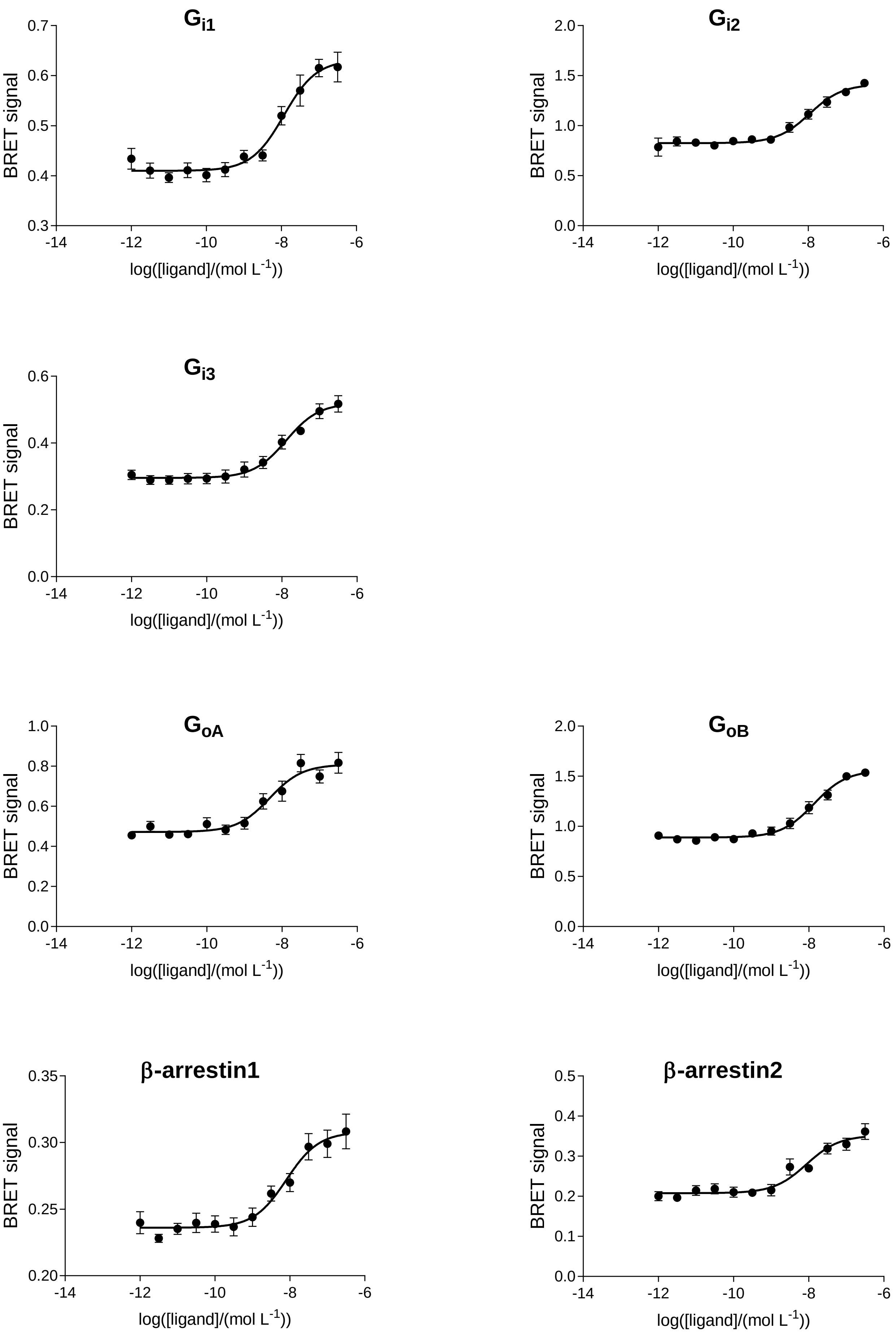
